# Biocontainment attenuation of mobile DNA host range in a wastewater microbiome

**DOI:** 10.64898/2026.07.13.738295

**Authors:** Malyn A. Selinidis, Travis R. Seamons, Lauren B. Stadler, Jonathan J. Silberg, James Chappell

## Abstract

Biocontainment systems designed to attenuate the spread of mobile DNA are challenging to evaluate within microbiomes of engineered environments. To better understand how toxin-based biocontainment systems affect horizontal gene transfer (HGT) in a microbiome, we evaluated the host range of pairs of plasmids using orthogonal catalytic RNA (cat-RNA) that amend distinct barcodes to 16S rRNA following HGT. We show that mobilizable (5 kb) and self-mobilizable (60 kb) plasmids, which use the same RP4 transfer machinery but different origins of replication, overlap in their host range when conjugated in parallel into a wastewater community, with 127 of the 143 amplicon sequence variants (ASVs) presenting barcoding signals from both plasmids (89%). We also find that mobilizable plasmids with or without the *Escherichia coli* CcdB toxin overlap in host range in a wastewater community. Among the two most abundant orders, CcdB attenuated the barcoding signal in Aeromonadales more consistently than Enterobacteriales, which have F plasmids containing the CcdB-CcdA toxin-antitoxin system used for biocontainment. Also, CcdB decreased the abundance of the mobilizable plasmid by >100-fold and yielded mutations in 85% of the reads. Together, these findings reveal how pairs of plasmids expressing orthogonal cat-RNA can be used to monitor the effects of plasmid-encoded traits on mobile DNA persistence following HGT. They also highlight challenges when using biocontainment systems containing genes related to those found in the microbiomes targeted for engineering.

## INTRODUCTION

Engineered microbes are being developed for a wide range of applications, including agriculture [1, 2], animal husbandry [3, 4], bioremediation [5, 6], biomanufacturing [7, 8], and health [9, 10]. These biotechnologies have tremendous promise to overcome challenges related to feeding a growing population [11], achieving a circular economy [12], and creating therapeutics that overcome antimicrobial resistance and treat disease [13]. To date, only a small number of these technologies have successfully transitioned to real-world ecological settings in open environments, such as biofertilizers and probiotic health supplements [14, 15], although numerous environmental release studies have been performed to assess commercialization potential of other products [16]. One challenge with evaluating these engineered technologies in the context of open environments has been the lack of simple tools for monitoring the fate, transport, and risk of engineered DNA following introduction into the environment [17]. Only a small number of studies have evaluated microbial persistence within open environments [16]. While these studies have shown that engineered microbe persistence can vary [18–20], most have not assessed the fate of the synthetic DNA used for microbial engineering, which could persist beyond the engineered microbes when exchanged using horizontal gene transfer (HGT).

To enable the safe use of engineered microbes across different operating environments, a range of genetically-encoded biocontainment strategies have been developed [21–23]. Many of these approaches are designed to control the duration and locations where engineered microbes persist, such as engineered metabolic dependencies [24, 25], synthetic kill switches [26–28], and genetically-recoded genomes [29–31]. Other biocontainment approaches are designed to inhibit HGT between engineered and environmental microbes [32–35]. Persistence studies have revealed that these latter systems vary in their performance. In one study, engineered DNA was observed in an environmental microbe years after the introduction of an engineered microbe [18]. Other studies have been unable to detect engineered DNA after only one year [19]. As the ecological controls on HGT in many environments remain poorly defined [17], there is a need to go beyond standard measurements of engineered DNA abundance in environmental samples and to directly measure how well biocontainment strategies restrict HGT across each strain in the microbiomes targeted.

HGT is commonly monitored in microbiomes using culture-dependent methods, such as visual reporters or selectable markers [36, 37]. However, these methods are limited in their ability to detect HGT in communities that contain unculturable microbes [38, 39]. Culture-independent methods have recently emerged to overcome these limitations. These approaches covalently link mobile DNA and host nucleic acids [40–42], thereby allowing for the detection of mobile DNA host range using targeted sequencing. With some of these technologies, HGT-dependent barcoding can be accomplished using genetically-encoded technologies [43]. Herein, we leverage one technology, called RNA addressable modification (RAM), to study DNA-encoded controls on conjugation in a wastewater microbiome [44–46]. With RAM, a catalytic RNA (cat-RNA) is used to report on HGT host range in the community by barcoding 16S ribosomal RNA (rRNA) in recipient cells that take up mobile DNA. Herein, we use orthogonal cat-RNA to compare the conjugative host ranges of pairs of plasmids in wastewater communities, including mobilizable and self-mobilizable plasmids that contain different origins of replication and pairs of mobilizable plasmids that contain or lack a genetic biocontainment circuit that is designed to express the DNA gyrase inhibitor CcdB following HGT. We observe a large overlap in the host ranges across each pair of plasmids, although the relative RAM signals from each plasmid pair varied across ASVs, illustrating how this culture-independent method can reveal how variations in mobile DNA host range in specific taxonomic groups.

## MATERIALS AND METHODS

### Plasmids and strains

All plasmids and strains are described in Tables S1-S2. Genome integration was performed at the *lacZ* operon [47].

### Growth medium

Lysogeny broth (LB) was used for all experiments. To complement the growth defect of *E. coli* strain MFDpir [48], LB and LB agar were supplemented with diaminopimelic acid (DAP) (0.3 or 0.6 mM). Note that for selections, DAP was not added to prevent donor growth. Individual strain cultures were supplemented with appropriate antibiotics and washed to remove antibiotics prior to conjugation.

### Wastewater collection

Wastewater was acquired from the West University Wastewater Treatment Plant in Houston, Texas, transported on ice for <1 hour, and used immediately for conjugation following pelleting (1 mL) and resuspension in phosphate-buffered saline (PBS) (1 mL).

### Filter mating

To conjugate from *E. coli* to *P. putida*, individual cultures were washed in LB, normalized to OD_600_ of 0.5 and mixed at a 7:3 donor:recipient ratio, plated on LB agar (60 µL), and incubated overnight at room temperature. To conjugate plasmids with and without CcdB from *E. coli* MFDpir to *E. coli* JEC088, an uracil auxotrophic mutant, strains were independently cultured, washed in LB, normalized to OD_600_ of 0.5, mixed 1:1, plated on LB-agar (50 µL), and incubated for 24 hours at room temperature. To conjugate plasmids into wastewater communities, donor strains were grown individually, washed, normalized to an OD_600_ of 0.5, and mixed with wastewater at a 1:1:2 ratio (donor-1:donor-2:wastewater). After mixing, samples were incubated on filter paper set on LB-agar plates for 24 hours prior to RNA extraction.

### RNA extraction

A Kingfisher Apex (Thermo Fisher Scientific) was used for extractions from conjugations involving pairs of microbes, while a Maxwell RSC 48 (Promega) was used for communities. With the Apex system, a MagMAX^TM^ Microbiome Ultra Nucleic Acid Isolation Kit (Thermo Fisher Scientific) was used. A modified protocol was implemented that involved a nucleic acid binding step, two washes, DNAse treatment, a second nucleic acid binding step, two additional washes, and elution into a volume of 50 µL/well. The DNAse treatment used a mixture of RNAse free water (170 µL/well), 20 µL/well DNAse buffer (Invitrogen AM2239), and 10 µL/well TURBO DNAse (Invitrogen AM2239). Following elution, RNA was frozen in three aliquots to minimize freeze-thaw cycles required. With the Maxwell, the Maxwell® RSC PureFood GMO and Authentication Kit (Promega) was used as described [44].

### Quantitative PCR

As described [44], RT-qPCR was used to measure barcoded-rRNA and 16S rRNA using the conditions and primers in Tables S3-S4. For qPCR reactions, an Applied Biosystems^TM^ MicroAmp® Fast 96-Well Reaction Plate, 0.1 mL (4346907) and Optical Adhesive Film (4311971) were used with a QuantStudio 5 PCR System (Thermo Fisher Scientific). Standard curves were used to quantify RNA copies per µL (Figure S1).

### Amplicon sequencing

Following RNA extraction, cDNA was prepared in a 6.75 µL reaction containing: 450 ng RNA, 0.1 µM reverse primer, and 0.5 mM dNTPs. This reaction was incubated for 5 minutes at 65°C and 5 minutes on ice. For the barcode-rRNA reactions, 0.25 µM of a blocking oligo was added to prevent amplification of unspliced ribozyme using 2 reactions for each sample. A master mix (3.25 µL) was added to each sample, which contained: 0.5 µL of DTT (100 mM), 0.5 µL of RNAseOUT (Thermo Fisher Scientific, 10777019), and 0.25 µL SuperScript^TM^ IV reverse transcriptase, and 2 µL of corresponding 5x buffer. Samples were mixed and incubated at 55°C for 20 minutes and 80°C for 10 minutes. All PCR reactions performed used Q5 polymerase and buffer (NEB) and dNTPs at a final concentration of 200 µM. The cDNA was amplified using a 25 µL reaction containing 2.5 µL of cDNA, 0.1 µM of blocking oligo (barcoded reaction only), and 1.25 µL primers (10 µM). Amplification was performed by incubating reactions at 98°C (5 minutes), then 25 to 30 PCR cycles (30 seconds at 98°C, 45 seconds at 60°C, and 60 seconds at 72°C), followed by 5 minutes at 72°C. Products were separated using 2% agarose gels and extracted. Two PCR protocols were used to add Illumina adaptors. To amplify native 16S rRNA, a single reaction (30 µL) was performed using gel extract (1.5 µL) as template and 1.5 µL of 10 µM primers. To amplify barcoded-rRNA, two parallel reactions (30 µL) were performed for each sample using a larger amount of gel extract (10 µL), no blocking oligo, and 1.5 µL of each primer (10 µM). Amplification was performed using 10 to 12 PCR cycles. The products were purified, yields were quantified using a Qubit, and the resulting products were sequenced by Genewiz using their Amplicon-EZ service.

### Transconjugant selection

To quantify transconjugants after incubating the different *E. coli* donor strains with *E. coli* JEC088, the cell mixtures (∼50 µL before growth at OD = 0.5) were resuspended in LB (1 mL), normalized by OD_600_, and serially diluted. Aliquots (6 µL) of each dilution were spotted on LB-agar containing kanamycin (100 µg/mL) to select for plasmid and 5-FOA (1 mg/mL) to select for the receiver strain (*E. coli* JEC088). After overnight incubation at 37°C, colony-forming units (cfu) were quantified.

### Isolating wastewater plasmids

Following incubation of *E. coli* and wastewater for 24 hours on filter paper placed on LB-agar, cells were harvested and resuspended in PBS (500 µL), and an aliquot was spread on LB agar containing kanamycin (100 µg/mL). Following 18 hours at 23°C, cells were harvested and resuspended in PBS. Extracted plasmid DNA (Qiagen Miniprep Kit) was analyzed using nanopore sequencing.

### Sequence analysis

A previously described protocol was used with QIIME 2 2024.10 to generate a filtered ASV feature table and assign taxonomy [44]. This was mapped onto a rooted tree using a custom python script and the interactive tree of life online tool [49]. The custom script is on GitHub (https://github.com/MalynSelinidis/Manuscript-3-RAM-Biocontainment/tree/main).

### Statistics

For RT-qPCR data, Cq were converted to RNA copies/µL using standard curves, and values were compared using a student t-test. For community NGS data, the host range was evaluated using a Principal Coordinates Analysis (PCoA) followed by permutational multivariate analysis of variance (PERMANOVA) using 999 permutations and data from all biological replicates. To calculate deviations from barcode ratios, the Analysis of Composition of Microbiomes with Bias Correction (ANCOMBC) QIIME plugin was used with Holm’s p-value adjustment [50].

## RESULTS

### Monitoring DNA mobility using RNA barcoding

Initial microbiome studies using RAM leveraged this barcoding tool to report on the host range of a small mobilizable conjugative plasmid [44]. To understand the sensitivity of RAM for studying the exchange of a wider range of DNA commonly engineered for synthetic biology, we encoded the same cat-RNA into three types of DNA elements (Figure 1A), including: (i) a 60 kb RP4 plasmid containing the incP origin of replication and incP origin of transfer (OriT) [51], (ii) a 5 kb plasmid with the pBBR1 and mobP OriT that requires strains with the RP4 transfer machinery for conjugation [44], and (iii) the genome of *Escherichia coli* MG1655. The pBBR1 plasmid was transformed into *E. coli* MFDpir and *E. coli* K12 to create strains with mobilizable and non-mobile plasmids, respectively, while the RP4 plasmid was transformed into *E. coli* K12 to create a strain with a self-mobilizable plasmid. In each strain, the cat-RNA was transcribed using a CymR repressible promoter (P_cym_).

**Figure 1.**
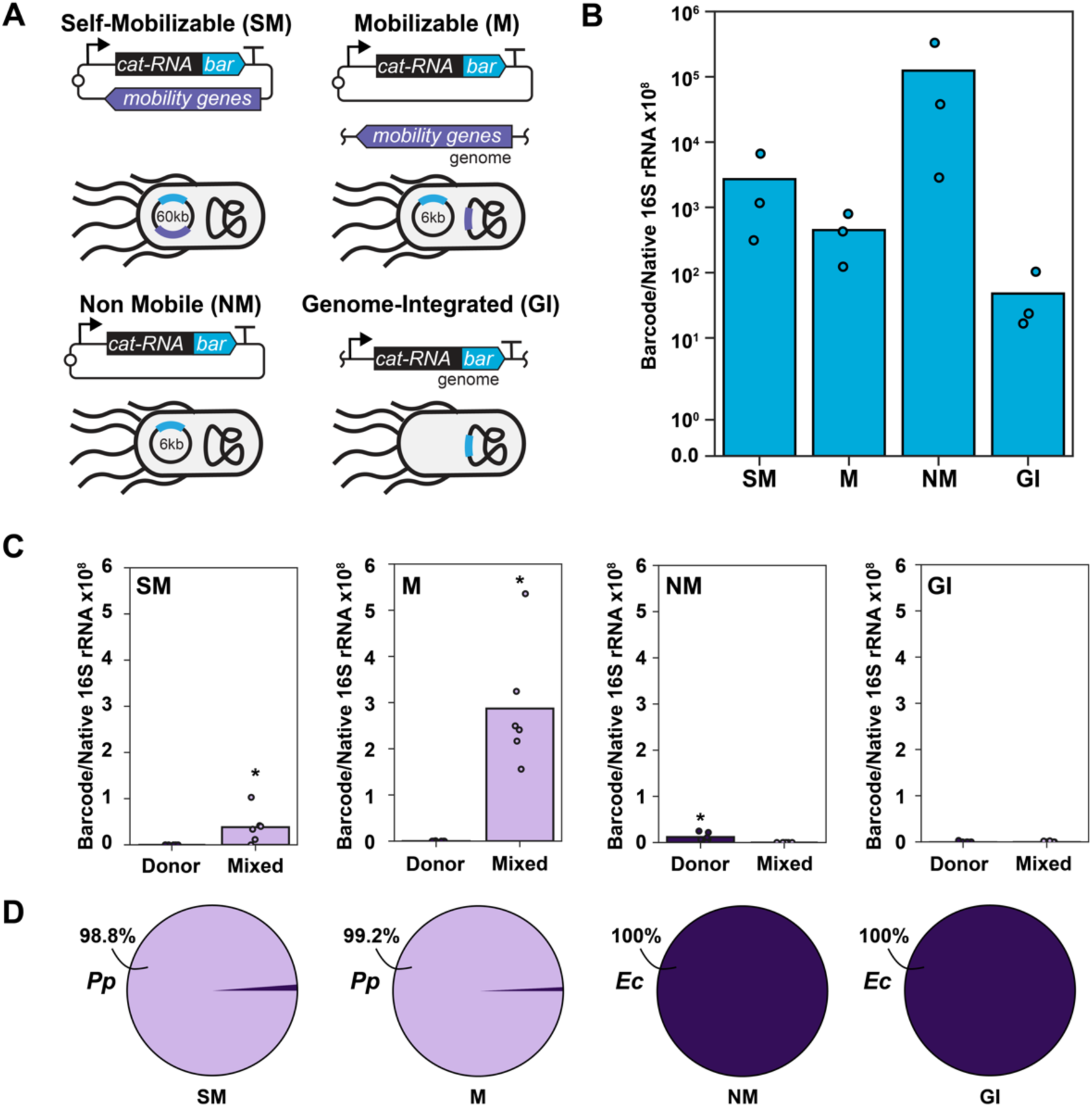
Monitoring the transfer of different DNA using cat-RNA. (**A**) *E. coli* strains created to transcribe cat-RNA from different DNA. The self-mobilizable (SM) plasmid has the mobility genes within the plasmid while the mobilizable (M) plasmid can only be conjugated if those genes are in the donor strain genome. (**B**) All four strains present a barcoded-rRNA signal, which was normalized to total 16S rRNA for comparison. *E. coli* lacking cat-RNA was below the detection limit. (**C**) The barcoded-rRNA signal following incubation of the *E. coli* strains (donor) alone and with *P. putida* (mixed). Only the mobilizable and self-mobilizable plasmids presented significantly higher signals with P. putida compared to the donor alone (asterisks; Student’s t test, p < 0.05). (**D**) Targeted sequencing of barcoded-rRNA from the mixtures yielded the relative frequencies of *E. coli* donor self-barcoding (brown) and *P. putida* barcoding arising from conjugation (purple).

To evaluate whether cat-RNA are competent for barcoding 16S rRNA when coded into these four different types of engineered DNA, each strain was grown in the presence of cumate, which derepresses the P_cym_ promoter, and barcoding was evaluated using RT-qPCR. Barcoded-rRNA was observed with each strain (Figure S2), while *E. coli* MG1655 lacking cat-RNA had no detectable signal. Normalization of the barcoded-rRNA to native 16S rRNA revealed variation in the barcoding signal across each strain (Figure 1B). These results show that a cat-RNA signal can be detected in strains that transcribe cat-RNA from different types of engineered DNA, including plasmids (self-mobilizable, mobilizable, and non-mobile) and the chromosome, whose signals span four orders of magnitude.

To establish whether cat-RNA can report on HGT when encoded on different DNA types, we evaluated conjugation with *Pseudomonas putida*. For these experiments, cat-RNA expression was repressed in each *E. coli* donor strain using the CymR repressor (Figure S3). Following a one-day incubation of each *E. coli* and *P. putida* mixture, RT-qPCR was used to quantify barcoded-rRNA (Figure S4). With *E. coli* containing the self-mobilizable and mobilizable plasmids (Figure 1C), the barcoding observed with *P. putida* exceeded the signal in the donor strain alone. In contrast, barcoding was not detected when *P. putida* was mixed with *E. coli* containing the non-mobile plasmid or the genome-encoded cat-RNA. Nanopore sequencing revealed that >98% of the barcoded-rRNA counts were from *P. putida* when mixed with *E. coli* having the self-mobilizable and mobilizable plasmids (Figure 1D), while *P. putida* sequences were not observed when mixed with *E. coli* containing the non-mobile plasmid and genome-encoded cat-RNA. These findings show that cat-RNA can report on the transfer of the self-mobilizable and mobilizable plasmids from *E. coli* to *P. putida* using either RT-qPCR or targeted sequencing.

### Mobilizable and self-mobilizable plasmid host ranges in wastewater

The mobilizable and self-mobilizable plasmids both use RP4 transfer machinery for conjugation, although they differ in their origins of replication (incP versus pBBR1) and the ways they encode the RP4 machinery (plasmid versus genome encoded). To investigate whether these differences affect their host range, we conjugated them in parallel into a wastewater community [52]. To allow direct comparison of plasmid host range in the same community, we added a 5 bp sequence to the cat-RNA barcode within the self-mobilizable plasmid so that the two barcodes can be differentiated when sequencing barcoded-rRNA (Figure 2A). In a prior study, these orthogonal cat-RNA presented similar signals in *E. coli* [44].

**Figure 2.**
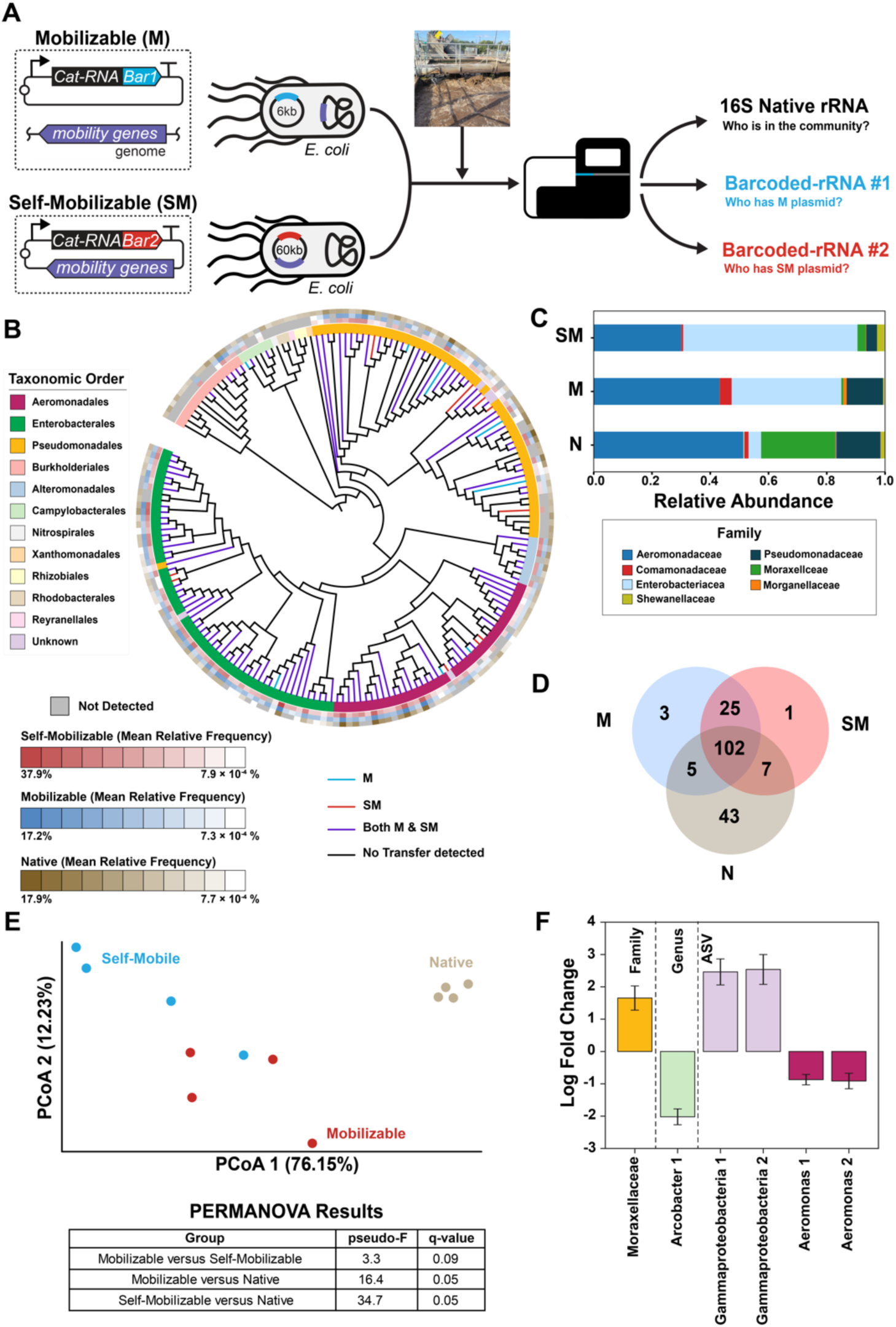
Comparing mobilizable and self-mobilizable plasmid host ranges in a wastewater community. (**A**) Mobilizable (M) and self-mobilizable (SM) plasmids with cat-RNA that splice orthogonal barcodes onto 16S rRNA were conjugated in parallel from *E. coli* into a wastewater community (4 biological replicates). (**B**) A phylogenetic tree is colored to indicate which ASVs had barcodes arising from conjugation. Outer leaflets show total rRNA (brown), M barcode (blue), and SM barcode (red) abundances. (**C**) Family-level comparison of total 16S rRNA and barcoded-rRNAs. (**D**) Venn diagram comparing ASVs in total 16S rRNA and barcoded-rRNAs. (**E**) PCoA of ASVs using a weighted unique fraction (unifrac) distance matrix and PERMANOVA. (**F**) Taxonomic groups that present significant differences (q<0.05) in the log fold ratios of the SM/M NGS read counts using an ANCOMBC analysis with Holm’s p-value correction. Bars colored by taxonomic order.

*E. coli* transformed with the mobilizable and self-mobilizable plasmids were mixed at equal titers and used as donors for conjugation into a wastewater sludge community. Following mating for one day, RT-qPCR was used to quantify the barcoded-rRNA in each sample (Figure S5). All four biological replicates presented strong barcoded-rRNA and total 16S rRNA signals compared with wastewater samples mixed with *E. coli* lacking a cat-RNA. To compare plasmid host ranges, the barcoded-rRNA and native 16S rRNA were analyzed using NGS (Figure 2B). With native 16S rRNA, 157 ASVs were observed, with *Aeromonadaceae*, *Moraxellaceae*, and *Pseudomonadaceae* representing the most abundant families (Figure 2C). With the barcoded-rRNA sequencing, a smaller number of ASVs were observed (n = 143), with *Aeromonadaceae* and *Enterobacteriaceae* representing the most abundant ASVs. In total, 127 of these ASVs presented barcoding signals for both the mobilizable and self-mobilizable plasmids (Figure 2D). For the six most abundant families observed with barcoded-rRNA, which spanned four classes (*Gammaproteobacteria*, *Alphaproteobacteria*, *Campylobacteria*, and *Nitrospiria*), more than half of their ASVs presented a barcoded-rRNA signal (Figure S6). These results show that orthogonal cat-RNA can be used to report on mobilizable and self-mobilizable plasmid conjugation in parallel.

To determine whether there are significant differences in the host ranges of the mobilizable and self-mobilizable plasmids, we performed a PCoA using our different biological replicates. A weighted unique fraction distance matrix followed by a PERMANOVA revealed that the ASVs observed with native 16S rRNA sequencing were significantly different from the ASVs observed with each barcode (Figure 2E). In contrast, the ASVs arising from the mobilizable and self-mobilizable plasmids were not significantly different. To investigate whether variations in plasmid abundances could be observed in specific taxonomic groups, we evaluated how the fraction of self-mobilizable reads for each ASV relates to their abundances within each order (Figure S7). With this analysis, many orders presented similar average signals for the two barcodes (Figure 3A), with large variation across individual ASVs from each order (Figure 3B). At lower taxonomic levels, significant variation in barcoding was observed across different taxonomic levels (Figures 2F, S8). The ASVs from the *Moraxellaceae* family presented significantly lower average abundance for the barcode from the mobilizable plasmid. Overall, this analysis reveals that the mobilizable and self-mobilizable plasmids exhibit similar host ranges across all six orders presenting barcoded-rRNA signals in the community, although differences at lower taxonomic levels are observed.

**Figure 3.**
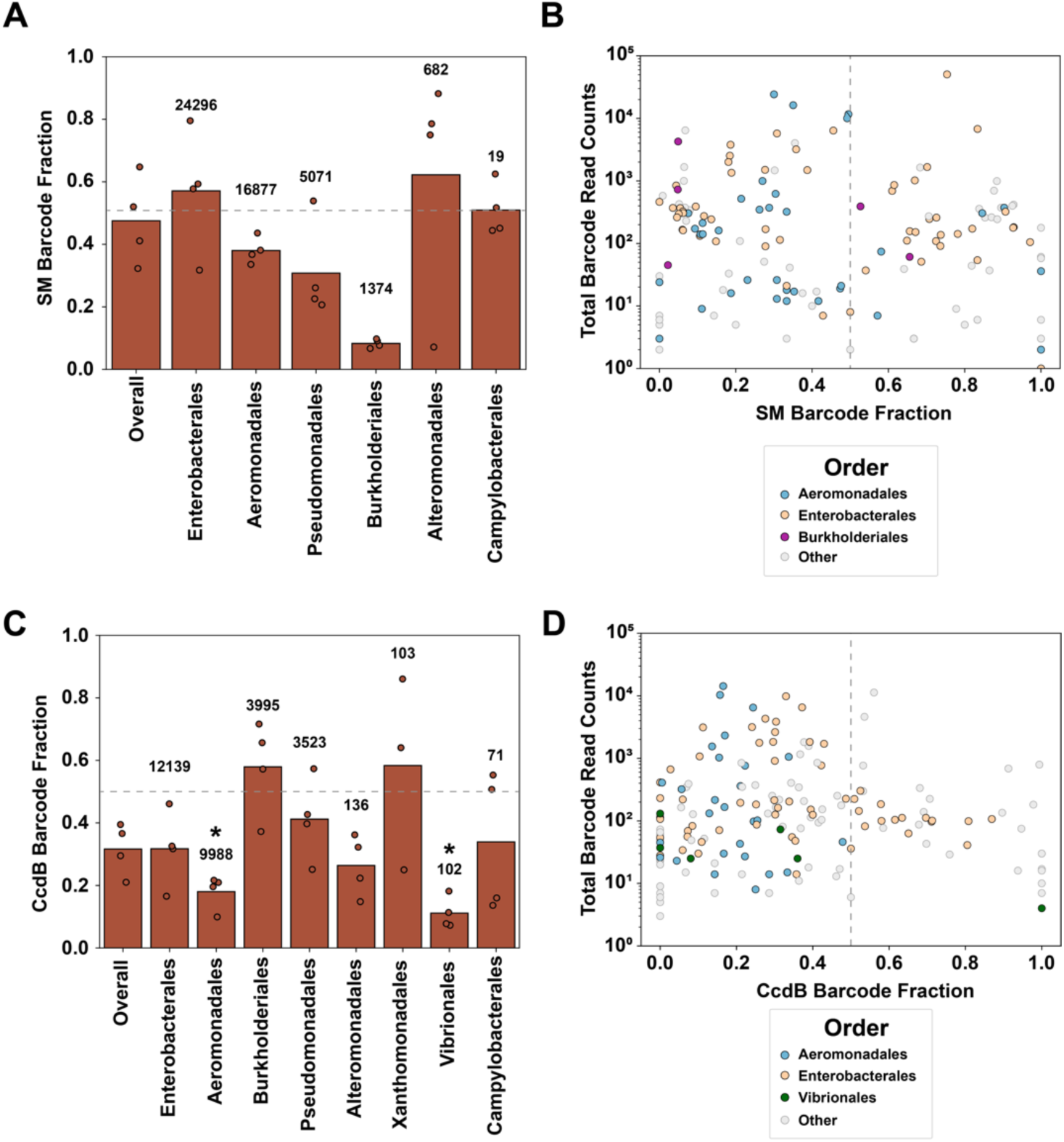
Comparing the barcoded-rRNA signals from conjugation of pairs of plasmids into wastewater communities. (**A**) For conjugations comparing mobilizable (M) and self-mobilizable (SM) plasmids, the SM plasmid read fraction is shown at the order level, and (**B**) the average ASV counts from replicates are shown relative to the SM barcode fraction in the barcoded-rRNA reads (=SM / M + SM counts). (**C**) For the conjugations comparing mobilizable plasmids ±CcdB, the CcdB plasmid read fraction is shown at the order level, and (**D**) the individual ASV counts are shown relative to the CcdB plasmid fraction (+CcdB / -CcdB + CcdB counts). In panels A and C, the average barcoded-rRNA counts from replicates are shown above bars, the dashed line represents the value expected when both plasmids yield similar barcoding signals, and the asterisks indicate significant differences (q<0.05) in the counts from the two plasmid types, calculated using an ANCOMBC with Holm p-value correction.

### Effect of a genetic biocontainment on HGT in a microbial community

To control the spread of engineered DNA in communities, genetic biocontainment strategies have been developed that conditionally express toxins following HGT into microbiomes [26–28], thereby killing any native microbes that take up the engineered DNA. To allow the engineered microbe to persist, protein toxin activity is inhibited using antitoxins. Currently, it is unclear how the host range of commonly used toxins and their genetic stability varies across communities, since their performance is typically evaluated using a small number of species.

To create a plasmid for studying toxin-based biocontainment in wastewater (Figure 4A), we modified the mobilizable plasmid so that it constitutively expresses the *E. coli* DNA gyrase inhibitor CcdB [53]. We also cloned an expression cassette for *E. coli* CcdA, a CcdB antitoxin, into the plasmid used to express CymR in the donor strain [54]. This toxin-antitoxin system was chosen because it is found in F plasmids that are widespread in *Enterobacteriales* [55], an abundant order in wastewater communities [44], and because the protein target for CcdB (DNA gyrase) is highly conserved (∼75% identity) in other wastewater orders like *Aeromonadales*. While related toxin-antitoxin systems are found in *Aeromonadales* [56], these systems have divergent sequences (≤35% identity), suggesting that *E. coli* CcdB may function more consistently in those orders compared with *Enterobacteriales*.

**Figure 4.**
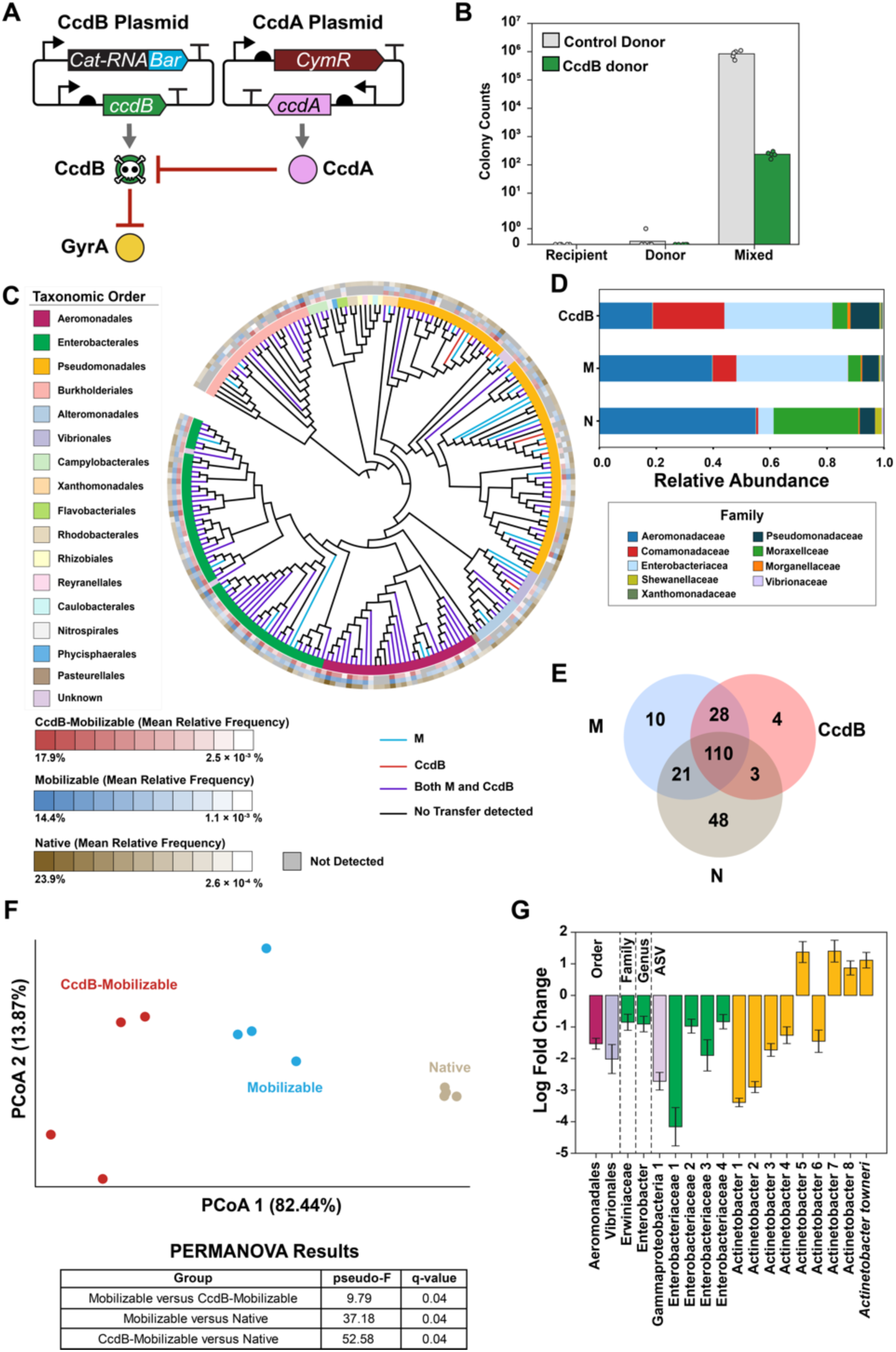
Effect of CcdB on mobilizable plasmid host range in a community. (**A**) A pair of mobilizable plasmids with and without the toxin CcdB, whose inhibition DNA gyrase (GyrA) is repressed by CcdA. (**B**) Following conjugation between *E. coli* strains, transconjugants (Mixed) were quantified for donor strains with (CcdB donor) and without (Control Donor) CcdB (5 biological replicates). Donor and recipient cells alone are shown as controls. (**C**) A phylogenetic tree showing the results of conjugating both strains in parallel from *E. coli* into a wastewater community (4 biological replicates). The branch colors indicate which ASVs have barcodes representing the mobilizable plasmid with (M) or without (CcdB) the toxin. Outer leaflets show total rRNA (brown), M barcode (blue), and CcdB barcode (red) abundances. (**D**) Family-level comparison and (**E**) Venn diagram showing total 16S rRNA and barcoded-rRNAs. (**F**) PCoA of ASVs using a weighted unique fraction (unifrac) distance matrix and PERMANOVA. (**G**) Taxonomic groups, colored by order, presenting significant variation (q<0.05) in the log fold ratios of the CcdB/M NGS read counts using an ANCOMBC analysis with Holm’s p-value correction.

The toxicity of the CcdB plasmid was first assessed in *E. coli* MG1655 by evaluating cfu with and without CcdA plasmids (Figure S9). In the absence of CcdA, colonies were not observed. In contrast, colonies were observed with cells expressing CcdA, albeit with 10-fold fewer cfu for the best design compared with cells lacking both CcdB and CcdA. Similar results were observed with the *E. coli* MFDpir donor strain used for conjugation with communities (Figure S10). These experiments establish a genetic biocontainment system that can be used with cat-RNA to study the effects of toxins on the host range of an engineered mobilizable plasmid.

To first characterize biocontainment, an *E. coli* recipient strain (JEC088) was mixed with *E. coli* MFDpir containing mobilizable plasmids ±CcdB and a plasmid that expresses CcdA. To obtain transconjugants, we used antibiotics to select for strains containing the mobilizable plasmid using plates lacking DAP, which the donor strain requires to grow [48]. When conjugation was performed using a mobilizable plasmid lacking a toxin, ∼10^6^ cfu were observed (Figure 4B), while colonies could not be detected with the individual donor and recipient strains. When conjugation was performed using the mobilizable plasmid that expresses CcdB, the number of transconjugants was diminished by ∼10^4^ fold. Whole-plasmid sequencing revealed that the transconjugants containing plasmid arose from the fusion of the mobilizable-CcdB and the CymR-CcdA plasmids (Figure S11). These results show that expression of CcdB toxin from the mobilizable plasmid decreases HGT arising from conjugation by ∼10,000 fold, and they reveal how escape from this biocontainment can occur.

To evaluate biocontainment in a microbiome, orthogonal cat-RNA were encoded in mobilizable plasmids ±CcdB, and *E. coli* containing each were mixed at equal titers and used as donors for conjugation into a wastewater community. Following a one-day incubation, a barcoded-rRNA signal was observed using RT-qPCR (Figure S12). To evaluate the effect of biocontainment on the host range of conjugation, barcoded-rRNA and native 16S rRNA were sequenced. NGS revealed that 69% of all barcoded-rRNA reads for all replicates (83,904 out of 121,591) were from the plasmid lacking biocontainment while only 31% of barcoded-rRNA reads from the plasmid with CcdB (Figure S13), with 9 orders containing ASVs that presented barcoded-rRNA signals (Figure 4C). Taxonomic analysis of the barcoded-rRNA revealed a broader host range for the plasmid lacking CcdB (169 ASVs) compared to the plasmid containing CcdB (145 ASVs), with the most abundant families for both plasmids being from *Aeromonadaceae, Comamonadaceae,* and *Enterobacteriaceae* (Figure 4D). In total, 95% of the ASVs having a barcode from the CcdB plasmid were observed in the ASVs barcoded by the plasmid lacking CcdB (Figure 4E), while only 81% of the ASVs having a barcode from the plasmid lacking CcdB were observed with the biocontainment plasmid. These results suggest that CcdB reduces the magnitude of conjugation into the wastewater community.

To determine whether biocontainment affects the plasmid host range, we performed a PCoA using a weighted unique fraction distance matrix followed by a PERMANOVA test (Figure 4F). The community composition, determined using native 16S rRNA, was significantly different from the composition of the ASVs having barcoded-rRNAs. The two plasmids barcoded ASVs that were significantly different, which was interpreted as reflecting differences in conjugation host range and transfer frequency. These observations show that CcdB biocontainment significantly alters conjugative plasmid host range in a wastewater community.

To better understand how biocontainment affects barcoding at the order level, we evaluated how the fraction of barcode reads from the CcdB plasmid for each ASV relates to their abundances within each order (Figure S14) With this analysis, six of the eight most abundant orders presented a lower fraction of barcode-rRNA reads for the CcdB plasmid compared with the plasmid lacking the toxin (Figure 3C), with two presenting statistically significantly lower signals. Among the most abundant orders, the ASVs exhibited a wide range of variation in biocontainment (Figure 3D). With *Aeromonadales*, all the ASVs presented a barcoded-rRNA signal from the CcdB plasmid that was less than half of the reads from both plasmids. The *Aeromonadaceae* family had a significantly lower signal for the CcdB plasmid (q = 4E-16) as did the *Aeromonas* genera (q = 1E-24). Within this genus, 48% of the unique ASVs (15/31) also had significantly lower signals for the plasmid with CcdB (q < 0.05) (Figure S15). Like *Aeromonadales*, many *Enterobacteriales* ASVs had a lower fraction of barcoded-rRNA counts from the CcdB plasmid. However, greater variability in signals from the two plasmids was observed across the *Enterobacteriales*. Only 1 of the 4 families (*Erwiniaceae*) had a significantly lower signal for the CcdB plasmid (q = 0.02), 2 of the 10 genera (*Enterobacter* and *Pantoeae*) presented significantly lower signals for the CcdB plasmid (q ≤ 0.009), and 5 of the 59 ASVs had significantly lower signals for the CcdB plasmid (q < 0.05) (Figures 4G, S15). At the family, genus, and ASV levels, additional groups presented significant variation in signals from the two plasmids (Figures 4G, S16). These results show that CcdB significantly reduces plasmid levels in a subset of taxonomic groups following HGT, and it suggests that biocontainment performance is more consistent across Aeromonadales compared with Enterobacteriales.

To better understand how wastewater microbes persist following acquisition of the CcdB plasmid, we conjugated the mobilizable ±CcdB (Figures 5A-B) into wastewater communities in separate reactions and selected for microbes that grew on kanamycin, which selects for the conjugative plasmids. Agarose gel analysis of plasmids in wastewater revealed DNA with a range of sizes (Figure S17). Following conjugation of the mobilizable plasmid lacking CcdB, an additional plasmid was observed whose size is consistent with the mobilizable plasmid. In contrast, the mobilizable plasmid with CcdB could not be visualized. We next sequenced the purified plasmid DNA from all three samples. Nanopore sequencing revealed that CcdB attenuated the abundance of the mobilizable plasmid (Figures 5C-D, S18). With conjugation of the CcdB plasmid, only 0.17% of reads (13 of 7,469 reads) were assigned to this engineered plasmid. In contrast, with the mobilizable plasmid lacking CcdB, 22.8% of all reads (1,608 of 7,037 reads) were assigned to the engineered plasmid. Mutational analysis of the CcdB plasmid revealed mutations in a subset (85%) of the plasmids (Figure 5E), none of which arose from recombination of the CcdB and CcdA donor strain plasmids (Figure S11) as observed in two-species conjugation reactions. In contrast, mutations were not observed with the plasmid lacking CcdB. Taken together, these results show that biocontainment escape can arise with and without mutations to the engineered plasmid.

**Figure 5.**
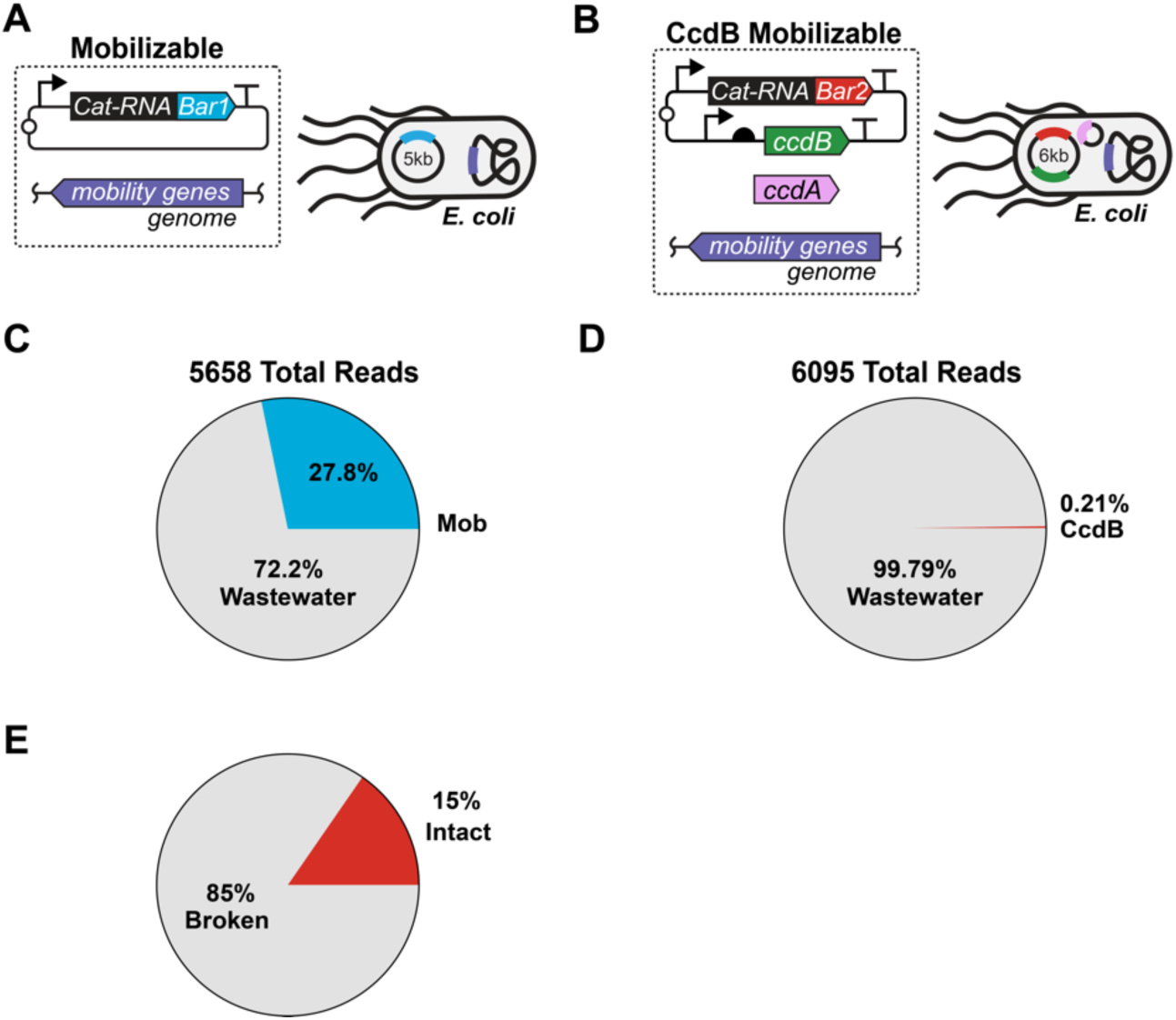
CcdB decreases the abundance of a mobilizable plasmid in community. Conjugation in wastewater was performed using an *E. coli* donor strain containing mobilizable plasmids (**A**) lacking CcdB or (**B**) containing CcdB. Following incubation for one day followed by selection for kanamycin resistance on plates, total plasmids were isolated and sequenced. Among all plasmids sequenced, the relative abundances of the mobilizable plasmids (**C**) lacking CcdB and (**D**) containing CcdB are shown. (**E**) Fraction of the CcdB plasmid sequences with mutations.

## DISCUSSION

Here, we show that RAM can be used to evaluate the effects of plasmid sequence variation on HGT in microbiomes. Importantly, we find that the conditional expression of a protein toxin differentially affects the abundance of a mobilizable plasmid across different taxonomic groups immediately following a one-day mating assay. While numerous toxin-antitoxin biocontainment systems have been described [32–35], only a small fraction have been tested in microbiomes. These tests have largely focused on measuring their biocontainment efficiencies [57–60], rather than plasmid host range and lethality across different taxonomic groups. Our findings provide evidence that the lethality of toxin-antitoxin biocontainment systems varies across the taxonomic groups that participate in HGT, critical information for establishing the safety of these biotechnologies in open environment [2, 17, 61]. They also show how RAM will be useful for benchmarking the performance of different biocontainment systems and for probing their performance across longer time scales and a wider spectrum of environmental conditions [21–23].

Our biocontainment measurements also reveal variability across different taxonomic orders in wastewater. The CcdB expression cassette decreased the average barcoding signal from the mobilizable plasmids across three of the four most abundant orders. *Aeromonadales* presented the most consistent reduction in signal from the CcdB plasmid regardless of the total amount of barcoded-rRNA observed for each ASV. *Enterobacteriales* presented a larger variance in CcdB plasmid signal, even among the ASVs with the highest numbers of reads. Several parameters could underlie this variability. First, heterogeneity in the abundance of related CcdA antitoxins across the community could lead to variation in *E. coli* CcdB toxicity. For example, the larger variance in CcdB plasmid signal within *Enterobacteriales* compared with *Aeromonadales* could arise because a subset of *Enterobacteriales* contain F plasmids that encode closely-related CcdB/CcdA toxin-antitoxin systems used for biocontainment [62]. Second, the expression of *E. coli* CcdB following HGT is expected to vary widely across non-native hosts, as observed with synthetic promoters and fluorescent reporters [63, 64]. Third, DNA gyrases in transconjugants may bind *E. coli* CcdB with varying affinities if sequence variation occurs in the gyrase-CcdB interface [65]. Finally, variation in growth rates across transconjugants could affect CcdB toxicity, since the CcdB-gyrase interaction is toxic in cells that are actively replicating their DNA [66].

In future studies, RAM is expected to be useful for probing controls on mobile DNA host range and persistence in microbiomes. Models are emerging to anticipate both plasmid and phage host ranges in communities [67–71]. To date, these models have largely been trained using snapshots of mobile DNA host range acquired using high-throughput chromosome conformation capture, which requires >1000-fold more sequencing than RAM to generate host range data. This requirement limits the throughput for studying how the addition or removal of individual genes from mobile DNA affects host range *in situ*. In contrast, our results show that RAM is compatible with studies of plasmids that differ by a single genetic characteristic in microbiomes. By multiplexing RAM beyond pairs of conjugative plasmids, this technology should be able to generate large data sets that capture how sequence variation in plasmids relate to host range in microbiomes. Such information will be critical for building artificial intelligence models that accurately anticipate plasmid host range in communities [67–69, 72, 73], as well as controls on mobile DNA persistence across a range of ecological settings.

## AUTHOR CONTRIBUTIONS

Conceptualization: M.A.S., J.J.S., and J.C.; Investigation: M.A.S. and T.S.; Analysis: M.A.S., J.J.S., and J.C.; Drafting manuscript: M.A.S., J.J.S., and J.C.; Manuscript review and editing: M.A.S., T.S., L.B.S., J.J.S., and J.C.; Visualization: M.A.S., J.J.S., and J.C.; Supervision: J.J.S. and J.C.; and Funding; J.C., J.J.S., and L.B.S.

## SUPPLEMENTARY MATERIALS

Supplementary figures and tables provide more details on the RAM methodology, RT-qPCR biological replicates, NGS sequence analysis from community experiments, PCA analysis of community data, conjugation assays using plating, strain development, and plasmid sequences.

## FUNDING

This research was supported by the US Department of Agriculture (USDA) National Institute of Food and Agriculture (NIFA) Biotechnology Risk Assessment Grants (BRAG) program under award 2025-33522-45287 (J.C., J.J.S., and L.B.S) and National Science Foundation grants 2237052 (L.B.S.), 2237512 (J.C.), 2227526 (J.C. and J.J.S), and 2223678 (J.J.S. and L.B.S.) and the Welch Foundation grant C-2183-20240404 (J.C.) McSherry Poe Research Award (M.A.S). Nanopore sequencing of plasmid DNA from wastewater was sequenced by the Genetic Design and Engineering Center (GDEC) at Rice University, which is funded by CPRIT RP210116.

## DATA AVAILABILITY

The sequence datasets generated and analyzed for this study can be found at the Sequence Read Archive under accession number PRJNA1483784.

## ACKNOWLEDGEMENTS

The authors would like to thank Darren Seet and Lavanya Karinje for reagents and support troubleshooting; Lin Fang, Lily Metsker, and Maddie Wolken for wastewater acquisition support; and August Staubus and Matt Dysart for their experimental guidance throughout this project.

## Notes

### Competing Interest Statement

The authors have declared no competing interest.

