## Supplemental Figures 1 to 18 for "Biocontainment attenuation of mobile DNA host range in a wastewater microbiome": CcdB_RAM_SI_071326_3p.pdf

### TABLE OF CONTENTS


| Item | Title | Page |
|---|---|---|
| Fig. S1 | Standard curves for 16S rRNA and barcoded-rRNA | 3 |
| Fig. S2 | Barcoded-rRNA in strains having RAM coded in different DNA types | 4 |
| Fig. S3 | Repression system used to prevent self-barcoding in donor cells | 5 |
| Fig. S4 | Total rRNA and barcoded-rRNA for two-microbe conjugations | 6 |
| Fig. S5 | RAM presents a strong RT-qPCR signal in wastewater communities | 7 |
| Fig. S6 | Barcoded-rRNA reads for each plasmid type across taxonomic groups | 8 |
| Fig. S7 | Individual ASV NGS reads vs self-mobilizable plasmid fraction | 9 |
| Fig. S8 | Ratio of self-mobilizable to mobilizable RAM reads | 10 |
| Fig. S9 | Tuning CcdB expression in *E. coli* | 11 |
| Fig. S10 | CcdA complementation of *E. coli* expressing CcdB | 12 |
| Fig. S11 | Features in CcdB plasmids that escape biocontainment | 13 |
| Fig. S12 | Mobilizable plasmids ±CcdB yield a RAM signal in wastewater | 14 |
| Fig. S13 | Barcoded-rRNA reads for plasmids ±CcdB across taxonomic groups. | 15 |
| Fig. S14 | Individual ASV NGS reads vs fraction of CcdB plasmid reads | 16 |
| Fig. S15 | Ratio of NGS reads ±CcdB across ASVs | 17 |
| Fig. S16 | Ratio of NGS reads ±CcdB in orders, families, and genera | 18 |
| Fig. S17 | Agarose gel analysis of wastewater plasmids | 19 |
| Fig. S18 | Plasmid length distributions from NGS | 20 |
| Sequences | Plasmid sequences with annotations | 21 |

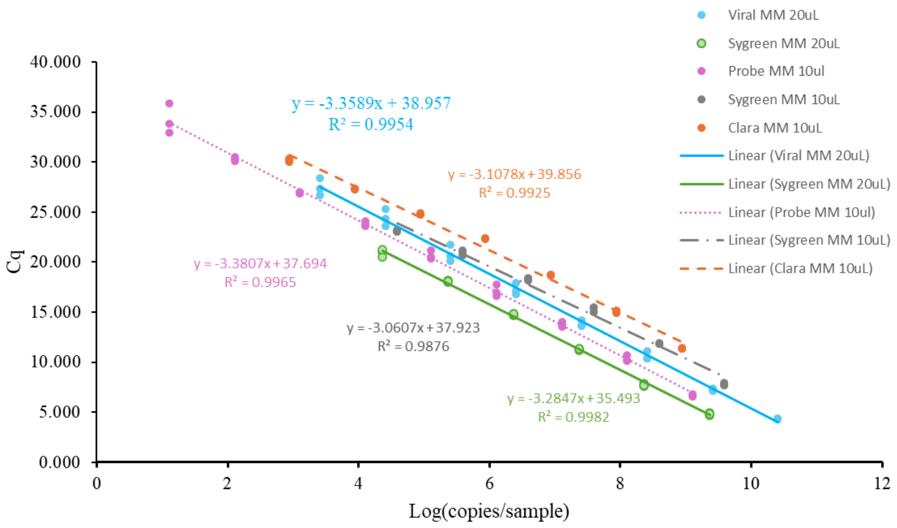


**Figure S1. Standard curves for 16S rRNA and barcoded-rRNA.** Amplicons of 16S rRNA and barcoded-rRNA were generated, their concentrations were determined using a Qubit, and serial dilutions of each were analyzed using qPCR (3 replicates). Standard curves are shown as copies/sample to account for variability at lower dilutions to allow for most accurate quantification.

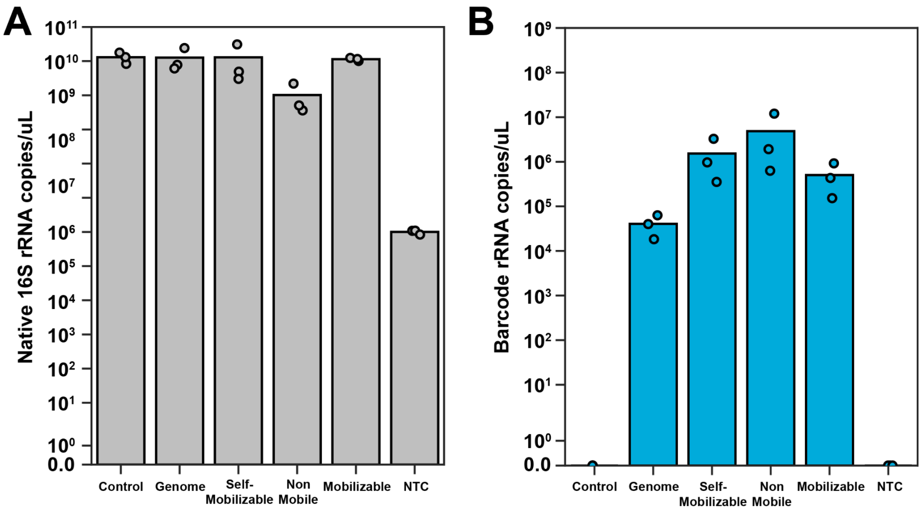


**Figure S2. Barcoded-rRNA in strains having RAM coded in different DNA types.** (**A**) Native 16S rRNA and (**B**) barcoded-rRNA detected in *E. coli* using RT-qPCR that transcribe RAM from the genome and mobilizable, self-mobilizable, non-mobile plasmids. Cells lacking RAM (Control) and a no template control (NTC) are shown as well. Data represents 3 biological replicates for cells grown in the presence of 100 μM cumate.

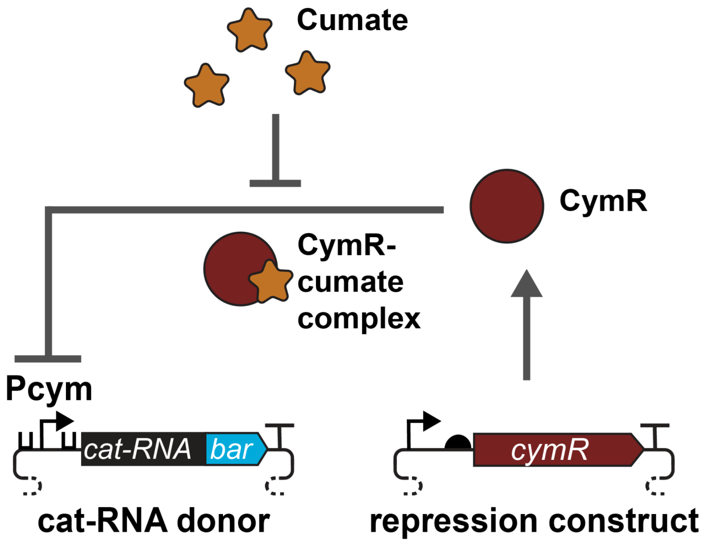


**Figure S3. Repression system used to prevent self-barcoding in donor cells.** To repress barcoding in strains having each DNA element type (self-mobilizable, mobilizable, non-mobile, and genome-integrated), the $P_{cym}$ promoter was used to regulate cat-RNA transcription. In cells that express CymR, transcription of the cat-RNA is repressed, unless cumate is provided which derepresses CymR.

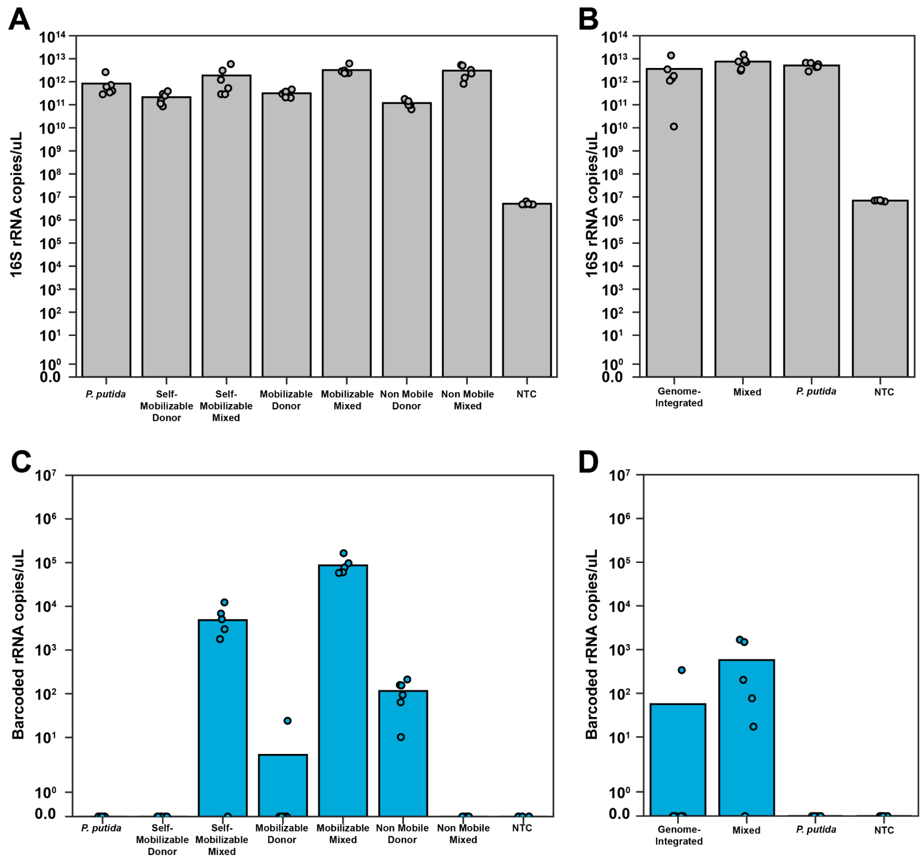


**Figure S4. Total rRNA and barcoded-rRNA for two-microbe conjugations.** The amount of total 16S rRNA was analyzed using RT-qPCR with (**A**) each *E. coli* strain having RAM coded in different plasmid types (Donor) and following incubation of those donor strains with *P. putida* (Mixed), and (**B**) the *E. coli* having a genome-integrated RAM alone and following incubation with *P. putida* (Mixed). The amount of barcoded-rRNA generated by RAM was also analyzed for (**C**) each *E. coli* donor alone and following incubation with *P. putida*, and (**D**) for the genome-integrated RAM. The recipient cell (*P. putida*) and no template control (NTC) are included to show that the barcoded-rRNA signal is dependent on RAM. Points represent 6 biological replicates.

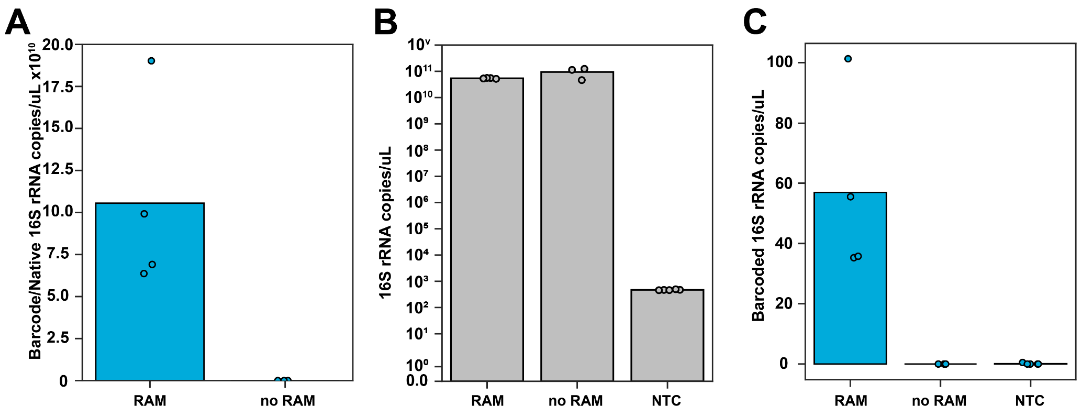


**Figure S5. RAM presents a strong RT-qPCR signal in wastewater communities.** (**A**) The barcoded-rRNA signal normalized to total 16S rRNA (n ≥ 3 biological replicates) following incubations of wastewater with either a mixture of *E. coli* donors having the self-mobilizable and mobilizable plasmids (RAM) or *E. coli* lacking RAM (no RAM). The RAM samples have significantly higher signals than the no RAM [student's t test ($t = 3.04$, $p = 0.028 < 0.05$)]. (**B**) The total 16S rRNA and (**C**) barcoded-rRNA is compared to a no template control (NTC).

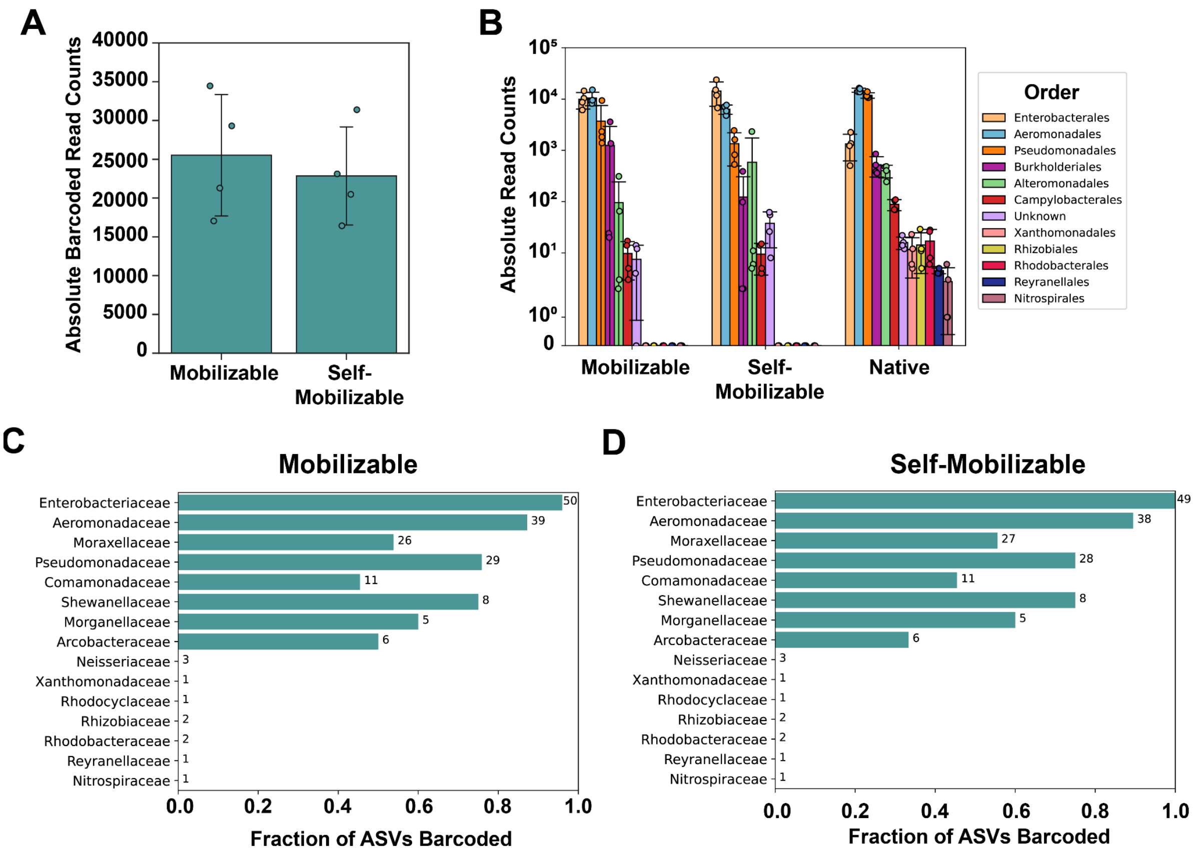


**Figure S6. Barcoded-rRNA reads for each plasmid type across taxonomic groups.** A comparison of the total barcoded-rRNA reads for (**A**) the two plasmid types conjugated into wastewater, and (**B**) across orders. The fraction of unique ASVs in each family that presented a barcoded-rRNA signal for the (**C**) mobilizable and (**D**) self-mobilizable plasmids. The numbers next to each bar represent the total number of ASVs observed for each family when sequencing barcoded-rRNA and total 16S rRNA. The data represents 4 biological replicates, shown as points in A and B. In panel A, the total plasmid counts were similar across both plasmid types [t test $t(6) = 0.53$, $p = 0.6$].

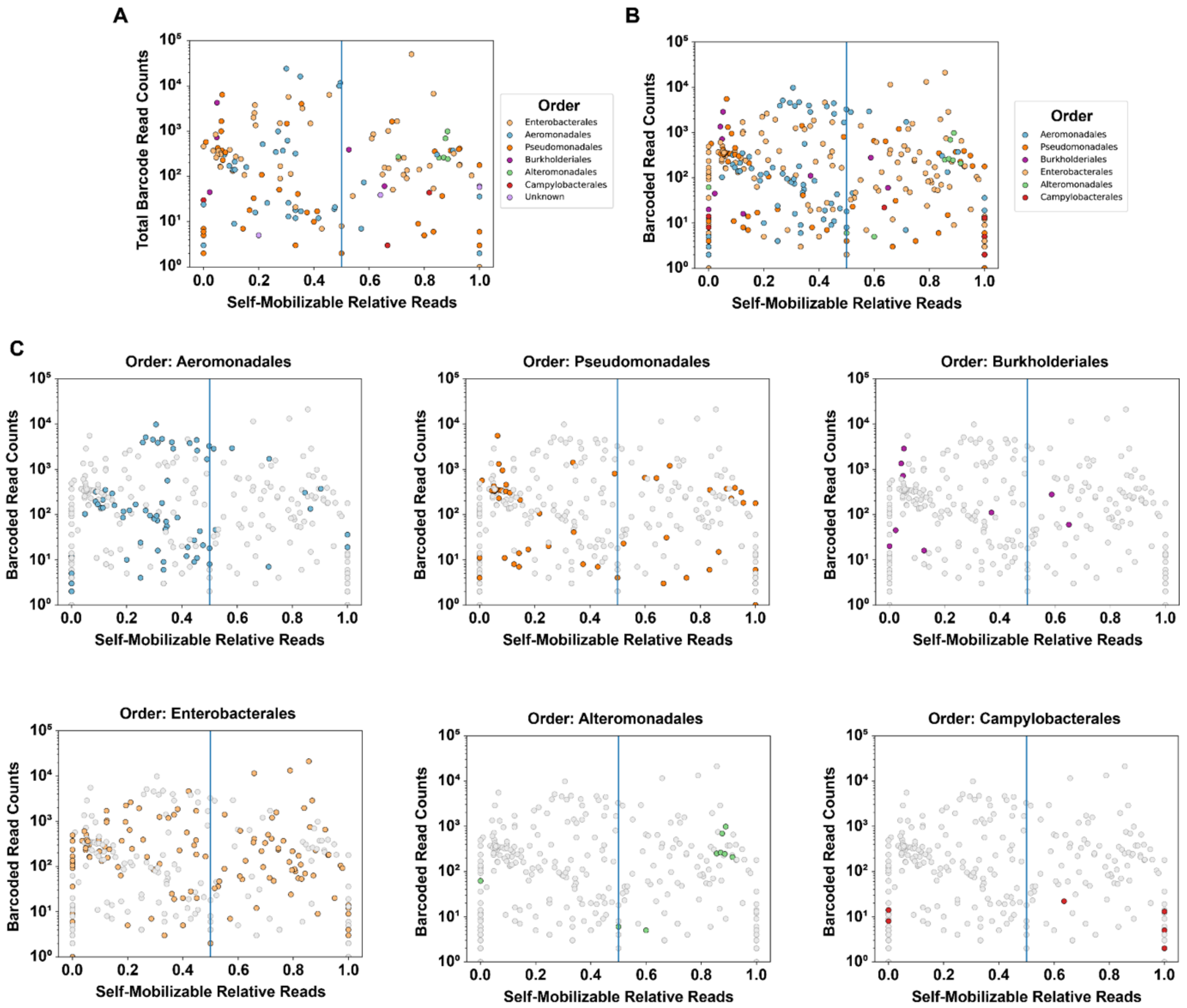


**Figure S7. Individual ASV NGS reads vs self-mobilizable plasmid fraction.** Data showing the trends when (**A**) summing the NGS reads for all ASVs and (**B**) plotting the data for each ASV from the different biological replicates (n = 4). (**C**) Data for ASVs from different orders are shown (colored points) versus all other orders (gray).

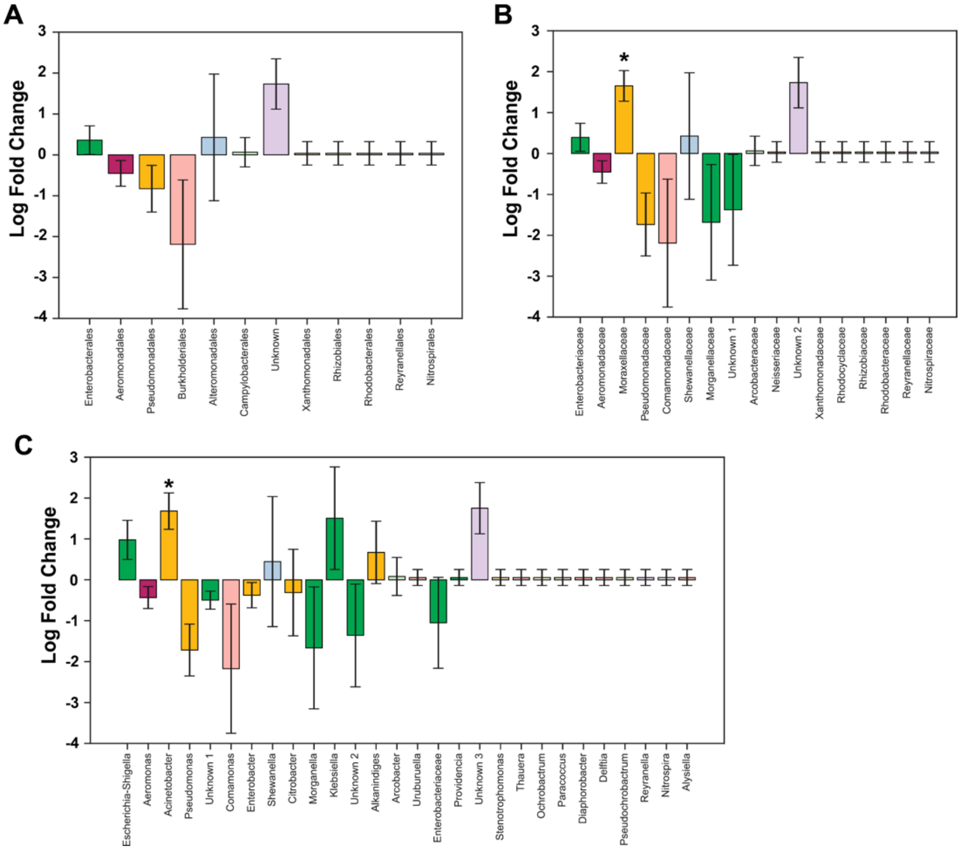


**Figure S8. Ratio of self-mobilizable to mobilizable RAM reads.** The log fold ratios of the ratio of self-mobilizable/mobilizable NGS reads from 4 biological replicates are shown for ASVs at the (**A**) order, (**B**) family, and (**C**) genus levels. Positive values represent a higher abundance of self-mobilizable reads, while negative values indicate a greater number of mobilizable reads. Bars colored by taxonomic order and error bars display standard error. Asterisk indicates significance ($q<0.05$) based on Analysis of Compositions of Microbiomes with Bias Correction (ANCOMBC) statistical analysis with Holm's p-value correction.

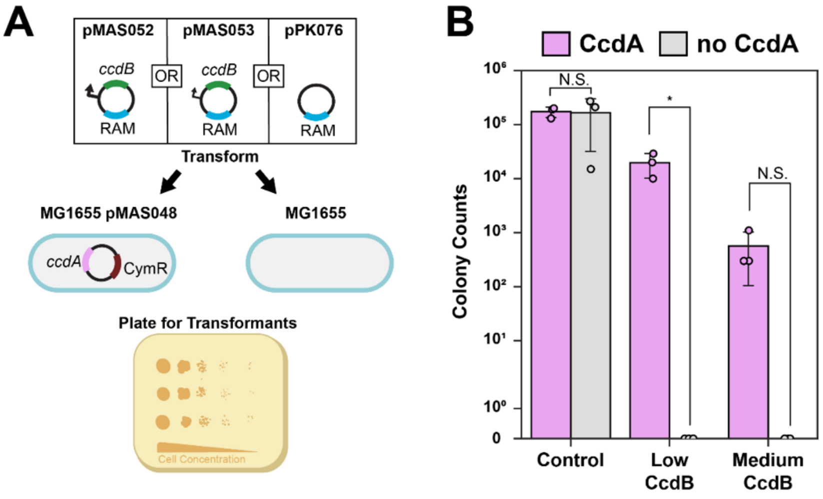


**Figure S9. Tuning CcdB expression in *E. coli*.** (**A**) Plasmids (pMAS052, pMAS053) that use low (Low CcdB) and medium (Medium CcdB) promoter strengths to express CcdB were transformed into *E. coli* MG1655 with or without a CcdA plasmid (pMAS048) and compared with transformation of a plasmid (pPK076) lacking CcdB (Control). (**B**) Colony counts observed in cells with CcdA (pink) and without CcdA (gray). The biological replicates ($n = 3$) are shown as points while error bars show standard deviation. The Low CcdB presents a significantly higher signal with CcdA compared to in the absence of the antitoxin [two-sample independent t test, $p < 0.05$]. The Low plasmid was used for all subsequent experiments.

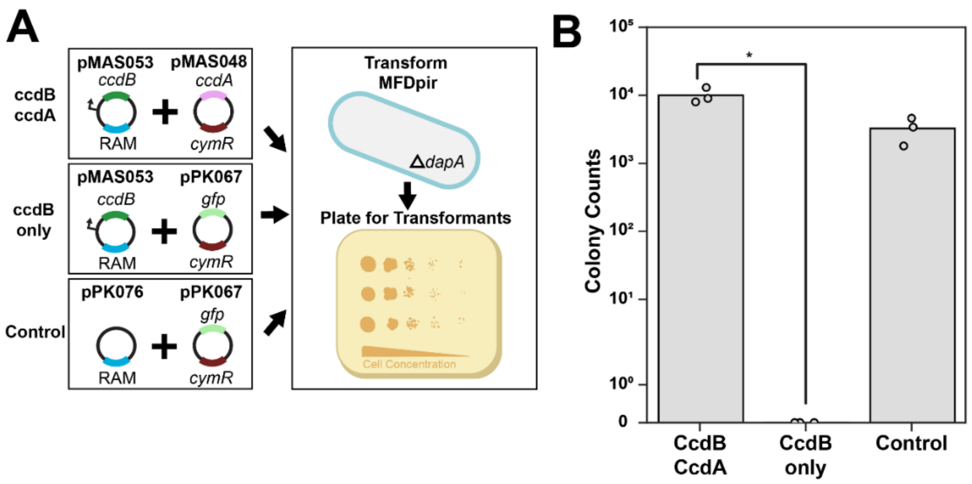


**Figure 10. CcdA complementation of *E. coli* expressing CcdB.** (**A**) Scheme for evaluating effects of CcdB and CcdA on *E. coli* MFDpir fitness. (**B**) Colony counts from *E. coli* co-transformed with pMAS053 and pMAS048 (CcdB/CcdA), pMAS053 and pPK067 that lacks CcdA (CcdB only), and a pair of vectors that lack CcdB and CcdA (Control). The biological replicates are shown as points (n=3). CcdB/CcdA has a significantly larger number of counts compared with CcdB [two-sample independent t test revealed, $p < 0.05$].

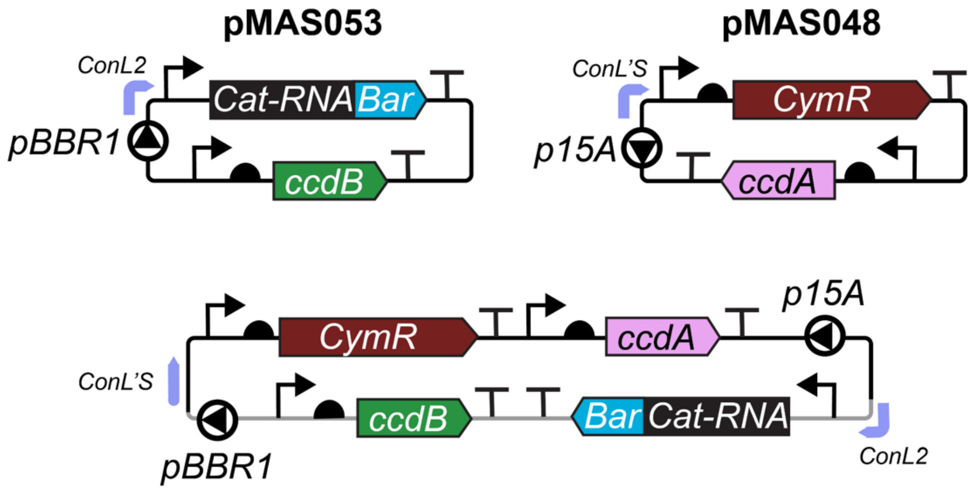


**Figure S11. Features in CcdB plasmids that escape biocontainment.** All three plasmids purified from colonies that escaped biocontainment contained a plasmid (*bottom*) that arose from recombination of the conjugative plasmid expressing CcdB (pMAS053) and the CcdA plasmid that enables donor *E. coli* to grow (pMAS048). The regions (ConL2 and ConL'S) where recombination occurs contain modified bidirectional terminators and multiple restriction sites.

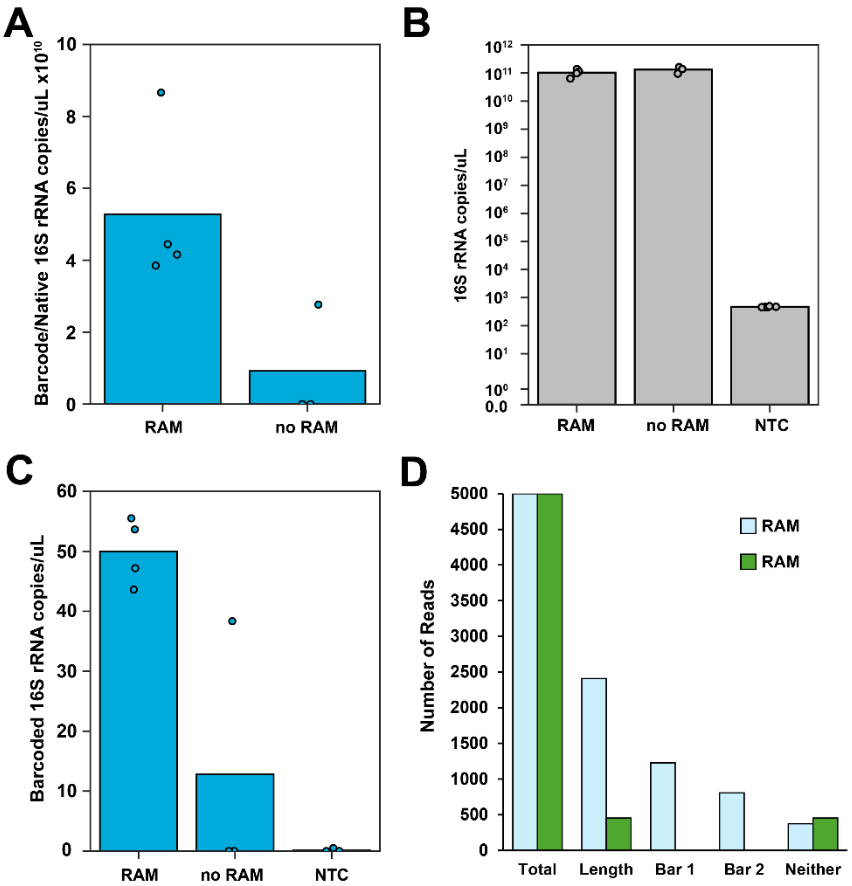


**Figure S12. Mobilizable plasmids ±CcdB yield a RAM signal in wastewater.** (**A**) The RT-qPCR signal for barcoded-rRNA normalized to native 16S rRNA following incubation of wastewater with a mixture of two strains (RAM), one with the mobilizable plasmid lacking CcdB and one with the mobilizable plasmid containing CcdB, or with an *E. coli* strain lacking those plasmids (no RAM). (**B**) The total 16S rRNA signals and (**C**) barcoded-rRNA are compared to a no template control (NTC). (**D**) The no RAM replicate that presented a signal was analyzed using nanopore sequencing and compared to a RAM replicate. With this analysis, the same number of total reads were returned. However, only the RAM sample had large numbers of barcoded-rRNA reads of the correct length with expected primers (Length) and with the different orthogonal barcodes (Bar 1, Bar 2) in the two different conjugative plasmids used for conjugation. Thus, a RAM signal could not be detected by nanopore with no RAM sample even though there was a small RT-qPCR signal, which we interpret as contamination.

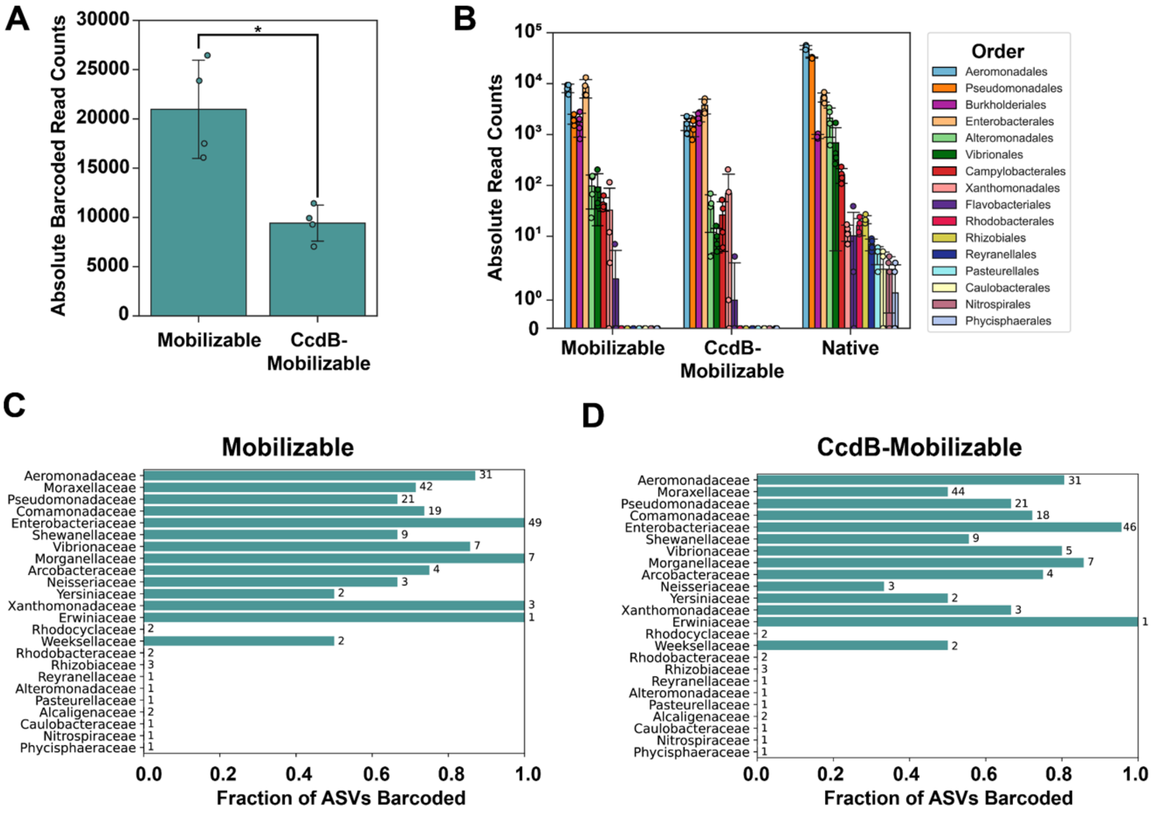


**Figure S13. Barcoded-rRNA reads for plasmids ±CcdB across taxonomic groups.** A comparison of the total barcoded-rRNA counts for (**A**) the mobilizable plasmids ±CcdB, and (**B**) across orders. In panel A, the total plasmid counts were significantly lower for the plasmid with CcdB biocontainment [t test $t(6) = 4.359$, $p = 0.005$]. The fraction of unique ASVs in each family that presented a barcoded-rRNA signal for the mobilizable plasmid (**C**) without CcdB and (**D**) with CcdB. The numbers next to each bar represent the total number of ASVs observed for each family. The data represents 4 biological replicates, shown as points in A and B.

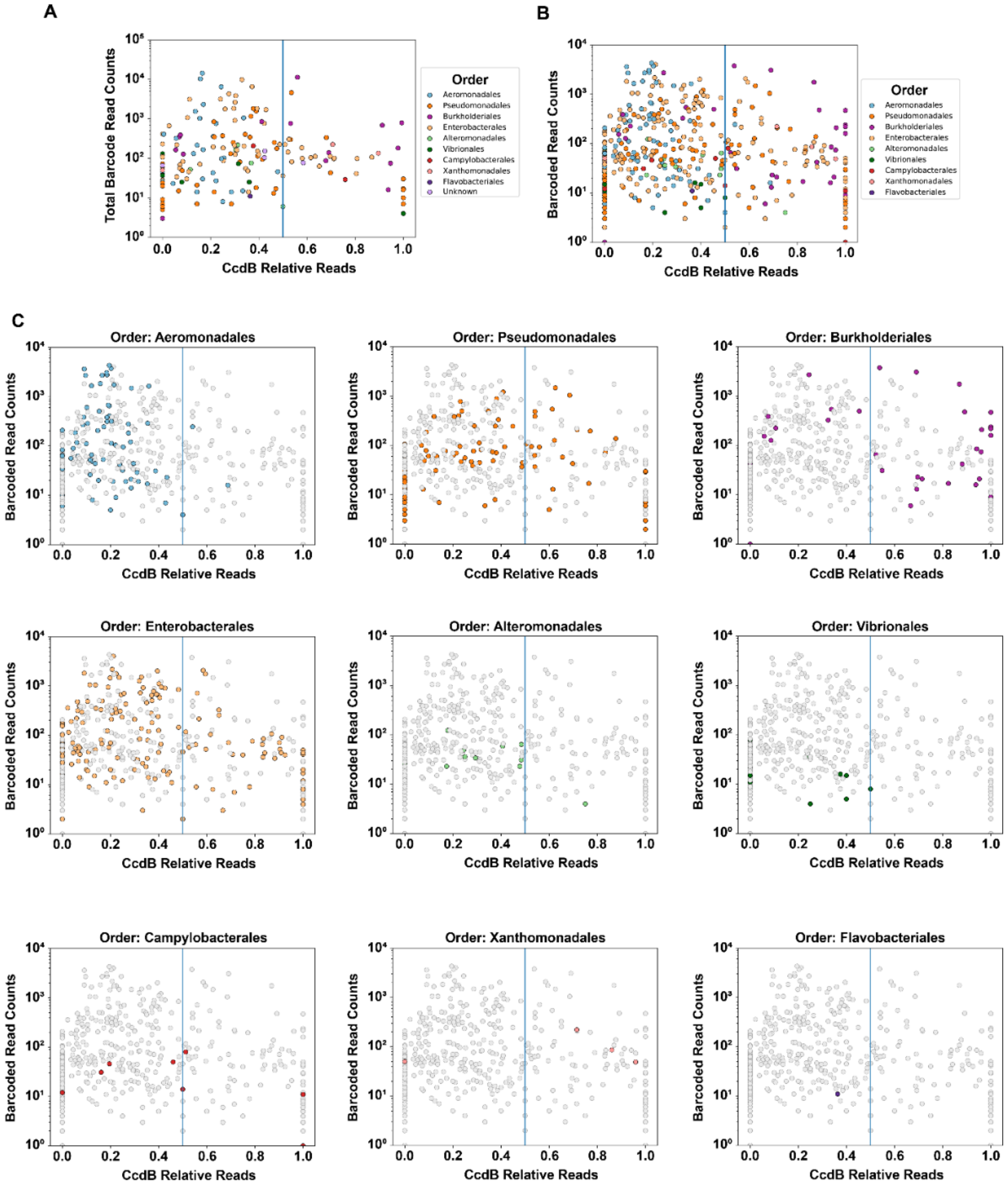


**Figure S14. Individual ASV NGS reads vs fraction of CcdB plasmid reads.** Data showing the trends when (**A**) summing the NGS reads for all ASVs and (**B**) plotting the data for each ASV from the different biological replicates (n = 4). (**C**) Data for ASVs from different orders are shown (colored points) versus all other orders (gray) to allow visualization of biocontainment performance.

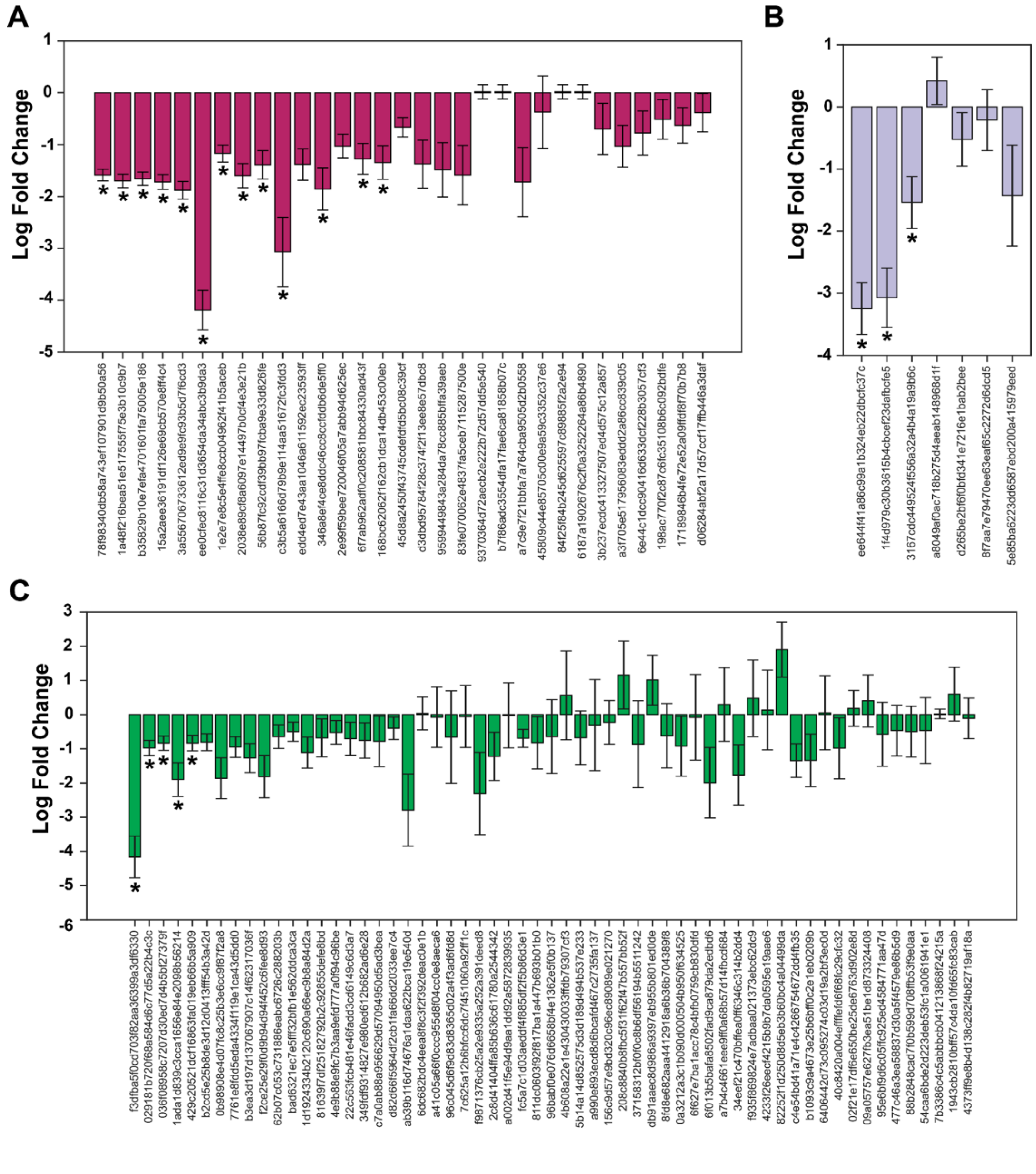


**Figure S15. Ratio of NGS reads ±CcdB across ASVs.** The log fold ratios of mobilizable plasmid reads (±CcdB) from 4 biological replicates are shown for ASVs within the orders (**A**) *Aeromonadales*, (**B**) *Vibrionales*, and (**C**) *Enterobacterales*. Asterisks indicate significance (q<0.05) based on ANCOMBC statistical analysis with Holm's p-value correction. Negative values indicate a larger fraction of the plasmid lacking CcdB, while positive values indicate a higher fraction of reads for the plasmid designed to express CcdB. The bars, which are colored by taxonomic order, show standard error.

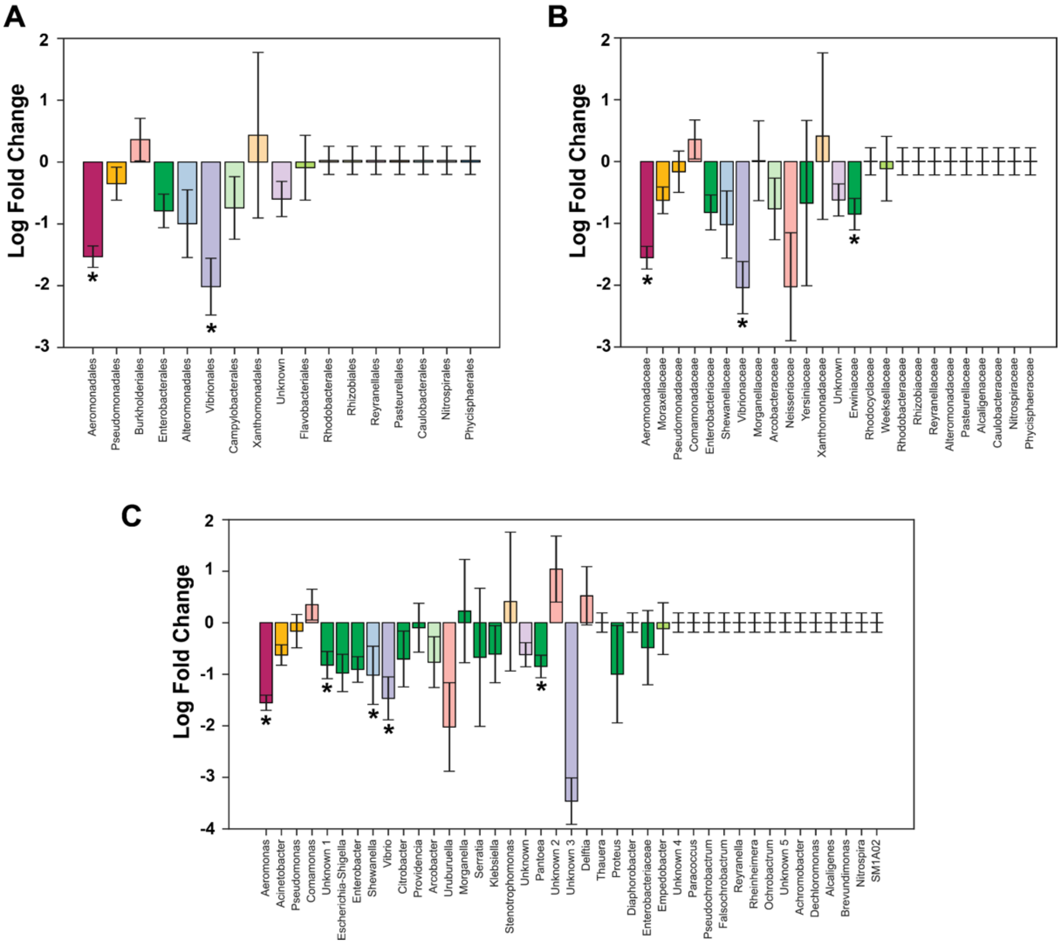


**Figure S16. Ratio of NGS reads ±CcdB in orders, families, and genera.** The log fold ratios of mobilizable plasmid reads (±CcdB) at the (**A**) order, (**B**) family, and (**C**) genus levels. Negative values indicate a larger fraction of the plasmid lacking CcdB, while positive values indicate a higher fraction of reads for the plasmid designed to express CcdB. Bars colored by taxonomic order represent data from 4 biological replicates, with error bars showing standard error. Asterisk indicates significance ($q<0.05$) based on ANCOMBC statistical analysis with Holm's p-value correction.

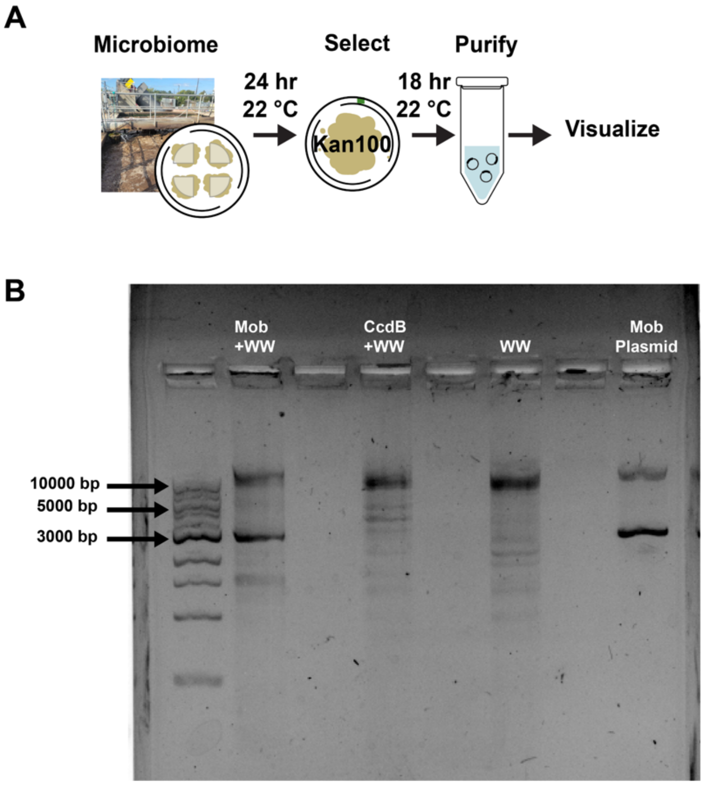


**Figure S17. Agarose gel analysis of wastewater plasmids.** (**A**) Schematic of conjugation protocol prior to analysis. Briefly, donor cells were incubated with a wastewater for 24 hours, the mixture was selected on plates containing kanamycin (100 µg/mL) for 22 hours, and plasmids were extracted. (**B**) An agarose gel showing the size distributions of the mobilizable plasmid (Mob Plasmid), the plasmids purified from untreated wastewater (WW), and the plasmids obtained following conjugation experiments in wastewater that used the mobilizable plasmid lacking (Mob+WW) and containing CcdB (CcdB+WW). Molecular weight markers are shown as a frame of reference.

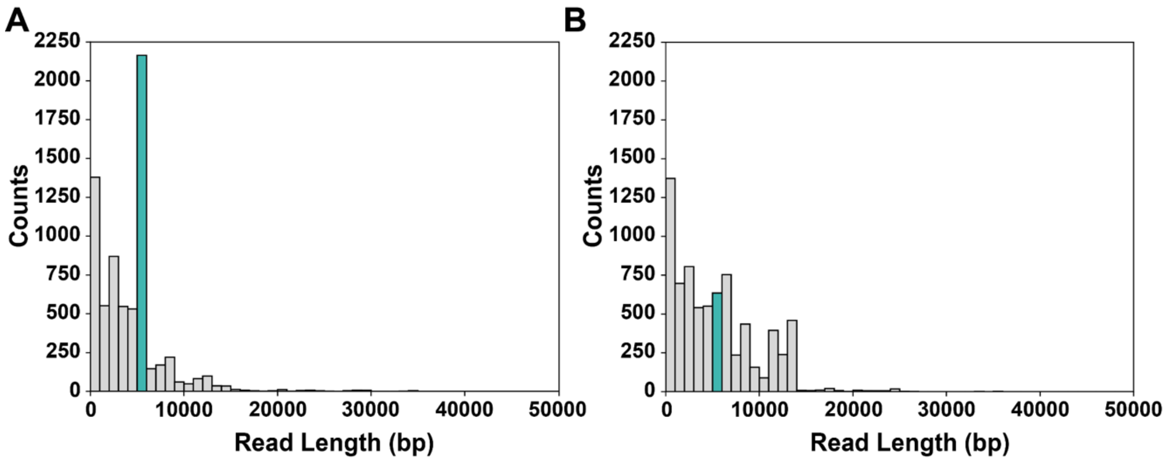


**Figure S18. Plasmid length distributions from NGS.** Data from plasmids isolated from wastewater samples that had been conjugated with the mobilizable plasmid (**A**) lacking CcdB and (**B**) containing CcdB. The bin corresponding to the conjugative plasmids is in teal, while all other wastewater plasmids are gray. Reads having lengths ≤1000 bp are not shown.

**Plasmid sequences with annotations.** The sequences of nine new plasmid are provided below using Genbank formatting. These include the self-mobilizable plasmid that transcribes RAM with barcode 1 (pDSZ114) and barcode 2 (pMAS046), the mobilizable plasmids that transcribes RAM with CcdB and barcode 1 (pMAS052, pMAS053) and barcode 2 (pMAS054), the plasmid that expresses CcdA (pMAS048), the plasmid that arose from recombination during conjugation (pESCccdB), and plasmids used for engineering (pMAS047, pMAS045).

**pDSZ114**

```
LOCUS       pDSZ114    61555 bp ds-DNA     circular     30-APR-2025
DEFINITION  Shuttle vector suitable for mobilizing plasmids to Anabaena
            (Nostoc) sp. strain PCC 7120 and other cyanobacterial strains..
SOURCE      synthetic DNA construct
  ORGANISM  synthetic DNA construct
COMMENT     Imported using the Genbank importer. File name:
            addgene-plasmid-70261-sequence-348706.gbk
FEATURES             Location/Qualifiers
     gene            587..1546
                     /label="trbB gene"
                     /ApEinfo_revcolor="#9eafd2"
                     /ApEinfo_fwdcolor="#9eafd2"
     primer          complement(764..776)
                     /label="AOS62B"
                     /note="sequence:
GACTGGAGTTCAGACGTGTGCTCTTCCGATCTGTTTCATGTGATCCGGATAAC"
                     /ApEinfo_revcolor="#b1ff67"
                     /ApEinfo_fwdcolor="#b1ff67"
     gene            1559..1996
                     /label="trbC gene"
                     /ApEinfo_revcolor="#b1ff67"
                     /ApEinfo_fwdcolor="#b1ff67"
     gene            1999..2310
                     /label="trbD gene"
                     /ApEinfo_revcolor="#9eafd2"
                     /ApEinfo_fwdcolor="#9eafd2"
     gene            2307..4865
                     /label="trbE gene"
                     /ApEinfo_revcolor="#84b0dc"
                     /ApEinfo_fwdcolor="#84b0dc"
     gene            4862..5620
                     /label="trbF gene"
                     /ApEinfo_revcolor="#faac61"
                     /ApEinfo_fwdcolor="#faac61"
     gene            5632..6525
                     /label="trbG gene"
                     /ApEinfo_revcolor="#c6c9d1"
                     /ApEinfo_fwdcolor="#c6c9d1"
     gene            6529..7011
                     /label="trbH gene"
                     /ApEinfo_revcolor="#ffef86"
                     /ApEinfo_fwdcolor="#ffef86"
     gene            7016..8407
                     /label="trbI gene"
                     /ApEinfo_revcolor="#b7e6d7"
```

```
                     /ApEinfo_fwdcolor="#b7e6d7"
     gene            8424..9200
                     /label="trbJ gene"
                     /ApEinfo_revcolor="#75c6a9"
                     /ApEinfo_fwdcolor="#75c6a9"
     gene            9212..9421
                     /label="trbK gene"
                     /ApEinfo_revcolor="#85dae9"
                     /ApEinfo_fwdcolor="#85dae9"
     gene            9428..11014
                     /label="trbL gene"
                     /ApEinfo_revcolor="#ff9ccd"
                     /ApEinfo_fwdcolor="#ff9ccd"
     gene            11038..11637
                     /label="trbM gene"
                     /ApEinfo_revcolor="#f58a5e"
                     /ApEinfo_fwdcolor="#f58a5e"
     gene            11653..12357
                     /label="trbN gene"
                     /ApEinfo_revcolor="#ff9ccd"
                     /ApEinfo_fwdcolor="#ff9ccd"
     gene            12388..12651
                     /label="trbO gene"
                     /ApEinfo_revcolor="#c7b0e3"
                     /ApEinfo_fwdcolor="#c7b0e3"
     gene            13873..14568
                     /label="fiwA gene"
                     /ApEinfo_revcolor="#d6b295"
                     /ApEinfo_fwdcolor="#d6b295"
     gene            14604..15266
                     /label="upf32.8 gene"
                     /ApEinfo_revcolor="#84b0dc"
                     /ApEinfo_fwdcolor="#84b0dc"
     operon          complement(15268..16770)
                     /label="operon"
                     /ApEinfo_revcolor="#d6b295"
                     /ApEinfo_fwdcolor="#d6b295"
     gene            complement(15290..15949)
                     /label="parA gene"
                     /ApEinfo_revcolor="#f8d3a9"
                     /ApEinfo_fwdcolor="#f8d3a9"
     gene            complement(15910..16443)
                     /label="parB gene"
                     /ApEinfo_revcolor="#9eafd2"
                     /ApEinfo_fwdcolor="#9eafd2"
     gene            complement(16440..16700)
                     /label="parC gene"
                     /ApEinfo_revcolor="#f8d3a9"
                     /ApEinfo_fwdcolor="#f8d3a9"
     gene            16884..17135
                     /label="parD gene"
                     /ApEinfo_revcolor="#84b0dc"
                     /ApEinfo_fwdcolor="#84b0dc"
     gene            17132..17443
                     /label="parE gene"
                     /ApEinfo_revcolor="#c7b0e3"
                     /ApEinfo_fwdcolor="#c7b0e3"
```

```
     gene            17521..17970
                     /label="upf35.8 gene"
                     /ApEinfo_revcolor="#d59687"
                     /ApEinfo_fwdcolor="#d59687"
     gene            19244..20041
                     /label="istB gene"
                     /ApEinfo_revcolor="#d59687"
                     /ApEinfo_fwdcolor="#d59687"
     misc_feature    20329..20360
                     /label="AphA-5'-IGR gRNA target"
                     /ApEinfo_revcolor="#b7e6d7"
                     /ApEinfo_fwdcolor="#b7e6d7"
     gene            21341..21369
                     /label="aphA gene"
                     /ApEinfo_revcolor="#f8d3a9"
                     /ApEinfo_fwdcolor="#f8d3a9"
     operon          complement(21341..30485)
                     /label="operon"
                     /ApEinfo_revcolor="#85dae9"
                     /ApEinfo_fwdcolor="#85dae9"
     gene            complement(21376..21666)
                     /label="traA gene"
                     /ApEinfo_revcolor="#ffef86"
                     /ApEinfo_fwdcolor="#ffef86"
     gene            complement(21674..22114)
                     /label="traB gene"
                     /ApEinfo_revcolor="#85dae9"
                     /ApEinfo_fwdcolor="#85dae9"
     gene            complement(22130..24370)
                     /label="traC2 gene"
                     /ApEinfo_revcolor="#b1ff67"
                     /ApEinfo_fwdcolor="#b1ff67"
     gene            complement(22130..25315)
                     /label="traC1 gene"
                     /ApEinfo_revcolor="#b4abac"
                     /ApEinfo_fwdcolor="#b4abac"
     gene            complement(25322..25585)
                     /label="traD gene"
                     /ApEinfo_revcolor="#f8d3a9"
                     /ApEinfo_fwdcolor="#f8d3a9"
     gene            complement(25591..27804)
                     /label="traE gene"
                     /ApEinfo_revcolor="#f8d3a9"
                     /ApEinfo_fwdcolor="#f8d3a9"
     gene            complement(27819..28352)
                     /label="traF gene"
                     /ApEinfo_revcolor="#faac61"
                     /ApEinfo_fwdcolor="#faac61"
     gene            complement(28349..30256)
                     /label="traG gene"
                     /ApEinfo_revcolor="#faac61"
                     /ApEinfo_fwdcolor="#faac61"
     gene            complement(30253..32451)
                     /label="traI gene"
                     /ApEinfo_revcolor="#b4abac"
                     /ApEinfo_fwdcolor="#b4abac"
     operon          complement(30253..33086)
```

```
                     /label="operon"
                     /ApEinfo_revcolor="#b1ff67"
                     /ApEinfo_fwdcolor="#b1ff67"
     gene            complement(30552..30911)
                     /label="traH gene"
                     /ApEinfo_revcolor="#b4abac"
                     /ApEinfo_fwdcolor="#b4abac"
     gene            complement(32448..32489)
                     /label="traX gene"
                     /ApEinfo_revcolor="#b7e6d7"
                     /ApEinfo_fwdcolor="#b7e6d7"
     gene            complement(32486..32857)
                     /label="traJ gene"
                     /ApEinfo_revcolor="#faac61"
                     /ApEinfo_fwdcolor="#faac61"
     CDS             complement(32486..32857)
                     /label="traJ"
                     /ApEinfo_revcolor="#84b0dc"
                     /ApEinfo_fwdcolor="#84b0dc"
                     /gene="traJ"
                     /product="oriT-recognizing protein"

/translation="MADETKPTRKGSPPIKVYCLPDERRAIEEKAAAAGMSLSAYLLAVGQGYKITGVVDYEHVRE
LARINGDLGRLGGLLKLWLTDDPRTARFGDATILALLAKIEEKQDELGKVMMGVVRPRAEP*"
     oriT            32890..32999
                     /label="incP origin of transfer"
                     /ApEinfo_revcolor="#84b0dc"
                     /ApEinfo_fwdcolor="#84b0dc"
     gene            33225..33629
                     /label="traK gene"
                     /ApEinfo_revcolor="#c6c9d1"
                     /ApEinfo_fwdcolor="#c6c9d1"
     gene            33629..34354
                     /label="traL gene"
                     /ApEinfo_revcolor="#84b0dc"
                     /ApEinfo_fwdcolor="#84b0dc"
     gene            34351..34788
                     /label="traM gene"
                     /ApEinfo_revcolor="#b7e6d7"
                     /ApEinfo_fwdcolor="#b7e6d7"
     gene            complement(36621..36657)
                     /label="krfA gene"
                     /ApEinfo_revcolor="#f8d3a9"
                     /ApEinfo_fwdcolor="#f8d3a9"
     gene            complement(37833..38360)
                     /label="korG gene"
                     /ApEinfo_revcolor="#f58a5e"
                     /ApEinfo_fwdcolor="#f58a5e"
     gene            complement(38944..40020)
                     /label="korB gene"
                     /ApEinfo_revcolor="#b7e6d7"
                     /ApEinfo_fwdcolor="#b7e6d7"
     gene            complement(40017..41111)
                     /label="incC gene"
                     /ApEinfo_revcolor="#85dae9"
                     /ApEinfo_fwdcolor="#85dae9"
     gene            complement(40793..41098)
```

```
                     /label="korA gene"
                     /ApEinfo_revcolor="#b4abac"
                     /ApEinfo_fwdcolor="#b4abac"
     gene            complement(41272..42225)
                     /label="klaC gene"
                     /ApEinfo_revcolor="#ffef86"
                     /ApEinfo_fwdcolor="#ffef86"
     operon          complement(41272..44222)
                     /label="operon"
                     /ApEinfo_revcolor="#84b0dc"
                     /ApEinfo_fwdcolor="#84b0dc"
     gene            complement(42222..43358)
                     /label="klaB gene"
                     /ApEinfo_revcolor="#faac61"
                     /ApEinfo_fwdcolor="#faac61"
     gene            complement(43376..44149)
                     /label="klaA gene"
                     /ApEinfo_revcolor="#85dae9"
                     /ApEinfo_fwdcolor="#85dae9"
     gene            complement(44272..44586)
                     /label="kleF gene"
                     /ApEinfo_revcolor="#b7e6d7"
                     /ApEinfo_fwdcolor="#b7e6d7"
     operon          complement(44272..45592)
                     /label="operon"
                     /ApEinfo_revcolor="#84b0dc"
                     /ApEinfo_fwdcolor="#84b0dc"
     gene            complement(44614..44937)
                     /label="kleE gene"
                     /ApEinfo_revcolor="#d6b295"
                     /ApEinfo_fwdcolor="#d6b295"
     gene            complement(45049..45267)
                     /label="kleD gene"
                     /ApEinfo_revcolor="#faac61"
                     /ApEinfo_fwdcolor="#faac61"
     gene            complement(45283..45513)
                     /label="kleC gene"
                     /ApEinfo_revcolor="#9eafd2"
                     /ApEinfo_fwdcolor="#9eafd2"
     gene            complement(45666..45881)
                     /label="kleB gene"
                     /ApEinfo_revcolor="#c6c9d1"
                     /ApEinfo_fwdcolor="#c6c9d1"
     operon          complement(45666..46247)
                     /label="operon"
                     /ApEinfo_revcolor="#f58a5e"
                     /ApEinfo_fwdcolor="#f58a5e"
     gene            complement(45930..46163)
                     /label="kleA gene"
                     /ApEinfo_revcolor="#c6c9d1"
                     /ApEinfo_fwdcolor="#c6c9d1"
     gene            complement(46283..46308)
                     /label="korC gene"
                     /ApEinfo_revcolor="#d59687"
                     /ApEinfo_fwdcolor="#d59687"
     gene            complement(46389..46646)
                     /label="korC gene"
```

```
                     /ApEinfo_revcolor="#b4abac"
                     /ApEinfo_fwdcolor="#b4abac"
     gene            complement(46669..47343)
                     /label="klcB gene"
                     /ApEinfo_revcolor="#b4abac"
                     /ApEinfo_fwdcolor="#b4abac"
     gene            complement(47495..48135)
                     /label="Disrupted bla gene 3'"
                     /ApEinfo_revcolor="#75c6a9"
                     /ApEinfo_fwdcolor="#75c6a9"
     primer          47939..47960
                     /label="DSZ20A"
                     /note="sequence: CGTTGTCAGAAGTAAGTTGGCC"
                     /ApEinfo_revcolor="#9eafd2"
                     /ApEinfo_fwdcolor="#9eafd2"
     misc_feature    48057..48088
                     /label="bla CDS g5"
                     /ApEinfo_revcolor="#c6c9d1"
                     /ApEinfo_fwdcolor="#c6c9d1"
     misc_feature    48131..48135
                     /label="Target site duplication"
                     /ApEinfo_revcolor="#ff9ccd"
                     /ApEinfo_fwdcolor="#ff9ccd"
     misc_feature    48136..48143
                     /label="END"
                     /ApEinfo_revcolor="#84b0dc"
                     /ApEinfo_fwdcolor="#84b0dc"
                     /note="ApEinfo_revcolor: #84b0dc"
     misc_feature    48136..48260
                     /label="VchINT R-end"
                     /ApEinfo_revcolor="#ffef86"
                     /ApEinfo_fwdcolor="#ffef86"
                     /note="ApEinfo_revcolor: #ffef86"
     misc_feature    48144..48163
                     /label="R1 tnsB binding site (VCHE45 Tn6677)"
                     /ApEinfo_revcolor="#c7b0e3"
                     /ApEinfo_fwdcolor="#c7b0e3"
     misc_feature    48144..48246
                     /label="RE (partial) (VCHE45 Tn6677)"
                     /ApEinfo_revcolor="#c6c9d1"
                     /ApEinfo_fwdcolor="#c6c9d1"
     misc_feature    48164..48183
                     /label="R2 tnsB binding site (VCHE45 Tn6677)"
                     /ApEinfo_revcolor="#c7b0e3"
                     /ApEinfo_fwdcolor="#c7b0e3"
     misc_feature    48187..48206
                     /label="R3 tnsB binding site (VCHE45 Tn6677)"
                     /ApEinfo_revcolor="#c7b0e3"
                     /ApEinfo_fwdcolor="#c7b0e3"
     primer          48246..48283
                     /label="DSZ79C"
                     /note="sequence:
aatgaagactatttcCCAATTATTGAAGGCCGCTAACG"
                     /ApEinfo_revcolor="#b4abac"
                     /ApEinfo_fwdcolor="#b4abac"
     primer          48257..48309
                     /label="DSZ51C"
```

```
                     /note="sequence:
tttcCCAATTATTGAAGGCCGCTAACGCGGCCTTTTTTTGTTTCTGGTCTGCC"
                     /ApEinfo_revcolor="#ff9ccd"
                     /ApEinfo_fwdcolor="#ff9ccd"
     terminator      48261..48309
                     /label="tVoigtS5, L3S3P22 47C>G"
                     /ApEinfo_revcolor="#c6c9d1"
                     /ApEinfo_fwdcolor="#c6c9d1"
     primer          complement(48261..48313)
                     /label="DSZ52C"
                     /note="sequence:
aggaGGCAGACCAGAAACAAAAAAAGGCCGCGTTAGCGGCCTTCAATAATTGG"
                     /ApEinfo_revcolor="#9eafd2"
                     /ApEinfo_fwdcolor="#9eafd2"
     primer          48310..48335
                     /label="DSZ76B"
                     /note="sequence: aatgaagacattcctCCCTGCAGGCTTCAACAAACAG"
                     /ApEinfo_revcolor="#84b0dc"
                     /ApEinfo_fwdcolor="#84b0dc"
     misc_feature    48315..48322
                     /label="SbfI"
                     /ApEinfo_revcolor="#85dae9"
                     /ApEinfo_fwdcolor="#85dae9"
     misc_feature    48327..48358
                     /label="CymR operator"
                     /ApEinfo_revcolor="#ffef86"
                     /ApEinfo_fwdcolor="#ffef86"
     promoter        48327..48416
                     /label="P.CymRC (Marionette)"
                     /ApEinfo_revcolor="#ffef86"
                     /ApEinfo_fwdcolor="#ffef86"
     misc_feature    48350..48355
                     /label=-35
                     /ApEinfo_revcolor="#ffef86"
                     /ApEinfo_fwdcolor="#ffef86"
     misc_feature    48373..48378
                     /label=-10
                     /ApEinfo_revcolor="#ffef86"
                     /ApEinfo_fwdcolor="#ffef86"
     misc_feature    48385..48385
                     /label="TSS"
                     /ApEinfo_revcolor="#ffef86"
                     /ApEinfo_fwdcolor="#ffef86"
     misc_feature    48385..48416
                     /label="CymR operator"
                     /ApEinfo_revcolor="#ffef86"
                     /ApEinfo_fwdcolor="#ffef86"
     misc_feature    48421..48476
                     /label="U64 RNA Guide"
                     /ApEinfo_revcolor="#faac61"
                     /ApEinfo_fwdcolor="#faac61"
     misc_feature    48421..49381
                     /label="U64 Ribozyme transcript"
                     /ApEinfo_revcolor="#d59687"
                     /ApEinfo_fwdcolor="#d59687"
     misc_feature    48471..48476
                     /label="IGS"
```

```
                    /ApEinfo_revcolor="#c7b0e3"
                    /ApEinfo_fwdcolor="#c7b0e3"
     misc_feature   48477..48863
                    /label="Group I Intron Ribozyme (from p-OiRS3GG)"
                    /ApEinfo_revcolor="#b4abac"
                    /ApEinfo_fwdcolor="#b4abac"
     misc_feature   48864..48879
                    /label="for cutadapt remove after splice (NGS
Analysis)"
                    /ApEinfo_revcolor="#b4abac"
                    /ApEinfo_fwdcolor="#b4abac"
     misc_feature   48864..49381
                    /label="Barcode (sfGFP_2)"
                    /ApEinfo_revcolor="#84b0dc"
                    /ApEinfo_fwdcolor="#84b0dc"
     misc_feature   48880..48884
                    /label="barcode 1"
                    /ApEinfo_revcolor="#b7e6d7"
                    /ApEinfo_fwdcolor="#b7e6d7"
     misc_feature   48885..48905
                    /label="AOS62B binds here"
                    /ApEinfo_revcolor="#faac61"
                    /ApEinfo_fwdcolor="#faac61"
     misc_feature   49294..49299
                    /label="BamHI-BglII Scar"
                    /ApEinfo_revcolor="#ac1dff"
                    /ApEinfo_fwdcolor="#ac1dff"
     terminator     49386..49438
                    /label="tVoigtS4 L3S3P21 51C>G"
                    /ApEinfo_revcolor="#f58a5e"
                    /ApEinfo_fwdcolor="#f58a5e"
     primer         complement(49425..49451)
                    /label="DSZ40A"
                    /note="sequence: aatgaagacattaagCGCAGCGGCAGACCAGAAAC"
                    /ApEinfo_revcolor="#85dae9"
                    /ApEinfo_fwdcolor="#85dae9"
     primer         complement(49437..49469)
                    /label="DSZ80C"
                    /note="sequence: aatgaagactaaggaCTGCAGTAAGCGCAGCGG"
                    /ApEinfo_revcolor="#d6b295"
                    /ApEinfo_fwdcolor="#d6b295"
     misc_feature   complement(49445..49589)
                    /label="LE (partial) (VCHE45 Tn6677)"
                    /ApEinfo_revcolor="#b4abac"
                    /ApEinfo_fwdcolor="#b4abac"
     misc_feature   complement(49445..49589)
                    /label="VchINT L-end"
                    /ApEinfo_revcolor="#ffef86"
                    /ApEinfo_fwdcolor="#ffef86"
                    /note="ApEinfo_revcolor: #ffef86"
     misc_feature   complement(49482..49501)
                    /label="L3 tnsB binding site (VCHE45 Tn6677)"
                    /ApEinfo_revcolor="#f58a5e"
                    /ApEinfo_fwdcolor="#f58a5e"
     misc_feature   complement(49508..49527)
                    /label="L2 tnsB binding site (VCHE45 Tn6677)"
                    /ApEinfo_revcolor="#f58a5e"
```

```
                     /ApEinfo_fwdcolor="#f58a5e"
     misc_feature    complement(49562..49581)
                     /label="L1 tnsB binding site (VCHE45 Tn6677)"
                     /ApEinfo_revcolor="#f58a5e"
                     /ApEinfo_fwdcolor="#f58a5e"
     misc_feature    complement(49582..49589)
                     /label="END"
                     /ApEinfo_revcolor="#84b0dc"
                     /ApEinfo_fwdcolor="#84b0dc"
                     /note="ApEinfo_revcolor: #84b0dc"
     CDS             complement(49584..49814)
                     /label="Disrupted bla 5' "
                     /ApEinfo_revcolor="#84b0dc"
                     /ApEinfo_fwdcolor="#84b0dc"

/translation="MSIQHFRVALIPFFAAFCLPVFAHPETLVKVKDAEDKLGARVGYIELDLNSGKILESFRPEE
RFPMMSTFKVLLCC*"
     misc_feature    49590..49594
                     /label="Target site dupication"
                     /ApEinfo_revcolor="#ff9ccd"
                     /ApEinfo_fwdcolor="#ff9ccd"
     gene            complement(49595..49814)
                     /label="Disrupted bla 5'"
                     /ApEinfo_revcolor="#c7b0e3"
                     /ApEinfo_fwdcolor="#c7b0e3"
     misc_feature    complement(49709..49740)
                     /label="bla CDS g4"
                     /ApEinfo_revcolor="#75c6a9"
                     /ApEinfo_fwdcolor="#75c6a9"
     primer          complement(49788..49807)
                     /label="DSZ19A"
                     /note="sequence: TTCAACATTTCCGTGTCGCC"
                     /ApEinfo_revcolor="#d59687"
                     /ApEinfo_fwdcolor="#d59687"
     promoter        complement(49815..49919)
                     /label="AmpR promoter"
                     /ApEinfo_revcolor="#f58a5e"
                     /ApEinfo_fwdcolor="#f58a5e"
     gene            complement(49997..50554)
                     /label="tnpR gene"
                     /ApEinfo_revcolor="#b4abac"
                     /ApEinfo_fwdcolor="#b4abac"
     gene            complement(53757..54467)
                     /label="klcB gene"
                     /ApEinfo_revcolor="#c6c9d1"
                     /ApEinfo_fwdcolor="#c6c9d1"
     gene            complement(54513..54953)
                     /label="klcA gene"
                     /ApEinfo_revcolor="#ff9ccd"
                     /ApEinfo_fwdcolor="#ff9ccd"
     rep_origin      55425..56136
                     /label="oriV"
                     /ApEinfo_revcolor="#b7e6d7"
                     /ApEinfo_fwdcolor="#b7e6d7"
                     /note="incP origin of replication"
     gene            complement(56610..57260)
                     /label="tetR gene"
```

```
                     /ApEinfo_revcolor="#ff9ccd"
                     /ApEinfo_fwdcolor="#ff9ccd"
     CDS             complement(56610..57260)
                     /label="TetR"
                     /ApEinfo_revcolor="#84b0dc"
                     /ApEinfo_fwdcolor="#84b0dc"
                     /gene="tetR"
                     /product="tetracycline resistance regulatory protein"

/translation="MTKLQPNTVIRAALDLLNEVGVDGLTTRKLAERLGVQQPALYWHFRNKRALLDALAEAMLAE
NHTHSVPRADDDWRSFLIGNARSFRQALLAYRDGARIHAGTRPGAPQMETADAQLRFLCEAGFSAGDAVNALMTIS
YFTVGAVLEEQAGDSDAGERGGTVEQAPLSPLLRAAIDAFDEAGPDAAFEQGLAVIVDGLAKRRLVVRNVEGPRKG
DD*"
     gene            57366..58565
                     /label="tetA gene"
                     /ApEinfo_revcolor="#75c6a9"
                     /ApEinfo_fwdcolor="#75c6a9"
     CDS             57366..58565
                     /label="TcR"
                     /ApEinfo_revcolor="#84b0dc"
                     /ApEinfo_fwdcolor="#84b0dc"
                     /note="confers resistance to tetracycline"
                     /gene="tetA"
                     /product="tetracycline efflux protein"

/translation="VKPNIPLIVILSTVALDAVGIGLIMPVLPGLLRDLVHSNDVTAHYGILLALYALVQFACAPV
LGALSDRFGRRPILLVSLAGATVDYAIMATAPFLWVLYIGRIVAGITGATGAVAGAYIADITDGDERARHFGFMSA
CFGFGMVAGPVLGGLMGGFSPHAPFFAAAALNGLNFLTGCFLLPESHKGERRPLRREALNPLASFRWARGMTVVAA
LMAVFFIMQLVGQVPAALWVIFGEDRFHWDATTIGISLAAFGILHSLAQAMITGPVAARLGERRALMLGMIADGTG
YILLAFATRGWMAFPIMVLLASGGIGMPALQAMLSRQVDEERQGQLQGSLAALTSLTSIVGPLLFTAIYAASITTW
NGWAWIAGAALYLLCLPALRRGLWSGAGQRADR*"
     primer          58658..58677
                     /label="DSZ23A"
                     /note="sequence: CAGATACTCCCGATCACGAG"
                     /ApEinfo_revcolor="#9eafd2"
                     /ApEinfo_fwdcolor="#9eafd2"
     misc_feature    58743..58774
                     /label="TetA-3'-IGR gRNA target"
                     /ApEinfo_revcolor="#b4abac"
                     /ApEinfo_fwdcolor="#b4abac"
     primer          complement(59055..59073)
                     /label="DSZ24A"
                     /note="sequence: CCAAACGCAGCGCTAGATC"
                     /ApEinfo_revcolor="#ff9ccd"
                     /ApEinfo_fwdcolor="#ff9ccd"
     gene            complement(59838..60986)
                     /label="trfA gene"
                     /ApEinfo_revcolor="#9eafd2"
                     /ApEinfo_fwdcolor="#9eafd2"
     CDS             complement(59838..60986)
                     /label="trfA"
                     /ApEinfo_revcolor="#84b0dc"
                     /ApEinfo_fwdcolor="#84b0dc"
                     /product="trans-acting replication protein that binds
to and activates oriV"

/translation="MNRTFDRKAYRQELIDAGFSAEDAETIASRTVMRAPRETFQSVGSMVQQATAKIERDSVQLA
```

```
PPALPAPSAAVERSRRLEQEAAGLAKSMTIDTRGTMTTKKRKTAGEDLAKQVSEAKQAALLKHTKQQIKEMQLSLF
DIAPWPDTMRAMPNDTARSALFTTRNKKIPREALQNKVIFHVNKDVKITYTGVELRADDDELVWQQVLEYAKRTPI
GEPITFTFYELCQDLGWSINGRYYTKAEECLSRLQATAMGFTSDRVGHLESVSLLHRFRVLDRGKKTSRCQVLIDE
EIVVLFAGDHYTKFIWEKYRKLSPTARRMFDYFSSHREPYPLKLETFRLMCGSDSTRVKKWREQVGEACEELRGSG
LVEHAWVNDDLVHCKR*"
     gene            complement(61035..61385)
                     /label="ssb gene"
                     /ApEinfo_revcolor="#ffef86"
                     /ApEinfo_fwdcolor="#ffef86"
     gene            61506..316
                     /label="trbA gene"
                     /ApEinfo_revcolor="#c6c9d1"
                     /ApEinfo_fwdcolor="#c6c9d1"
ORIGIN
        1 gggcatgacg aaacatgagc tgtcggagag ggcaggggtt tcaatttcgt ttttatcaga
       61 cttaaccaac ggtaaggcca acccctcgtt gaaggtgatg gaggccattg ccgacgccct
      121 ggaaactccc ctacctcttc tcctggagtc caccgacctt gaccgcgagg cactcgcgga
      181 gattgcgggt catcctttca agagcagcgt gccgcccgga tacgaacgca tcagtgtggt
      241 tttgccgtca cataaggcgt ttatcgtaaa gaaatggggc gacgacaccc gaaaaaagct
      301 gcgtggaagg ctctgacgcc aagggttagg gcttgcactt ccttctttag ccgctaaaac
      361 ggccccttct ctgcgggccg tcggctcgcg catcatatcg acatcctcaa cggaagccgt
      421 gccgcgaatg gcatcgggcg ggtgcgcttt gacagttgtt ttctatcaga acccctacgt
      481 cgtgcggttc gattagctgt ttgtcttgca ggctaaacac tttcggtata tcgtttgcct
      541 gtgcgataat gttgctaatg atttgttgcg taggggttac tgaaaagtga gcgggaaaga
      601 agagtttcag accatcaagg agcgggccaa gcgcaagctg gaacgcgaca tgggtgcgga
      661 cctgttggcc gcgctcaacg acccgaaaac cgttgaagtc atgctcaacg cggacggcaa
      721 ggtgtggcac gaacgccttg gcgagccgat gcggtacatc tgcgacatgc ggcccagcca
      781 gtcgcaggcg attatagaaa cggtggccgg attccacggc aaagaggtca cgcggcattc
      841 gcccatcctg gaaggcgagt tcccctcgga tggcagccgc tttgccggcc aattgccgcc
      901 ggtcgtggcc gcgccaacct ttgcgatccg caagcgcgcg gtcgccatct tcacgctgga
      961 acagtacgtc gaggcgggca tcatgacccg cgagcaatac gaggtcatta aaagcgccgt
     1021 cgcggcgcat cgaaacatcc tcgtcattgg cggtactggc tcgggcaaga ccacgctcgt
     1081 caacgcgatc atcaatgaaa tggtcgcctt caacccgtct gagcgcgtcg tcatcatcga
     1141 ggacaccggc gaaatccagt gcgccgcaga gaacgccgtc caataccaca ccagcatcga
     1201 cgtctcgatg acgctgctgc tcaagacaac gctgcgtatg cgccccgacc gcatcctggt
     1261 cggtgaggta cgtggccccg aagcccttga tctgttgatg gcctggaaca ccgggcatga
     1321 aggaggtgcc gccaccctgc acgcaaacaa ccccaaagcg ggcctgagcc ggctcgccat
     1381 gcttatcagc atgcacccgg attcaccgaa acccattgag ccgctgattg gcgaggcggt
     1441 tcatgtggtc gtccatatcg ccaggacccc tagcggccgt cgagtgcaag aaattctcga
     1501 agttcttggt tacgagaacg gccagtacat caccaaaacc ctgtaaggag tatttccaat
     1561 gacaacggct gttccgttcc gtctgaccat gaatcgcggc attttgttct accttgccgt
     1621 gttcttcgtt ctcgctctcg cgttatccgc gcatccggcg atggcctcgg aaggcaccgg
     1681 cggcagcttg ccatatgaga gctggctgac gaacctgcgc aactccgtaa ccggcccggt
     1741 ggccttcgcg ctgtccatca tcggcatcgt cgtcgccggc ggcgtgctga tcttcggcgg
     1801 cgaactcaac gccttcttcc gaaccctgat cttcctggtt ctggtgatgg cgctgctggt
     1861 cggcgcgcag aacgtgatga gcaccttctt cggtcgtggt gccgaaatcg cggccctcgg
     1921 caacggggcg ctgcaccagg tgcaagtcgc ggcggcggat gccgtgcgtg cggtagcggc
     1981 tggacggctc gcctaatcat ggctctgcgc acgatcccca tccgtcgcgc aggcaaccga
     2041 gaaaacctgt tcatgggtgg tgatcgtgaa ctggtgatgt tctcgggcct gatggcgttt
     2101 gcgctgattt tcagcgccca agagctgcgg gccaccgtgg tcggtctgat cctgtggttc
     2161 ggggcgctct atgcgttccg aatcatggcg aaggccgatc cgaagatgcg gttcgtgtac
     2221 ctgcgtcacc gccggtacaa gccgtattac ccggcccgct cgaccccgtt ccgcgagaac
     2281 accaatagcc aagggaagca ataccgatga tccaagcaat tgcgattgca atcgcgggcc
     2341 tcggcgcgct tctgttgttc atcctctttg cccgcatccg cgcggtcgat gccgaactga
     2401 aactgaaaaa gcatcgttcc aaggacgccg gcctggccga tctgctcaac tacgccgctg
     2461 tcgtcgatga cggcgtaatc gtgggcaaga acggcagctt tatggctgcc tggctgtaca
     2521 agggcgatga caacgcaagc agcaccgacc agcagcgcga agtagtgtcc gcccgcatca
```

2581 accaggccct cgcgggcctg ggaagtgggt ggatgatcca tgtggacgcc gtgcggcgtc
2641 ctgctccgaa ctacgcggag cggggcctgt cggcgttccc tgaccgtctg acggcagcga
2701 ttgaagaaga gcgccggcgg catttcgaga gcctgggaac gatgtacgag ggctatttcg
2761 tcctcacctt gacctggttc ccgccgctgc tcgcccagcg caagttcgtc gagctgatgt
2821 ttgacgacga cgcgaccgca ccggatcgca aggcgcgcac gcggggcctc atcgaccaat
2881 tcaagcgtga cgtgcgcagc atcgagtcgc gcctgtcgtc ggccgtgtcg ctcactcgct
2941 tgaaggggca caagatcgtc aacgaggacg gcacgaccgt cacgcatgac gacttcctgc
3001 gctggctgca attctgcgtg acgggcctgc accatccggt gcagctcccc agcaacccga
3061 tgtacctgga cgccctggtc ggcggacagg aaatgtgggg cggggtagtg cccaaggtcg
3121 gccgcaagtt cgtccaggtg gtcgctctcg aaggcttccc cttggagtcc tatcccggca
3181 tcctgacggc gctcggcgag ctgccctgcg agtatcggtg gtcgagccgg ttcatcttca
3241 tggaccagca cgaagccgtg aagcacctcg acaagttccg caagaagtgg cggcagaaga
3301 ttcgcggctt cttcgaccag gtgttcaaca cgaacaccgg cccggtcgat caggacgcgc
3361 tttcgatggt ggccgatgct gaggcggcca ttgccgaagt caacagcggc atcgtggccg
3421 tgggctacta caccagcgtc gtcgtgctga tggatgagga ccgcacgcgc ctggaagctg
3481 cggcccgcga tgttgaaaag gccgtcaacc ggttgggctt tgccgcgcgc atcgagtcca
3541 tcaacaccct ggacgccttc cttggtagtt tgccgggcca cggcgtggaa aacgtccgcc
3601 ggccgctcat caacacgatg aacctggccg acctgctgcc gaccagcacc atctggaccg
3661 gcaacgcgaa cgcgccatgc ccgatgtacc cgccgctgtc gccggcgctc atgcactgcg
3721 tcacgcaagg atcaacgccg ttccggctga acctgcacgt gcgcgacctc ggccacacct
3781 ttatgttcgg gccgaccggc gcaggtaaat cgacgcacct ggcgatcctc gccgcgcagc
3841 tccgtcgcta tgccggcatg tcgatcttcg cctttgacaa gggcatgtcg atgtacccgc
3901 tggccgccgg catccgtgcg gccacgaagg gcaccagcgg cctgcacttc accgtggcgg
3961 ccgacgacga acgcctggcg ttctgcccgt tgcagttcct gagcaccaag ggcgaccgtg
4021 cttgggcgat ggagtggatc gacaccatcc tggcgttgaa cggcgtcgaa acgacccggg
4081 cccagcgcaa cgaaatcggc aacgcgatca tgagcatgca cgccagcggc gcgcgcacgc
4141 tctccgagtt cagcgtgacg attcaggatg aggcgatccg cgaggcgatc cgccagtaca
4201 ccgtcgatgg cgcaatgggc catctgctcg acgccgaaga ggacggcttg gcgctgtccg
4261 actttacagt gttcgagatc gaagagctga tgaacctcgg cgagaaattc gccctgcctg
4321 tgttgctcta cctgttccgc cgtatcgagc gcgccctgac gggccagccg gccgtcatca
4381 tcctggacga agcctggttg atgctcggcc acccggcatt ccgcgcgaag atcagggaat
4441 ggctcaaggt gctgcgtaag gccaactgcc ttgtgctgat ggcaacgcag agcctgtccg
4501 acgccgccaa cagcggcatc ctggacgtga tcgtggaatc gaccgcgacc aagatttcc
4561 tgccgaatat ttacgccagg gatgaggaca cggcggccct gtaccgccgc atgggcctga
4621 acgctcgcca gatcgagatt ctggcccagg ccgttcccaa gcgtcagtac tactacgtgt
4681 cggaaaacgg ccgccgtctc tacgacctgg cacttggccc gctcgcgctc gcgttcgtcg
4741 gcgcatccga caaggaatcc gtcgccatca tcaagaacct ggaagccaag ttcggcgacc
4801 agtgggtgga tgaatggctg cgtggccggg gcctcgccct tgatgaatac ctggaggcag
4861 catgagtttt gcagacacga tcaagggctt gatcttcaag aagaagcccg caacggccgc
4921 agcagcggcg acgccggccg cgaccggccc gcaaaccgac aacccgtacc tgacggcgcg
4981 gcgcacctgg aacgaccacg ttggttccgt tgtgtcgcaa aagcagacct ggcaggttgt
5041 cggcatcctt tcgctgatga tcgtcctcgc ggcggtcggc ggcatcatcc acatcggcag
5101 ccagtcgaag ttcgtgccct atgtctacga ggtagacaag ctcgggcaga cggccgccgt
5161 ggggccgatg accagggcgt cgaaagccga tccgcgtgtc attcacgcct cggtggctga
5221 gttcgtcggc gatgctcgcc tggtgacgcc ggacgtagct ttgcagcgca aggccgtcta
5281 ccgcctctat gccaagctcg ggccgaatga cccggccacc gccaagatga acgaatggct
5341 caacggcacc gccgacgcca gcccgttcgc tcgcgcggcc gtcgaaacgg tcagcaccga
5401 aatcacttcc gtaatcccgc agacgcccga cacctggcag gtcgattggg tcgagacgac
5461 gcgcgacagg caaggcgtgg tgaaaggcca gcccgtgcgc atgcgggcct tggtgacggt
5521 ctacgtcgtc gagccgacgg cggacaccaa ggaagaacaa ctgcgaaaca acccggccgg
5581 gatctacgtc cgggacttct cctggtcgag acttctgtga ggcactgaat tatgaaaaag
5641 gaactgtttg ctttggtcct ggccgcgtcc gttagcgtgc ctgcatttgc cgccgatccc
5701 ggcgcggacc tgactgacct ctatttttcc ggcaagaacc cggagctgac cgcgcaagag
5761 cgggcggcca tcgccatcgc caagaagtgg gaggcgggta ccgccggcat gcggccggtg
5821 gccggccccg gtggttcggt gcgcttcctg ttcggcgcgc agcagccgag catcgtatgc
5881 gccgtgctgc aagtgtgcga cgtggccctg caacccggcg agcaagtcaa ctcgatcaac
5941 ctgggcgaca ccgcccgttg gacggtcgag ccggccatta ccggcagcgg cgcgaacgaa

6001 acccagcacc tcatcatcaa gccgatggat gtgggcctgg aaaccagcct ggtcgtgacc
6061 acggaccgcc gcagctacca catgcgcctg cgctcgcatc gcacgcagta catgccgcag
6121 gtgtcgttca cctacccgga agatgccctt gcgaagtggg acgccatcaa gaaccgcgaa
6181 cagcgggatc gcgtcgagaa aaccattccg cagaccggcg agtacctggg caacctgagc
6241 ttcaactact ccgtcagcgg gtccacgtcg tggaagccgg tgcgcgtcta caacgacggc
6301 aagaaaacca tcatccagat gccgcactcg atggaacaga ccgaagcgcc gacgctcctg
6361 gtcgttcgca gggagggcgg cctgttctcc gacgatgaaa cggtgatggt caactaccgg
6421 gtccagggcg accgctacat cgtcgatacg attttcgaca aggccatcct catcgcgggc
6481 gtgggcagca gccaggaccg cgtgaccatt tcaaggggga actaaaccat gcgtaagatt
6541 ctgaccgtca tcgcactcgc ggccacgttg gccggctgcg cgacctccaa gtacggcagc
6601 ttcgtccagg acgcgccggc cgcctacaac cagaccattg cgaccgacgc ggtgaagcag
6661 ctcgtcaagc tctacccgcc ggcgcaaacc aagctggaat tgcagcaggc tacgcccgat
6721 ccgttcggca ttgccctggt cactgacctt cgcgcccagg gctatgctgt catggagtac
6781 aagcccgacg gcaacgcggc cgcagctccg gctgctgcgt cctcggccgc tgcgaagccg
6841 gcaacgccgc aagcccaggg cggctatccg ctgcgctacg tgctggacca attcagcgac
6901 agcaacctgt atcgcctgac cgtcatggtc ggctctcaat cgctcacgcg cgcctacctc
6961 gcccaaaaca acacgatggt cccggccggc gcatgggttc ggaaggagta agccaatgag
7021 cgaagatcaa atggcaccgg acgcatcgcc agatgcggtc aagccgaaaa gcggggttcg
7081 ccgcgtcaac aacatgccga tgtacctcat cggcggtgtg ctcggcatct tcctgctggt
7141 gatggccctg gttgctgcgg atcgcgctgc gcagcagaac cagccgggag ctgcgaaggc
7201 tgagaaggcc ggcagcacca gcatgtttgc cgacgaaatt gccggcaaac agcaggacgg
7261 catcatcaag gccaagccgc tggagattcc gccggaacaa accgcccagc aaccgacgac
7321 ggagctgacg ccagccccgg cgcagggaac gactatcacg gtcgcacggc ccgagaacct
7381 ggaccagccc ccgacgccgc cgcagggtgc gcgcaacgag gacctggacc gcatccgcat
7441 ggcgaagttg cagatgctgg aagaggcgat caaggccaag acgacggtgc gcatcgacgc
7501 gccgcgcagc cagggcagcg ccggcggcgg tgctccgcag ggccgcgagg aaacccttgc
7561 gcgcatccag gagctgcgtc ggcaggctga gaacgcccgc gccaccgatc cgaccgccgc
7621 ctatcaggcc gcgcttgcgc aggctcgcac gatgggcggc gcggcagggg gtggcggtat
7681 gggcggctcg ggtgcgccga ccctcgtgca gacctcgaac cgcagtggtg gcggcgctgg
7741 ctatgggtcg ttcgacaacc gcagcgaggg cgaccgttgg cggctcgact cccagccgga
7801 agcacctgca acgccctatg tgctgcgcgc tggcttcgtc gttccggcta cgcttatctc
7861 gggcatcaac tccgatctgc caggccaaat catggcccag gtatcgcagt cggtgtacga
7921 cacggcgacc ggcaagcaca tgctcatccc ccaaggctcg cgcctggtgg gcagctactc
7981 gaacgatgtg gcctacgggc agaagcgcgt tctggtggca tggcagcgca tcatcttccc
8041 cgacggcaag gcaatggaca ttggggccat gccgggcggc gatagcgctg ggtatgcagg
8101 cttcaacgac aaggtcaaca accactactt ccgcaccttc gcatcggcat tcctcatgtc
8161 gggcgtcgtt gcgggcatca gcttgagtca ggaccgtggc aacagcaaca gcggttacgg
8221 acgacaagac gcgggttccg cgatgagtga agcgttgggt caacagctcg gccaagtaac
8281 ggcgcagatg atcgccaaaa acttgaatat cgcgccgacg ctggaaatcc gtccgggcta
8341 tcgcttcaac gtcattgtca cgaaagacat gacgttttct aagccctacc aggcgtttga
8401 ctattaactc caaggagtaa cttatgaaga agctcgctaa gaatgtttta gccgctaaag
8461 tagctctggt gctggccctc tcggtcggca ccttggcggt cacgcctgcg caagcgggca
8521 ttccggtcat cgacggcacc aacctgtcac aaaccactgt caccgcgatt cagcaggttg
8581 cgcaggtcca gaagcaaatc gaggaatacc ggacgcagtt gcagcagtac gaaaacatgc
8641 tgcaaaacac ggtggccccg gccgcctacg tgtgggacca ggcgcagtcc accatcaacg
8701 gcctgatgag cgccgttgat accctgaact actacaagaa ccaggcgggc agcatcgacg
8761 cttacctggg caagttcaag gacgtgtcct actacaaggg gtcgccgtgc ttctccctgt
8821 cgggctgctc ggaaagcgag cgcaaggcga tggaagagaa ccgccgcctg gcgtccgaat
8881 cgcagaaaaa ggccaacgat gcgctgttcc gtggcctcga tcagcagcag agcaacctca
8941 agtccgacgc cgccacgctg gagcaattga agggcaaggc gacgacggcg cagggccagt
9001 tggaagccct cggctacgcc aaccagttcg ccagccagca ggccaaccag ctcatgcaaa
9061 tccgtggcct tctgcttgcg cagcagaacg ccatcgccac gcagatgcag gcccagcagg
9121 accggcaggc ccagcaggac gctgcgggcg cgaagctgcg cgagggttcg taccgcgcaa
9181 gcccgtctaa gacctggtga ggggaggcgc gatgaagaaa tccaacttca tcgcagttgc
9241 cgcgctggcc gccgtcatgg cggccagcct ggcaggctgc gacaacaagc ccgacaccga
9301 caagctgacc tgcgccgatc tgccgaaggt cacggatgcc gctcaacgcg cggagctgtt
9361 gaagaagtgc ccgcgcggag aaccgggagg cttcaagccc agcgaaaaga aagagtggtg

9421 atgacgtatg aaaatccaga ctagagctgc cgcgctcgcg gtcctgatgc tggccttgat
9481 gccggtagcg gcatacgccc aaatcgacaa ttcgggcatc ctcgacaacg tattgcagcg
9541 ctaccagaac gccgcgagcg gctgggccac tgtcgtccag aacgccgcaa cctggctgtt
9601 ctggaccttg accgtgatta gcatggtctg gaccttcggc atgatggcac tgcgcaaggc
9661 cgacattggc gagttcttcg ccgagttcgt gcggttcacc atcttcaccg gcttcttctg
9721 gtggctgctg accaacggcc cgaatttcgc gtcgtccatc tatgcgtccc tgcggcagat
9781 tgcaggccag gcaacggggt tggggcaggg gctttcgccg tccggcatcg tcgatgttgg
9841 cttcgagatt ttcttcaagg tgatggacga aacctcgtac tggtcgccgg tcgatagctt
9901 cgtcggtgcc tcgttggcgg ccgccatcct ctgcatcctg gccctggtcg gcgtgaatat
9961 gcttctgctc ctggcgtccg gatggattct tgcctacggc ggtgtgttct tcctgggctt
10021 cggcggctcg cgctggacct cggacatggc gatcaactac tacaagaccg tcctcggggt
10081 cgccgcgcag ctcttcgcaa tggtgctgct cgtaggcatc ggcaagacct tcctcgatga
10141 ctactacagc cgcatgagcg aaggcatcaa cttcaaggaa cttggagtga tgctgatcgt
10201 cggcctgatc ctgctcgttc tggtcaacaa ggtgccgcag ctcatcgccg gcatcatcac
10261 cggcgcgagc gtcggcggtg ctggtatcgg ccagttcggc gctggcacgc tcgtcggtgc
10321 ggccgcgacg gccggcgcgg caatcgcaac tggcggcgca tctatcgcgg ccggcgctgc
10381 ggcggcggcc ggtggcgcgc aggccatcat ggcggccgcg tcgaaggcca gcgataacgt
10441 ctctgccggc actgacattc tgtcgagcat gatgggcggc ggcggtggcg gcggcggtgg
10501 tagcgccggc accagcggcg gcgacggcgg cggctcgggt ggcggcggtg gctcgggcgg
10561 cggtgaaacc ccgatggcct cggccgccgg cgacaacagc agcggcgcac gcggcggcag
10621 ttcgggcggc ggctcgggtg gtggccgttc gtctggcggt atcggtgcca cggcggccaa
10681 gggcggccgg atcgcggccg ataccgtcgc caacctggcg aaaggtgccg gctcgattgc
10741 caaggccaag gccggcgaaa tgcgcgcatc ggcccaggaa cgcatcggcg ataccgtagg
10801 cggcaagatc gcgcaggcaa ttcgcggcgc gggtgcggcg gcgcagaccg ctgcaaccgt
10861 cgccgatagc aacagccagg cgcaggaaca acctgcaccg gcacccgcac cgtcgttcga
10921 cgacaacagc ctttccgcaa gcaacaacag ggaagcggcc gccgacgcgg attccgaagt
10981 ggcgagcttc gtcaacaagc ccgcccaatc ctgaaacgac tcttaggagc tacgaccatg
11041 caactgaaaa aagcgttctc gtcggccgcc ctggtggtgg ccttgggcct cggcgcaact
11101 ggctcggcca gcgcgcaaga cgtgctgacg ggcgataccc gcctggcctg cgaggccatt
11161 ctgtgcctgt ccacgggcag ccggcccagc gagtgcagcc cgtcgctctc gcggtacttc
11221 ggcatccaca agcgcaagct gtcggacacg ctcaaggcgc ggctgaactt cctcaacctc
11281 tgcccggtat cgaaccagac gccggaaatg cagacgctcg tttcctcgat ttcgcgcggg
11341 gccgggcgct gcgatgcgtc ctcgctgaac tccgtgctgc gtgagtggcg gagctgggac
11401 gaccagttct acatcggcaa ccgcctgccg gactactgcg cggcctacac cggccatgcc
11461 tataccgact tcaacacgac cgcgccgcgc tacgtcggca cgccggaaga gggcggctat
11521 tggatcgagg cggccgacta cgaccgcgcg ctcaaggagt acgaggcgaa gctgaaagag
11581 cggcagcagc agtacggtcg ctatggcagc gacgcctacc gtcggttcga gcggtaaggg
11641 gaggggatag cgatgccgtt tgccaagctg ctggcacgga acgctctgcc ggtggtcgcc
11701 ctggtggcgg ccactggctt cggtgcggcg gatgcgaccg ccgcacggct cttccccgat
11761 ctgtcggaac agatggaaga gcgcgttgtg tgctcggtgt ctgcggccgc gaagtacgag
11821 attccggcca acattcttct cgccattcgg gaaaaggagg gcggcaagcc gggccagtgg
11881 gtcaagaaca ccaatggcac ctatgacgtg ggcgagctgc aattcaacac cgcctacctg
11941 ggcgacctgg cgaagtatgg gatcacggcc caggacgttg ctgcggcagg ctgctatccc
12001 tatgacctgg cggcctggcg gttgcgcggg cacattcgca acgacagggg cgatctgtgg
12061 acacgcgccg ctaactatca ctcgcgcacg ccgtcgaaga acgcgatcta tcgcgccgat
12121 ctgatggtga aggccgacaa gtgggcgaag tggctggatg cgcgtttcgt caccgtcaac
12181 tatggcccca gctcgccggc gcagccggca gggaagggga ccacacttgc ggccgctgat
12241 acgtcggcag cagcgccggc cgaagcgcag ccgatgaagc aaggccggat cacccgcacc
12301 agcctccgca gctcgggtta cgtaccccgg cagctcatca tcaacaacac gccataagga
12361 ggaacggccg tttagcggct aaagcctatg ggcattcgca acctgacgca gcgatacatg
12421 aacggggcca gggcctacgc ggcctgggcg gcatcgcagg cgaaagcgcc gtttgatctt
12481 ctggtactgg gcatcgggcc tgtcatcgtc tttggcctgg tcgcgcatac gctgctcgcg
12541 ttcctgccca catgggccat gtacgccgcc ggcgctctgc tggtcctcgc ggccctgcct
12601 ttggcgctgc acgtcctccg ggaatacgcg ctgcgctatg ggcgcaaata gcgccctgca
12661 gggcgttctt actccaaggg ggagggcatg aatacacgcg ccatgaacga cgccagcggc
12721 cgggcctcgc tgcctgccat ggtgatcgcc gacggcacca ttgaagcctt gaagtggctc
12781 gccttgcttg ccatgaccgg ggatcacgtc aacaagtacc tgttcaacgg tacgctgcca

12841 tatctgttcg aggcggggcg cttggccctg cctcttttcg ttttcgtcct ggcgtacaac
12901 ctcgcccgcc cgggcgcgct cgagcgcggt ttgtacgggc gagcgatgaa acgcctgttg
12961 gccttcggcc tggtcgcctc ggtcccgttc attgcgttgg gtggagtggt gggcggatgg
13021 tggccgctga acgtcatgtt cacgctgttg gccgcaaccg cgatgctcta cctggtcgag
13081 cgcggccgct cggtcgctcc tatagcgctg ttcgtcgtgg ccggcggcct ggtcgagttc
13141 tgttggccgg cgctgctgct ggccgcgtct gtctggttgt acctcaagcg cccgacgtgg
13201 gcggccgcgt tgatggcgct gctgtcttgc gcgtccctgt ggtacatcaa tggcaacctt
13261 tgggcgcttg ctgttgtgcc cctggtgatc gtcgccgccg gcgtcgatct tcgtgtcccg
13321 cgcctgcgct gggcctttta cacgtactac ccgctgcatc ttgccgctct ttggctgatc
13381 cgcattccga tgcgcgaggc gggctacttg tttttcacct gacctttgag attccaatat
13441 gcaattgctc aagaaatgca ccatcgcggc cctgccgctg ctcgccctgt ccggctgcgc
13501 actgctgaac atccccatgc cgacgccgcc cggttcgacc ccgccggaaa tgctgaccgt
13561 gccagtggcg caaatctgcc gcgacgctga caagaaccct gttcgggcaa cggagctgta
13621 cggcaagaaa gggttgtcgg ccaccggcaa ggtgcaggtg atttccgaag gcttcaagcc
13681 tcgctatcgg gtgctgctgc gcgctggcag cgcctcggtc catgctggga ccgataacca
13741 gctcgccatc aagtcggttt ccaccggcca gaccacgcgc gtcactggca ccgtgaagga
13801 cgtgtcctac gaccataacg gctgctcgat ctcgcttgac gatgcgaagt tctactgagg
13861 ggagggcggc ggatgctgac acggttgaag ggcttccttg ctcgtcgccg cgagttgaag
13921 gaactggatg tgtccgtggt gagccggccc cggccggctc cggcggaatt ggtccaggtt
13981 gatgcacgcg aggccgtttg gcgcgtgccg gtgcccggcc aggccgaccg cttcatgtcg
14041 gccaagcctg gcgcgatcaa cgatgaaatg ttcgtggttc gggtggacac cgaagcgttc
14101 tatcgggctt ggctgcgcag cagctcgacg ggccgcgaaa cgcggtcgga caactgcccg
14161 ctgcgctcgg aaatgccgca ggactacaag ttcaagcacg ccgtccaggg cttcgcgcac
14221 ggcagggaaa atcctgtgcc gctggccttc gccggcgcgc accaggagcg ccaccgggtg
14281 gacattggtt tcagcaacgg ggtcacgcgc tcgttctggc tgattgccaa caaggctccg
14341 tcgttcccga tccaggtcca cggccgggag tcggccgagc tgctgaacaa ggtttgcggc
14401 ctcgatcctg cgccgctgtc gttcacggaa ctgttcgcgc aggcccaacg ccaggctccg
14461 caggtcgcca caccggcccg gcctgcgccg gcagcggcca cccggccagc tcccaaggtg
14521 cagccacgcc ccggccgaag cggcccgcgc aaaggccgcg gactctgact acaaccgtgc
14581 gcaaggcgca ttagggagga tgtatgtatg taatcgcctg cggcatcgtt gccggcttgg
14641 cggctgcggt ggccctgttg ggcttcacgc cgatgatgga ggcgcttgcc gccggcgaac
14701 gccgcaaggc actcgcgcaa tggacgcgga cgatgttcct ggtgctgctg cctgtcgtgc
14761 tgatgtgcgc gcccatcggg tccagcattt acgacgccgt gcaagcggac gctggcaagc
14821 ccatcgcttt ccacaacggc cggatcacgg tcgtcatggc cctggtcggc agcttggccg
14881 ttgtcctggt cgcggctgcg cgtgcggtgg tcaaccgcaa gcatgccagc ttctggttcg
14941 tcggctgggt gatggcgtcg gttttggccg gcggcgtcgg cgcgatcgcc agcgcgaagc
15001 aactggcgtt cctcggcgaa catagcggca tggtggcctt cggcttcttc cgcgaccagg
15061 tgaaggacat gcactgcgat gcggacgtga tcctggcccg gtgggatgaa aaggcgaact
15121 cgccggtggt ctaccgctgc ccgaaggcgt acctgctcaa caggttcgca tccgcgccct
15181 tcgtgccctg gccggactac accgaggggg aaagcgagga tctaggtagg gcgctcgcag
15241 cggccctgcg ggacgcgaaa aggtgagaaa agccgggcac tgcccggctt tatttttgct
15301 gctgcgcgtt ccaggccgcc cacactcgtt tgacctggct cgggctgcat ccgaccagct
15361 tggccgtctt ggcaatgctc gatccgccgg agcgaagcgt gatgatgcgg tcgtgcatgc
15421 cggcgtcacg tttgcggccg gtgtagcggc cggcggcctt cgccaactgg acaccctgac
15481 gttgacgctc gcgccgatcc tcgtagtcgt cgcgggccat ctgcaaggcg agcttcaaaa
15541 gcatgtcctg gacggattcc agaacgattt tcgccactcc gttcgcctcg gcggccagct
15601 ccgacaggtc caccacgcca ggcacggcca gcttggcccc tttggcccgg atcgacgcaa
15661 ccaggcgctc ggcctcggcc aacggcaagc ggctgatgcg gtcgatcttc tccgcaacga
15721 cgacttcacc aggttgcagg tccgcgatca tgcgcagcag ctcgggccgg tcggcgcgtg
15781 cgccggacgc cttctcgcgg tagatgccgg cgacgtagta cccggcggcc cgcgtggccg
15841 ctacaaggct ctcctggcgt tcaagattct gctcgtccgt actggcgcgc aggtagatgc
15901 gggcgacctt caaccttcgt ccctccggtt gttgctctcg cgtcgccatt tccacggctc
15961 gacggcgtgc ggatcggacc agaggccgac gcgcttgcct cgcgcctcct gttcgagccg
16021 cagcatttca gggtcggccg cgcggccgtg gaagcgatag gcccacgcca tgccctggtg
16081 aaccatcgcg gcgttgacgt tgcgcggctg cggcggccgg ctggccagct ccatgttgac
16141 ccacacggtg cccagcgtgc ggccgtaacg gtcggtgtcc ttctcgtcga ccaggacgtg
16201 ccggcggaac accatgccgg ccagcgcctg gcgcgcacgt tcgccgaagg cttgccgctt

16261 ttccggcgcg tcaatgtcca ccaggcgcac gcgcaccggc tgcttgtcta ccagcacgtc
16321 gatggtgtcg ccgtcgatga tgcgcacgac ctcgccgcgc agctcggccc atgccggcga
16381 ggcaacgacc aggacggcca gcgcggcagc ggcgcgcagc atggcgtagc ttcggcgctt
16441 catgcgtggc cccattgctg atgatcgggg tacgccaggt gcagcactgc atcgaaattg
16501 gccttgcagt agccgtccag cgccacccgc gagccgaacg ccggcgaaag gtactcgacc
16561 aggccgggcc ggtcgcggac ctcgcgcccc aggacgtgga tgcgccggcc gcgtgtgccg
16621 tcgggtccag gcacgaaggc cagcgcctcg atgttgaagt cgatggatag aagttgtcgg
16681 tagtgcttgg ccgccctcat cgcgtccccc ttggtcaaat tgggtatacc catttgggcc
16741 tagtctagcc ggcatggcgc attacagcaa tacgcaattt aaatgcgcct agcgcatttt
16801 cccgacctta atgcgcctcg cgctgtagcc tcacgcccac atatgtgcta atgtggttac
16861 gtgtatttta tggaggttat ccaatgagcc gcctgacaat cgacatgacg gaccagcagc
16921 accagagcct gaaagccctg gccgccttgc agggcaagac cattaagcaa tacgccctcg
16981 aacgtctgtt ccccggtgac gctgatgccg atcaggcatg gcaggaactg aaaaccatgc
17041 tggggaaccg catcaacgat gggcttgccg gcaaggtgtc caccaagagc gtcggcgaaa
17101 ttcttgatga agaactcagc ggggatcgcg cttgacggcc tacatcctca cggctgaggc
17161 cgaagccgat ctacgcggca tcatccgcta cacgcgccgg gagtggggcg cggcgcaggt
17221 gcgccgctat atcgctaagc tggaacaggg catagccagg cttgccgccg gcgaaggccc
17281 gtttaaggac atgagcgaac tctttcccgc gctgcggatg gcccgctgcg aacaccacta
17341 cgtttttttgc ctgccgcgtg cgggcgaacc cgcgttggtc gtggcgatcc tgcatgagcg
17401 catggacctc atgacgcgac ttgccgacag gctcaagggc tgatttcagc cgctaaaaat
17461 cgcgccactc acaacgtcct gatggcgtac ttacccaaag aacagctagg agaatcattt
17521 atgctcagca cacttccaca agctcatgca actttcttga accgcatccg cgatgcggtc
17581 gcttccgatg ttcgcttccg cgctcttctg atcggcggct cttacgttca cggaggactc
17641 gatgagcact ccgatttgga tttcgacatc gttgttgagg acaactgcta cgcagatgtc
17701 ttgtctacac gcaaggattt tgccgaggca ctgcccggct tcctcaacgc gttcaccggc
17761 gaacatgtag gagaaccgcg ccttctgatc tgcctatatg gtccgccact gctacacatc
17821 gatttgaagt tttctcttgc ttccgatctc gaccagcaaa tcgagcggcg ggcggttctg
17881 tttgctcgtg atccggcaga gatcgagaag cgcattgagg cggcagcggt ggcatggcca
17941 aaccgtccct ccgagtggtt cgaagcacgt tgtcagcgcc agtgatataa gacggtaatt
18001 caccatttgg attgtccgct ccacccaaca tgttgtttcc ttaaggttct cacaccagaa
18061 aggacatcaa catgctgagc agagaggact tttacatgat aaagcaaatg cgccagcagg
18121 gcgcgtacat tgtcgatatt gcgactcaga ttggttgctc tgaacggacg gtcagacgct
18181 acctcaaata ccctgaaccg ccagccagaa agacccgcca caaaatggtt aagctgaaac
18241 cgtttatgga ttacatcgac atgcgcctgg cagagaatgt ctggaatagt gaggttatct
18301 ttgcggagat taaggcaatg ggttatacgg gcggacgttc catgctgcgt tactacatcc
18361 agcccaaacg taaaatgcgt ccgtcaaaaa gaacagttcg cttcgaaact cagcctggat
18421 accagctcca gcacgactgg ggcgaagttg aggtggaggt tgccgggcaa cggtgcaaag
18481 ttaactttgc ggttaatacg ctggggttct cccgccgctt ccatgtcttc gccgcaccaa
18541 aacaggatgc tgagcatacc tacgaatcac tggttcgcgc cttccgctac ttcggtggtt
18601 gtgtgaaaac ggtgctggtt gataaccaga aggctgcggt gctgaagaat aacaacggga
18661 aagtcgtgtt caactccgga ttcctgttgc tggccgacca ctataacttc ctgccacggg
18721 catgccgtcc acgcagggcc agaacaaaag gtaaggttga gcggatggtg aaatacctca
18781 aggagaactt cttcgttcgg taccgcaggt tcgacagctt cactcatgtt aatcaacaac
18841 tggagcaatg gatagccgat gtggctgaca aacgggaact tcgccagttc aaagaaacgc
18901 cggaacagcg cttcgcgctg gagcaggaac atctgcagcc gttaccggat acggacttcg
18961 ataccagtta cttcgacatc cgccatgtgt cctgggacag ctatatcgag gttggtggta
19021 atcgttacag cgttcccgaa gcgctgtgtg gtcagccggt atcgatacga atatcgctgg
19081 atgacgagtt gcggatctac agtaatgaga aactggtggc ctcacatcgc ctctgttcgg
19141 catcgtctgg ctggcagaca gtgccggagc atcacgcccc gctctggcag caggtcagtc
19201 aggtggaaca tcgaccactg agtgcctatg aggagctgtt gtgatgcatg agctggaagt
19261 cctgctgagt cgcctgaaaa tggagcatct gagttatcac gttgaaagcc tgctggaaca
19321 ggcagctaaa aaagagctga actaccggga gttcctgtgc atggcgctac agcaggaatg
19381 gaacggcagg catcagcgcg gtatggagtc caggctgaag caggctcgtc tgccgtgggt
19441 caaaacgctg gagcagttcg actttacctt ccagccgggc atcgaccgta aggttgtccg
19501 ggaactggct ggtctggcgt tcgtggagcg cagcgaaaac gtgatcctgc tgggacctcc
19561 tggtgtcgga aaaactcatc tggccatagc tcttggcgtg aaagcggtgg atgcgggaca
19621 tcgggtactg tttatgccac tggacagact gatcgcgaca ctgatgaaag cgaaacagga

19681 aaaccggctg gagcgtcagc tgcagcaact gagttatgcc cgggtgttga tcctggatga
19741 aataggctat ctgccgatga acagagagga agccagcctg ttcttccggc tactgaaccg
19801 tcgatatgaa aaagcgagca tcatactgac gtcaaacaaa gggttcgcag actggggaga
19861 aatgttcgga gatcacgtgc tggcaacagc gatactggat cggttgctac atcactcaac
19921 cacgctgaat atcaaaggag agagttaccg gttaaaagag aaacgtaaag ctggagtgct
19981 gaccaaaaac acaacgccaa tcagtgatga tgaaatggtg aaaagcggac agcatcagta
20041 acgaaagtat cttagcgggc atgaaaatgg caaataacgg tcaaacatcg tggcgttgac
20101 aacgtgcctg gatctggcta cactatgcgg ccaccaagct cgcccgtggc gagctttacg
20161 aagcgatcgg catgctcggt ttcttccgtg agcaagtgtt aggacctttg ctctaccgtc
20221 gcgctggaaa ggaccagcgc ggagtgaggc gattggaaac ccttcgactg gatgaagagc
20281 gcagactagc caccaccatt gcgctgcacg atgcgttgtc tgtcagggat gccatcaaag
20341 catctgcctc catctatctc gacctccgag ccgccgatcc gtcgttggaa ccgacaacgc
20401 atatgccagg tcttctgtac gacttaatag aacgtgcggt accaggcacg cctaaccgtc
20461 agtgagattg gatgagtgaa cgatattgat cgagaagagc cctgcgcagc cgctgccgtg
20521 cccgagagca tggcggctca cgtgatggga tacaaatggg cgcgtgataa ggttggtcag
20581 tccggctgcg cggtctatcg gctgcatagc aagtcaggcg gctccgactt gtttctgaag
20641 cacggcaaag atgcttttgc cgacgacgtg actgatgaaa tggtgagatt gcgttggctg
20701 gcgggggcaca tttctgtgcc ctccgttgta agcttcgttc gcacgcccaa tcaggcatgg
20761 ctcctgacaa cagcaataca tggaaaaacg gcatatcaag tgctgaaatc ggatttcgga
20821 gcccgtctcg ttgttgttga cgcattggcg gcgttcatgc gccgactgca tgcgatccca
20881 gtgagcgaat gctccttcaa cagtgaccac gcatgcaggc ttgcccgagc gcgggagcgt
20941 atcgaggcgg ggggttgttg atgtcgatga cttcgataag gagcgcgaag ggtggacggc
21001 cgaacaggtt tgggaggcga tgcatcgcct cctaccgctc gcgccggacc cagtcgtgac
21061 gcacggcgat ttttcactcg ataatctact tatcgtcgaa ggtaaggtag tcggctgcat
21121 cgacgttggg cgggctggta ttgctgatcg ataccaagac cttgccgtgt tatggaactg
21181 tcttgaggag ttcgaacctt cgcttcagga gaggcttgtt gcgcaatatg gcattgccga
21241 tccggatagg cgcaagctgc aatttcatct cctgctggac gaacttttct aaggcgatgc
21301 cccctcgacc tcgatcaggg aggcgttcag gacgactcac aaagaaagcc gggcaatgcc
21361 cggctttttc tgctgctacc tccgtagtcg taaggtcgtt gcaggtgctc gggtgcggta
21421 caactcgccg gtcgccagct caagcgcgat cacgtcgttg ccgtcgtagt tgacgatgat
21481 gctgttgggc cgactgtcct cacgcttcgc agggagaggc cagccttcaa tcgaagccgg
21541 cgcaagctcg tagtgcttcc cggtttcgac gctgcgcagc gtccaggtcc tgcaaccggc
21601 cacgccggtc gcagaaacca cggcgagcga gccgcgaaaa tcgtgcgggt acgcctcgat
21661 gttcatacgc ctcctagatc gagcgcgagc gtttctgctc ggccttggcc gcctgttcct
21721 gggacacctc gccgatgacc ttgccctggc cccggctgta ggcgatttcg tagttcttgc
21781 cgacaaccgg cggcttctca aagatgcccc ggctgtgttt cacgatcccg ccttcgctga
21841 actggtagac gttgcgccca tcgtcgtgca gcacctggcc gacgtgcttg tgcgggtgga
21901 cgtttttgct tgcgtccttc gcatcgctca actggtgaat gcctttcggt agccccgcct
21961 cgggcaacac cttcatggtc agccattcgc cgttcactac ctggtccacc tggcggctgc
22021 cgttcatgac ggcgatcttg acgctgccct cgggtttcat gatgacgcca gggcttgccg
22081 atgtgcgtgg tgccccgatc tgtactttgt tcatacgctc tagttctcct tagtaggttc
22141 tcgcgcggcg ttgccgctgt tcttgctgct cgatgtcttg ctgcttgagc tgctgcacct
22201 tctgccgctg gccctcgtcg agaagcacct tgccgacagc acttctcacc tggcgttcaa
22261 ccccgtcctt gcccaggctg ctgcgctcgg acaggtcgtt gaaatcggtg tgcttcttca
22321 tgttcgacag ggcggcgagc tggccatcgt tcaacagcga ttccttgagc ttggccgtgt
22381 cggcctcgct caactggacc ttgccggccg ccgcgtccgc gaggcgcttt tcagcgtgca
22441 aatggttgcg gtagttctcc ggggtgatcg gcggcagctc cttcgggtag gcgttctcgc
22501 ccggcgcgaa gattgggaag atggccttgc cgccgaccgc cttggcggcc tcctgtgcct
22561 tcgtcctgcc gggattcacg ccctgggtga tctgcacctg gcggtcgtcg tcgccggcga
22621 tcacaacggg cttgtccggg aatttcgcgt gcagggcctc ggcaacagcc tgtaggttgc
22681 cggaatcgaa cgcggcgaca gtcgcgtgcc ccagcgcttc ggccactgtg gcggcggtgg
22741 catagccttc gccgatcacc agcgccggcg cggccgcgag cgcatccatg ccaccgacga
22801 catggaagca tccttccttg cggctgtcct tggcgaagcg cttggtgccg tcctcctgga
22861 tgtactgcat ggtccattgc ttgccgtcgg cgtcgtaggc cgggatgtag gtttctggc
22921 cctcctggtc ggtaaggacg ccggcgtgca cctgtagacc cttgtcgcgc aggtacggcg
22981 tcggttccgt gatgggaacc aggctttgcg cctggcggcc gatgcgctgc gccgtggctt
23041 cgtgctggcg ttcttgttcc tcggcacgcg cggccagctt ggccgccgcc tcggcctgca

23101 tcttggcctt ctcggcgggg tccagggcgt agcccttggc cttccacttc atttcgacgc
23161 cggtgcggtt gtttttgatg taaccggccg ggtggccgtc gaggtggccg acgtagaagc
23221 ccgacttctc gcccttcttg tcgccctcgg tctcgatgcg gtgcttcttg ccgtccatga
23281 tggggtgctc gccgcctggg gtgacgacgc agcccatgct tttcagggcc tccgcgaact
23341 catcttcggg ggtgacggcc ggggattgct gggtgggcac gttgtccggc agccagcgtt
23401 gcagcttgcc catgtcggcg ttcggtccgg cgtaccagga cttggccacc ttgtcccact
23461 gcgcgccggc cgccttggca acctggcgct cgccgtaggg cacggccagg tagacgcgct
23521 cctgggccgc gttggggcgc tcggccgtgg gttgggtagg ctgggcctcg gcgcgggcct
23581 ctacggccgc tgtagcgccc tcgcgcgccc atttggcgaa cggggcaggg tcaacccctg
23641 ccggaacgta ccaggcgcgt tcctggcggt cccagcgcgc tccaagggcc ttcacctcgt
23701 ctttctcctt gaacggcacg ttcaagtagg cgcgctcggg cttggcgggg gcttgagcgg
23761 ccgccggctg ctcggcggcg ttcatggcct gggccatttc ctgctgctcg cgctcgtagt
23821 cggcgatccg gcgctgtagg tcctcgtcgt gcagcatggc cgtgccctcg gcggccttgc
23881 gcgcctcctt ggcggcaacg cggtcctcgt cggtgctgtt gggatcgcgg cgaactcgct
23941 cttcatggat gcgggcgaac ttcgcggcct gctcgtactc gttggccgtc gcgtaagcgt
24001 cgatcacggc caggcggtcg gcgagcgctt cggcattggt ctgcgcgtcg ggccggcga
24061 agtcggcaag ccattggtgg ccgcccccacg catggttcgc atagacgccc caaaactccg
24121 gctctcggtc gccggccggc accacggacc gttcgccgtc gtgctcgacc tcgacgttgg
24181 cctggacctg gacgcggccg gtccaatcgg caggcagctc aaagcccagc gtggtttcgg
24241 tcagcgcggc cagcgattgg ttgccttccg ccggctccgc gccggcgcgg tacatgcgca
24301 gggtctgcgc gatcagctcg tcggccggcg caatggccgg tcgtgccacc tggtcttgct
24361 gttgctccat agttgccccc tgcgcaggct cgatggcctg ctgggtcgtt tgttcttgaa
24421 tttgcttctg ctcgaacgcc aggacgaaat cctggatctt ctccgcgtcg gcggccgcgc
24481 ggaaaatctc tagcgggtcc tcttgtagcg ccttgatcca cgatccgaca taggccgcgt
24541 gctggccggg gtcgtggccg atgcccagct cgtcgcccag gatcatgctg gcaatctcgg
24601 cccgcagctc ttccttggcg taccccctcgc tcccgaaggg atgcgccagg tcgcggtcca
24661 gccgcgacgg gtggccggtc cagtgcccca gctcatggag cgcggttgcg tagtagttgt
24721 cggcgctcgg gaactggcct ttgtcgggca gatggatgct gtccgtggac ggccgataaa
24781 acgcgcggtc gtgctcgccg tggcggatgg tggcacctga cgccgcaagg atgtgctcgg
24841 cccgctcgac ggcgctccaa gtctgttcct tgcgttccaa cggcggcagg ccgtcgatct
24901 gctccgcatt gaacacggtg gcgaagaaca cgcgcgggcg ttcgagctgc accgtcacct
24961 tgaccggatc gccgttggca tcgaggaccg gcttgccggt ctgctcgtcg gtcttggtct
25021 gctcttcgct gaacttccaa tactggatcg gcgtgccttt ctcgccgcga cgcacctgtg
25081 cgccggcggc agcggcctgc ttgtaggtca tccagcgcgg gtccgcatgg ccctgggcca
25141 tgagctgaat cgcgttgatg cccttgtaac gcttcccggt agtcgggttg agcgggatga
25201 aggagccggg catgcccggt tcccacggtt tttgccacgg cgcagtgccg gctttcagtt
25261 gctcaatgag gcgttcggca acctgctcgt ggaacggctt tttgacctct gccatagcca
25321 attacctccc gtcattggcg gccgcggtcg tcgtgtcctc gggcacggtc gcgtccacca
25381 ggtcaatgtc gctctcggcg gcgtcctgct cgctcaaggc gtcctcgggg aaggcccccgg
25441 ccttctccgc ttcttccggg tcgaactcga cctggaagcc gggcgtcatc gcatcggcga
25501 gcttttcacg cagggcggcg gcggcgttcg cggtatcgtc gaacggcgca aagtcgtcct
25561 gctgcacggt cttgggatca ttcatcgctt tcactcctgg ttggtgccgt tacggccttt
25621 gctgtagtcc ggcctgcctt tcaggtcggg gtatgtctgc ttgcacgtcg ggaagttgct
25681 gcacccccac cagaacatgc cgcgcttctt gccaggccga cgggaaaggc cgtggccgca
25741 ggccatgcac ttgtgcagct cggagacttt cggggcctcg cgcgggacgg gcttgccgcc
25801 cttgtcgtcg cacgcgaact tgcagccgtc ggcaaagccg gtgcagcccc aaaagtattc
25861 gttcttgtcc ttcttcttga ggcgtcgcag cggcttgccg caggacgggc aagggtgcgt
25921 gtcgatcttc atgttgaggc cgttgtcttt gatgttggcg acctcggcgc cgatgtattc
25981 catcagctcg ttgacgaacg acagcgtgtc gcgctcgccg gcctggatgg ccttctgctg
26041 ctcatgccag agcgcggtca tgtcggggaa tctggccgtg tcgggcagtg cgtcgtacag
26101 ctcttcgccg gtcggcgtgg acacgatgtg cttgcccttc tccaccaggt agccgcgctc
26161 gaaaagcgtg gcgatgatgg agtctcgcgt tgccggcgtg ccgatcccgc cgtgctcgcc
26221 ttgcttgccc ttgtcctttt cgatcaagat tttccgcagg cggtcatcgc ggatgtattt
26281 cgcaacgcgg gtaaggtccg acagcagaga ttccatcgtg tacagcggct gcggtttcgt
26341 ctcctgctgc tcggccttcg catcggtgca ggtgccggcc tggccgtcac gcagcttgcg
26401 caggtcctgt tcaatgtcgt cggcattgcc ttccaggtcc tcgttgccgg cgtcgttctt
26461 gtagagaatc ttccagcccg gcgacgtggt gacgttcgag cgcacgccga aacgatgatc

26521 gccgacctgg gcaagcacgt cggtctggtc atacagatgc ttcggccaga actgcgcgac
26581 gtaggcgcgc gcgatcagca ggtaaatctt ctgctcggca tcggtgagct tcgacaggtc
26641 ggccgtgctt tcggtcggga tgatcgcgtg gtgcgcggaa accttggacg agttgaaggc
26701 gcggctcttg atcgtcggat tggcgcgctg cgcagcagcg gccagcatgg gggccgtctg
26761 tgcgatggcc gccagcacgc ccggcgcatc gccgtgctgt tcctcgctca agtattcgca
26821 gtcggaacgg ttgtaggtga tgagcttgtg cttctcgcgc agggcctgcg taatgtcctt
26881 cacctggtcc ggcttgaagc cgaacttgcg cgaggcgtcc atttgcagtt tcagcaggtt
26941 gtagggcagc ggcgcagccg cttccttcgc cttggtggtc acggacacga tgcgggcggg
27001 ttggccgctc acggcggccg cgatgccctc ggcgtgctcc ttgttgctga ggcggccttt
27061 ctcgtccacc ggatcgccgt cggcgacctg gtaacgggcc gggaactgaa tgccctcgac
27121 ctcgaactgg ccgttcacca ggtagtagta ggttttctgg tgggccgcgt tctcgcggca
27181 acggcgcacg acaaggccca ggatcggagt ctgcacgcgc cccacgctca acagcccctg
27241 atagcccttc gcgcgtgccg caagcgtgta caggcgcgtg atgttgaagc cgtatagctg
27301 gtcgccgacg ctgcgggcct cggccgcagc ggacaggccg gcgaactcgc ggttgtcgcg
27361 catcgcggcg agctgccggc gcacgatctt cacgttgttg tcgttgataa gcagccgctg
27421 caccggcaga cggcagttgg cgtattccag gatttcatcg accagaagct ggccttcgtc
27481 gtccgggtcg ccggcgtgaa ccacgctttt cgcctgcttc aacaggctga ggatggtctt
27541 gaactgagct ttcgcaccgg catcgccgga cggtttcttg cgccagggaa tatggacgat
27601 gggcaggtcg gccatgttcc agttggcgta gcgctcgtcg tagtcctccg ggtctagcaa
27661 ggccagcatg tgaccgtagc accaggtcac gcggtcggag ccgcattcgt aatagccgtc
27721 cttgcggctg ccgccgccca ggccctcgac gatggctttt gccagctccg gtttttcagc
27781 gattacaagg cgttcaaatt gcatatatcc ccctaccctc accaggtcag aaccggcctg
27841 atgacggtga tgatttgcga acgattgaca ggcccgaagt agcggccgtc gaaagacgtg
27901 tcgcttacgt cggacataag cagaacctcg gcggtcccca gggtgtagct gtcggactga
27961 taacgaggca gcggccgtcc tgatggatcg gccttgatga gcgcgctgtg aggcagcagc
28021 ccgccattca cgcgcacgcc ggcgtcggtg atggcaacct cgtcgccttt agcggctaaa
28081 actcgcttca tcatgtagcc gtagtcgccg gggcagaaac cgccggcgat gtagccccgc
28141 tccttggcgt ccgaaaacac gccgacttgc ggcgggcaga acatgacgta agccccccttc
28201 tccaccggcg cattcgattt ccagtacagg ccgaccggaa tgcttttggt ggtgttgacc
28261 ttcgcgccgg cgagataggc cgcgccggcg agcaacaagg ccgcgccgcc tccgatggcg
28321 acgtacttgg tgaggcgctg gaagcggctc atatcgtgat cccctcccct tcctcgacgg
28381 tggccgtctg gatcagcttg tcgctgacct tcggagccgg tacggccgcg cgggcctgga
28441 atatcgggtc tttgaagtag agcggctgct tgccgtagat cgcgggatag ccggcgacgt
28501 acacaaccat gtcgcccgcc tcttcaatgc tgccgtcggc gctcttcttc ggccccggca
28561 tgcgcaggca ttcatcgggg gtcagcaatg gccgctgcac ttcctggaag gtccgcgaga
28621 cgttgcccaa cagcgccgac gtgcggcggc cgctcgtcgt gatctgctcc ttcacgatgg
28681 tcgtggtgcc tgtcagtttt gacaggtgct cggccgtctc cacgcggttc ggcgggtagg
28741 cgttctgcac gtggcagttc gacgtgatgc tttcgtcgtg gccgtagccg gtttcgcggc
28801 tcttgagctg gttaatgtcc tggcagatga ggtagcactt gatgccgtag ccggcgacga
28861 aggcaaggga ctcttgcagg atttcgagct tgcccaggct ggggaactcg tcgagcatca
28921 tcagcagacg atgcttgtag tgcgcgacag gacggccgtt ctcgaagtcc atcttgtcgg
28981 ccagcagccg gacgatcatg ttgaccatga cgcgcaccag aggccgcaga cgggccttgt
29041 cgttgggctg cgtcacgatg aacaggctta ccgggtcgtc gtggtgcatc agttgcttga
29101 tgcggaagtc ggacttgctg acgttgcggg ccacaaccgg gtcgcggtac agggccaggt
29161 aggacttggc ggtggacagc acggaaccgg attcttcctc cgggcggtcc atcatgtcgc
29221 gggccgcaga gccgaccgca gggtggttct gcccgtcaac gtggccgtag gtggtcattt
29281 ccatccaaag ctcgcccacg tcgcggttcg ggtcggcaag catgccgtcc accgacggca
29341 gggtggccgg cgtaccctcg ttcttagcct tgtagagcgc gtgcaggatg acgccgacaa
29401 gcagcgcctg gctggttttc tgccagtgcg attccaggcc cttgccgtcc ggatcgacga
29461 tcagggtggc aaggttctgc acgtcgccaa cctcgtactc ggtccccaag cggatttcat
29521 cgagcgggtt ccagcacgcg ctaccctgcg cggatgccgg ctcaaagcgc acgaccttgt
29581 tgcgggcatg cttcttccgc cagccggcgg tcagcgccca caactcgcct ttcaggtcgg
29641 tgatgacggc gctgtgcgcc caggaaagca gcgtcggaac gaccaggccg acgcccttgc
29701 cggagcgcgt cggcgcgtag gtcaagacgt gctcggggcc gttgtgccgc aggtagtgga
29761 acttgccgtc cttgtcctgc cagccgccca catagacgcc gctggaagtg ggcgggtgtt
29821 tgcctgacac cagctcgacg acggtgcgcg gccggggcag caggccggcg gcctgtatgt
29881 ccttcttgtc ggcccagcgg gccgaaccgt gcagatagtc gttcgccttg ccggtgttcg

29941 ccttgaccat ctgcgtgacg gccgtgccca gcaggcccac ggtcgaaacg accataccca
30001 tgctggccgc gcgcatgaaa tcgtcgggat attggccgta ccacttgccg gcccattgaa
30061 ggatcgacca gggcgtgtag acgtggttga tattccagcc aagtccggcc tgatactgga
30121 aggaatgggc gaaatattgc gtcgcggtct gcaagcctgc cccaagggac aggccggcga
30181 ggatgggaac ggtcttgctg gccttcggtt ttttcgcccg tatctgtggc cccacggcgt
30241 tgtttcggtt cttcatctac tcctacctcg ggtagtttta agggagcctc gcggggtcac
30301 ggtgacggga tcaccgatgg cgaggcgctt catgcgttgc accgtggcct tatcgacggg
30361 cagcaccaga atctcgtcgt tttctttcct caacagggcc agcgcctggt cctcgacgtt
30421 ccgggtgcct gcataggaca gcgcaccaac ataatcagta tatcgtgcat gcttcggtat
30481 atcgaagccg tttagccgct tttgctcgcg ctcggcaaca tatttctcgg ccgccgcgat
30541 ctgttcgggc tttagccctc ttcctggccc agaaactccc cgtcgcagtg cgtgagctgg
30601 ttcggctcct tgctgctcca cgtgaccagg aacatcacgc ggcaatagca tttcagctcc
30661 gccggcgatg cgaaccacac cgagttggga cagcgctcgc aaacggtttt ggctttgggg
30721 cggcggcttt cgtccaatgc gtccaacgtt gggcttgcgg agtgcgacgg ttccgccggc
30781 gctgacggcg cgagcgtccc gtcggtcgcc gtcgccgcct gtggcgttga gggtggttct
30841 ggctgcggca ggtcgaatgc ctccatcgcc gccgcgatct cttcgtccgt catttcgttc
30901 gggttgctca tgtgcttgct ccttcgtcag tagttcttga cggcggcgct caagggcggc
30961 gtcgtcaaag gtgattgcca gacggccagc ggcggccgcc tgcgcgatcc gctccttgaa
31021 ctctgctgtg ccgttgacgg tgatccggtc gccgaagcgc tccattgcca ggcgcagggc
31081 ggcgtccagg ccgtccgtgg tggcctcgcg cgagacttgc aggcggtcgc cgtcgtcgcg
31141 gacggcgctg ctgccgacgc gatagatgat ggttcccttc ttcgtgatgt tgtccgtcac
31201 ggccgcatgg cccggcttgg cctcgccgct gccctggatg gtgttgccct tgaggtcgct
31261 gcggccctcg cgtgcgcgca gcgcggccag ggccttgtcg tcgcccttca tcgcctcggc
31321 cttgagccag tcggcccacg cgcggcgctg cgtgcgctcc tggaccgcct gacggccctg
31381 ccggtactcg cggttgatct tgtccaggtc ggcgcgcaga gccttgtgcg cctgcgcgta
31441 catcagtcgc tttgcaatgc gcccctcgcc cagcagcttg atagcggcgc ggcgcagccg
31501 gttgctgcgc atcgcggctt caatcaggcg gtcacgacgc cggcgcagcg tgtccagctc
31561 gcccttgcgc acggccccca tttcctggcg ttcagactga taccgggcgt atagctcggt
31621 ggtgtcgatg cgggtcttga gcggcttcgc tcgatactcc cgccgccggg gggcttcgcc
31681 gccctcggct ggcgtgaatg ccccgaatcg ggcttcgagc ttcggcttgg acaggtcgcg
31741 cgaaacggtg ctggccttga ccgtcgtgcc gtcgccggcc tcgaagatga agccgtttcc
31801 gcgctcgcgc agcttaagcc cgttttcccg caggacgcgg tgcaggtcct cccaggattg
31861 cgccgcttgc agctccggca ggcattcgcg cttgatccag ccgaccaggc tttccacgcc
31921 cgcgtgccgc tccatgtcgt tcgcgcggtt ctcggaaacg cgctgccgcg tttcgtgatt
31981 gtcacgctca agcccgtagt cccgttcgag cgtcgcgcag aggtcagcga gggcgcggta
32041 ggcccgatac ggctcatgga tggtgtttcg ggtcgggtga atcttgttga tggcgatatg
32101 gatgtgcagg ttgtcggtgt cgtgatgcac ggcactgacg cgctgatgct cggcgaagcc
32161 aagcccagcg cagatgcggt cctcaatcgc gcgcaacgtc tccgcgtcgg gcttctctcc
32221 cgcgcggaag ctaaccagca ggtgataggt cttgtcggcc tcggaacggg tgttgccgtg
32281 ctgggtcgcc atcacctcgg ccatgacagc gggcagggtg tttgcctcgc agttcgtgac
32341 gcgcacgtga cccaggcgct cggtcttgcc ttgctcgtcg gtgatgtact tcaccagctc
32401 cgcgaagtcg ctcttcttga tggagcgcat ggggacgtgc ttggcaatca cgcgcacccc
32461 ccggccgttt tagcggctaa aaaagtcatg gctctgccct cgggcggacc acgcccatca
32521 tgaccttgcc aagctcgtcc tgcttctctt cgatcttcgc cagcagggcg aggatcgtgg
32581 catcaccgaa ccgcgccgtg cgcgggtcgt cggtgagcca gagtttcagc aggccgccca
32641 ggcggcccag gtcgccattg atgcgggcca gctcgcggac gtgctcatag tccacgacgc
32701 ccgtgatttt gtagccctgg ccgacggcca gcaggtaggc cgacaggctc atgccggccg
32761 ccgccgcctt ttcctcaatc gctcttcgtt cgtctggaag gcagtacacc ttgataggtg
32821 ggctgccctt cctggttggc ttggtttcat cagccatccg cttgccctca tctgttacgc
32881 cggcggtagc cggccagcct cgcagagcag gattcccgtt gagcaccgcc aggtgcgaat
32941 aagggacagt gaagaaggaa cacccgctcg cgggtgggcc tacttcacct atcctgcccg
33001 gctgacgccg ttggatacac caaggaaagt ctacacgaac cctttggcaa aatcctgtat
33061 atcgtgcgaa aaaggatgga tataccgaaa aaatcgctat aatgaccccg aagcagggtt
33121 atgcagcgga aaagcgctgc ttccctgctg ttttgtggaa tatctaccga ctggaaacag
33181 gcaaatgcag gaaattactg aactgagggg acaggcgaga gacgatgcca aagagctaca
33241 ccgacgagct ggccgagtgg gttgaatccc gcgcggccaa gaagcgccgg cgtgatgagg
33301 ctgcggttgc gttcctggcg gtgagggcgg atgtcgaggc ggcgttagcg tccggctatg

33361 cgctcgtcac catttgggag cacatgcggg aaacggggaa ggtcaagttc tcctacgaga
33421 cgttccgctc gcacgccagg cggcacatca aggccaagcc cgccgatgtg cccgcaccgc
33481 aggccaaggc tgcggaaccc gcgccggcac ccaagacgcc ggagccacgg cggccgaagc
33541 agggggggcaa ggctgaaaag ccggcccccg ctgcggcccc gaccggcttc accttcaacc
33601 caacaccgga caaaaaggat ctactgtaat ggcgaaaatt cacatggttt tgcagggcaa
33661 gggcggggtc ggcaagtcgg ccatcgccgc gatcattgcg cagtacaaga tggacaaggg
33721 gcagacaccc ttgtgcatcg acaccgaccc ggtgaacgcg acgttcgagg gctacaaggc
33781 cctgaacgtc cgccggctga acatcatggc cggcgacgaa attaactcgc gcaacttcga
33841 caccctggtc gagctgattg cgccgaccaa ggatgacgtg gtgatcgaca acggtgccag
33901 ctcgttcgtg cctctgtcgc attacctcat cagcaaccag gtgccggctc tgctgcaaga
33961 aatggggcat gagctggtca tccataccgt cgtcaccggc ggccaggctc tcctggacac
34021 ggtgagcggc ttcgcccagc tcgccagcca gttcccggcc gaagcgcttt tcgtggtctg
34081 gctgaacccg tattgggggc ctatcgagca tgagggcaag agctttgagc agatgaaggc
34141 gtacacggcc aacaaggccc gcgtgtcgtc catcatccag attccggccc tcaaggaaga
34201 aacctacggc cgcgatttca gcgacatgct gcaagagcgg ctgacgttcg accaggcgct
34261 ggccgatgaa tcgctcacga tcatgacgcg gcaacgcctc aagatcgtgc ggcgcggcct
34321 gtttgaacag ctcgacgcgg cggccgtgct atgagcgacc agattgaaga gctgatccgg
34381 gagattgcgg ccaagcacgg catcgccgtc ggccgcgacg acccggtgct gatcctgcat
34441 accatcaacg cccggctcat ggccgacagt gcggccaagc aagaggaaat ccttgccgcg
34501 ttcaaggaag agctggaagg gatcgcccat cgttggggcg aggacgccaa ggccaaagcg
34561 gagcggatgc tgaacgcggc cctggcggcc agcaaggacg caatggcgaa ggtaatgaag
34621 gacagcgccg cgcaggcggc cgaagcgatc cgcagggaaa tcgacgacgg ccttggccgc
34681 cagctcgcgg ccaaggtcgc ggacgcgcgg cgcgtggcga tgatgaacat gatcgccggc
34741 ggcatggtgt tgttcgcggc cgccctggtg gtgtgggcct cgttatgaat cgcagaggcg
34801 cagatgaaaa agcccggcgt tgccgggctt tgtttttgcg ttagctgggc ttgtttgaca
34861 ggcccaagct ctgactgcgc ccgcgctcgc gctcctgggc ctgtttcttc tcctgctcct
34921 gcttgcgcat cagggcctgg tgccgtcggg ctgcttcacg catcgaatcc cagtcgccgg
34981 ccagctcggg atgctccgcg cgcatcttgc gcgtcgccag ttcctcgatc ttgggcgcgt
35041 gaatgcccat gccttccttg atttcgcgca ccatgtccag ccgcgtgtgc agggtctgca
35101 agcgggcttg ctgttgggcc tgctgctgct gccaggcggc ctttgtacgc ggcagggaca
35161 gcaagccggg ggcattggac tgtagctgct gcaaacgcgc ctgctgacgg tctacgagct
35221 gttctaggcg gtcctcgatg cgctccacct ggtcatgctt tgcctgcacg tagagcgcaa
35281 gggtctgctg gtaggtctgc tcgatgggcg cggattctaa gagggcctgc tgttccgtct
35341 cggcctcctg ggccgcctgt agcaaatcct cgccgctgtt gccgctggac tgctttactg
35401 ccggggactg ctgttgccct gctcgcgccg tcgtcgcagt tcggcttgcc cccactcgat
35461 tgactgcttc atttcgagcc gcagcgatgc gatctcggat tgcgtcaacg gacggggcag
35521 cgcggaggtg tccggcttct ccttgggtga gtcggtcgat gccatagcca aaggtttcct
35581 tccaaaatgc gtccattgct ggaccgtgtt tctcattgat gcccgcaagc atcttcggct
35641 tgaccgccag gtcaagcgcg ccttcatggg cggtcatgac ggacgccgcc atgaccttgc
35701 cgccgttgtt ctcgatgtag ccgcgtaatg aggcaatggt gccgcccatc gtcagcgtgt
35761 catcgacaac gatgtacttc tggccgggga tcacctcccc ctcgaaagtc gggttgaacg
35821 ccaggcgatg atctgaaccg gctccggttc gggcgacctt ctcccgctgc acaatgtccg
35881 tttcgacctc aaggccaagg cggtcggcca gaacgaccgc catcatggcc ggaatcttgt
35941 tgttccccgc cgcctcgacg gcgaggactg gaacgatgcg gggcttgtcg tcgccgatca
36001 gcgtcttgag ctgggcaaca gtgtcgtccg aaatcaggcg ctcgaccaaa ttaagcgccg
36061 cttccgcgtc gccctgcttc gcagcctggt attcaggctc gttggtcaaa gaaccaaggt
36121 cgccgttgcg aaccaccttc gggaagtctc cccacggtgc gcgctcggct ctgctgtagc
36181 tgctcaagac gcctcccttt ttagccgcta aaactctaac gagtgcgccc gcgactcaac
36241 ttgacgcttt cggcacttac ctgtgccttg ccacttgcgt cataggtgat gcttttcgca
36301 ctcccgattt caggtacttt atcgaaatct gaccgggcgt gcattacaaa gttcttcccc
36361 acctgttggt aaatgctgcc gctatctgcg tggacgatgc tgccgtcgtg gcgctgcgac
36421 ttatcggcct tttgggccat atagatgttg taaatgccag gtttcagggc cccggcttta
36481 tctaccttct ggttcgtcca tgcgccttgg ttctcggtct ggacaattct ttgcccattc
36541 atgaccagga ggcggtgttt cattgggtga ctcctgacgg ttgcctctgg tgttaaacgt
36601 gtcctggtcg cttgccggct aaaaaaaagc cgacctcggc agttcgaggc cggctttccc
36661 tagagccggg cgcgtcaagg ttgttccatc tattttagtg aactgcgttc gatttatcag
36721 ttactttcct cccgctttgt gtttcctccc actcgtttcc gcgtctagcc gacccctcaa

36781 catagcggcc tcttcttggg ctgcctttgc ctcttgccgc gcttcgtcac gctcggcttg
36841 caccgtcgta aagcgctcgg cctgcctggc cgcctcttgc gccgccaact tcctttgctc
36901 ctggtgggcc tcggcgtcgg cctgcgcctt cgctttcacc gctgccaact ccgtgcgcaa
36961 actctccgct tcgcgcctgg tggcgtcgcg ctcgccgcga agcgcctgca tttcctggtt
37021 ggccgcgtcc agggtcttgc ggctctcttc tttgaatgcg cgggcgtcct ggtgagcgta
37081 gtccagctcg gcgcgcagct cctgcgctcg acgctccacc tcgtcggccc gctgcgtcgc
37141 cagcgcggcc cgctgctcgg ctcctgccag ggcggtgcgt gcttcggcca gggcttgccg
37201 ctggcgtgcg gccagctcgg ccgcctcggc ggcctgctgc tctagcaatg taacgcgcgc
37261 ctgggcttct tccagctcgc gggcctgcgc ctcgaaggcg tcggccagct ccccgcgcac
37321 ggcttccaac tcgttgcgct cacgatccca gccggcttgc gctgcctgca acgattcatt
37381 ggcaagggcc tgggcggctt gccagagggc ggccacggcc tggttgccgg cctgctgcac
37441 cgcgtccggc acctggactg ccagcggggc ggcctgcgcc gtgcgctggc gtcgccattc
37501 gcgcatgccg gcgctggcgt cgttcatgtt gacgcgggcg gccttacgca ctgcatccac
37561 ggtcgggaag ttctcccggt cgccttgctc gaacagctcg tccgcagccg caaaaatgcg
37621 gtcgcgcgtc tctttgttca gttccatgtt ggctccggta attggtaaga ataataatac
37681 tcttacctac cttatcagcg caagagttta gctgaacagt tctcgactta acggcaggtt
37741 ttttagcggc tgaagggcag gcaaaaaaag ccccgcacgg tcggcggggg caaagggtca
37801 gcgggaaggg gattagcggg cgtcgggctt cttcatgcgt cggggccgcg cttcttggga
37861 tggagcacga cgaagcgcgc acgcgcatcg tcctcggccc tatcggcccg cgtcgcggtc
37921 aggaacttgt cgcgcgctag gtcctccctg gtgggcacca ggggcatgaa ctcggcctgc
37981 tcgatgtagg tccactccat gaccgcatcg cagtcgaggc cgcgttcctt caccgtctct
38041 tgcaggtcgc ggtacgcccg ctcgttgagc ggctggtaac gggccaattg gtcgtaaatg
38101 gctgtcggcc atgagcggcc tttcctgttg agccagcagc cgacgacgaa gccggcaatg
38161 caggcccctg gcacaaccag gccgacgccg gggcagggg atggcagcag ctcgccaacc
38221 aggaaccccg ccgcgatgat gccgatgccg gtcaaccagc ccttgaaact atccggcccc
38281 gaaacacccc tgcgcattgc ctggatgctg cgccggatag cttgcaacat caggagccgt
38341 ttcttttgtt cgtcagtcat ggtccgccct caccagttgt tcgtatcggt gtcggacgaa
38401 ctgaaatcgc aagagctgcc ggtatcggtc cagccgctgt ccgtgtcgct gctgccgaag
38461 cacggcgagg ggtccgcgaa cgccgcagac ggcgtatccg gccgcagcgc atcgcccagc
38521 atggccccgg tcagcgagcc gccggccagg tagcccagca tggtgctgtt ggtcgccccg
38581 gccaccaggg ccgacgtgac gaaatcgccg tcattccctc tggattgttc gctgctcggc
38641 ggggcagtgc gccgcgccgg cggcgtcgtg gatggctcgg gttggctggc ctgcgacggc
38701 cggcgaaagg tgcgcagcag ctcgttatcg accggctgcg gcgtcggggc cgccgccttg
38761 cgctgcggtc ggtgttcctt cttcggctcg cgcagcttga acagcatgat cgcggaaacc
38821 agcagcaacg ccgcgcctac gcctcccgcg atgtagaaca gcatcggatt cattcttcgg
38881 tcctccttgt agcggaaccg ttgtctgtgc ggcgcgggtg gcccgcgccg ctgtctttgg
38941 ggatcagccc tcgatgagcg cgaccagttt cacgtcggca aggttcgcct cgaactcctg
39001 gccgtcgtcc tcgtacttca accaggcata gccttccgcc ggcggccgac ggttgaggat
39061 aaggcgggca gggcgctcgt cgtgctcgac ctggacgatg gcctttttca gcttgtccgg
39121 gtccggctcc ttcgcgccct tttccttggc gtccttaccg tcctggtcgc cgtcctcgcc
39181 gtcctggccg tcgccggcct ccgcgtcacg ctcggcatca gtctggccgt tgaaggcatc
39241 gacggtgttg ggatcgcggc ccttctcgtc caggaactcg cgcagcagct tgaccgtgcc
39301 gcgcgtgatt tcctgggtgt cgtcgtcaag ccacgcctcg acttcctccg ggcgcttctt
39361 gaaggccgtc accagctcgt tcaccacggt cacgtcgcgc acgcggccgg tgttgaacgc
39421 atcggcgatc ttctccggca ggtccagcag cgtgacgtgc tgggtgatga acgccggcga
39481 cttgccgatt tccttggcga tatcgccttt cttcttgccc ttcgccagct cgcggccaat
39541 gaagtcggca atttcgcgcg gggtcagctc gttgcgttgc aggttctcga taacctggtc
39601 ggcttcgttg tagtcgttgt cgatgaacgc cgggatggac ttcttgccgg cccacttcga
39661 gccacggtag cggcgggcgc cgtgattgat gatatagcgg cccggctgct cctggttctc
39721 gcgcaccgaa atgggtgact tcaccccgcg ctctttgatc gtggcaccga tttccgcgat
39781 gctctccggg gaaaagccgg ggttgtcggc cgtccgcggc tgatgcggat cttcgtcgat
39841 caggtccagg tccagctcga tagggccgga accgccctga gacgccgcag gagcgtccag
39901 gaggctcgac aggtcgccga tgctatccaa ccccaggccg gacggctgcg ccgcgcctgc
39961 ggcttcctga gcggccgcag cggtgttttt cttggtggtc ttggcttgag ccgcagtcat
40021 tgggaaatct ccatcttcgt gaacacgtaa tcagccaggg cgcgaacctc tttcgatgcc
40081 ttgcgcgcgg ccgttttctt gatcttccag accggcacac cggatgcgag ggcatcggcg
40141 atgctgctgc gcaggccaac ggtggccgga atcatcatct tggggtacgc ggccagcagc

40201 tcggcttggt ggcgcgcgtg gcgcggattc cgcgcatcga ccttgctggg caccatgcca

40261 aggaattgca gcttggcgtt cttctggcgc acgttcgcaa tggtcgtgac catcttcttg

40321 atgccctgga tgctgtacgc ctcaagctcg atgggggaca gcacatagtc ggccgcgaag

40381 agggcggccg ccaggccgac gccaagggtc ggggccgtgt cgatcaggca cacgtcgaag

40441 ccttggttcg ccagggcctt gatgttcgcc ccgaacagct cgcgggcgtc gtccagcgac

40501 agccgttcgg cgttcgccag taccgggttg gactcgatga gggcgaggcg cgcggcctgg

40561 ccgtcgccgg ctgcgggtgc ggtttcggtc cagccgccgg cagggacagc gccgaacagc

40621 ttgcttgcat gcaggccggt agcaaagtcc ttgagcgtgt aggacgcatt gccctggggg

40681 tccaggtcga tcacggcaac ccgcaagccg cgctcgaaaa agtcgaaggc aagatgcaca

40741 agggtcgaag tcttgccgac gccgcctttc tggttggccg tgaccaaagt tttcatcgtt

40801 tggtttcctg tttttcttg gcgtccgctt cccacttccg gacgatgtac gcctgatgtt

40861 ccggcagaac cgccgttacc cgcgcgtacc cctcgggcaa gttcttgtcc tcgaacgcgg

40921 cccacacgcg atgcaccgct tgcgacactg cgcccctggt cagtcccagc gacgttgcga

40981 acgtcgcctg tggcttccca tcgactaaga cgccccgcgc tatctcgatg gtctgctgcc

41041 ccacttccag cccctggatc gcctcctgga actggctttc ggtaagccgt ttcttcatgg

41101 ataacaccca taatttgctc cgcgccttgg ttgaacatag cggtgacagc cgccagcaca

41161 tgagagaagt ttagctaaac atttctcgca cgtcaacacc tttagccgct aaaactcgtc

41221 cttggcgtaa caaaacaaaa gcccggaaac cgggctttcg tctcttgccg cttatggctc

41281 tgcacccggc tccatcacca acaggtcgcg cacgcgcttc actcggttgc ggatcgacac

41341 tgccagccca acaaagccgg ttgccgccgc cgccaggatc gcgccgatga tgccggccac

41401 accggccatc gcccaccagg tcgccgcctt ccggttccat tcctgctggt actgcttcgc

41461 aatgctggac ctcggctcac cataggctga ccgctcgatg gcgtatgccg cttctcccct

41521 tggcgtaaaa cccagcgccg caggcggcat tgccatgctg cccgccgctt tcccgaccac

41581 gacgcgcgca ccaggcttgc ggtccagacc ttcggccacg gcgagctgcg caaggacata

41641 atcagccgcc gacttggctc cacgcgcctc gatcagctct tgcactcgcg cgaaatcctt

41701 ggcctccacg gccgccatga atcgcgcacg cggcgaaggc tccgcagggc cggcgtcgtg

41761 atcgccgccg agaatgccct tcaccaagtt cgacgacacg aaaatcatgc tgacggctat

41821 caccatcatg cagacggatc gcacgaaccc gctgaattca cccccgaaca cgagcacggc

41881 acccgcgacc actatgccaa gaatgcccaa ggtaaaaatt gccggccccg ccatgaagtc

41941 cgtgaatgcc ccgacggccg aagtgaaggg caggccgcca cccaggccgc cgccctcact

42001 gcccggcacc tggtcgctga atgtcgatgc cagcacctgc ggcacgtcaa tgcttccggg

42061 cgtcgcgctc gggctgatcg cccatcccgt tactgccccg atcccggcaa tggcaaggac

42121 tgccagcgcc gcgatgagga agcgggtgcc ccgcttcttc atcttcgcgc ctcgggcctc

42181 gaggccgcct acctgggcga aaacatcggt gtttgtggca ttcatacgga ctcctgttgg

42241 gccagctcgc gcacgggctg gcgggtcagc ttggcttgaa gatcgccacg cattgcggcg

42301 atctgcttct cggcatcctt gcgcttctgc acgccttcct gctggatgcg aataacgtcc

42361 tcgacggtct tgatgagcgt cgtctgaacc tgcttgagcg tgtccacgtc gatcaccagg

42421 cgttggttct ccttcgccgt ctcgacggac gtgcgatgca gcagggccgc attgcgcttc

42481 atcaggtcgt tggtggtgtc gtcgatggcc gtggccagtt cgacggcgtt cttctgctcg

42541 ttgaggctca aggccagcat gaattgccgc ttccacgccg gcacggtgat ttcgcggatg

42601 gtgtggaatt tatcgaccag catctggttg ttggcctgga tcatgcggat ggtcggcagg

42661 ctctgcatgg ccgaatgttg caaggcgatc aggtcgccga tgcgcttgtc caggttggca

42721 accatcgcat cgaggtcggc cagctcctgc acgcggcccg ggtcgttccc gacattgccg

42781 cgcagaccct cggcctgctc gcgcagctcg gcaaggcgga ccttgccggc cgcgatgtgg

42841 acgccaagaa ggcggtgttc ctcgcgcacg gctgcgaaca tttcgtcgag cgaggcattg

42901 cgctgcgcga tgccttgctg ggtggtctgc acttcgctga ccaggtgttc gatctgctcg

42961 cgggtcgtgt cgaagcgcgc catgaagccc gtcgaacgga cgcggaagcg gtcgatcagc

43021 gggccaatca ggggcaggcg ggaacggttg tcggacaaag ggccgacgtt cagggaacgg

43081 gccttggcga caacctgggt cagtttctcg cctgcttcgt ccaggtcgct gttgcgcacc

43141 tggtccagca ggctatcggc gtagcgggac gtgtgctcgg ccacgtcgcg gccgaactcg

43201 gcaacggtct gcggactgcc gacctcgatc cgctgcgcga ccgcatggac ttccggcacg

43261 tcgctttcct gcaagcccag ctcgcgcagg gttgccgggg tcatgtcgaa ggcgacgata

43321 ggggccttgg cgtcgtgcgt cgttttcagt gcgttcatag ggttctcccg ccgtgttatt

43381 ggttgatgcc ttccaggctc tgcgaaaggc tccgcatgag cgcctggtga gctttggccg

43441 cctcggcgac cattgccgga ttcatgttct tggtggtgat gagcgcgagg gtgtgctgac

43501 gccagacggg caccaggacg gatgccgttt cagagaagcg gtccagcatg tccacggcct

43561 gcgcccgcgt gagcttcatc tgagtgacgc tcatttcatg ggacgccatg agggttgcca

43621 ggttggcgag cttgcgcgcg aagcgttcgc gcggcttgtc gaactcgatc acgccggcct
43681 tggccgcgcc ggcctcgggg ttctcgtcca ggaactcgcg cccggcttga atgtaggctc
43741 tgagccggtc tacctcggcc tcatgcgtat tgagcatgtc atccaaggcg cgcaacgtgt
43801 cccgcacgcg ctgcgctacg ccctcggctt cgtccagcaa ctggtcgagc gtcttgcggg
43861 cgacctgata cctcacctgg cgttcaacct cacggccaag catcttctcg aaccaggtag
43921 gcttttccgc gatcttgcgg gggtccgcgt cggccagctt cgccacgatc tggctgattt
43981 tgtcggccag cgcggcaact gcgccgtgct ccatcagatt cgacagctcg ttgagggaat
44041 ccgccccgtc gatgccggcc ccgtactcgc caatcgtcgc cggcgacgcg aagagggcgg
44101 gcaaaacctc ccccttcaat cgcgccatgt tcacgctttg ttcttccatt cgatacaccc
44161 tcgcggtggg ttaattgctt ttcgatggaa gaagtttagc taaactttct atccctcgtc
44221 aacaccttta gccgctaaaa tttggggaca ggtcatttac agaaagccag ctcactcctg
44281 gcgttgcccc ttgagcgccg ctaggcgcgc agcatccttc gcgctgagaa agaacgtcat
44341 cagcggcccg accgtcttgc ttgaaccgtc ggcaaagcaa acatccatcg aacagccttg
44401 cgtgtggggg tccacgcctt cgaccagttt ccaagggtcc atgccccagc cctcggcctc
44461 cggattgaac cagtacgccg atgcgtcgcc gtttaggtcg ctgtcggcgt agtccttgac
44521 cagcacggcc acccgcccgg tcgtcgggca cacgtagccc ggctgcttag gttcctgtct
44581 tggcattgct caaagctcct tgaaggggcc gctctacagc cccttgggct tgtagagcga
44641 cacgaaatag gtgagtgcgg tcagtaccgc gaaatgcacc aggaacgtcc agccggcatg
44701 aacgccaagg gtgttccagt ggtacagcat ccgcaggaac tgaaagaaaa cgtcgataga
44761 gatgatccac ttcgccaccg gccacaccag gacagtaacg acccatacaa agcggaccag
44821 ggcctggaca cccttggcaa aagtgaaccg gggcggggcc ttgctcgggg cctcaacgcg
44881 cggggcaggg gcctcggcct ccacttccac gcctgggaac ttgataatct tcgacattgc
44941 ttgaccctcc acggcgatgc gtgttcaatt cgtccagcgc tcgcgcgcct agaccgtgat
45001 gtgacagcat cgaggtcaag cgccccggag aaatccgggg cgtcatccct atgccccgtc
45061 caactcggga accggctttt ccctggtgct caacctggcc ggctcgaccc acttggtgac
45121 ttgctgccac tcgttaccac tgcgaacggc tacccgaatc tgcacacgtt tagccgcgat
45181 cttcgtcact atgccagcaa cacaaacgga atacccgtac ccgccgcgcg gggtgtgctg
45241 ccagttcacc ctatctccta cttgcatcat catcccctgg cgtcagtgac cggcccggaa
45301 tttcgccaag tcgatttcgt tgaaggtcca gcgctgtttc gcccgttcct ccaacctcga
45361 cgactcccgc atgacctcga tgcgcaggcg ctcgacctgg tgcatcagct catcagcgcg
45421 ccgctgcttc tcggcaatag cattccgctg cgcgaccagc tcctggtcta cgttcggcag
45481 ctcgtcgatc cacggcatga acttatcggt catcggattg gcctccggta attgacctgg
45541 gaatctaccc ggcctcaaaa caagaatagg gcataatgcc ctaacttgtc aagcaatttt
45601 agctaaacaa ttgaggggat tcagcgaggc gtcatgcttg aaaacacctt tcccctggcg
45661 tgcaatcagc ttgtcccggc ggcagcgcac tgccgcagcg cggccagcaa ggtgccttgc
45721 tcgatccgct gtcgttcctc cggcgtaatg caatcgccgc agataccgcc cagctcttcg
45781 cgcagggcgt ccgccccgat caggctgcgg ctaagggtcg aaatggactt accgcagcgg
45841 cggcaattgg tggtgacaat ctcgatcttg cggttgggca tggccctatc tccttgagag
45901 aggcccgacc gtagccgggc ctcgttccgt taccagcctg cgcgagcctt cgcgcgttcg
45961 acttcctggg tggaccagtg gccctttgct tcaccctcca gggtgcaacc ttgcaggtac
46021 gccttgtgcc gcagctcttc ggcctggcgg gtaagctctg ccgcctgctc catcagcgcc
46081 gcgatctggt cacgacgggc cagaaaatcc gtggcggccg cgccgggcag ctcatcgagc
46141 caagacatga tcttgctttt catcggggtt gatcctccgg ttgctgacct gggcgaagtg
46201 cccggccttg gattgctata ttagggcatt atgccctaga tagtcaagga aatttagcta
46261 aacaatttgc ggcgggcgca cgaaaaaacc cggcttgcgg gccgggctgc tggcagcata
46321 tcgcaacgat cagggttggc ggttttagcc gctaaagtcc tctcccttgg cgtaaagtcc
46381 tgcgggcgtc agccctggcc tttccagatc gccccaatcc ccgctagatc gcaaaggatc
46441 gcccaggcgg catagggggat cggcgaatcc tcgccaaccc aacgccgcac cgtgcggtcg
46501 cccttggcac ccaagcccaa gatgcgcgca gcctgtccgc cggtgaggcc ggccaagtgc
46561 aagacttccc ggatttctgc gccggtcggc tgcacccagc gttccgccgg gcgcaggcac
46621 tcaagccgga tattcacgtc gctcatgctg cttttctcct aatcgttatc aaatggcggc
46681 ccgcaattgg tcgtagccgt agcacgactc gatgcaacgc gggtcatatt cacaaacctt
46741 tctaccgttg cgattgatgc gatagcgcgg cggcttgtcg tcgtgcgccc acacctgtcg
46801 cgtttcggga ctgatcttcg taatgatgac gtgcttcccg atttccttgg caaaaaccgg
46861 gtgatgcacg ctcacaatct cgcaccgcag gccaggaaag aacgtcgccg gccacgccgg
46921 cgcgtcctcg tccttggccg gctccggctc cggcttggcc ggtgtaaccg gctccctgcg
46981 ctgcccggcc ggctcctgag ctttcgcacg cgcggccgcg agcttcgcct tggcgctggc

47041 cgcccatttg cgggcgatga atgcctggtg cgccggcagc acggcatcaa cccaggcgaa
47101 gcccggcggc agttcgtccg gcttccgcca gtcctcgacc acggcccgca cgcggcgcgg
47161 cggtggcgtg atacccgcga gccacgccgg cagtggctcg tcggcaaggc ggtcctggtc
47221 gcgggcgtcc agataccggc gatgcgagcg caacagggcg cgcaaaaccc cgtccgctac
47281 ctggtccacc gtcatgccgc cgcgcgcatg gtcgtaatgg gaccgatagc ccgtttcgga
47341 aataaaaggg gtctgacgct cagtggaacg aaaactcacg ttaagggatt ttggtcatga
47401 gattatcaaa aaggatcttc acctagatcc ttttaaatta aaaatgaagt tttaaatcaa
47461 tctaaagtat atatgagtaa acttggtctg acagttacca atgcttaatc agtgaggcac
47521 ctatctcagc gatctgtcta tttcgttcat ccatagttgc ctgactcccc gtcgtgtaga
47581 taactacgat acgggagggc ttaccatctg gccccagtgc tgcaatgata ccgcgagatc
47641 cacgctcacc ggctccagat ttatcagcaa taaaccagcc agccggaagg gccgagcgca
47701 gaagtggtcc tgcaacttta tccgcctcca tccagtctat taattgttgc cgggaagcta
47761 gagtaagtag ttcgccagtt aatagtttgc gcaacgttgt tgccattgct gcaggcatcg
47821 tggtgtcacg ctcgtcgttt ggtatggctt cattcagctc cggttcccaa cgatcaaggc
47881 gggttacatg atcccccatg ttgtgcaaaa aagcggttag ctccttcggt cctccgatcg
47941 ttgtcagaag taagttggcc gcagtgttat cactcatggt tatggcagca ctgcataatt
48001 ctcttactgt catgccatcc gtaagatgct tttctgtgac tggtgagtac tcaaccaagt
48061 cattctgaga atagtgtatg cggcgaccga gttgctcttg cccggcgtca acacgggata
48121 ataccgcacc acataTGTTG ATACAACCAT AAAATGATAA TTACACCCAT AAATTGATAA
48181 TTATCACACC CATAAATTGA TATTGCCTCT TCATGGTCTA AACTTCAGTA AGTTTACGAC
48241 ATTTTCCTCG AGGTCATTTC ccaattattg aaggccgcta acgcggcctt tttttgtttc
48301 tggtctgcct cctCCCTGCA GGCTTCAACA AACAGacaat ctggtctgtt tgtattatgg
48361 aaaatttttc tgtataatag attcaacaaa cagacaatct ggtctgtttg tattatcgac
48421 CAACCCACTC CCATGGTGTG ACGGGCGGTG TGTACAAGGC CCGGGAACGT gTTCACAAAA
48481 GTTATCAGGC ATGCACCTGG TAGCTAGTCT TTAAACCAAT AGATTGCATC GGTTTAAAAG
48541 GCAAGACCGT CAAATTGCGG GAAAGGGGTC AACAGCCGTT CAGTACCAAG TCTCAGGGGA
48601 AACTTTGAGA TGGCCTTGCA AAGGGTATGG TAATAAGCTG ACGGACATGG TCCTAACCAC
48661 GCAGCCAAGT CCTAAGTCAA CAGATCTTCT GTTGATATGG ATGCAGTTCA CAGACTAAAT
48721 GTCGGTCGGG GAAGATGTAT TCTTCTCATA AGATATAGTC GGACCTCTCC TTAATGGGAG
48781 CTAGCGGATG AAGTGATGCA ACACTGGAGC CGCTGGGAAC TAATTTGTAT GCGAAAGTAT
48841 ATTGATTAGT TTTGGAGTAC TCGATGGTGT TCAATGCTTT TCCCGTTATC CGGATCACAT
48901 GAAACGGCAT GACTTTTTCA AGAGTGCCAT GCCCGAAGGT TATGTACAGG AACGCACTAT
48961 ATCTTTCAAA GATGACGGGA CCTACAAGAC GCGTGCTGAA GTCAAGTTTG AAGGTGATAC
49021 CCTTGTTAAT CGTATCGAGT TAAAGGGTAT TGATTTTAAA GAAGATGGAA ACATTCTTGG
49081 ACACAAACTC GAGTACAACT TTAACTCACA CAATGTATAC ATCACGGCAG ACAAACAAAA
49141 GAATGGAATC AAAGCTAACT TCAAAATTCG CCACAACGTT GAAGATGGTT CCGTTCAACT
49201 AGCAGACCAT TATCAACAAA ATACTCCAAT TGGCGATGGC CCTGTCCTTT TACCAGACAA
49261 CCATTACCTG TCGACACAAT CTGTCCTTTC GAAAGATCCC AACGAAAAGC GTGACCACAT
49321 GGTCCTTCTT GAGTTTGTAA CTGCTGCTGG GATTACACAT GGCATGGATG AGCTCTACAA
49381 ATGGCccaat tattgaaggc ctccctaacg gggggccttt ttttGTTTCT GGTCTGCCGC
49441 TGCGCTTACT GCAGTAGTTT TGCTGAAATA CTCGATTCAC AAAAATATCA ACTTATGGTT
49501 GTTTTGTGAG ATATCAATAT ATGGTTGTTT TGTGGTTAAG TTGCTGATTA TAAATAATTA
49561 TTAAATATCA CTTTATGGTT GCATCAACAa catagcagaa ctttaaaagt gctcatcatt
49621 ggaaaacgtt cttcggggcg aaaactctca aggatcttac cgctgttgag atccagctcg
49681 atgtaaccca ctcgtgcacc caacttatct tcagcatctt ttactttcac cagcgtttct
49741 gggtgagcaa aaacaggaag gcaaaatgcc gcaaaaaagg gaataagggc gacacggaaa
49801 tgttgaatac tcatactctt cctttttcaa tattattgaa gcatttatca gggttattgt
49861 ctcatgagcg gatacatatt tgaatgtatt tagaaaaata aacaaatagg ggttccgcgc
49921 acatttcccc gaaaagtgcc acctgacgtc taagaaacca ttattatcat gacattaacc
49981 tataaaaata agcgtatcac gaggcccttt cgtcttcaag aattttataa accgtggagc
50041 gggcaatact gagctgatga gcaatttccg ttgcaccagt gcccttctga tgaagcgtca
50101 gcacgacgtt cctgtccacg gtacgcctgc ggccaaattt gattcctttc agctttgctt
50161 cctgtcggcc ctcattcgtg cgctctagga tcctccggcg ttcagcttgt gccacagccg
50221 acaggatggt gaccaccatt tgccccatat caccgtcggt actgatcccg tcgtcaataa
50281 accgaaccgc gacaccctga gcatcaaact cttttatcag ttggatcatg tcggcggtgt
50341 cgcggccaag acggtcgagc ttcttcacca gaatgacatc accttcctcc accttcatcc
50401 tcagcaaatc cagcccttcc cgatctgttg aactgccgga tgccttgtcg gtaaagatgc

50461 ggttagcttt taccccctgca tctttgagcg ctctgatctg aatatcgagg gactgctggc
50521 tggttgagac ccgcgcataa ccaaaaattc gcataaaatg taccttaaat cgaatatcag
50581 acacgatgtg tctattatgc caaaatgacg atttaatgga cactcaaacg aagccgtttt
50641 actatgtctg ataatttata acatttcgga cggttgcaaa aatgttacta aatgcccgtc
50701 aggcagggag gccgatatgc ccgttgactt tctgaccact gagcagactg aaagctatgg
50761 cagattcacc ggtgaaccgg atgagcttca gctggcacga tattttcatc ttgatgaagc
50821 agacaaggaa tttatcggaa aaagcagagg tgatcacaac cgtctgggca ttgccctgca
50881 aattggatgt gtccgttttc tgggcacctt cctcaccgat atgaatcata ttccttccgg
50941 cgtccggcat tttaccgcca gacagctcgg gattcgtgat atcaccgttc ttgcagaata
51001 cggtcagagg gaaaataccc gccgtgagca tgcagcgctg atacgtcagc actatcagta
51061 tcgtgaattt gcctggccct ggacatttcg ccttacccgt cttttatata cccggagctg
51121 gataagcaac gaacgtcctg gcctgctttt cgatctggcg acagggtggc ttatgcaaca
51181 tcgtattatt ctccccggag ccactacgct gacccggttg atttcagagg taagggaaaa
51241 ggcgacgttg cgcctgtgga acaaactggc actgataccg tcagccgaac aacgttcaca
51301 gctggagatg ctgctggggc caactgattg cagccgcctg tctttactgg aatcactgaa
51361 aaaaggccct gtgaccatca gtggtccggc gtttaatgaa gcaattgaac gctggaaaac
51421 tctgaacgat tttggcttgc atgctgacaa cctgagtaca ctcccggctg tgcgcctgaa
51481 aaatctcgca cgttatgctg gtatgacttc ggtgttcaat attgccagga tgtcaccgca
51541 gaaaaggatg gcggttctgg ttgcctttgt ccttgcatgg gaaacgctgg cgttggatga
51601 tgcactggac gttctggacg ccatgctggc cgttatcatc cgtgacgcca gaaagattgg
51661 gcagaaaaaa cggctccgct cgctgaagga tctggataaa tctgcattgg cgctcgccag
51721 cgcatgttcg tacctgctga aagaagaaac accggacgaa tcgattcgtg ctgaggtgtt
51781 cagctacatc cccaggcaaa agctggctga aatcatcacg cttgtccgtg aaattgcccg
51841 gccctcagac gataattttc atgaagaaat ggtggagcag tacgggcgcg ttcgtcgttt
51901 cctgccccat ctgctgaata ccgttaaatt ttcatccgca cctgccgggg ttaccactct
51961 gaatgcctgt gactacctca gccgggagtt cagctcacgg cggcagtttt ttgacgacgc
52021 accaacggaa attatcagtc ggtcatggaa acggctggtg attaacaagg aaaaacatat
52081 cacccgcagg ggatacacgc tctgctttct cagtaaactg caggatagtc tgaggcggag
52141 ggatgtctac gttaccggca gtaaccggtg gggagatccc cgagcaagat tactacaggg
52201 tgctgactgg caggcaaacc ggattaaggt ttatcgttct ctgggacacc cgacagaccc
52261 gcaggaagca ataaaatctc tgggtcatca gcttgatagt cgttacagac aggttgctgc
52321 acgtcttggc gaaaatgagg ctgtcgaact cgatgtttct ggcccgaagc cccggttgac
52381 aatttctccc ctcgccagtc ttgatgagcc ggacagtctg aaacgactga gcaaaatgat
52441 cagtgatctg ctccctccgg tggatttaac ggagttgctg ctcgaaatta acgcccatac
52501 cggatttgct gatgagtttt tccatgctag tgaagccagt gccagagttg atgatctgcc
52561 cgtcagcatc agcgccgtgc tgatggctga agcctgcaat atcggtctgg aaccactgat
52621 cagatcaaat gttcctgcac tgacccgaca ccggctgaac tggacaaaag cgaactatct
52681 gcgggctgaa actatcacca gcgctaatgc cagactggtt gattttcagg caacgctgcc
52741 actggcacag atatggggtg gcggagaagt ggcatctgca gatggaatgc gctttgttac
52801 gccagtcaga acaatcaatg ccggaccgaa ccgcaaatac tttggtaata acagagggat
52861 cacctggtac aactttgtgt ccgatcagta ttccggcttt catggcatcg ttataccggg
52921 gacgctgagg gactctatct ttgtgctgga aggccttctg gaacaggaga ccgggctgaa
52981 tccaaccgaa attatgaccg atacagcagg taccagcgaa cttgtctttg gcctttttctg
53041 gctgctggga taccagtttt ctccacgcct ggctgatgcc ggtgcttcgg ttttctggcg
53101 aatgggccat gatgccaact atggcgtgct gaatgatatt gccagagggc aatcagatcc
53161 ccgaaaaata gtccttcagt gggacgaaat gatccggacc gctggctccc tgaaactggg
53221 caaagtacag gcttcagtgc tggtccgttc attgctgaaa agtgaacgtc cttccggact
53281 gactcaggca atcattgaag tggggcgcat caacaaaacg ctgtatctgc ttaattatat
53341 tgatgatgaa gattaccgcc ggcgcattct gacccagctt aatcggggag aaagccgcca
53401 tgccgttgcc agagccatct gtcacggtca aaaaggtgag ataagaaaac gatataccga
53461 cggtcaggaa gatcaactgg gcgcactggg gctggtcact aacgccgtcg tgttatggaa
53521 cactatgtat atgcaggcag ccctggatca tctccgggcg caaggtgaaa cactgaatga
53581 tgaagatatc gcacgcctct ccccgctttg ccacggacat atcaatatgc tcggccatta
53641 ttccttcacg ctggcagaac tggtgaccaa aggacatctg agaccattaa aagaggcgtc
53701 agaggtagaa aacgttgctt aacgtgagtt ttcgttccac tgagcgtcag acccctaaaa
53761 gggccgtcca ggtccacggc gtggaactgg aaatgcaccg tggttaggcc gccgtagccg
53821 ctttcgacct ccacccatac gcgcatcccg tccacggaag ccaggaagtc gccttcctgg

53881 ccccacatcg ggacgacgcc cggcgtcgcg cggcaatggc gctctatgac cctctcggcc
53941 gcatcctggt cgccggcgca accgaagaac gtgcccccgt tgagcttcca caccgccgcc
54001 gcgtaacgct cgccggccac gtcggccacc aaatcggcac gattgagcac tgcgtcatgc
54061 accgccgcta ccgcgtccgc cgcaacggcc agcaggccgg cgcgatcctc gggcagcgtc
54121 gccgccagcg ctgcaatgcg cgcgtccttg tcctcggtct gcggcttgct tttgcgcgcc
54181 ttcgtccgcc ggcgagccgc cggggtggtc gcctcgggcg gcgtcgggtt ggctaccagg
54241 taggccgcga tgcaatccgg tttccggtag tcatcgcaga acttcaagcc gtccaggacg
54301 cagcaggtcg gcaacgggta ctgcatccag tattcgggtt ccgggaacgg atgcagcaac
54361 tcgccgcagt ggtggcatcg ctctgtctcg cggtcctgca ccgaaccgct gaacaagcgg
54421 ccggcaacga tggcatcgtt gcggcgcttg atgttgtcgt cttgcatgct cgcctccaac
54481 ggcccccagc ctttcggccg gggcctgcc ccttagtcga tggcccggta gatcgccgcc
54541 gcctcggcgt gctcgctcgc aaacgccgcc aggtggtggt agcggtcgat cagggcgtcc
54601 gcgtcggccg tgccggccag ctccgcgcag agctggttca gggcgaagag cgtggcgacg
54661 atgccggcgg cgtcggccga cagctcgccc gcgaagccgt tgccatcgac ctcgatgcgc
54721 aagcggccgt catgctccgg ggccatgtag aagccaccgt tcgagagggt gtagaagtgc
54781 cagaaaccgc cgtggtaggc agcgcaaagg cggcccatcc agccgaatac cagggcctcg
54841 ccgcgcagca tgcgcagcat cgagggggccg aagtacgtcg gcagaaaccg caggcgatcc
54901 gcctcggcga cgcgggtggc aacgataggg gaggggattt gaacatcagt catcggactt
54961 tctccggtag ttgacctggg cggaatgccc ggcctcaaga ccaatattag ggcattatgc
55021 cctagcctgt caagcaaatt tagctaaact atgccgccgg ccgtacccga ttctgcggtt
55081 acagctcggt cctggtcccg accgggcttt gccaggcggg cggatggaac gccaaggggg
55141 agcgcggccg ttgaatggtg atcgccatca cgtttcattg acacttgagg ggcgtttaga
55201 gcgagccagg aaagccgacc ccctccttgg agtaaaaacc cttgcggcgt tgcagccggc
55261 acggatcttc cgatcgggcg cggtggtggc cgcgtctgtg acctaaaaag gggggagtcc
55321 agagggggcgc agccccttttg ggcatagcgc agcgtaatcg gagacgtaat tgagcatttc
55381 caggcgcttg cgcctggtca acgaaagagt cagcgccgta ggcgctgcca tttttggggt
55441 gaggccgttc gcggccgagg ggcgcagccc ctggggggat gggaggcccg cgttagcggg
55501 ccgggagggt tcgagaaggg ggggcacccc ccttcggcgt gcgcggtcac gcgcacaggg
55561 cgcagccctg gttaaaaaca aggtttataa atattggttt aaaagcaggt taaaagacag
55621 gttagcggtg gccgaaaaac gggcggaaac ccttgcaaat gctggatttt ctgcctgtgg
55681 acagcccctc aaatgtcaat aggtgcgccc ctcatctgtc agcactctgc ccctcaagtg
55741 tcaaggatcg cgcccctcat ctgtcagtag tcgcgcccct caagtgtcaa taccgcaggg
55801 cacttatccc caggcttgtc cacatcatct gtgggaaact cgcgtaaaat caggcgtttt
55861 cgccgatttg cgaggctggc cagctccacg tcgccggccg aaatcgagcc tgcccctcat
55921 ctgtcaacgc cgcgccgggt gagtcggccc ctcaagtgtc aacgtccgcc cctcatctgt
55981 cagtgagggc caagttttcc gcgaggtatc cacaacgccg gcggccgcgg tgtctcgcac
56041 acggcttcga cggcgtttct ggcgcgtttg cagggccata gacggccgcc agcccagcgg
56101 cgagggcaac cagcccggtg agcgtcggaa aggcgctgga agccccgtag cgacgcggag
56161 aggggcgaga caagccaagg gcgcaggctc gatgcgcagc acgacatagc cggttctcgc
56221 aaggacgaga atttccctgc ggtgcccctc aagtgtcaat gaaagtttcc aacgcgagcc
56281 attcgcgaga gccttgagtc cacgctagat ctatctcatc tgcgcaaggc agaacgtgaa
56341 gacggccgcc ctggacctcg cccgcgagcg ccaggcgcac gaggccggcg cgcggacccg
56401 cgccacggcc cacgagcgga cgccgcagca ggagcgccag aaggccgcca gagaggccga
56461 gcgcggccgt gaggcttgga cgctagggca gggcatgaaa aagcccgtag cgggctgcta
56521 cgggcgtctg acgcggtgga aagggggagg ggatgttgtc tacatggctc tgctgtagtg
56581 agtgggttgc gctccggcag cggtcctgat caatcgtcac cctttctcgg tccttcaacg
56641 ttcctgacaa cgagcctcct tttcgccaat ccatcgacaa tcaccgcgag tccctgctcg
56701 aacgctgcgt ccggaccggc ttcgtcgaag gcgtctatcg cggcccgcaa cagcggcgag
56761 agcggagcct gttcaacggt gccgccgcgc tcgccggcat cgctgtcgcc ggcctgctcc
56821 tcaagcacgg ccccaacagt gaagtagctg attgtcatca gcgcattgac ggcgtccccg
56881 gccgaaaaac ccgcctcgca gaggaagcga agctgcgcgt cggccgtttc catctgcggt
56941 gcgcccggtc gcgtgccggc atggatgcgc gcgccatcgc ggtaggcgag cagcgcctgc
57001 ctgaagctgc gggcattccc gatcagaaat gagcgccagt cgtcgtcggc tctcggcacc
57061 gaatgcgtat gattctccgc cagcatggct tcggccagtg cgtcgagcag cgcccgcttg
57121 ttcctgaagt gccagtaaag cgccggctgc tgaaccccca accgttccgc cagtttgcgt
57181 gtcgtcagac cgtctacgcc gacctcgttc aacaggtcca gggcggcacg gatcactgta
57241 ttcggctgca actttgtcat gcttgacact ttatcactga taaacataat atgtccacca

57301 acttatcagt gataaagaat ccgcgcgttc aatcggacca gcggaggctg gtccggaggc
57361 cagacgtgaa acccaacata cccctgatcg taattctgag cactgtcgcg ctcgacgctg
57421 tcggcatcgg cctgattatg ccggtgctgc cgggcctcct gcgcgatctg gttcactcga
57481 acgacgtcac cgcccactat ggcattctgc tggcgctgta tgcgttggtg caatttgcct
57541 gcgcacctgt gctgggcgcg ctgtcggatc gtttcgggcg gcggccaatc ttgctcgtct
57601 cgctggccgg cgccactgtc gactacgcca tcatggcgac agcgcctttc ctttgggttc
57661 tctatatcgg gcggatcgtg gccggcatca ccggggcgac tggggcggta gccggcgctt
57721 atattgccga tatcactgat ggcgatgagc gcgcgcggca cttcggcttc atgagcgcct
57781 gtttcgggtt cgggatggtc gcgggacctg tgctcggtgg gctgatgggc ggtttctccc
57841 cccacgctcc gttcttcgcc gcggcagcct tgaacggcct caatttcctg acgggctgtt
57901 tccttttgcc ggagtcgcac aaaggcgaac gccggccgtt acgccgggag gctctcaacc
57961 cgctcgcttc gttccggtgg gcccggggca tgaccgtcgt cgccgccctg atggcggtct
58021 tcttcatcat gcaacttgtc ggacaggtgc cggccgcgct ttgggtcatt ttcggcgagg
58081 atcgctttca ctgggacgcg accacgatcg gcatttcgct tgccgcattt ggcattctgc
58141 attcactcgc ccaggcaatg atcaccggcc ctgtagccgc ccggctcggc gaaaggcggg
58201 cactcatgct cggaatgatt gccgacggca caggctacat cctgcttgcc ttcgcgacac
58261 ggggatggat ggcgttcccg atcatggtcc tgcttgcttc gggtggcatc ggaatgccgg
58321 cgctgcaagc aatgttgtcc aggcaggtgg atgaggaacg tcaggggcag ctgcaaggct
58381 cactggcggc gctcaccagc ctgacctcga tcgtcggacc cctcctcttc acggcgatct
58441 atgcggcttc tataacaacg tggaacgggt gggcatggat tgcaggcgct gccctctact
58501 tgctctgcct gccggcgctg cgtcgcgggc tttggagcgg cgcagggcaa cgagccgatc
58561 gctgatcgtg gaaacgatag gcctatgcca tgcgggtcaa ggcgacttcc ggcaagctat
58621 acgcgcccta ggagtgcggt tggaacgttg gcccagccag atactcccga tcacgagcag
58681 gacgccgatg atttgaagcg cactcagcgt ctgatccaag aacaaccatc ctagcaacac
58741 ggcggtcccc gggctgagaa agcccagtaa ggaaacaact gtaggttcga gtcgcgagat
58801 cccccggaac caaaggaagt aggttaaacc cgctccgatc aggccgagcc acgccaggcc
58861 gagaacattg gttcctgtag gcatcgggat tggcggatca aacactaaag ctactggaac
58921 gagcagaagt cctccggccg ccagttgcca ggcggtaaag gtgagcagag gcacgggagg
58981 ttgccacttg cgggtcagca cggttccgaa cgccatggaa accgcccccg ccaggcccgc
59041 tgcgacgccg acaggatcta gcgctgcgtt tggtgtcaac accaacagcg ccacgcccgc
59101 agttccgcaa atagccccca ggaccgccat caatcgtatc gggctaccta gcagagcggc
59161 agagatgaac acgaccatca gcggctgcac agcgcctacc gtcgccgcga ccccgcccgg
59221 caggcggtag accgaaataa acaacaagct ccagaatagc gaaatattaa gtgcgccgag
59281 gatgaagatg cgcatccacc agattcccgt tggaatctgt cggacgatca tcacgagcaa
59341 taaacccgcc ggcaacgccc gcagcagcat accggcgacc cctcggcctc gctgttcggg
59401 ctccacgaaa acgccggaca gatgcgcctt gtgagcgtcc ttggggccgt cctcctgttt
59461 gaagaccgac agcccaatga tctcgccgtc gatgtaggcg ccgaatgcca cggcatctcg
59521 caaccgttca gcgaacgcct ccatgggctt tttctcctcg tgctcgtaaa cggacccgaa
59581 catctctgga gctttcttca gggccgacaa tcggatctcg cggaaatcct gcacgtcggc
59641 cgctccaagc cgtcgaatct gagccttaat cacaattgtc aattttaatc ctctgtttat
59701 cggcagttcg tagagcgcgc cgtgcgtccc gagcgatact gagcgaagca agtgcgtcga
59761 gcagtgcccg cttgttcctg aaatgccagt aaagcgctgg ctgctgaacc cccagccgga
59821 actgacccca caaggcccta gcgtttgcaa tgcaccaggt catcattgac ccaggcgtgt
59881 tccaccaggc cgctgcctcg caactcttcg caggcttcgc cgacctgctc gcgccacttc
59941 ttcacgcggg tggaatccga tccgcacatg aggcggaagg tttccagctt gagcgggtac
60001 ggctcccggt gcgagctgaa atagtcgaac atccgtcggg ccgtcggcga cagcttgcgg
60061 tacttctccc atatgaattt cgtgtagtgg tcgccagcaa acagcacgac gatttcctcg
60121 tcgatcagga cctggcaacg ggacgttttc ttgccacggt ccaggacgcg gaagcggtgc
60181 agcagcgaca ccgattccag gtgcccaacg cggtcggacg tgaagcccat cgccgtcgcc
60241 tgtaggcgcg acaggcattc ctcggccttc gtgtaatacc ggccattgat cgaccagccc
60301 aggtcctggc aaagctcgta gaacgtgaag gtgatcggct cgccgatagg ggtgcgcttc
60361 gcgtactcca acacctgctg ccacaccagt tcgtcatcgt cggcccgcag ctcgacgccg
60421 gtgtaggtga tcttcacgtc cttgttgacg tggaaaatga ccttgttttg cagcgcctcg
60481 cgcgggattt tcttgttgcg cgtggtgaac agggcagagc gggccgtgtc gtttggcatc
60541 gctcgcatcg tgtccggcca cggcgcaata tcgaacaagg aaagctgcat ttccttgatc
60601 tgctgcttcg tgtgtttcag caacgcggcc tgcttggcct cgctgacctg ttttgccagg
60661 tcctcgccgg cggtttttcg cttcttggtc gtcatagttc ctcgcgtgtc gatggtcatc

```
    60721 gacttcgcca aacctgccgc ctcctgttcg agacgacgcg aacgctccac ggcggccgat
    60781 ggcgcgggca gggcaggggg agccagttgc acgctgtcgc gctcgatctt ggccgtagct
    60841 tgctggacca tcgagccgac ggactggaag gtttcgcggg gcgcacgcat gacggtgcgg
    60901 cttgcgatgg tttcggcatc ctcggcggaa aaccccgcgt cgatcagttc ttgcctgtat
    60961 gccttccggt caaacgtccg attcattcac cctccttgcg ggattgcccc gactcacgcc
    61021 ggggcaatgt gcccttattc ctgatttgac ccgcctggtg ccttggtgtc cagataatcc
    61081 accttatcgg caatgaagtc ggtcccgtag accgtctggc cgtccttctc gtacttggta
    61141 ttccgaatct tgccctgcac gaataccagc gaccccttgc ccaaatactt gccgtgggcc
    61201 tcggcctgag agccaaaaca cttgatgcgg aagaagtcgg tgcgctcctg cttgtcgccg
    61261 gcatcgttgc gccactcttc attaaccgct atatcgaaaa ttgcttgcgg cttgttagaa
    61321 ttgccatgac gtacctcggt gtcacgggta agattaccga taaactggaa ctgattatgg
    61381 ctcatatcga aagtctcctt gagaaaggag actctagttt agctaaacat tggttccgct
    61441 gtcaagaact ttagcggcta aaattttgcg ggccgcgacc aaaggtgcga ggggcggctt
    61501 ccgctgtgta caaccagata tttttcacca acatccttcg tctgctcgat gagcg
//
```

**pMAS046**

```
LOCUS       pMAS046                61560 bp ds-DNA     circular     24-FEB-
2026
DEFINITION  .
FEATURES             Location/Qualifiers
     gene            587..1546
                     /label="trbB gene"
                     /ApEinfo_revcolor="#9eafd2"
                     /ApEinfo_fwdcolor="#9eafd2"
     gene            1559..1996
                     /label="trbC gene"
                     /ApEinfo_revcolor="#b1ff67"
                     /ApEinfo_fwdcolor="#b1ff67"
     gene            1999..2310
                     /label="trbD gene"
                     /ApEinfo_revcolor="#9eafd2"
                     /ApEinfo_fwdcolor="#9eafd2"
     gene            2307..4865
                     /label="trbE gene"
                     /ApEinfo_revcolor="#84b0dc"
                     /ApEinfo_fwdcolor="#84b0dc"
     gene            4862..5620
                     /label="trbF gene"
                     /ApEinfo_revcolor="#faac61"
                     /ApEinfo_fwdcolor="#faac61"
     gene            5632..6525
                     /label="trbG gene"
                     /ApEinfo_revcolor="#c6c9d1"
                     /ApEinfo_fwdcolor="#c6c9d1"
     gene            6529..7011
                     /label="trbH gene"
                     /ApEinfo_revcolor="#ffef86"
                     /ApEinfo_fwdcolor="#ffef86"
     gene            7016..8407
                     /label="trbI gene"
                     /ApEinfo_revcolor="#b7e6d7"
                     /ApEinfo_fwdcolor="#b7e6d7"
     gene            8424..9200
                     /label="trbJ gene"
                     /ApEinfo_revcolor="#75c6a9"
                     /ApEinfo_fwdcolor="#75c6a9"
     gene            9212..9421
                     /label="trbK gene"
                     /ApEinfo_revcolor="#85dae9"
                     /ApEinfo_fwdcolor="#85dae9"
     gene            9428..11014
                     /label="trbL gene"
                     /ApEinfo_revcolor="#ff9ccd"
                     /ApEinfo_fwdcolor="#ff9ccd"
     gene            11038..11637
                     /label="trbM gene"
                     /ApEinfo_revcolor="#f58a5e"
                     /ApEinfo_fwdcolor="#f58a5e"
     gene            11653..12357
                     /label="trbN gene"
```

```
                     /ApEinfo_revcolor="#ff9ccd"
                     /ApEinfo_fwdcolor="#ff9ccd"
     gene            12388..12651
                     /label="trbO gene"
                     /ApEinfo_revcolor="#c7b0e3"
                     /ApEinfo_fwdcolor="#c7b0e3"
     gene            13873..14568
                     /label="fiwA gene"
                     /ApEinfo_revcolor="#d6b295"
                     /ApEinfo_fwdcolor="#d6b295"
     gene            14604..15266
                     /label="upf32.8 gene"
                     /ApEinfo_revcolor="#84b0dc"
                     /ApEinfo_fwdcolor="#84b0dc"
     operon          complement(15268..16770)
                     /label="operon"
                     /ApEinfo_revcolor="#d6b295"
                     /ApEinfo_fwdcolor="#d6b295"
     gene            complement(15290..15949)
                     /label="parA gene"
                     /ApEinfo_revcolor="#f8d3a9"
                     /ApEinfo_fwdcolor="#f8d3a9"
     gene            complement(15910..16443)
                     /label="parB gene"
                     /ApEinfo_revcolor="#9eafd2"
                     /ApEinfo_fwdcolor="#9eafd2"
     gene            complement(16440..16700)
                     /label="parC gene"
                     /ApEinfo_revcolor="#f8d3a9"
                     /ApEinfo_fwdcolor="#f8d3a9"
     gene            16884..17135
                     /label="parD gene"
                     /ApEinfo_revcolor="#84b0dc"
                     /ApEinfo_fwdcolor="#84b0dc"
     gene            17132..17443
                     /label="parE gene"
                     /ApEinfo_revcolor="#c7b0e3"
                     /ApEinfo_fwdcolor="#c7b0e3"
     gene            17521..17970
                     /label="upf35.8 gene"
                     /ApEinfo_revcolor="#d59687"
                     /ApEinfo_fwdcolor="#d59687"
     gene            19244..20041
                     /label="istB gene"
                     /ApEinfo_revcolor="#d59687"
                     /ApEinfo_fwdcolor="#d59687"
     misc_feature    20329..20360
                     /label="AphA-5'-IGR gRNA target"
                     /ApEinfo_revcolor="#b7e6d7"
                     /ApEinfo_fwdcolor="#b7e6d7"
     gene            21341..21369
                     /label="aphA gene"
                     /ApEinfo_revcolor="#f8d3a9"
                     /ApEinfo_fwdcolor="#f8d3a9"
     operon          complement(21341..30485)
                     /label="operon"
                     /ApEinfo_revcolor="#85dae9"
```

```
                     /ApEinfo_fwdcolor="#85dae9"
     gene            complement(21376..21666)
                     /label="traA gene"
                     /ApEinfo_revcolor="#ffef86"
                     /ApEinfo_fwdcolor="#ffef86"
     gene            complement(21674..22114)
                     /label="traB gene"
                     /ApEinfo_revcolor="#85dae9"
                     /ApEinfo_fwdcolor="#85dae9"
     gene            complement(22130..24370)
                     /label="traC2 gene"
                     /ApEinfo_revcolor="#b1ff67"
                     /ApEinfo_fwdcolor="#b1ff67"
     gene            complement(22130..25315)
                     /label="traC1 gene"
                     /ApEinfo_revcolor="#b4abac"
                     /ApEinfo_fwdcolor="#b4abac"
     gene            complement(25322..25585)
                     /label="traD gene"
                     /ApEinfo_revcolor="#f8d3a9"
                     /ApEinfo_fwdcolor="#f8d3a9"
     gene            complement(25591..27804)
                     /label="traE gene"
                     /ApEinfo_revcolor="#f8d3a9"
                     /ApEinfo_fwdcolor="#f8d3a9"
     gene            complement(27819..28352)
                     /label="traF gene"
                     /ApEinfo_revcolor="#faac61"
                     /ApEinfo_fwdcolor="#faac61"
     gene            complement(28349..30256)
                     /label="traG gene"
                     /ApEinfo_revcolor="#faac61"
                     /ApEinfo_fwdcolor="#faac61"
     gene            complement(30253..32451)
                     /label="traI gene"
                     /ApEinfo_revcolor="#b4abac"
                     /ApEinfo_fwdcolor="#b4abac"
     operon          complement(30253..33086)
                     /label="operon"
                     /ApEinfo_revcolor="#b1ff67"
                     /ApEinfo_fwdcolor="#b1ff67"
     gene            complement(30552..30911)
                     /label="traH gene"
                     /ApEinfo_revcolor="#b4abac"
                     /ApEinfo_fwdcolor="#b4abac"
     gene            complement(32448..32489)
                     /label="traX gene"
                     /ApEinfo_revcolor="#b7e6d7"
                     /ApEinfo_fwdcolor="#b7e6d7"
     gene            complement(32486..32857)
                     /label="traJ gene"
                     /ApEinfo_revcolor="#faac61"
                     /ApEinfo_fwdcolor="#faac61"
     oriT            32890..32999
                     /label="incP origin of transfer"
                     /ApEinfo_revcolor="#84b0dc"
                     /ApEinfo_fwdcolor="#84b0dc"
```

```
     gene            33225..33629
                     /label="traK gene"
                     /ApEinfo_revcolor="#c6c9d1"
                     /ApEinfo_fwdcolor="#c6c9d1"
     gene            33629..34354
                     /label="traL gene"
                     /ApEinfo_revcolor="#84b0dc"
                     /ApEinfo_fwdcolor="#84b0dc"
     gene            34351..34788
                     /label="traM gene"
                     /ApEinfo_revcolor="#b7e6d7"
                     /ApEinfo_fwdcolor="#b7e6d7"
     gene            complement(36621..36657)
                     /label="krfA gene"
                     /ApEinfo_revcolor="#f8d3a9"
                     /ApEinfo_fwdcolor="#f8d3a9"
     gene            complement(37833..38360)
                     /label="korG gene"
                     /ApEinfo_revcolor="#f58a5e"
                     /ApEinfo_fwdcolor="#f58a5e"
     gene            complement(38944..40020)
                     /label="korB gene"
                     /ApEinfo_revcolor="#b7e6d7"
                     /ApEinfo_fwdcolor="#b7e6d7"
     gene            complement(40017..41111)
                     /label="incC gene"
                     /ApEinfo_revcolor="#85dae9"
                     /ApEinfo_fwdcolor="#85dae9"
     gene            complement(40793..41098)
                     /label="korA gene"
                     /ApEinfo_revcolor="#b4abac"
                     /ApEinfo_fwdcolor="#b4abac"
     gene            complement(41272..42225)
                     /label="klaC gene"
                     /ApEinfo_revcolor="#ffef86"
                     /ApEinfo_fwdcolor="#ffef86"
     operon          complement(41272..44222)
                     /label="operon"
                     /ApEinfo_revcolor="#84b0dc"
                     /ApEinfo_fwdcolor="#84b0dc"
     gene            complement(42222..43358)
                     /label="klaB gene"
                     /ApEinfo_revcolor="#faac61"
                     /ApEinfo_fwdcolor="#faac61"
     gene            complement(43376..44149)
                     /label="klaA gene"
                     /ApEinfo_revcolor="#85dae9"
                     /ApEinfo_fwdcolor="#85dae9"
     gene            complement(44272..44586)
                     /label="kleF gene"
                     /ApEinfo_revcolor="#b7e6d7"
                     /ApEinfo_fwdcolor="#b7e6d7"
     operon          complement(44272..45592)
                     /label="operon"
                     /ApEinfo_revcolor="#84b0dc"
                     /ApEinfo_fwdcolor="#84b0dc"
     gene            complement(44614..44937)
```

```
                     /label="kleE gene"
                     /ApEinfo_revcolor="#d6b295"
                     /ApEinfo_fwdcolor="#d6b295"
     gene            complement(45049..45267)
                     /label="kleD gene"
                     /ApEinfo_revcolor="#faac61"
                     /ApEinfo_fwdcolor="#faac61"
     gene            complement(45283..45513)
                     /label="kleC gene"
                     /ApEinfo_revcolor="#9eafd2"
                     /ApEinfo_fwdcolor="#9eafd2"
     gene            complement(45666..45881)
                     /label="kleB gene"
                     /ApEinfo_revcolor="#c6c9d1"
                     /ApEinfo_fwdcolor="#c6c9d1"
     operon          complement(45666..46247)
                     /label="operon"
                     /ApEinfo_revcolor="#f58a5e"
                     /ApEinfo_fwdcolor="#f58a5e"
     gene            complement(45930..46163)
                     /label="kleA gene"
                     /ApEinfo_revcolor="#c6c9d1"
                     /ApEinfo_fwdcolor="#c6c9d1"
     gene            complement(46283..46308)
                     /label="korC gene"
                     /ApEinfo_revcolor="#d59687"
                     /ApEinfo_fwdcolor="#d59687"
     gene            complement(46389..46646)
                     /label="korC gene"
                     /ApEinfo_revcolor="#b4abac"
                     /ApEinfo_fwdcolor="#b4abac"
     gene            complement(46669..47343)
                     /label="klcB gene"
                     /ApEinfo_revcolor="#b4abac"
                     /ApEinfo_fwdcolor="#b4abac"
     gene            complement(47495..48135)
                     /label="Disrupted bla gene 3'"
                     /ApEinfo_revcolor="#75c6a9"
                     /ApEinfo_fwdcolor="#75c6a9"
     misc_feature    48057..48088
                     /label="bla CDS g5"
                     /ApEinfo_revcolor="#c6c9d1"
                     /ApEinfo_fwdcolor="#c6c9d1"
     misc_feature    48131..48135
                     /label="Target site duplication"
                     /ApEinfo_revcolor="#ff9ccd"
                     /ApEinfo_fwdcolor="#ff9ccd"
     misc_feature    48136..48143
                     /label="END"
                     /ApEinfo_revcolor="#84b0dc"
                     /ApEinfo_fwdcolor="#84b0dc"
                     /note="ApEinfo_revcolor: #84b0dc"
     misc_feature    48136..48260
                     /label="VchINT R-end"
                     /ApEinfo_revcolor="#ffef86"
                     /ApEinfo_fwdcolor="#ffef86"
                     /note="ApEinfo_revcolor: #ffef86"
```

```
     misc_feature    48144..48163
                     /label="R1 tnsB binding site (VCHE45 Tn6677)"
                     /ApEinfo_revcolor="#c7b0e3"
                     /ApEinfo_fwdcolor="#c7b0e3"
     misc_feature    48144..48246
                     /label="RE (partial) (VCHE45 Tn6677)"
                     /ApEinfo_revcolor="#c6c9d1"
                     /ApEinfo_fwdcolor="#c6c9d1"
     misc_feature    48164..48183
                     /label="R2 tnsB binding site (VCHE45 Tn6677)"
                     /ApEinfo_revcolor="#c7b0e3"
                     /ApEinfo_fwdcolor="#c7b0e3"
     misc_feature    48187..48206
                     /label="R3 tnsB binding site (VCHE45 Tn6677)"
                     /ApEinfo_revcolor="#c7b0e3"
                     /ApEinfo_fwdcolor="#c7b0e3"
     terminator      48261..48309
                     /label="tVoigtS5, L3S3P22 47C>G"
                     /ApEinfo_revcolor="#c6c9d1"
                     /ApEinfo_fwdcolor="#c6c9d1"
     misc_feature    48315..48322
                     /label="SbfI"
                     /ApEinfo_revcolor="#85dae9"
                     /ApEinfo_fwdcolor="#85dae9"
     misc_feature    48327..48358
                     /label="CymR operator"
                     /ApEinfo_revcolor="#ffef86"
                     /ApEinfo_fwdcolor="#ffef86"
     promoter        48327..48416
                     /label="P.CymRC (Marionette)"
                     /ApEinfo_revcolor="#ffef86"
                     /ApEinfo_fwdcolor="#ffef86"
     misc_feature    48350..48355
                     /label=-35
                     /ApEinfo_revcolor="#ffef86"
                     /ApEinfo_fwdcolor="#ffef86"
     misc_feature    48373..48378
                     /label=-10
                     /ApEinfo_revcolor="#ffef86"
                     /ApEinfo_fwdcolor="#ffef86"
     misc_feature    48385..48385
                     /label="TSS"
                     /ApEinfo_revcolor="#ffef86"
                     /ApEinfo_fwdcolor="#ffef86"
     misc_feature    48385..48416
                     /label="CymR operator"
                     /ApEinfo_revcolor="#ffef86"
                     /ApEinfo_fwdcolor="#ffef86"
     misc_feature    48421..48476
                     /label="U64 RNA Guide"
                     /ApEinfo_revcolor="#faac61"
                     /ApEinfo_fwdcolor="#faac61"
     misc_feature    48421..48780
                     /label="U64 Ribozyme transcript"
                     /ApEinfo_revcolor="#d59687"
                     /ApEinfo_fwdcolor="#d59687"
     misc_feature    48471..48476
```

```
                     /label="IGS"
                     /ApEinfo_revcolor="#c7b0e3"
                     /ApEinfo_fwdcolor="#c7b0e3"
     misc_feature    48477..48780
                     /label="Group I Intron Ribozyme (from p-OiRS3GG)"
                     /ApEinfo_revcolor="#b4abac"
                     /ApEinfo_fwdcolor="#b4abac"
     misc_feature    48781..48863
                     /label="Group I Intron Ribozyme (from p-OiRS3GG)"
                     /ApEinfo_revcolor="#b4abac"
                     /ApEinfo_fwdcolor="#b4abac"
     misc_feature    48781..49375
                     /label="U64 Ribozyme transcript"
                     /ApEinfo_revcolor="#d59687"
                     /ApEinfo_fwdcolor="#d59687"
     misc_feature    48864..48884
                     /label="Probe Binding Site"
                     /ApEinfo_revcolor="#b4abac"
                     /ApEinfo_fwdcolor="#b4abac"
     misc_feature    48864..48885
                     /label="Barcode (sfGFP_2)"
                     /ApEinfo_revcolor="#84b0dc"
                     /ApEinfo_fwdcolor="#84b0dc"
     misc_feature    48885..48889
                     /label="Internal Barcode"
                     /ApEinfo_revcolor="#b1ff67"
                     /ApEinfo_fwdcolor="#b1ff67"
     misc_feature    48890..49375
                     /label="Barcode (sfGFP_2)"
                     /ApEinfo_revcolor="#84b0dc"
                     /ApEinfo_fwdcolor="#84b0dc"
     misc_feature    49376..49386
                     /label="Barcode (sfGFP_2)"
                     /ApEinfo_revcolor="#84b0dc"
                     /ApEinfo_fwdcolor="#84b0dc"
     misc_feature    49376..49386
                     /label="U64 Ribozyme transcript"
                     /ApEinfo_revcolor="#d59687"
                     /ApEinfo_fwdcolor="#d59687"
     terminator      49391..49443
                     /label="tVoigtS4 L3S3P21 51C>G"
                     /ApEinfo_revcolor="#f58a5e"
                     /ApEinfo_fwdcolor="#f58a5e"
     misc_feature    complement(49450..49594)
                     /label="VchINT L-end"
                     /ApEinfo_revcolor="#ffef86"
                     /ApEinfo_fwdcolor="#ffef86"
                     /note="ApEinfo_revcolor: #ffef86"
     misc_feature    complement(49450..49594)
                     /label="LE (partial) (VCHE45 Tn6677)"
                     /ApEinfo_revcolor="#b4abac"
                     /ApEinfo_fwdcolor="#b4abac"
     misc_feature    complement(49487..49506)
                     /label="L3 tnsB binding site (VCHE45 Tn6677)"
                     /ApEinfo_revcolor="#f58a5e"
                     /ApEinfo_fwdcolor="#f58a5e"
     misc_feature    complement(49513..49532)
```

```
                     /label="L2 tnsB binding site (VCHE45 Tn6677)"
                     /ApEinfo_revcolor="#f58a5e"
                     /ApEinfo_fwdcolor="#f58a5e"
     misc_feature    complement(49567..49586)
                     /label="L1 tnsB binding site (VCHE45 Tn6677)"
                     /ApEinfo_revcolor="#f58a5e"
                     /ApEinfo_fwdcolor="#f58a5e"
     misc_feature    complement(49587..49594)
                     /label="END"
                     /ApEinfo_revcolor="#84b0dc"
                     /ApEinfo_fwdcolor="#84b0dc"
                     /note="ApEinfo_revcolor: #84b0dc"
     misc_feature    49595..49599
                     /label="Target site dupication"
                     /ApEinfo_revcolor="#ff9ccd"
                     /ApEinfo_fwdcolor="#ff9ccd"
     gene            complement(49600..49819)
                     /label="Disrupted bla 5'"
                     /ApEinfo_revcolor="#c7b0e3"
                     /ApEinfo_fwdcolor="#c7b0e3"
     misc_feature    complement(49714..49745)
                     /label="bla CDS g4"
                     /ApEinfo_revcolor="#75c6a9"
                     /ApEinfo_fwdcolor="#75c6a9"
     promoter        complement(49820..49924)
                     /label="AmpR promoter"
                     /ApEinfo_revcolor="#f58a5e"
                     /ApEinfo_fwdcolor="#f58a5e"
     gene            complement(50002..50559)
                     /label="tnpR gene"
                     /ApEinfo_revcolor="#b4abac"
                     /ApEinfo_fwdcolor="#b4abac"
     gene            complement(53762..54472)
                     /label="klcB gene"
                     /ApEinfo_revcolor="#c6c9d1"
                     /ApEinfo_fwdcolor="#c6c9d1"
     gene            complement(54518..54958)
                     /label="klcA gene"
                     /ApEinfo_revcolor="#ff9ccd"
                     /ApEinfo_fwdcolor="#ff9ccd"
     rep_origin      55430..56141
                     /label="oriV"
                     /ApEinfo_revcolor="#b7e6d7"
                     /ApEinfo_fwdcolor="#b7e6d7"
                     /note="incP origin of replication"
     gene            complement(56615..57265)
                     /label="tetR gene"
                     /ApEinfo_revcolor="#ff9ccd"
                     /ApEinfo_fwdcolor="#ff9ccd"
     gene            57371..58570
                     /label="tetA gene"
                     /ApEinfo_revcolor="#75c6a9"
                     /ApEinfo_fwdcolor="#75c6a9"
     misc_feature    58748..58779
                     /label="TetA-3'-IGR gRNA target"
                     /ApEinfo_revcolor="#b4abac"
                     /ApEinfo_fwdcolor="#b4abac"
```

```
     gene            complement(59843..60991)
                     /label="trfA gene"
                     /ApEinfo_revcolor="#9eafd2"
                     /ApEinfo_fwdcolor="#9eafd2"
     gene            complement(61040..61390)
                     /label="ssb gene"
                     /ApEinfo_revcolor="#ffef86"
                     /ApEinfo_fwdcolor="#ffef86"
     gene            61511..316
                     /label="trbA gene"
                     /ApEinfo_revcolor="#c6c9d1"
                     /ApEinfo_fwdcolor="#c6c9d1"
ORIGIN
        1 gggcatgacg aaacatgagc tgtcggagag ggcaggggtt tcaatttcgt ttttatcaga
       61 cttaaccaac ggtaaggcca acccctcgtt gaaggtgatg gaggccattg ccgacgccct
      121 ggaaactccc ctacctcttc tcctggagtc caccgacctt gaccgcgagg cactcgcgga
      181 gattgcgggt catcctttca agagcagcgt gccgcccgga tacgaacgca tcagtgtggt
      241 tttgccgtca cataaggcgt ttatcgtaaa gaaatggggc gacgacaccc gaaaaaagct
      301 gcgtggaagg ctctgacgcc aagggttagg gcttgcactt ccttctttag ccgctaaaac
      361 ggccccttct ctgcgggccg tcggctcgcg catcatatcg acatcctcaa cggaagccgt
      421 gccgcgaatg gcatcgggcg ggtgcgcttt gacagttgtt ttctatcaga acccctacgt
      481 cgtgcggttc gattagctgt ttgtcttgca ggctaaacac tttcggtata tcgtttgcct
      541 gtgcgataat gttgctaatg atttgttgcg taggggttac tgaaaagtga gcgggaaaga
      601 agagtttcag accatcaagg agcgggccaa gcgcaagctg gaacgcgaca tgggtgcgga
      661 cctgttggcc gcgctcaacg acccgaaaac cgttgaagtc atgctcaacg cggacggcaa
      721 ggtgtggcac gaacgccttg gcgagccgat gcggtacatc tgcgacatgc ggcccagcca
      781 gtcgcaggcg attatagaaa cggtggccgg attccacggc aaagaggtca cgcggcattc
      841 gcccatcctg gaaggcgagt tcccccttgga tggcagccgc tttgccggcc aattgccgcc
      901 ggtcgtggcc gcgccaacct ttgcgatccg caagcgcgcg gtcgccatct tcacgctgga
      961 acagtacgtc gaggcgggca tcatgacccg cgagcaatac gaggtcatta aaagcgccgt
     1021 cgcggcgcat cgaaacatcc tcgtcattgg cggtactggc tcgggcaaga ccacgctcgt
     1081 caacgcgatc atcaatgaaa tggtcgcctt caacccgtct gagcgcgtcg tcatcatcga
     1141 ggacaccggc gaaatccagt gcgccgcaga gaacgccgtc caataccaca ccagcatcga
     1201 cgtctcgatg acgctgctgc tcaagacaac gctgcgtatg cgccccgacc gcatcctggt
     1261 cggtgaggta cgtggccccg aagcccttga tctgttgatg gcctggaaca ccgggcatga
     1321 aggaggtgcc gccaccctgc acgcaaacaa ccccaaagcg ggcctgagcc ggctcgccat
     1381 gcttatcagc atgcacccgg attcaccgaa acccattgag ccgctgattg gcgaggcggt
     1441 tcatgtggtc gtccatatcg ccaggacccc tagcggccgt cgagtgcaag aaattctcga
     1501 agttcttggt tacgagaacg gccagtacat caccaaaacc ctgtaaggag tatttccaat
     1561 gacaacggct gttccgttcc gtctgaccat gaatcgcggc attttgttct accttgccgt
     1621 gttcttcgtt ctcgctctcg cgttatccgc gcatccggcg atggcctcgg aaggcaccgg
     1681 cggcagcttg ccatatgaga gctggctgac gaacctgcgc aactccgtaa ccggcccggt
     1741 ggccttcgcg ctgtccatca tcggcatcgt cgtcgccggc ggcgtgctga tcttcggcgg
     1801 cgaactcaac gccttcttcc gaaccctgat cttcctggtt ctggtgatgg cgctgctggt
     1861 cggcgcgcag aacgtgatga gcaccttctt cggtcgtggt gccgaaatcg cggccctcgg
     1921 caacggggcg ctgcaccagg tgcaagtcgc ggcggcggat gccgtgcgtg cggtagcggc
     1981 tggacggctc gcctaatcat ggctctgcgc acgatcccca tccgtcgcgc aggcaaccga
     2041 gaaaacctgt tcatgggtgg tgatcgtgaa ctggtgatgt tctcgggcct gatggcgttt
     2101 gcgctgattt tcagcgccca agagctgcgg gccaccgtgg tcggtctgat cctgtggttc
     2161 ggggcgctct atgcgttccg aatcatggcg aaggccgatc cgaagatgcg gttcgtgtac
     2221 ctgcgtcacc gccggtacaa gccgtattac ccggcccgct cgaccccgtt ccgcgagaac
     2281 accaatagcc aagggaagca ataccgatga tccaagcaat tgcgattgca atcgcgggcc
     2341 tcggcgcgct tctgttgttc atcctctttg cccgcatccg cgcggtcgat gccgaactga
     2401 aactgaaaaa gcatcgttcc aaggacgccg gcctggccga tctgctcaac tacgccgctg
     2461 tcgtcgatga cggcgtaatc gtgggcaaga acggcagctt tatggctgcc tggctgtaca
     2521 agggcgatga caacgcaagc agcaccgacc agcagcgcga agtagtgtcc gcccgcatca
     2581 accaggccct cgcgggcctg ggaagtgggt ggatgatcca tgtggacgcc gtgcggcgtc
```

2641 ctgctccgaa ctacgcggag cggggcctgt cggcgttccc tgaccgtctg acggcagcga
2701 ttgaagaaga gcgccggcgg catttcgaga gcctgggaac gatgtacgag ggctatttcg
2761 tcctcacctt gacctggttc ccgccgctgc tcgcccagcg caagttcgtc gagctgatgt
2821 ttgacgacga cgcgaccgca ccggatcgca aggcgcgcac gcggggcctc atcgaccaat
2881 tcaagcgtga cgtgcgcagc atcgagtcgc gcctgtcgtc ggccgtgtcg ctcactcgct
2941 tgaaggggca caagatcgtc aacgaggacg gcacgaccgt cacgcatgac gacttcctgc
3001 gctggctgca attctgcgtg acgggcctgc accatccggt gcagctcccc agcaacccga
3061 tgtacctgga cgccctggtc ggcggacagg aaatgtgggg cggggtagtg cccaaggtcg
3121 gccgcaagtt cgtccaggtg gtcgctctcg aaggcttccc cttggagtcc tatcccggca
3181 tcctgacggc gctcggcgag ctgccctgcg agtatcggtg gtcgagccgg ttcatcttca
3241 tggaccagca cgaagccgtg aagcacctcg acaagttccg caagaagtgg cggcagaaga
3301 ttcgcggctt cttcgaccag gtgttcaaca cgaacaccgg cccggtcgat caggacgcgc
3361 tttcgatggt ggccgatgct gaggcggcca ttgccgaagt caacagcggc atcgtggccg
3421 tgggctacta caccagcgtc gtcgtgctga tggatgagga ccgcacgcgc ctggaagctg
3481 cggcccgcga tgttgaaaag gccgtcaacc ggttgggctt tgccgcgcgc atcgagtcca
3541 tcaacaccct ggacgccttc cttggtagtt tgccgggcca cggcgtggaa aacgtccgcc
3601 ggccgctcat caacacgatg aacctggccg acctgctgcc gaccagcacc atctggaccg
3661 gcaacgcgaa cgcgccatgc ccgatgtacc cgccgctgtc gccggcgctc atgcactgcg
3721 tcacgcaagg atcaacgccg ttccggctga acctgcacgt gcgcgacctc ggccacacct
3781 ttatgttcgg gccgaccggc gcaggtaaat cgacgcacct ggcgatcctc gccgcgcagc
3841 tccgtcgcta tgccggcatg tcgatcttcg cctttgacaa gggcatgtcg atgtacccgc
3901 tggccgccgg catccgtgcg gccacgaagg gcaccagcgg cctgcacttc accgtggcgg
3961 ccgacgacga acgcctggcg ttctgcccgt tgcagttcct gagcaccaag ggcgaccgtg
4021 cttgggcgat ggagtggatc gacaccatcc tggcgttgaa cggcgtcgaa acgaccccgg
4081 cccagcgcaa cgaaatcggc aacgcgatca tgagcatgca cgccagcggc gcgcgcacgc
4141 tctccgagtt cagcgtgacg attcaggatg aggcgatccg cgaggcgatc cgccagtaca
4201 ccgtcgatgg cgcaatgggc catctgctcg acgccgaaga ggacggcttg gcgctgtccg
4261 actttacagt gttcgagatc gaagagctga tgaacctcgg cgagaaattc gccctgcctg
4321 tgttgctcta cctgttccgc cgtatcgagc gcgccctgac gggccagccg gccgtcatca
4381 tcctggacga agcctggttg atgctcggcc acccggcatt ccgcgcgaag atcagggaat
4441 ggctcaaggt gctgcgtaag gccaactgcc ttgtgctgat ggcaacgcag agcctgtccg
4501 acgccgccaa cagcggcatc ctggacgtga tcgtggaatc gaccgcgacc aagattttcc
4561 tgccgaatat ttacgccagg gatgaggaca cggcggccct gtaccgccgc atgggcctga
4621 acgctcgcca gatcgagatt ctggcccagg ccgttcccaa gcgtcagtac tactacgtgt
4681 cggaaaacgg ccgccgtctc tacgacctgg cacttggccc gctcgcgctc gcgttcgtcg
4741 gcgcatccga caaggaatcc gtcgccatca tcaagaacct ggaagccaag ttcggcgacc
4801 agtgggtgga tgaatggctg cgtggccggg gcctcgccct tgatgaatac ctggaggcag
4861 catgagtttt gcagacacga tcaagggctt gatcttcaag aagaagcccg caacggccgc
4921 agcagcggcg acgccggccg cgaccggccc gcaaaccgac aacccgtacc tgacggcgcg
4981 gcgcacctgg aacgaccacg ttggttccgt tgtgtcgcaa aagcagacct ggcaggttgt
5041 cggcatcctt tcgctgatga tcgtcctcgc ggcggtcggc ggcatcatcc acatcggcag
5101 ccagtcgaag ttcgtgccct atgtctacga ggtagacaag ctcgggcaga cggccgccgt
5161 ggggccgatg accagggcgt cgaaagccga tccgcgtgtc attcacgcct cggtggctga
5221 gttcgtcggc gatgctcgcc tggtgacgcc ggacgtagct ttgcagcgca aggccgtcta
5281 ccgcctctat gccaagctcg ggccgaatga cccggccacc gccaagatga acgaatggct
5341 caacggcacc gccgacgcca gcccgttcgc tcgcgcggcc gtcgaaacgg tcagcaccga
5401 aatcacttcc gtaatcccgc agacgcccga cacctggcag gtcgattggg tcgagacgac
5461 gcgcgacagg caaggcgtgg tgaaaggcca gcccgtgcgc atgcgggcct tggtgacggt
5521 ctacgtcgtc gagccgacgg cggacaccaa ggaagaacaa ctgcgaaaca acccggccgg
5581 gatctacgtc cgggacttct cctggtcgag acttctgtga ggcactgaat tatgaaaaag
5641 gaactgtttg ctttggtcct ggccgcgtcc gttagcgtgc ctgcatttgc cgccgatccc
5701 ggcgcggacc tgactgacct ctatttttcc ggcaagaacc cggagctgac cgcgcaagag
5761 cgggcggcca tcgccatcgc caagaagtgg gaggcgggta ccgccggcat gcggccggtg
5821 gccggccccg gtggttcggt gcgcttcctg ttcggcgcgc agcagccgag catcgtatgc
5881 gccgtgctgc aagtgtgcga cgtggccctg caacccggcg agcaagtcaa ctcgatcaac
5941 ctgggcgaca ccgcccgttg gacggtcgag ccggccatta ccggcagcgg cgcgaacgaa
6001 acccagcacc tcatcatcaa gccgatggat gtgggcctgg aaaccagcct ggtcgtgacc

```
6061 acggaccgcc gcagctacca catgcgcctg cgctcgcatc gcacgcagta catgccgcag
6121 gtgtcgttca cctacccgga agatgccctt gcgaagtggg acgccatcaa gaaccgcgaa
6181 cagcgggatc gcgtcgagaa aaccattccg cagaccggcg agtacctggg caacctgagc
6241 ttcaactact ccgtcagcgg gtccacgtcg tggaagccgg tgcgcgtcta caacgacggc
6301 aagaaaacca tcatccagat gccgcactcg atggaacaga ccgaagcgcc gacgctcctg
6361 gtcgttcgca gggagggcgg cctgttctcc gacgatgaaa cggtgatggt caactaccgg
6421 gtccagggcg accgctacat cgtcgatacg attttcgaca aggccatcct catcgcgggc
6481 gtgggcagca gccaggaccg cgtgaccatt tcaaggggga actaaaccat gcgtaagatt
6541 ctgaccgtca tcgcactcgc ggccacgttg gccggctgcg cgacctccaa gtacggcagc
6601 ttcgtccagg acgcgccggc cgcctacaac cagaccattg cgaccgacgc ggtgaagcag
6661 ctcgtcaagc tctacccgcc ggcgcaaacc aagctggaat tgcagcaggc tacgcccgat
6721 ccgttcggca ttgccctggt cactgacctt cgcgcccagg gctatgctgt catggagtac
6781 aagcccgacg gcaacgcggc cgcagctccg gctgctgcgt cctcggccgc tgcgaagccg
6841 gcaacgccgc aagcccaggg cggctatccg ctgcgctacg tgctggacca attcagcgac
6901 agcaacctgt atcgcctgac cgtcatggtc ggctctcaat cgctcacgcg cgcctacctc
6961 gcccaaaaca acacgatggt cccggccggc gcatgggttc ggaaggagta agccaatgag
7021 cgaagatcaa atggcaccgg acgcatcgcc agatgcggtc aagccgaaaa gcggggttcg
7081 ccgcgtcaac aacatgccga tgtacctcat cggcggtgtg ctcggcatct tcctgctggt
7141 gatggccctg gttgctgcgg atcgcgctgc gcagcagaac cagccgggag ctgcgaaggc
7201 tgagaaggcc ggcagcacca gcatgtttgc cgacgaaatt gccggcaaac agcaggacgg
7261 catcatcaag gccaagccgc tggagattcc gccggaacaa accgcccagc aaccgacgac
7321 ggagctgacg ccagccccgg cgcagggaac gactatcacg gtcgcacggc ccgagaacct
7381 ggaccagccc ccgacgccgc cgcagggtgc gcgcaacgag gacctggacc gcatccgcat
7441 ggcgaagttg cagatgctgg aagaggcgat caaggccaag acgacggtgc gcatcgacgc
7501 gccgcgcagc cagggcagcg ccggcggcgg tgctccgcag ggccgcgagg aaacccttgc
7561 gcgcatccag gagctgcgtc ggcaggctga gaacgcccgc gccaccgatc cgaccgccgc
7621 ctatcaggcc gcgcttgcgc aggctcgcac gatgggcggc gcggcagggg gtggcggtat
7681 gggcggctcg ggtgcgccga ccctcgtgca gacctcgaac cgcagtggtg gcggcgctgg
7741 ctatgggtcg ttcgacaacc gcagcgaggg cgaccgttgg cggctcgact cccagccgga
7801 agcacctgca acgccctatg tgctgcgcgc tggcttcgtc gttccggcta cgcttatctc
7861 gggcatcaac tccgatctgc caggccaaat catggcccag gtatcgcagt cggtgtacga
7921 cacggcgacc ggcaagcaca tgctcatccc ccaaggctcg cgcctggtgg gcagctactc
7981 gaacgatgtg gcctacgggc agaagcgcgt tctggtggca tggcagcgca tcatcttccc
8041 cgacggcaag gcaatggaca ttggggccat gccgggcggc gatagcgctg ggtatgcagg
8101 cttcaacgac aaggtcaaca accactactt ccgcaccttc gcatcggcat tcctcatgtc
8161 gggcgtcgtt gcgggcatca gcttgagtca ggaccgtggc aacagcaaca gcggttacgg
8221 acgacaagac gcgggttccg cgatgagtga agcgttgggt caacagctcg gccaagtaac
8281 ggcgcagatg atcgccaaaa acttgaatat cgcgccgacg ctggaaatcc gtccgggcta
8341 tcgcttcaac gtcattgtca cgaaagacat gacgttttct aagccctacc aggcgtttga
8401 ctattaactc caaggagtaa cttatgaaga agctcgctaa gaatgtttta gccgctaaag
8461 tagctctggt gctggccctc tcggtcggca ccttggcggt cacgcctgcg caagcgggca
8521 ttccggtcat cgacggcacc aacctgtcac aaaccactgt caccgcgatt cagcaggttg
8581 cgcaggtcca gaagcaaatc gaggaatacc ggacgcagtt gcagcagtac gaaaacatgc
8641 tgcaaaacac ggtggccccg gccgcctacg tgtgggacca ggcgcagtcc accatcaacg
8701 gcctgatgag cgccgttgat accctgaact actacaagaa ccaggcgggc agcatcgacg
8761 cttacctggg caagttcaag gacgtgtcct actacaaggg gtcgccgtgc ttctccctgt
8821 cgggctgctc ggaaagcgag cgcaaggcga tggaagagaa ccgccgcctg gcgtccgaat
8881 cgcagaaaaa ggccaacgat gcgctgttcc gtggcctcga tcagcagcag agcaacctca
8941 agtccgacgc cgccacgctg gagcaattga agggcaaggc gacgacggcg cagggccagt
9001 tggaagccct cggctacgcc aaccagttcg ccagccagca ggccaaccag ctcatgcaaa
9061 tccgtggcct tctgcttgcg cagcagaacg ccatcgccac gcagatgcag gcccagcagg
9121 accggcaggc ccagcaggac gctgcgggcg cgaagctgcg cgagggttcg taccgcgcaa
9181 gcccgtctaa gacctggtga ggggaggcgc gatgaagaaa tccaacttca tcgcagttgc
9241 cgcgctggcc gccgtcatgg cggccagcct ggcaggctgc gacaacaagc ccgacaccga
9301 caagctgacc tgcgccgatc tgccgaaggt cacggatgcc gctcaacgcg cggagctgtt
9361 gaagaagtgc ccgcgcggag aaccgggagg cttcaagccc agcgaaaaga aagagtggtg
9421 atgacgtatg aaaatccaga ctagagctgc cgcgctcgcg gtcctgatgc tggccttgat
```

9481 gccggtagcg gcatacgccc aaatcgacaa ttcgggcatc ctcgacaacg tattgcagcg
9541 ctaccagaac gccgcgagcg gctgggccac tgtcgtccag aacgccgcaa cctggctgtt
9601 ctggaccttg accgtgatta gcatggtctg gaccttcggc atgatggcac tgcgcaaggc
9661 cgacattggc gagttcttcg ccgagttcgt gcggttcacc atcttcaccg gcttcttctg
9721 gtggctgctg accaacggcc cgaatttcgc gtcgtccatc tatgcgtccc tgcggcagat
9781 tgcaggccag gcaacggggt tggggcaggg gctttcgccg tccggcatcg tcgatgttgg
9841 cttcgagatt ttcttcaagg tgatggacga aacctcgtac tggtcgccgg tcgatagctt
9901 cgtcggtgcc tcgttggcgg ccgccatcct ctgcatcctg gccctggtcg gcgtgaatat
9961 gcttctgctc ctggcgtccg gatggattct tgcctacggc ggtgtgttct tcctgggctt
10021 cggcggctcg cgctggacct cggacatggc gatcaactac tacaagaccg tcctcggggt
10081 cgccgcgcag ctcttcgcaa tggtgctgct cgtaggcatc ggcaagacct tcctcgatga
10141 ctactacagc cgcatgagcg aaggcatcaa cttcaaggaa cttggagtga tgctgatcgt
10201 cggcctgatc ctgctcgttc tggtcaacaa ggtgccgcag ctcatcgccg gcatcatcac
10261 cggcgcgagc gtcggcggtg ctggtatcgg ccagttcggc gctggcacgc tcgtcggtgc
10321 ggccgcgacg gccggcgcgg caatcgcaac tggcggcgca tctatcgcgg ccggcgctgc
10381 ggcggcggcc ggtggcgcgc aggccatcat ggcggccgcg tcgaaggcca gcgataacgt
10441 ctctgccggc actgacattc tgtcgagcat gatgggcggc ggcggtggcg gcggcggtgg
10501 tagcgccggc accagcggcg gcgacggcgg cggctcgggt ggcggcggtg gctcgggcgg
10561 cggtgaaacc ccgatggcct cggccgccgg cgacaacagc agcggcgcac gcggcggcag
10621 ttcgggcggc ggctcgggtg gtggccgttc gtctggcggt atcggtgcca cggcggccaa
10681 gggcggccgg atcgcggccg ataccgtcgc caacctggcg aaaggtgccg gctcgattgc
10741 caaggccaag gccggcgaaa tgcgcgcatc ggcccaggaa cgcatcggcg ataccgtagg
10801 cggcaagatc gcgcaggcaa ttcgcggcgc gggtgcggcg gcgcagaccg ctgcaaccgt
10861 cgccgatagc aacagccagg cgcaggaaca acctgcaccg gcacccgcac cgtcgttcga
10921 cgacaacagc ctttccgcaa gcaacaacag ggaagcggcc gccgacgcgg attccgaagt
10981 ggcgagcttc gtcaacaagc ccgcccaatc ctgaaacgac tcttaggagc tacgaccatg
11041 caactgaaaa aagcgttctc gtcggccgcc ctggtggtgg ccttgggcct cggcgcaact
11101 ggctcggcca gcgcgcaaga cgtgctgacg ggcgataccc gcctggcctg cgaggccatt
11161 ctgtgcctgt ccacgggcag ccggcccagc gagtgcagcc cgtcgctctc gcggtacttc
11221 ggcatccaca agcgcaagct gtcggacacg ctcaaggcgc ggctgaactt cctcaacctc
11281 tgcccggtat cgaaccagac gccggaaatg cagacgctcg tttcctcgat ttcgcgcggg
11341 gccgggcgct gcgatgcgtc ctcgctgaac tccgtgctgc gtgagtggcg gagctgggac
11401 gaccagttct acatcggcaa ccgcctgccg gactactgcg cggcctacac cggccatgcc
11461 tataccgact tcaacacgac cgcgccgcgc tacgtcggca cgccggaaga gggcggctat
11521 tggatcgagg cggccgacta cgaccgcgcg ctcaaggagt acgaggcgaa gctgaaagag
11581 cggcagcagc agtacggtcg ctatggcagc gacgcctacc gtcggttcga gcggtaaggg
11641 gaggggatag cgatgccgtt tgccaagctg ctggcacgga acgctctgcc ggtggtcgcc
11701 ctggtggcgg ccactggctt cggtgcggcg gatgcgaccg ccgcacggct cttccccgat
11761 ctgtcggaac agatggaaga gcgcgttgtg tgctcggtgt ctgcggccgc gaagtacgag
11821 attccggcca acattcttct cgccattcgg gaaaaggagg gcggcaagcc gggccagtgg
11881 gtcaagaaca ccaatggcac ctatgacgtg ggcgagctgc aattcaacac cgcctacctg
11941 ggcgacctgg cgaagtatgg gatcacggcc caggacgttg ctgcggcagg ctgctatccc
12001 tatgacctgg cggcctggcg gttgcgcggg cacattcgca acgacagggg cgatctgtgg
12061 acacgcgccg ctaactatca ctcgcgcacg ccgtcgaaga acgcgatcta tcgcgccgat
12121 ctgatggtga aggccgacaa gtgggcgaag tggctggatg cgcgtttcgt caccgtcaac
12181 tatggcccca gctcgccggc gcagccggca gggaagggga ccacacttgc ggccgctgat
12241 acgtcggcag cagcgccggc cgaagcgcag ccgatgaagc aaggccggat cacccgcacc
12301 agcctccgca gctcgggtta cgtaccccgg cagctcatca tcaacaacac gccataagga
12361 ggaacggccg tttagcggct aaagcctatg ggcattcgca acctgacgca gcgatacatg
12421 aacggggcca gggcctacgc ggcctgggcg gcatcgcagg cgaaagcgcc gtttgatctt
12481 ctggtactgg gcatcgggcc tgtcatcgtc tttggcctgg tcgcgcatac gctgctcgcg
12541 ttcctgccca catgggccat gtacgccgcc ggcgctctgc tggtcctcgc ggccctgcct
12601 ttggcgctgc acgtcctccg ggaatacgcg ctgcgctatg ggcgcaaata gcgccctgca
12661 gggcgttctt actccaaggg ggagggcatg aatacacgcg ccatgaacga cgccagcggc
12721 cgggcctcgc tgcctgccat ggtgatcgcc gacggcacca ttgaagcctt gaagtggctc
12781 gccttgcttg ccatgaccgg ggatcacgtc aacaagtacc tgttcaacgg tacgctgcca
12841 tatctgttcg aggcggggcg cttggccctg cctcttttcg ttttcgtcct ggcgtacaac

12901 ctcgcccgcc cgggcgcgct cgagcgcggt ttgtacgggc gagcgatgaa acgcctgttg
12961 gccttcggcc tggtcgcctc ggtcccgttc attgcgttgg gtggagtggt gggcggatgg
13021 tggccgctga acgtcatgtt cacgctgttg gccgcaaccg cgatgctcta cctggtcgag
13081 cgcggccgct cggtcgctcc tatagcgctg ttcgtcgtgg ccggcggcct ggtcgagttc
13141 tgttggccgg cgctgctgct ggccgcgtct gtctggttgt acctcaagcg cccgacgtgg
13201 gcggccgcgt tgatggcgct gctgtcttgc gcgtccctgt ggtacatcaa tggcaacctt
13261 tgggcgcttg ctgttgtgcc cctggtgatc gtcgccgccg gcgtcgatct tcgtgtcccg
13321 cgcctgcgct gggcctttta cacgtactac ccgctgcatc ttgccgctct ttggctgatc
13381 cgcattccga tgcgcgaggc gggctacttg tttttcacct gacctttgag attccaatat
13441 gcaattgctc aagaaatgca ccatcgcggc cctgccgctg ctcgccctgt ccggctgcgc
13501 actgctgaac atccccatgc cgacgccgcc cggttcgacc ccgccggaaa tgctgaccgt
13561 gccagtggcg caaatctgcc gcgacgctga caagaaccct gttcgggcaa cggagctgta
13621 cggcaagaaa gggttgtcgg ccaccggcaa ggtgcaggtg atttccgaag gcttcaagcc
13681 tcgctatcgg gtgctgctgc gcgctggcag cgcctcggtc catgctggga ccgataacca
13741 gctcgccatc aagtcggttt ccaccggcca gaccacgcgc gtcactggca ccgtgaagga
13801 cgtgtcctac gaccataacg gctgctcgat ctcgcttgac gatgcgaagt tctactgagg
13861 ggagggcggc ggatgctgac acggttgaag ggcttccttg ctcgtcgccg cgagttgaag
13921 gaactggatg tgtccgtggt gagccggccc cggccggctc cggcggaatt ggtccaggtt
13981 gatgcacgcg aggccgtttg gcgcgtgccg gtgcccggcc aggccgaccg cttcatgtcg
14041 gccaagcctg gcgcgatcaa cgatgaaatg ttcgtggttc gggtggacac cgaagcgttc
14101 tatcgggctt ggctgcgcag cagctcgacg ggccgcgaaa cgcggtcgga caactgcccg
14161 ctgcgctcgg aaatgccgca ggactacaag ttcaagcacg ccgtccaggg cttcgcgcac
14221 ggcagggaaa atcctgtgcc gctggccttc gccggcgcgc accaggagcg ccaccgggtg
14281 gacattggtt tcagcaacgg ggtcacgcgc tcgttctggc tgattgccaa caaggctccg
14341 tcgttcccga tccaggtcca cggccgggag tcggccgagc tgctgaacaa ggtttgcggc
14401 ctcgatcctg cgccgctgtc gttcacggaa ctgttcgcgc aggcccaacg ccaggctccg
14461 caggtcgcca caccggcccg gcctgcgccg gcagcggcca cccggccagc tcccaaggtg
14521 cagccacgcc ccggccgaag cggcccgcgc aaaggccgcg gactctgact acaaccgtgc
14581 gcaaggcgca ttagggagga tgtatgtatg taatcgcctg cggcatcgtt gccggcttgg
14641 cggctgcggt ggccctgttg ggcttcacgc cgatgatgga ggcgcttgcc gccggcgaac
14701 gccgcaaggc actcgcgcaa tggacgcgga cgatgttcct ggtgctgctg cctgtcgtgc
14761 tgatgtgcgc gcccatcggg tccagcattt acgacgccgt gcaagcggac gctggcaagc
14821 ccatcgcttt ccacaacggc cggatcacgg tcgtcatggc cctggtcggc agcttggccg
14881 ttgtcctggt cgcggctgcg cgtgcggtgg tcaaccgcaa gcatgccagc ttctggttcg
14941 tcggctgggt gatggcgtcg gttttggccg gcggcgtcgg cgcgatcgcc agcgcgaagc
15001 aactggcgtt cctcggcgaa catagcggca tggtggcctt cggcttcttc cgcgaccagg
15061 tgaaggacat gcactgcgat gcggacgtga tcctggcccg gtgggatgaa aaggcgaact
15121 cgccggtggt ctaccgctgc ccgaaggcgt acctgctcaa caggttcgca tccgcgccct
15181 tcgtgccctg gccggactac accgaggggg aaagcgagga tctaggtagg gcgctcgcag
15241 cggccctgcg ggacgcgaaa aggtgagaaa agccgggcac tgcccggctt tatttttgct
15301 gctgcgcgtt ccaggccgcc cacactcgtt tgacctggct cggctgcat ccgaccagct
15361 tggccgtctt ggcaatgctc gatccgccgg agcgaagcgt gatgatgcgg tcgtgcatgc
15421 cggcgtcacg tttgcggccg gtgtagcggc cggcggcctt cgccaactgg acaccctgac
15481 gttgacgctc gcgccgatcc tcgtagtcgt cgcgggccat ctgcaaggcg agcttcaaaa
15541 gcatgtcctg gacggattcc agaacgattt tcgccactcc gttcgcctcg gcggccagct
15601 ccgacaggtc caccacgcca ggcacggcca gcttggcccc tttggcccgg atcgacgcaa
15661 ccaggcgctc ggcctcggcc aacggcaagc ggctgatgcg gtcgatcttc tccgcaacga
15721 cgacttcacc aggttgcagg tccgcgatca tgcgcagcag ctcgggccgg tcggcgcgtg
15781 cgccggacgc cttctcgcgg tagatgccgg cgacgtagta cccggcggcc cgcgtggccg
15841 ctacaaggct ctcctggcgt tcaagattct gctcgtccgt actggcgcgc aggtagatgc
15901 gggcgacctt caaccttcgt ccctccggtt gttgctctcg cgtcgccatt tccacggctc
15961 gacggcgtgc ggatcggacc agaggccgac gcgcttgcct cgcgcctcct gttcgagccg
16021 cagcatttca gggtcggccg cgcggccgtg gaagcgatag gcccacgcca tgccctggtg
16081 aaccatcgcg gcgttgacgt tgcgcggctg cggcggccgg ctggccagct ccatgttgac
16141 ccacacggtg cccagcgtgc ggccgtaacg gtcggtgtcc ttctcgtcga ccaggacgtg
16201 ccggcggaac accatgccgg ccagcgcctg gcgcgcacgt tcgccgaagg cttgccgctt
16261 ttccggcgcg tcaatgtcca ccaggcgcac gcgcaccggc tgcttgtcta ccagcacgtc

16321 gatggtgtcg ccgtcgatga tgcgcacgac ctcgccgcgc agctcggccc atgccggcga
16381 ggcaacgacc aggacggcca gcgcggcagc ggcgcgcagc atggcgtagc ttcggcgctt
16441 catgcgtggc cccattgctg atgatcgggg tacgccaggt gcagcactgc atcgaaattg
16501 gccttgcagt agccgtccag cgccacccgc gagccgaacg ccggcgaaag gtactcgacc
16561 aggccgggcc ggtcgcggac ctcgcgcccc aggacgtgga tgcgccggcc gcgtgtgccg
16621 tcgggtccag gcacgaaggc cagcgcctcg atgttgaagt cgatggatag aagttgtcgg
16681 tagtgcttgg ccgccctcat cgcgtccccc ttggtcaaat tgggtatacc catttgggcc
16741 tagtctagcc ggcatggcgc attacagcaa tacgcaattt aaatgcgcct agcgcatttt
16801 cccgacctta atgcgcctcg cgctgtagcc tcacgcccac atatgtgcta atgtggttac
16861 gtgtatttta tggaggttat ccaatgagcc gcctgacaat cgacatgacg gaccagcagc
16921 accagagcct gaaagccctg gccgccttgc agggcaagac cattaagcaa tacgccctcg
16981 aacgtctgtt ccccggtgac gctgatgccg atcaggcatg gcaggaactg aaaaccatgc
17041 tggggaaccg catcaacgat gggcttgccg gcaaggtgtc caccaagagc gtcggcgaaa
17101 ttcttgatga agaactcagc ggggatcgcg cttgacggcc tacatcctca cggctgaggc
17161 cgaagccgat ctacgcggca tcatccgcta cacgcgccgg gagtggggcg cggcgcaggt
17221 gcgccgctat atcgctaagc tggaacaggg catagccagg cttgccgccg gcgaaggccc
17281 gtttaaggac atgagcgaac tctttcccgc gctgcggatg gcccgctgcg aacaccacta
17341 cgtttttgc ctgccgcgtg cgggcgaacc cgcgttggtc gtggcgatcc tgcatgagcg
17401 catggacctc atgacgcgac ttgccgacag gctcaagggc tgatttcagc cgctaaaaat
17461 cgcgccactc acaacgtcct gatggcgtac ttacccaaag aacagctagg agaatcattt
17521 atgctcagca cacttccaca agctcatgca actttcttga accgcatccg cgatgcggtc
17581 gcttccgatg ttcgcttccg cgctcttctg atcggcggct cttacgttca cggaggactc
17641 gatgagcact ccgatttgga tttcgacatc gttgttgagg acaactgcta cgcagatgtc
17701 ttgtctacac gcaaggattt tgccgaggca ctgcccggct tcctcaacgc gttcaccggc
17761 gaacatgtag gagaaccgcg ccttctgatc tgcctatatg gtccgccact gctacacatc
17821 gatttgaagt tttctcttgc ttccgatctc gaccagcaaa tcgagcggcg ggcggttctg
17881 tttgctcgtg atccggcaga gatcgagaag cgcattgagg cggcagcggt ggcatggcca
17941 aaccgtccct ccgagtggtt cgaagcacgt tgtcagcgcc agtgatataa gacggtaatt
18001 caccatttgg attgtccgct ccacccaaca tgttgtttcc ttaaggttct cacaccagaa
18061 aggacatcaa catgctgagc agagaggact tttacatgat aaagcaaatg cgccagcagg
18121 gcgcgtacat tgtcgatatt gcgactcaga ttggttgctc tgaacggacg gtcagacgct
18181 acctcaaata ccctgaaccg ccagccagaa agacccgcca caaaatggtt aagctgaaac
18241 cgtttatgga ttacatcgac atgcgcctgg cagagaatgt ctggaatagt gaggttatct
18301 ttgcggagat taaggcaatg ggttatacgg gcggacgttc catgctgcgt tactacatcc
18361 agcccaaacg taaaatgcgt ccgtcaaaaa gaacagttcg cttcgaaact cagcctggat
18421 accagctcca gcacgactgg ggcgaagttg aggtggaggt tgccgggcaa cggtgcaaag
18481 ttaactttgc ggttaatacg ctggggttct cccgccgctt ccatgtcttc gccgcaccaa
18541 aacaggatgc tgagcatacc tacgaatcac tggttcgcgc cttccgctac ttcggtggtt
18601 gtgtgaaaac ggtgctggtt gataaccaga aggctgcggt gctgaagaat aacaacggga
18661 aagtcgtgtt caactccgga ttcctgttgc tggccgacca ctataacttc ctgccacggg
18721 catgccgtcc acgcagggcc agaacaaaag gtaaggttga gcggatggtg aaatacctca
18781 aggagaactt cttcgttcgg taccgcaggt tcgacagctt cactcatgtt aatcaacaac
18841 tggagcaatg gatagccgat gtggctgaca aacgggaact tcgccagttc aaagaaacgc
18901 cggaacagcg cttcgcgctg gagcaggaac atctgcagcc gttaccggat acggacttcg
18961 ataccagtta cttcgacatc cgccatgtgt cctgggacag ctatatcgag gttggtggta
19021 atcgttacag cgttcccgaa gcgctgtgtg gtcagccggt atcgatacga atatcgctgg
19081 atgacgagtt gcggatctac agtaatgaga aactggtggc ctcacatcgc ctctgttcgg
19141 catcgtctgg ctggcagaca gtgccggagc atcacgcccc gctctggcag caggtcagtc
19201 aggtggaaca tcgaccactg agtgcctatg aggagctgtt gtgatgcatg agctggaagt
19261 cctgctgagt cgcctgaaaa tggagcatct gagttatcac gttgaaagcc tgctggaaca
19321 ggcagctaaa aaagagctga actaccggga gttcctgtgc atggcgctac agcaggaatg
19381 gaacggcagg catcagcgcg gtatggagtc caggctgaag caggctcgtc tgccgtgggt
19441 caaaacgctg gagcagttcg actttacctt ccagccgggc atcgaccgta aggttgtccg
19501 ggaactggct ggtctggcgt tcgtggagcg cagcgaaaac gtgatcctgc tgggacctcc
19561 tggtgtcgga aaaactcatc tggccatagc tcttggcgtg aaagcggtgg atgcgggaca
19621 tcgggtactg tttatgccac tggacagact gatcgcgaca ctgatgaaag cgaaacagga
19681 aaaccggctg gagcgtcagc tgcagcaact gagttatgcc cgggtgttga tcctggatga

19741 aataggctat ctgccgatga acagagagga agccagcctg ttcttccggc tactgaaccg
19801 tcgatatgaa aaagcgagca tcatactgac gtcaaacaaa gggttcgcag actggggaga
19861 aatgttcgga gatcacgtgc tggcaacagc gatactggat cggttgctac atcactcaac
19921 cacgctgaat atcaaaggag agagttaccg gttaaaagag aaacgtaaag ctggagtgct
19981 gaccaaaaac acaacgccaa tcagtgatga tgaaatggtg aaaagcggac agcatcagta
20041 acgaaagtat cttagcgggc atgaaaatgg caaataacgg tcaaacatcg tggcgttgac
20101 aacgtgcctg gatctggcta cactatgcgg ccaccaagct cgcccgtggc gagctttacg
20161 aagcgatcgg catgctcggt ttcttccgtg agcaagtgtt aggacctttg ctctaccgtc
20221 gcgctggaaa ggaccagcgc ggagtgaggc gattggaaac ccttcgactg gatgaagagc
20281 gcagactagc caccaccatt gcgctgcacg atgcgttgtc tgtcagggat gccatcaaag
20341 catctgcctc catctatctc gacctccgag ccgccgatcc gtcgttggaa ccgacaacgc
20401 atatgccagg tcttctgtac gacttaatag aacgtgcggt accaggcacg cctaaccgtc
20461 agtgagattg gatgagtgaa cgatattgat cgagaagagc cctgcgcagc cgctgccgtg
20521 cccgagagca tggcggctca cgtgatggga tacaaatggg cgcgtgataa ggttggtcag
20581 tccggctgcg cggtctatcg gctgcatagc aagtcaggcg gctccgactt gtttctgaag
20641 cacggcaaag atgcttttgc cgacgacgtg actgatgaaa tggtgagatt gcgttggctg
20701 gcggggcaca tttctgtgcc ctccgttgta agcttcgttc gcacgcccaa tcaggcatgg
20761 ctcctgacaa cagcaataca tggaaaaacg gcatatcaag tgctgaaatc ggatttcgga
20821 gcccgtctcg ttgttgttga cgcattggcg gcgttcatgc gccgactgca tgcgatccca
20881 gtgagcgaat gctccttcaa cagtgaccac gcatgcaggc ttgcccgagc gcgggagcgt
20941 atcgaggcgg ggggttgttg atgtcgatga cttcgataag gagcgcgaag ggtggacggc
21001 cgaacaggtt tgggaggcga tgcatcgcct cctaccgctc gcgccggacc cagtcgtgac
21061 gcacggcgat ttttcactcg ataatctact tatcgtcgaa ggtaaggtag tcggctgcat
21121 cgacgttggg cgggctggta ttgctgatcg ataccaagac cttgccgtgt tatggaactg
21181 tcttgaggag ttcgaacctt cgcttcagga gaggcttgtt gcgcaatatg gcattgccga
21241 tccggatagg cgcaagctgc aatttcatct cctgctggac gaacttttct aaggcgatgc
21301 cccctcgacc tcgatcaggg aggcgttcag gacgactcac aaagaaagcc gggcaatgcc
21361 cggctttttc tgctgctacc tccgtagtcg taaggtcgtt gcaggtgctc gggtgcggta
21421 caactcgccg gtcgccagct caagcgcgat cacgtcgttg ccgtcgtagt tgacgatgat
21481 gctgttgggc cgactgtcct cacgcttcgc agggagaggc cagccttcaa tcgaagccgg
21541 cgcaagctcg tagtgcttcc cggtttcgac gctgcgcagc gtccaggtcc tgcaaccggc
21601 cacgccggtc gcagaaacca cggcgagcga gccgcgaaaa tcgtgcgggt acgcctcgat
21661 gttcatacgc ctcctagatc gagcgcgagc gtttctgctc ggccttggcc gcctgttcct
21721 gggacacctc gccgatgacc ttgccctggc cccggctgta ggcgatttcg tagttcttgc
21781 cgacaaccgg cggcttctca aagatgcccc ggctgtgttt cacgatcccg ccttcgctga
21841 actggtagac gttgcgccca tcgtcgtgca gcacctggcc gacgtgcttg tgcgggtgga
21901 cgtttttgct tgcgtccttc gcatcgctca actggtgaat gcctttcggt agccccgcct
21961 cgggcaacac cttcatggtc agccattcgc cgttcactac ctggtccacc tggcggctgc
22021 cgttcatgac ggcgatcttg acgctgccct cgggtttcat gatgacgcca gggcttgccg
22081 atgtgcgtgg tgccccgatc tgtactttgt tcatacgctc tagttctcct tagtaggttc
22141 tcgcgcggcg ttgccgctgt tcttgctgct cgatgtcttg ctgcttgagc tgctgcacct
22201 tctgccgctg gccctcgtcg agaagcacct tgccgacagc acttctcacc tggcgttcaa
22261 ccccgtcctt gcccaggctg ctgcgctcgg acaggtcgtt gaaatcggtg tgcttcttca
22321 tgttcgacag ggcggcgagc tggccatcgt tcaacagcga ttccttgagc ttggccgtgt
22381 cggcctcgct caactggacc ttgccggccg ccgcgtccgc gaggcgcttt tcagcgtgca
22441 aatggttgcg gtagttctcc ggggtgatcg gcggcagctc cttcgggtag gcgttctcgc
22501 ccggcgcgaa gattgggaag atggccttgc cgccgaccgc cttggcggcc tcctgtgcct
22561 tcgtcctgcc gggattcacg ccctgggtga tctgcacctg gcggtcgtcg tcgccggcga
22621 tcacaacggg cttgtccggg aatttcgcgt gcagggcctc ggcaacagcc tgtaggttgc
22681 cggaatcgaa cgcggcgaca gtcgcgtgcc ccagcgcttc ggccactgtg gcggcggtgg
22741 catagccttc gccgatcacc agcgccggcg cggccgcgag cgcatccatg ccaccgacga
22801 catggaagca tccttccttg cggctgtcct tggcgaagcg cttggtgccg tcctcctgga
22861 tgtactgcat ggtccattgc ttgccgtcgg cgtcgtaggc cgggatgtag gttttctggc
22921 cctcctggtc ggtaaggacg ccggcgtgca cctgtagacc cttgtcgcgc aggtacggcg
22981 tcggttccgt gatgggaacc aggctttgcg cctggcggcc gatgcgctgc gccgtggctt
23041 cgtgctggcg ttcttgttcc tcggcacgcg cggccagctt ggccgccgcc tcggcctgca
23101 tcttggcctt ctcggcgggg tccagggcgt agcccttggc cttccacttc atttcgacgc

23161 cggtgcggtt gtttttgatg taaccggccg ggtggccgtc gaggtggccg acgtagaagc
23221 ccgacttctc gcccttcttg tcgccctcgg tctcgatgcg gtgcttcttg ccgtccatga
23281 tggggtgctc gccgcctggg gtgacgacgc agcccatgct tttcagggcc tccgcgaact
23341 catcttcggg ggtgacggcc ggggattgct gggtgggcac gttgtccggc agccagcgtt
23401 gcagcttgcc catgtcggcg ttcggtccgg cgtaccagga cttggccacc ttgtcccact
23461 gcgcgccggc cgccttggca acctggcgct cgccgtaggg cacggccagg tagacgcgct
23521 cctgggccgc gttggggcgc tcggccgtgg gttgggtagg ctgggcctcg gcgcgggcct
23581 ctacggccgc tgtagcgccc tcgcgcgccc atttggcgaa cggggcaggg tcaacccctg
23641 ccggaacgta ccaggcgcgt tcctggcggt cccagcgcgc tccaagggcc ttcacctcgt
23701 ctttctcctt gaacggcacg ttcaagtagg cgcgctcggg cttggcgggg gcttgagcgg
23761 ccgccggctg ctcggcggcg ttcatggcct gggccatttc ctgctgctcg cgctcgtagt
23821 cggcgatccg gcgctgtagg tcctcgtcgt gcagcatggc cgtgccctcg gcggccttgc
23881 gcgcctcctt ggcggcaacg cggtcctcgt cggtgctgtt gggatcgcgg cgaactcgct
23941 cttcatggat gcgggcgaac ttcgcggcct gctcgtactc gttggccgtc gcgtaagcgt
24001 cgatcacggc caggcggtcg gcgagcgctt cggcattggt ctgcgcgtcg ggccggcgga
24061 agtcggcaag ccattggtgg ccgccccacg catggttcgc atagacgccc caaaactccg
24121 gctctcggtc gccggccggc accacggacc gttcgccgtc gtgctcgacc tcgacgttgg
24181 cctggacctg gacgcggccg gtccaatcgg caggcagctc aaagcccagc gtggtttcgg
24241 tcagcgcggc cagcgattgg ttgccttccg ccggctccgc gccggcgcgg tacatgcgca
24301 gggtctgcgc gatcagctcg tcggccggcg caatggccgg tcgtgccacc tggtcttgct
24361 gttgctccat agttgccccc tgcgcaggct cgatggcctg ctgggtcgtt tgttcttgaa
24421 tttgcttctg ctcgaacgcc aggacgaaat cctggatctt ctccgcgtcg gcggccgcgc
24481 ggaaaatctc tagcgggtcc tcttgtagcg ccttgatcca cgatccgaca taggccgcgt
24541 gctggccggg gtcgtggccg atgcccagct cgtcgcccag gatcatgctg gcaatctcgg
24601 cccgcagctc ttccttggcg taccccctcgc tcccgaaggg atgcgccagg tcgcggtcca
24661 gccgcgacgg gtggccggtc cagtgcccca gctcatggag cgcggttgcg tagtagttgt
24721 cggcgctcgg gaactggcct ttgtcgggca gatggatgct gtccgtggac ggccgataaa
24781 acgcgcggtc gtgctcgccg tggcggatgg tggcacctga cgccgcaagg atgtgctcgg
24841 cccgctcgac ggcgctccaa gtctgttcct tgcgttccaa cggcggcagg ccgtcgatct
24901 gctccgcatt gaacacggtg gcgaagaaca cgcgcgggcg ttcgagctgc accgtcacct
24961 tgaccggatc gccgttggca tcgaggaccg gcttgccggt ctgctcgtcg gtcttggtct
25021 gctcttcgct gaacttccaa tactggatcg gcgtgccttt ctcgccgcga cgcacctgtg
25081 cgccggcggc agcggcctgc ttgtaggtca tccagcgcgg gtccgcatgg ccctgggcca
25141 tgagctgaat cgcgttgatg cccttgtaac gcttcccggt agtcgggttg agcgggatga
25201 aggagccggg catgcccggt tcccacggtt tttgccacgg cgcagtgccg gctttcagtt
25261 gctcaatgag gcgttcggca acctgctcgt ggaacggctt tttgacctct gccatagcca
25321 attacctccc gtcattggcg gccgcggtcg tcgtgtcctc gggcacggtc gcgtccacca
25381 ggtcaatgtc gctctcggcg gcgtcctgct cgctcaaggc gtcctcgggg aaggccccgg
25441 ccttctccgc ttcttccggg tcgaactcga cctggaagcc gggcgtcatc gcatcggcga
25501 gcttttcacg cagggcggcg gcggcgttcg cggtatcgtc gaacggcgca aagtcgtcct
25561 gctgcacggt cttgggatca ttcatcgctt tcactcctgg ttggtgccgt tacggccttt
25621 gctgtagtcc ggcctgcctt tcaggtcggg gtatgtctgc ttgcacgtcg ggaagttgct
25681 gcaccccac cagaacatgc cgcgcttctt gccaggccga cgggaaaggc cgtggccgca
25741 ggccatgcac ttgtgcagct cggagacttt cggggcctcg cgcgggacgg gcttgccgcc
25801 cttgtcgtcg cacgcgaact tgcagccgtc ggcaaagccg gtgcagcccc aaaagtattc
25861 gttcttgtcc ttcttcttga ggcgtcgcag cggcttgccg caggacgggc aagggtgcgt
25921 gtcgatcttc atgttgaggc cgttgtcttt gatgttggcg acctcggcgc cgatgtattc
25981 catcagctcg ttgacgaacg acagcgtgtc gcgctcgccg gcctggatgg ccttctgctg
26041 ctcatgccag agcgcggtca tgtcggggaa tctggccgtg tcgggcagtg cgtcgtacag
26101 ctcttcgccg gtcggcgtgg acacgatgtg cttgcccttc tccaccaggt agccgcgctc
26161 gaaaagcgtg gcgatgatgg agtctcgcgt tgccggcgtg ccgatcccgc cgtgctcgcc
26221 ttgcttgccc ttgtcctttt cgatcaagat tttccgcagg cggtcatcgc ggatgtattt
26281 cgcaacgcgg gtaaggtccg acagcagaga ttccatcgtg tacagcggct gcggtttcgt
26341 ctcctgctgc tcggccttcg catcggtgca ggtgccggcc tggccgtcac gcagcttgcg
26401 caggtcctgt tcaatgtcgt cggcattgcc ttccaggtcc tcgttgccgg cgtcgttctt
26461 gtagagaatc ttccagcccg gcgacgtggt gacgttcgag cgcacgccga aacgatgatc
26521 gccgacctgg gcaagcacgt cggtctggtc atacagatgc ttcggccaga actgcgcgac

26581 gtaggcgcgc gcgatcagca ggtaaatctt ctgctcggca tcggtgagct tcgacaggtc
26641 ggccgtgctt tcggtcggga tgatcgcgtg gtgcgcggaa accttggacg agttgaaggc
26701 gcggctcttg atcgtcggat tggcgcgctg cgcagcagcg gccagcatgg gggccgtctg
26761 tgcgatggcc gccagcacgc ccggcgcatc gccgtgctgt tcctcgctca agtattcgca
26821 gtcggaacgg ttgtaggtga tgagcttgtg cttctcgcgc agggcctgcg taatgtcctt
26881 cacctggtcc ggcttgaagc cgaacttgcg cgaggcgtcc atttgcagtt tcagcaggtt
26941 gtagggcagc ggcgcagccg cttccttcgc cttggtggtc acggacacga tgcgggcggg
27001 ttggccgctc acggcggccg cgatgccctc ggcgtgctcc ttgttgctga ggcggccttt
27061 ctcgtccacc ggatcgccgt cggcgacctg gtaacgggcc gggaactgaa tgccctcgac
27121 ctcgaactgg ccgttcacca ggtagtagta ggttttctgg tgggccgcgt tctcgcggca
27181 acggcgcacg acaaggccca ggatcggagt ctgcacgcgc cccacgctca acagcccctg
27241 atagcccttc gcgcgtgccg caagcgtgta caggcgcgtg atgttgaagc cgtatagctg
27301 gtcgccgacg ctgcgggcct cggccgcagc ggacaggccg gcgaactcgc ggttgtcgcg
27361 catcgcggcg agctgccggc gcacgatctt cacgttgttg tcgttgataa gcagccgctg
27421 caccggcaga cggcagttgg cgtattccag gatttcatcg accagaagct ggccttcgtc
27481 gtccgggtcg ccggcgtgaa ccacgctttt cgcctgcttc aacaggctga ggatggtctt
27541 gaactgagct ttcgcaccgg catcgccgga cggtttcttg cgccagggaa tatggacgat
27601 gggcaggtcg gccatgttcc agttggcgta gcgctcgtcg tagtcctccg ggtctagcaa
27661 ggccagcatg tgaccgtagc accaggtcac gcggtcggag ccgcattcgt aatagccgtc
27721 cttgcggctg ccgccgccca ggccctcgac gatggctttt gccagctccg gtttttcagc
27781 gattacaagg cgttcaaatt gcatatatcc ccctaccctc accaggtcag aaccggcctg
27841 atgacggtga tgatttgcga acgattgaca ggcccgaagt agcggccgtc gaaagacgtg
27901 tcgcttacgt cggacataag cagaacctcg gcggtcccca gggtgtagct gtcggactga
27961 taacgaggca gcggccgtcc tgatggatcg gccttgatga gcgcgctgtg aggcagcagc
28021 ccgccattca cgcgcacgcc ggcgtcggtg atggcaacct cgtcgccttt agcggctaaa
28081 actcgcttca tcatgtagcc gtagtcgccg gggcagaaac cgccggcgat gtagccccgc
28141 tccttggcgt ccgaaaacac gccgacttgc ggcgggcaga acatgacgta agccccctc
28201 tccaccggcg cattcgattt ccagtacagg ccgaccggaa tgcttttggt ggtgttgacc
28261 ttcgcgccgg cgagataggc cgcgccggcg agcaacaagg ccgcgccgcc tccgatggcg
28321 acgtacttgg tgaggcgctg gaagcggctc atatcgtgat cccctcccct tcctcgacgg
28381 tggccgtctg gatcagcttg tcgctgacct tcggagccgg tacggccgcg cgggcctgga
28441 atatcgggtc tttgaagtag agcggctgct tgccgtagat cgcgggatag ccggcgacgt
28501 acacaaccat gtcgcccgcc tcttcaatgc tgccgtcggc gctcttcttc ggccccggca
28561 tgcgcaggca ttcatcgggg gtcagcaatg gccgctgcac ttcctggaag gtccgcgaga
28621 cgttgcccaa cagcgccgac gtgcggcggc cgctcgtcgt gatctgctcc ttcacgatgg
28681 tcgtggtgcc tgtcagtttt gacaggtgct cggccgtctc cacgcggttc ggcgggtagg
28741 cgttctgcac gtggcagttc gacgtgatgc tttcgtcgtg gccgtagccg gtttcgcggc
28801 tcttgagctg gttaatgtcc tggcagatga ggtagcactt gatgccgtag ccggcgacga
28861 aggcaaggga ctcttgcagg atttcgagct tgcccaggct ggggaactcg tcgagcatca
28921 tcagcagacg atgcttgtag tgcgcgacag gacggccgtt ctcgaagtcc atcttgtcgg
28981 ccagcagccg gacgatcatg ttgaccatga cgcgcaccag aggccgcaga cgggccttgt
29041 cgttgggctg cgtcacgatg aacaggctta ccgggtcgtc gtggtgcatc agttgcttga
29101 tgcggaagtc ggacttgctg acgttgcggg ccacaaccgg gtcgcggtac agggccaggt
29161 aggacttggc ggtggacagc acggaaccgg attcttcctc cgggcggtcc atcatgtcgc
29221 gggccgcaga gccgaccgca gggtggttct gcccgtcaac gtggccgtag gtggtcattt
29281 ccatccaaag ctcgcccacg tcgcggttcg ggtcggcaag catgccgtcc accgacggca
29341 gggtggccgg cgtaccctcg ttcttagcct tgtagagcgc gtgcaggatg acgccgacaa
29401 gcagcgcctg gctggttttc tgccagtgcg attccaggcc cttgccgtcc ggatcgacga
29461 tcagggtggc aaggttctgc acgtcgccaa cctcgtactc ggtccccaag cggatttcat
29521 cgagcgggtt ccagcacgcg ctaccctgcg cggatgccgg ctcaaagcgc acgaccttgt
29581 tgcgggcatg cttcttccgc cagccggcgg tcagcgccca caactcgcct ttcaggtcgg
29641 tgatgacggc gctgtgcgcc caggaaagca gcgtcggaac gaccaggccg acgcccttgc
29701 cggagcgcgt cggcgcgtag gtcaagacgt gctcggggcc gttgtgccgc aggtagtgga
29761 acttgccgtc cttgtcctgc cagccgccca catagacgcc gctggaagtg ggcgggtgtt
29821 tgcctgacac cagctcgacg acggtgcgcg gccggggcag caggccggcg gcctgtatgt
29881 ccttcttgtc ggcccagcgg gccgaaccgt gcagatagtc gttcgccttg ccggtgttcg
29941 ccttgaccat ctgcgtgacg gccgtgccca gcaggcccac ggtcgaaacg accataccca

30001 tgctggccgc gcgcatgaaa tcgtcgggat attggccgta ccacttgccg gcccattgaa
30061 ggatcgacca gggcgtgtag acgtggttga tattccagcc aagtccggcc tgatactgga
30121 aggaatgggc gaaatattgc gtcgcggtct gcaagcctgc cccaagggac aggccggcga
30181 ggatgggaac ggtcttgctg gccttcggtt ttttcgcccg tatctgtggc cccacggcgt
30241 tgtttcggtt cttcatctac tcctacctcg ggtagtttta agggagcctc gcggggtcac
30301 ggtgacggga tcaccgatgg cgaggcgctt catgcgttgc accgtggcct tatcgacggg
30361 cagcaccaga atctcgtcgt tttctttcct caacagggcc agcgcctggt cctcgacgtt
30421 ccgggtgcct gcataggaca gcgcaccaac ataatcagta tatcgtgcat gcttcggtat
30481 atcgaagccg tttagccgct tttgctcgcg ctcggcaaca tatttctcgg ccgccgcgat
30541 ctgttcgggc tttagccctc ttcctggccc agaaactccc cgtcgcagtg cgtgagctgg
30601 ttcggctcct tgctgctcca cgtgaccagg aacatcacgc ggcaatagca tttcagctcc
30661 gccggcgatg cgaaccacac cgagttggga cagcgctcgc aaacggtttt ggctttgggg
30721 cggcggcttt cgtccaatgc gtccaacgtt gggcttgcgg agtgcgacgg ttccgccggc
30781 gctgacggcg cgagcgtccc gtcggtcgcc gtcgccgcct gtggcgttga gggtggttct
30841 ggctgcggca ggtcgaatgc ctccatcgcc gccgcgatct cttcgtccgt catttcgttc
30901 gggttgctca tgtgcttgct ccttcgtcag tagttcttga cggcggcgct caagggcggc
30961 gtcgtcaaag gtgattgcca gacggccagc ggcggccgcc tgcgcgatcc gctccttgaa
31021 ctctgctgtg ccgttgacgg tgatccggtc gccgaagcgc tccattgcca ggcgcagggc
31081 ggcgtccagg ccgtccgtgg tggcctcgcg cgagacttgc aggcggtcgc cgtcgtcgcg
31141 gacggcgctg ctgccgacgc gatagatgat ggttcccttc ttcgtgatgt tgtccgtcac
31201 ggccgcatgg cccggcttgg cctcgccgct gccctggatg gtgttgccct tgaggtcgct
31261 gcggccctcg cgtgcgcgca gcgcggccag ggccttgtcg tcgcccttca tcgcctcggc
31321 cttgagccag tcggcccacg cgcggcgctg cgtgcgctcc tggaccgcct gacggccctg
31381 ccggtactcg cggttgatct tgtccaggtc ggcgcgcaga gccttgtgcg cctgcgcgta
31441 catcagtcgc tttgcaatgc gcccctcgcc cagcagcttg atagcggcgc ggcgcagccg
31501 gttgctgcgc atcgcggctt caatcaggcg gtcacgacgc cggcgcagcg tgtccagctc
31561 gcccttgcgc acggccccca tttcctggcg ttcagactga taccgggcgt atagctcggt
31621 ggtgtcgatg cgggtcttga gcggcttcgc tcgatactcc cgccgccggg gggcttcgcc
31681 gccctcggct ggcgtgaatg ccccgaatcg ggcttcgagc ttcggcttgg acaggtcgcg
31741 cgaaacggtg ctggccttga ccgtcgtgcc gtcgccggcc tcgaagatga agccgtttcc
31801 gcgctcgcgc agcttaagcc cgttttcccg caggacgcgg tgcaggtcct cccaggattg
31861 cgccgcttgc agctccggca ggcattcgcg cttgatccag ccgaccaggc tttccacgcc
31921 cgcgtgccgc tccatgtcgt tcgcgcggtt ctcggaaacg cgctgccgcg tttcgtgatt
31981 gtcacgctca agcccgtagt cccgttcgag cgtcgcgcag aggtcagcga gggcgcggta
32041 ggcccgatac ggctcatgga tggtgtttcg ggtcgggtga atcttgttga tggcgatatg
32101 gatgtgcagg ttgtcggtgt cgtgatgcac ggcactgacg cgctgatgct cggcgaagcc
32161 aagcccagcg cagatgcggt cctcaatcgc gcgcaacgtc tccgcgtcgg gcttctctcc
32221 cgcgcggaag ctaaccagca ggtgataggt cttgtcggcc tcggaacggg tgttgccgtg
32281 ctgggtcgcc atcacctcgg ccatgacagc gggcagggtg tttgcctcgc agttcgtgac
32341 gcgcacgtga cccaggcgct cggtcttgcc ttgctcgtcg gtgatgtact tcaccagctc
32401 cgcgaagtcg ctcttcttga tggagcgcat ggggacgtgc ttggcaatca cgcgcacccc
32461 ccggccgttt tagcggctaa aaaagtcatg gctctgccct cgggcggacc acgcccatca
32521 tgaccttgcc aagctcgtcc tgcttctctt cgatcttcgc cagcagggcg aggatcgtgg
32581 catcaccgaa ccgcgccgtg cgcgggtcgt cggtgagcca gagtttcagc aggccgccca
32641 ggcggcccag gtcgccattg atgcgggcca gctcgcggac gtgctcatag tccacgacgc
32701 ccgtgatttt gtagccctgg ccgacggcca gcaggtaggc cgacaggctc atgccggccg
32761 ccgccgcctt ttcctcaatc gctcttcgtt cgtctggaag gcagtacacc ttgataggtg
32821 ggctgccctt cctggttggc ttggtttcat cagccatccg cttgccctca tctgttacgc
32881 cggcggtagc cggccagcct cgcagagcag gattcccgtt gagcaccgcc aggtgcgaat
32941 aagggacagt gaagaaggaa cacccgctcg cgggtgggcc tacttcacct atcctgcccg
33001 gctgacgccg ttggatacac caaggaaagt ctacacgaac cctttggcaa aatcctgtat
33061 atcgtgcgaa aaaggatgga tataccgaaa aaatcgctat aatgaccccg aagcagggtt
33121 atgcagcgga aaagcgctgc ttccctgctg ttttgtggaa tatctaccga ctggaaacag
33181 gcaaatgcag gaaattactg aactgagggg acaggcgaga gacgatgcca aagagctaca
33241 ccgacgagct ggccgagtgg gttgaatccc gcgcggccaa gaagcgccgg cgtgatgagg
33301 ctgcggttgc gttcctggcg gtgagggcgg atgtcgaggc ggcgttagcg tccggctatg
33361 cgctcgtcac catttgggag cacatgcggg aaacggggaa ggtcaagttc tcctacgaga

33421 cgttccgctc gcacgccagg cggcacatca aggccaagcc cgccgatgtg cccgcaccgc
33481 aggccaaggc tgcggaaccc gcgccggcac ccaagacgcc ggagccacgg cggccgaagc
33541 agggggggcaa ggctgaaaag ccggcccccg ctgcggcccc gaccggcttc accttcaacc
33601 caacaccgga caaaaaggat ctactgtaat ggcgaaaatt cacatggttt tgcagggcaa
33661 gggcggggtc ggcaagtcgg ccatcgccgc gatcattgcg cagtacaaga tggacaaggg
33721 gcagacaccc ttgtgcatcg acaccgaccc ggtgaacgcg acgttcgagg gctacaaggc
33781 cctgaacgtc cgccggctga acatcatggc cggcgacgaa attaactcgc gcaacttcga
33841 caccctggtc gagctgattg cgccgaccaa ggatgacgtg gtgatcgaca acggtgccag
33901 ctcgttcgtg cctctgtcgc attacctcat cagcaaccag gtgccggctc tgctgcaaga
33961 aatgggcat gagctggtca tccataccgt cgtcaccggc ggccaggctc tcctggacac
34021 ggtgagcggc ttcgcccagc tcgccagcca gttcccggcc gaagcgcttt tcgtggtctg
34081 gctgaacccg tattgggggc ctatcgagca tgagggcaag agctttgagc agatgaaggc
34141 gtacacggcc aacaaggccc gcgtgtcgtc catcatccag attccggccc tcaaggaaga
34201 aacctacggc cgcgatttca gcgacatgct gcaagagcgg ctgacgttcg accaggcgct
34261 ggccgatgaa tcgctcacga tcatgacgcg gcaacgcctc aagatcgtgc ggcgcggcct
34321 gtttgaacag ctcgacgcgg cggccgtgct atgagcgacc agattgaaga gctgatccgg
34381 gagattgcgg ccaagcacgg catcgccgtc ggccgcgacg acccggtgct gatcctgcat
34441 accatcaacg cccggctcat ggccgacagt gcggccaagc aagaggaaat ccttgccgcg
34501 ttcaaggaag agctggaagg gatcgcccat cgttggggcg aggacgccaa ggccaaagcg
34561 gagcggatgc tgaacgcggc cctggcggcc agcaaggacg caatggcgaa ggtaatgaag
34621 gacagcgccg cgcaggcggc cgaagcgatc cgcagggaaa tcgacgacgg ccttggccgc
34681 cagctcgcgg ccaaggtcgc ggacgcgcgg cgcgtggcga tgatgaacat gatcgccggc
34741 ggcatggtgt tgttcgcggc cgccctggtg gtgtgggcct cgttatgaat cgcagaggcg
34801 cagatgaaaa agcccggcgt tgccgggctt tgtttttgcg ttagctgggc ttgtttgaca
34861 ggcccaagct ctgactgcgc ccgcgctcgc gctcctgggc ctgtttcttc tcctgctcct
34921 gcttgcgcat cagggcctgg tgccgtcggg ctgcttcacg catcgaatcc cagtcgccgg
34981 ccagctcggg atgctccgcg cgcatcttgc gcgtcgccag ttcctcgatc ttgggcgcgt
35041 gaatgcccat gccttccttg atttcgcgca ccatgtccag ccgcgtgtgc agggtctgca
35101 agcgggcttg ctgttgggcc tgctgctgct gccaggcggc ctttgtacgc ggcaggggaca
35161 gcaagccggg ggcattggac tgtagctgct gcaaacgcgc ctgctgacgg tctacgagct
35221 gttctaggcg gtcctcgatg cgctccacct ggtcatgctt tgcctgcacg tagagcgcaa
35281 gggtctgctg gtaggtctgc tcgatgggcg cggattctaa gagggcctgc tgttccgtct
35341 cggcctcctg ggccgcctgt agcaaatcct cgccgctgtt gccgctggac tgctttactg
35401 ccggggactg ctgttgccct gctcgcgccg tcgtcgcagt tcggcttgcc cccactcgat
35461 tgactgcttc atttcgagcc gcagcgatgc gatctcggat tgcgtcaacg gacggggcag
35521 cgcggaggtg tccggcttct ccttgggtga gtcggtcgat gccatagcca aggtttcct
35581 tccaaaatgc gtccattgct ggaccgtgtt tctcattgat gcccgcaagc atcttcggct
35641 tgaccgccag gtcaagcgcg ccttcatggg cggtcatgac ggacgccgcc atgaccttgc
35701 cgccgttgtt ctcgatgtag ccgcgtaatg aggcaatggt gccgcccatc gtcagcgtgt
35761 catcgacaac gatgtacttc tggccgggga tcacctcccc ctcgaaagtc gggttgaacg
35821 ccaggcgatg atctgaaccg gctccggttc gggcgacctt ctcccgctgc acaatgtccg
35881 tttcgacctc aaggccaagg cggtcggcca gaacgaccgc catcatggcc ggaatcttgt
35941 tgttccccgc cgcctcgacg gcgaggactg gaacgatgcg gggcttgtcg tcgccgatca
36001 gcgtcttgag ctgggcaaca gtgtcgtccg aaatcaggcg ctcgaccaaa ttaagcgccg
36061 cttccgcgtc gccctgcttc gcagcctggt attcaggctc gttggtcaaa gaaccaaggt
36121 cgccgttgcg aaccaccttc gggaagtctc cccacggtgc gcgctcggct ctgctgtagc
36181 tgctcaagac gcctcccttt ttagccgcta aaactctaac gagtgcgccc gcgactcaac
36241 ttgacgcttt cggcacttac ctgtgccttg ccacttgcgt cataggtgat gcttttcgca
36301 ctcccgattt caggtacttt atcgaaatct gaccgggcgt gcattacaaa gttcttcccc
36361 acctgttggt aaatgctgcc gctatctgcg tggacgatgc tgccgtcgtg gcgctgcgac
36421 ttatcggcct tttgggccat atagatgttg taaatgccag gtttcagggc cccggcttta
36481 tctaccttct ggttcgtcca tgcgccttgg ttctcggtct ggacaattct ttgcccattc
36541 atgaccagga ggcggtgttt cattgggtga ctcctgacgg ttgcctctgg tgttaaacgt
36601 gtcctggtcg cttgccggct aaaaaaaagc cgacctcggc agttcgaggc cggctttccc
36661 tagagccggg cgcgtcaagg ttgttccatc tattttagtg aactgcgttc gatttatcag
36721 ttactttcct cccgctttgt gtttcctccc actcgtttcc gcgtctagcc gacccctcaa
36781 catagcggcc tcttcttggg ctgcctttgc ctcttgccgc gcttcgtcac gctcggcttg

36841 caccgtcgta aagcgctcgg cctgcctggc cgcctcttgc gccgccaact tcctttgctc
36901 ctggtgggcc tcggcgtcgg cctgcgcctt cgctttcacc gctgccaact ccgtgcgcaa
36961 actctccgct tcgcgcctgg tggcgtcgcg ctcgccgcga agcgcctgca tttcctggtt
37021 ggccgcgtcc agggtcttgc ggctctcttc tttgaatgcg cgggcgtcct ggtgagcgta
37081 gtccagctcg gcgcgcagct cctgcgctcg acgctccacc tcgtcggccc gctgcgtcgc
37141 cagcgcggcc cgctgctcgg ctcctgccag ggcggtgcgt gcttcggcca gggcttgccg
37201 ctggcgtgcg gccagctcgg ccgcctcggc ggcctgctgc tctagcaatg taacgcgcgc
37261 ctgggcttct tccagctcgc gggcctgcgc ctcgaaggcg tcggccagct ccccgcgcac
37321 ggcttccaac tcgttgcgct cacgatccca gccggcttgc gctgcctgca acgattcatt
37381 ggcaagggcc tgggcggctt gccagagggc ggccacggcc tggttgccgg cctgctgcac
37441 cgcgtccggc acctggactg ccagcggggc ggcctgcgcc gtgcgctggc gtcgccattc
37501 gcgcatgccg gcgctggcgt cgttcatgtt gacgcgggcg gccttacgca ctgcatccac
37561 ggtcgggaag ttctcccggt cgccttgctc gaacagctcg tccgcagccg caaaaatgcg
37621 gtcgcgcgtc tctttgttca gttccatgtt ggctccggta attggtaaga ataataatac
37681 tcttacctac cttatcagcg caagagttta gctgaacagt tctcgactta acggcaggtt
37741 ttttagcggc tgaagggcag gcaaaaaaag ccccgcacgg tcggcggggg caaagggtca
37801 gcgggaaggg gattagcggg cgtcgggctt cttcatgcgt cggggccgcg cttcttggga
37861 tggagcacga cgaagcgcgc acgcgcatcg tcctcggccc tatcggcccg cgtcgcggtc
37921 aggaacttgt cgcgcgctag gtcctccctg gtgggcacca ggggcatgaa ctcggcctgc
37981 tcgatgtagg tccactccat gaccgcatcg cagtcgaggc cgcgttcctt caccgtctct
38041 tgcaggtcgc ggtacgcccg ctcgttgagc ggctggtaac gggccaattg gtcgtaaatg
38101 gctgtcggcc atgagcggcc tttcctgttg agccagcagc cgacgacgaa gccggcaatg
38161 caggcccctg gcacaaccag gccgacgccg gggcaggggg atggcagcag ctcgccaacc
38221 aggaaccccg ccgcgatgat gccgatgccg gtcaaccagc ccttgaaact atccggcccc
38281 gaaacacccc tgcgcattgc ctggatgctg cgccggatag cttgcaacat caggagccgt
38341 ttcttttgtt cgtcagtcat ggtccgccct caccagttgt tcgtatcggt gtcggacgaa
38401 ctgaaatcgc aagagctgcc ggtatcggtc cagccgctgt ccgtgtcgct gctgccgaag
38461 cacggcgagg ggtccgcgaa cgccgcagac ggcgtatccg gccgcagcgc atcgcccagc
38521 atggccccgg tcagcgagcc gccggccagg tagcccagca tggtgctgtt ggtcgccccg
38581 gccaccaggg ccgacgtgac gaaatcgccg tcattccctc tggattgttc gctgctcggc
38641 ggggcagtgc gccgcgccgg cggcgtcgtg gatggctcgg gttggctggc ctgcgacggc
38701 cggcgaaagg tgcgcagcag ctcgttatcg accggctgcg gcgtcggggc cgccgccttg
38761 cgctgcggtc ggtgttcctt cttcggctcg cgcagcttga acagcatgat cgcggaaacc
38821 agcagcaacg ccgcgcctac gcctcccgcg atgtagaaca gcatcggatt cattcttcgg
38881 tcctccttgt agcggaaccg ttgtctgtgc ggcgcgggtg gcccgcgccg ctgtctttgg
38941 ggatcagccc tcgatgagcg cgaccagttt cacgtcggca aggttcgcct cgaactcctg
39001 gccgtcgtcc tcgtacttca accaggcata gccttccgcc ggcggccgac ggttgaggat
39061 aaggcgggca gggcgctcgt cgtgctcgac ctggacgatg gcctttttca gcttgtccgg
39121 gtccggctcc ttcgcgccct tttccttggc gtccttaccg tcctggtcgc cgtcctcgcc
39181 gtcctggccg tcgccggcct ccgcgtcacg ctcggcatca gtctggccgt tgaaggcatc
39241 gacggtgttg ggatcgcggc ccttctcgtc caggaactcg cgcagcagct tgaccgtgcc
39301 gcgcgtgatt tcctgggtgt cgtcgtcaag ccacgcctcg acttcctccg ggcgcttctt
39361 gaaggccgtc accagctcgt tcaccacggt cacgtcgcgc acgcggccgg tgttgaacgc
39421 atcggcgatc ttctccggca ggtccagcag cgtgacgtgc tgggtgatga acgccggcga
39481 cttgccgatt tccttggcga tatcgccttt cttcttgccc ttcgccagct cgcggccaat
39541 gaagtcggca atttcgcgcg gggtcagctc gttgcgttgc aggttctcga taacctggtc
39601 ggcttcgttg tagtcgttgt cgatgaacgc cgggatggac ttcttgccgg cccacttcga
39661 gccacggtag cggcgggcgc cgtgattgat gatatagcgg cccggctgct cctggttctc
39721 gcgcaccgaa atgggtgact tcaccccgcg ctctttgatc gtggcaccga tttccgcgat
39781 gctctccggg gaaaagccgg ggttgtcggc cgtccgcggc tgatgcggat cttcgtcgat
39841 caggtccagg tccagctcga tagggccgga accgccctga gacgccgcag gagcgtccag
39901 gaggctcgac aggtcgccga tgctatccaa ccccaggccg gacggctgcg ccgcgcctgc
39961 ggcttcctga gcggccgcag cggtgttttt cttggtggtc ttggcttgag ccgcagtcat
40021 tgggaaatct ccatcttcgt gaacacgtaa tcagccaggg cgcgaacctc tttcgatgcc
40081 ttgcgcgcgg ccgttttctt gatcttccag accggcacac cggatgcgag ggcatcggcg
40141 atgctgctgc gcaggccaac ggtggccgga atcatcatct tggggtacgc ggccagcagc
40201 tcggcttggt ggcgcgcgtg gcgcggattc cgcgcatcga ccttgctggg caccatgcca

40261 aggaattgca gcttggcgtt cttctggcgc acgttcgcaa tggtcgtgac catcttcttg
40321 atgccctgga tgctgtacgc ctcaagctcg atgggggaca gcacatagtc ggccgcgaag
40381 agggcggccg ccaggccgac gccaagggtc ggggccgtgt cgatcaggca cacgtcgaag
40441 ccttggttcg ccagggcctt gatgttcgcc ccgaacagct cgcgggcgtc gtccagcgac
40501 agccgttcgg cgttcgccag taccgggttg gactcgatga gggcgaggcg cgcggcctgg
40561 ccgtcgccgg ctgcgggtgc ggtttcggtc cagccgccgg cagggacagc gccgaacagc
40621 ttgcttgcat gcaggccggt agcaaagtcc ttgagcgtgt aggacgcatt gccctggggg
40681 tccaggtcga tcacggcaac ccgcaagccg cgctcgaaaa agtcgaaggc aagatgcaca
40741 agggtcgaag tcttgccgac gccgcctttc tggttggccg tgaccaaagt tttcatcgtt
40801 tggtttcctg tttttcttg gcgtccgctt cccacttccg gacgatgtac gcctgatgtt
40861 ccggcagaac cgccgttacc cgcgcgtacc cctcgggcaa gttcttgtcc tcgaacgcgg
40921 cccacacgcg atgcaccgct tgcgacactg cgcccctggt cagtcccagc gacgttgcga
40981 acgtcgcctg tggcttccca tcgactaaga cgccccgcgc tatctcgatg gtctgctgcc
41041 ccacttccag cccctggatc gcctcctgga actggctttc ggtaagccgt ttcttcatgg
41101 ataacaccca taatttgctc cgcgccttgg ttgaacatag cggtgacagc cgccagcaca
41161 tgagagaagt ttagctaaac atttctcgca cgtcaacacc tttagccgct aaaactcgtc
41221 cttggcgtaa caaaacaaaa gcccggaaac cgggctttcg tctcttgccg cttatggctc
41281 tgcacccggc tccatcacca acaggtcgcg cacgcgcttc actcggttgc ggatcgacac
41341 tgccagccca acaaagccgg ttgccgccgc cgccaggatc gcgccgatga tgccggccac
41401 accggccatc gcccaccagg tcgccgcctt ccggttccat tcctgctggt actgcttcgc
41461 aatgctggac ctcggctcac cataggctga ccgctcgatg gcgtatgccg cttctcccct
41521 tggcgtaaaa cccagcgccg caggcggcat tgccatgctg cccgccgctt tcccgaccac
41581 gacgcgcgca ccaggcttgc ggtccagacc ttcggccacg gcgagctgcg caaggacata
41641 atcagccgcc gacttggctc cacgcgcctc gatcagctct tgcactcgcg cgaaatcctt
41701 ggcctccacg gccgccatga atcgcgcacg cggcgaaggc tccgcagggc cggcgtcgtg
41761 atcgccgccg agaatgccct tcaccaagtt cgacgacacg aaaatcatgc tgacggctat
41821 caccatcatg cagacggatc gcacgaaccc gctgaattca cccccgaaca cgagcacggc
41881 acccgcgacc actatgccaa gaatgcccaa ggtaaaaatt gccggccccg ccatgaagtc
41941 cgtgaatgcc ccgacggccg aagtgaaggg caggccgcca cccaggccgc cgccctcact
42001 gcccggcacc tggtcgctga atgtcgatgc cagcacctgc ggcacgtcaa tgcttccggg
42061 cgtcgcgctc gggctgatcg cccatcccgt tactgccccg atcccggcaa tggcaaggac
42121 tgccagcgcc gcgatgagga agcgggtgcc ccgcttcttc atcttcgcgc ctcgggcctc
42181 gaggccgcct acctgggcga aaacatcggt gtttgtggca ttcatacgga ctcctgttgg
42241 gccagctcgc gcacgggctg gcgggtcagc ttggcttgaa gatcgccacg cattgcggcg
42301 atctgcttct cggcatcctt gcgcttctgc acgccttcct gctggatgcg aataacgtcc
42361 tcgacggtct tgatgagcgt cgtctgaacc tgcttgagcg tgtccacgtc gatcaccagg
42421 cgttggttct ccttcgccgt ctcgacggac gtgcgatgca gcagggccgc attgcgcttc
42481 atcaggtcgt tggtggtgtc gtcgatggcc gtggccagtt cgacggcgtt cttctgctcg
42541 ttgaggctca aggccagcat gaattgccgc ttccacgccg gcacggtgat ttcgcggatg
42601 gtgtggaatt tatcgaccag catctggttg ttggcctgga tcatgcggat ggtcggcagg
42661 ctctgcatgg ccgaatgttg caaggcgatc aggtcgccga tgcgcttgtc caggttggca
42721 accatcgcat cgaggtcggc cagctcctgc acgcggcccg ggtcgttccc gacattgccg
42781 cgcagaccct cggcctgctc gcgcagctcg gcaaggcgga ccttgccggc cgcgatgtgg
42841 acgccaagaa ggcggtgttc ctcgcgcacg gctgcgaaca tttcgtcgag cgaggcattg
42901 cgctgcgcga tgccttgctg ggtggtctgc acttcgctga ccaggtgttc gatctgctcg
42961 cgggtcgtgt cgaagcgcgc catgaagccc gtcgaacgga cgcggaagcg gtcgatcagc
43021 gggccaatca ggggcaggcg ggaacggttg tcggacaaag ggccgacgtt cagggaacgg
43081 gccttggcga caacctgggt cagtttctcg cctgcttcgt ccaggtcgct gttgcgcacc
43141 tggtccagca ggctatcggc gtagcgggac gtgtgctcgg ccacgtcgcg gccgaactcg
43201 gcaacggtct gcggactgcc gacctcgatc cgctgcgcga ccgcatggac ttccggcacg
43261 tcgctttcct gcaagcccag ctcgcgcagg gttgccgggg tcatgtcgaa ggcgacgata
43321 ggggccttgg cgtcgtgcgt cgttttcagt gcgttcatag ggttctcccg ccgtgttatt
43381 ggttgatgcc ttccaggctc tgcgaaaggc tccgcatgag cgcctggtga gctttggccg
43441 cctcggcgac cattgccgga ttcatgttct tggtggtgat gagcgcgagg gtgtgctgac
43501 gccagacggg caccaggacg gatgccgttt cagagaagcg gtccagcatg tccacggcct
43561 gcgcccgcgt gagcttcatc tgagtgacgc tcatttcatg ggacgccatg agggttgcca
43621 ggttggcgag cttgcgcgcg aagcgttcgc gcggcttgtc gaactcgatc acgccggcct

43681 tggccgcgcc ggcctcgggg ttctcgtcca ggaactcgcg cccggcttga atgtaggctc
43741 tgagccggtc tacctcggcc tcatgcgtat tgagcatgtc atccaaggcg cgcaacgtgt
43801 cccgcacgcg ctgcgctacg ccctcggctt cgtccagcaa ctggtcgagc gtcttgcggg
43861 cgacctgata cctcacctgg cgttcaacct cacggccaag catcttctcg aaccaggtag
43921 gcttttccgc gatcttgcgg gggtccgcgt cggccagctt cgccacgatc tggctgattt
43981 tgtcggccag cgcggcaact gcgccgtgct ccatcagatt cgacagctcg ttgagggaat
44041 ccgccccgtc gatgccggcc ccgtactcgc caatcgtcgc cggcgacgcg aagagggcgg
44101 gcaaaacctc ccccttcaat cgcgccatgt tcacgctttg ttcttccatt cgatacaccc
44161 tcgcggtggg ttaattgctt ttcgatggaa gaagtttagc taaactttct atccctcgtc
44221 aacaccttta gccgctaaaa tttgggggaca ggtcatttac agaaagccag ctcactcctg
44281 gcgttgcccc ttgagcgccg ctaggcgcgc agcatccttc gcgctgagaa agaacgtcat
44341 cagcggcccg accgtcttgc ttgaaccgtc ggcaaagcaa acatccatcg aacagccttg
44401 cgtgtggggg tccacgcctt cgaccagttt ccaagggtcc atgccccagc cctcggcctc
44461 cggattgaac cagtacgccg atgcgtcgcc gtttaggtcg ctgtcggcgt agtccttgac
44521 cagcacggcc acccgcccgg tcgtcgggca cacgtagccc ggctgcttag gttcctgtct
44581 tggcattgct caaagctcct tgaaggggcc gctctacagc cccttgggct tgtagagcga
44641 cacgaaatag gtgagtgcgg tcagtaccgc gaaatgcacc aggaacgtcc agccggcatg
44701 aacgccaagg gtgttccagt ggtacagcat ccgcaggaac tgaaagaaaa cgtcgataga
44761 gatgatccac ttcgccaccg gccacaccag gacagtaacg acccatacaa agcggaccag
44821 ggcctggaca cccttggcaa aagtgaaccg gggcggggcc ttgctcgggg cctcaacgcg
44881 cggggcaggg gcctcggcct ccacttccac gcctgggaac ttgataatct tcgacattgc
44941 ttgaccctcc acggcgatgc gtgttcaatt cgtccagcgc tcgcgcgcct agaccgtgat
45001 gtgacagcat cgaggtcaag cgccccggag aaatccgggg cgtcatccct atgccccgtc
45061 caactcggga accggctttt ccctggtgct caacctggcc ggctcgaccc acttggtgac
45121 ttgctgccac tcgttaccac tgcgaacggc tacccgaatc tgcacacgtt tagccgcgat
45181 cttcgtcact atgccagcaa cacaaacgga atacccgtac ccgccgcgcg gggtgtgctg
45241 ccagttcacc ctatctccta cttgcatcat catcccctgg cgtcagtgac cggcccggaa
45301 tttcgccaag tcgatttcgt tgaaggtcca gcgctgtttc gcccgttcct ccaacctcga
45361 cgactcccgc atgacctcga tgcgcaggcg ctcgacctgg tgcatcagct catcagcgcg
45421 ccgctgcttc tcggcaatag cattccgctg cgcgaccagc tcctggtcta cgttcggcag
45481 ctcgtcgatc cacggcatga acttatcggt catcggattg gcctccggta attgacctgg
45541 gaatctaccc ggcctcaaaa caagaatagg gcataatgcc ctaacttgtc aagcaatttt
45601 agctaaacaa ttgaggggat tcagcgaggc gtcatgcttg aaaacacctt tcccctggcg
45661 tgcaatcagc ttgtcccggc ggcagcgcac tgccgcagcg cggccagcaa ggtgccttgc
45721 tcgatccgct gtcgttcctc cggcgtaatg caatcgccgc agataccgcc cagctcttcg
45781 cgcagggcgt ccgccccgat caggctgcgg ctaagggtcg aaatggactt accgcagcgg
45841 cggcaattgg tggtgacaat ctcgatcttg cggttgggca tggccctatc tccttgagag
45901 aggcccgacc gtagccgggc ctcgttccgt taccagcctg cgcgagcctt cgcgcgttcg
45961 acttcctggg tggaccagtg gccctttgct tcaccctcca gggtgcaacc ttgcaggtac
46021 gccttgtgcc gcagctcttc ggcctggcgg gtaagctctg ccgcctgctc catcagcgcc
46081 gcgatctggt cacgacgggc cagaaaatcc gtggcggccg cgccgggcag ctcatcgagc
46141 caagacatga tcttgctttt catcggggtt gatcctccgg ttgctgacct gggcgaagtg
46201 cccggccttg gattgctata ttagggcatt atgccctaga tagtcaagga aatttagcta
46261 aacaatttgc ggcgggcgca cgaaaaaacc cggcttgcgg gccgggctgc tggcagcata
46321 tcgcaacgat cagggttggc ggttttagcc gctaaagtcc tctcccttgg cgtaaagtcc
46381 tgcgggcgtc agccctggcc tttccagatc gccccaatcc ccgctagatc gcaaaggatc
46441 gcccaggcgg catagggggat cggcgaatcc tcgccaaccc aacgccgcac cgtgcggtcg
46501 cccttggcac ccaagcccaa gatgcgcgca gcctgtccgc cggtgaggcc ggccaagtgc
46561 aagacttccc ggatttctgc gccggtcggc tgcacccagc gttccgccgg gcgcaggcac
46621 tcaagccgga tattcacgtc gctcatgctg cttttctcct aatcgttatc aaatggcggc
46681 ccgcaattgg tcgtagccgt agcacgactc gatgcaacgc gggtcatatt cacaaacctt
46741 tctaccgttg cgattgatgc gatagcgcgg cggcttgtcg tcgtgcgccc acacctgtcg
46801 cgtttcggga ctgatcttcg taatgatgac gtgcttcccg atttccttgg caaaaaccgg
46861 gtgatgcacg ctcacaatct cgcaccgcag gccaggaaag aacgtcgccg gccacgccgg
46921 cgcgtcctcg tccttggccg gctccggctc cggcttggcc ggtgtaaccg gctccctgcg
46981 ctgcccggcc ggctcctgag ctttcgcacg cgcggccgcg agcttcgcct tggcgctggc
47041 cgcccatttg cgggcgatga atgcctggtg cgccggcagc acggcatcaa cccaggcgaa

47101 gcccggcggc agttcgtccg gcttccgcca gtcctcgacc acggcccgca cgcggcgcgg
47161 cggtggcgtg atacccgcga gccacgccgg cagtggctcg tcggcaaggc ggtcctggtc
47221 gcgggcgtcc agataccggc gatgcgagcg caacagggcg cgcaaaaccc cgtccgctac
47281 ctggtccacc gtcatgccgc cgcgcgcatg gtcgtaatgg gaccgatagc ccgtttcgga
47341 aataaaaggg gtctgacgct cagtggaacg aaaactcacg ttaagggatt ttggtcatga
47401 gattatcaaa aaggatcttc acctagatcc ttttaaatta aaaatgaagt tttaaatcaa
47461 tctaaagtat atatgagtaa acttggtctg acagttacca atgcttaatc agtgaggcac
47521 ctatctcagc gatctgtcta tttcgttcat ccatagttgc ctgactcccc gtcgtgtaga
47581 taactacgat acgggagggc ttaccatctg gccccagtgc tgcaatgata ccgcgagatc
47641 cacgctcacc ggctccagat ttatcagcaa taaaccagcc agccggaagg gccgagcgca
47701 gaagtggtcc tgcaacttta tccgcctcca tccagtctat taattgttgc cgggaagcta
47761 gagtaagtag ttcgccagtt aatagtttgc gcaacgttgt tgccattgct gcaggcatcg
47821 tggtgtcacg ctcgtcgttt ggtatggctt cattcagctc cggttcccaa cgatcaaggc
47881 gggttacatg atcccccatg ttgtgcaaaa aagcggttag ctccttcggt cctccgatcg
47941 ttgtcagaag taagttggcc gcagtgttat cactcatggt tatggcagca ctgcataatt
48001 ctcttactgt catgccatcc gtaagatgct tttctgtgac tggtgagtac tcaaccaagt
48061 cattctgaga atagtgtatg cggcgaccga gttgctcttg cccggcgtca acacgggata
48121 ataccgcacc acataTGTTG ATACAACCAT AAAATGATAA TTACACCCAT AAATTGATAA
48181 TTATCACACC CATAAATTGA TATTGCCTCT TCATGGTCTA AACTTCAGTA AGTTTACGAC
48241 ATTTTCCTCG AGGTCATTTC ccaattattg aaggccgcta acgcggcctt tttttgtttc
48301 tggtctgcct cctCCCTGCA GGCTTCAACA AACAGacaat ctggtctgtt tgtattatgg
48361 aaaatttttc tgtataatag attcaacaaa cagacaatct ggtctgtttg tattatcgac
48421 CAACCCACTC CCATGGTGTG ACGGGCGGTG TGTACAAGGC CCGGGAACGT gTTCACAAAA
48481 GTTATCAGGC ATGCACCTGG TAGCTAGTCT TTAAACCAAT AGATTGCATC GGTTTAAAAG
48541 GCAAGACCGT CAAATTGCGG GAAAGGGGTC AACAGCCGTT CAGTACCAAG TCTCAGGGGA
48601 AACTTTGAGA TGGCCTTGCA AAGGGTATGG TAATAAGCTG ACGGACATGG TCCTAACCAC
48661 GCAGCCAAGT CCTAAGTCAA CAGATCTTCT GTTGATATGG ATGCAGTTCA CAGACTAAAT
48721 GTCGGTCGGG GAAGATGTAT TCTTCTCATA AGATATAGTC GGACCTCTCC TTAATGGGAG
48781 ctagcggatg aagtgatgca acactggagc cgctgggaac taatttgtat gcgaaagtat
48841 attgattagt tttggagtac tcgatggtgt tcaatgcttt tcccGAGTGg ttatccggat
48901 cacatgaaac ggcatgactt tttcaagagt gccatgcccg aaggttatgt acaggaacgc
48961 actatatctt tcaaagatga cgggacctac aagacgcgtg ctgaagtcaa gtttgaaggt
49021 gatacccttg ttaatcgtat cgagttaaag ggtattgatt ttaaagaaga tggaaacatt
49081 cttggacaca aactcgagta caactttaac tcacacaatg tatacatcac ggcagacaaa
49141 caaaagaatg gaatcaaagc taacttcaaa attcgccaca acgttgaaga tggttccgtt
49201 caactagcag accattatca acaaaatact ccaattggcg atggccctgt ccttttacca
49261 gacaaccatt acctgtcgac acaatctgtc ctttcgaaag atcccaacga aaagcgtgac
49321 cacatggtcc ttcttgagtt tgtaactgct gctgggatta cacatggcat ggatgAGCTC
49381 TACAAATGGC ccaattattg aaggcctccc taacgggggg cctttttttG TTTCTGGTCT
49441 GCCGCTGCGC TTACTGCAGT AGTTTTGCTG AAATACTCGA TTCACAAAAA TATCAACTTA
49501 TGGTTGTTTT GTGAGATATC AATATATGGT TGTTTTGTGG TTAAGTTGCT GATTATAAAT
49561 AATTATTAAA TATCACTTTA TGGTTGCATC AACAacatag cagaacttta aaagtgctca
49621 tcattggaaa acgttcttcg gggcgaaaac tctcaaggat cttaccgctg ttgagatcca
49681 gctcgatgta acccactcgt gcacccaact tatcttcagc atcttttact ttcaccagcg
49741 tttctgggtg agcaaaaaca ggaaggcaaa atgccgcaaa aaagggaata agggcgacac
49801 ggaaatgttg aatactcata ctcttccttt ttcaatatta ttgaagcatt tatcagggtt
49861 attgtctcat gagcggatac atatttgaat gtatttagaa aaataaacaa ataggggttc
49921 cgcgcacatt tccccgaaaa gtgccacctg acgtctaaga aaccattatt atcatgacat
49981 taacctataa aaataagcgt atcacgaggc cctttcgtct tcaagaattt tataaaccgt
50041 ggagcgggca atactgagct gatgagcaat ttccgttgca ccagtgccct tctgatgaag
50101 cgtcagcacg acgttcctgt ccacggtacg cctgcggcca aatttgattc ctttcagctt
50161 tgcttcctgt cggccctcat tcgtgcgctc taggatcctc cggcgttcag cttgtgccac
50221 agccgacagg atggtgacca ccatttgccc catatcaccg tcggtactga tcccgtcgtc
50281 aataaaccga accgcgacac cctgagcatc aaactctttt atcagttgga tcatgtcggc
50341 ggtgtcgcgg ccaagacggt cgagcttctt caccagaatg acatcacctt cctccacctt
50401 catcctcagc aaatccagcc cttcccgatc tgttgaactg ccggatgcct tgtcggtaaa
50461 gatgcggtta gcttttaccc ctgcatcttt gagcgctctg atctgaatat cgagggactg

```
50521 ctggctggtt gagacccgcg cataaccaaa aattcgcata aaatgtacct taaatcgaat
50581 atcagacacg atgtgtctat tatgccaaaa tgacgattta atggacactc aaacgaagcc
50641 gttttactat gtctgataat ttataacatt tcggacggtt gcaaaaatgt tactaaatgc
50701 ccgtcaggca gggaggccga tatgcccgtt gactttctga ccactgagca gactgaaagc
50761 tatggcagat tcaccggtga accggatgag cttcagctgg cacgatattt tcatcttgat
50821 gaagcagaca aggaatttat cggaaaaagc agaggtgatc acaaccgtct gggcattgcc
50881 ctgcaaattg gatgtgtccg ttttctgggc accttcctca ccgatatgaa tcatattcct
50941 tccggcgtcc ggcattttac cgccagacag ctcgggattc gtgatatcac cgttcttgca
51001 gaatacggtc agagggaaaa tacccgccgt gagcatgcag cgctgatacg tcagcactat
51061 cagtatcgtg aatttgcctg gccctggaca tttcgcctta cccgtctttt atatacccgg
51121 agctggataa gcaacgaacg tcctggcctg cttttcgatc tggcgacagg gtggcttatg
51181 caacatcgta ttattctccc cggagccact acgctgaccc ggttgatttc agaggtaagg
51241 gaaaaggcga cgttgcgcct gtggaacaaa ctggcactga taccgtcagc cgaacaacgt
51301 tcacagctgg agatgctgct ggggccaact gattgcagcc gcctgtcttt actggaatca
51361 ctgaaaaaag gccctgtgac catcagtggt ccggcgttta atgaagcaat tgaacgctgg
51421 aaaactctga acgattttgg cttgcatgct gacaacctga gtacactccc ggctgtgcgc
51481 ctgaaaaatc tcgcacgtta tgctggtatg acttcggtgt tcaatattgc caggatgtca
51541 ccgcagaaaa ggatggcggt tctggttgcc tttgtccttg catgggaaac gctggcgttg
51601 gatgatgcac tggacgttct ggacgccatg ctggccgtta tcatccgtga cgccagaaag
51661 attgggcaga aaaaacggct ccgctcgctg aaggatctgg ataaatctgc attggcgctc
51721 gccagcgcat gttcgtacct gctgaaagaa gaaacaccgg acgaatcgat tcgtgctgag
51781 gtgttcagct acatccccag gcaaaagctg gctgaaatca tcacgcttgt ccgtgaaatt
51841 gcccggccct cagacgataa ttttcatgaa gaaatggtgg agcagtacgg gcgcgttcgt
51901 cgtttcctgc cccatctgct gaataccgtt aaattttcat ccgcacctgc cggggttacc
51961 actctgaatg cctgtgacta cctcagccgg gagttcagct cacggcggca gtttttttgac
52021 gacgcaccaa cggaaattat cagtcggtca tggaaacggc tggtgattaa caaggaaaaa
52081 catatcaccc gcaggggata cacgctctgc tttctcagta aactgcagga tagtctgagg
52141 cggagggatg tctacgttac cggcagtaac cggtggggag atccccgagc aagattacta
52201 cagggtgctg actggcaggc aaaccggatt aaggtttatc gttctctggg acacccgaca
52261 gacccgcagg aagcaataaa atctctgggt catcagcttg atagtcgtta cagacaggtt
52321 gctgcacgtc ttggcgaaaa tgaggctgtc gaactcgatg tttctggccc gaagccccgg
52381 ttgacaattt ctcccctcgc cagtcttgat gagccggaca gtctgaaacg actgagcaaa
52441 atgatcagtg atctgctccc tccggtggat ttaacggagt tgctgctcga aattaacgcc
52501 cataccggat ttgctgatga gtttttccat gctagtgaag ccagtgccag agttgatgat
52561 ctgcccgtca gcatcagcgc cgtgctgatg gctgaagcct gcaatatcgg tctggaacca
52621 ctgatcagat caaatgttcc tgcactgacc cgacaccggc tgaactggac aaaagcgaac
52681 tatctgcggg ctgaaactat caccagcgct aatgccagac tggttgattt tcaggcaacg
52741 ctgccactgg cacagatatg gggtggcgga gaagtggcat ctgcagatgg aatgcgcttt
52801 gttacgccag tcagaacaat caatgccgga ccgaaccgca aatactttgg taataacaga
52861 gggatcacct ggtacaactt tgtgtccgat cagtattccg gctttcatgg catcgttata
52921 ccggggacgc tgagggactc tatctttgtg ctggaaggcc ttctggaaca ggagaccggg
52981 ctgaatccaa ccgaaattat gaccgataca gcaggtacca gcgaacttgt ctttggcctt
53041 ttctggctgc tgggatacca gttttctcca cgcctggctg atgccggtgc ttcggttttc
53101 tggcgaatgg gccatgatgc caactatggc gtgctgaatg atattgccag agggcaatca
53161 gatccccgaa aaatagtcct tcagtgggac gaaatgatcc ggaccgctgg ctccctgaaa
53221 ctgggcaaag tacaggcttc agtgctggtc cgttcattgc tgaaaagtga acgtccttcc
53281 ggactgactc aggcaatcat tgaagtgggg cgcatcaaca aaacgctgta tctgcttaat
53341 tatattgatg atgaagatta ccgccggcgc attctgaccc agcttaatcg gggagaaagc
53401 cgccatgccg ttgccagagc catctgtcac ggtcaaaaag gtgagataag aaaacgatat
53461 accgacggtc aggaagatca actgggcgca ctggggctgg tcactaacgc cgtcgtgtta
53521 tggaacacta tgtatatgca ggcagccctg gatcatctcc gggcgcaagg tgaaacactg
53581 aatgatgaag atatcgcacg cctctccccg ctttgccacg gacatatcaa tatgctcggc
53641 cattattcct tcacgctggc agaactggtg accaaaggac atctgagacc attaaaagag
53701 gcgtcagagg tagaaaacgt tgcttaacgt gagttttcgt tccactgagc gtcagacccc
53761 taaaagggcc gtccaggtcc acggcgtgga actggaaatg caccgtggtt aggccgccgt
53821 agccgctttc gacctccacc catacgcgca tcccgtccac ggaagccagg aagtcgcctt
53881 cctggcccca catcgggacg acgcccggcg tcgcgcggca atggcgctct atgaccctct
```

```
53941 cggccgcatc ctggtcgccg gcgcaaccga agaacgtgcc cccgttgagc ttccacaccg
54001 ccgccgcgta acgctcgccg gccacgtcgg ccaccaaatc ggcacgattg agcactgcgt
54061 catgcaccgc cgctaccgcg tccgccgcaa cggccagcag gccggcgcga tcctcgggca
54121 gcgtcgccgc cagcgctgca atgcgcgcgt ccttgtcctc ggtctgcggc ttgcttttgc
54181 gcgccttcgt ccgccggcga gccgccgggg tggtcgcctc gggcggcgtc gggttggcta
54241 ccaggtaggc cgcgatgcaa tccggtttcc ggtagtcatc gcagaacttc aagccgtcca
54301 ggacgcagca ggtcggcaac gggtactgca tccagtattc gggttccggg aacggatgca
54361 gcaactcgcc gcagtggtgg catcgctctg tctcgcggtc ctgcaccgaa ccgctgaaca
54421 agcggccggc aacgatggca tcgttgcggc gcttgatgtt gtcgtcttgc atgctcgcct
54481 ccaacggccc ccagcctttc ggccgggggc ctgcccctta gtcgatggcc cggtagatcg
54541 ccgccgcctc ggcgtgctcg ctcgcaaacg ccgccaggtg gtggtagcgg tcgatcaggg
54601 cgtccgcgtc ggccgtgccg gccagctccg cgcagagctg gttcagggcg aagagcgtgg
54661 cgacgatgcc ggcggcgtcg gccgacagct cgcccgcgaa gccgttgcca tcgacctcga
54721 tgcgcaagcg gccgtcatgc tccggggcca tgtagaagcc accgttcgag agggtgtaga
54781 agtgccagaa accgccgtgg taggcagcgc aaaggcggcc catccagccg aataccaggg
54841 cctcgccgcg cagcatgcgc agcatcgagg ggccgaagta cgtcggcaga aaccgcaggc
54901 gatccgcctc ggcgacgcgg gtggcaacga taggggaggg gatttgaaca tcagtcatcg
54961 gactttctcc ggtagttgac ctgggcggaa tgcccggcct caagaccaat attagggcat
55021 tatgccctag cctgtcaagc aaatttagct aaactatgcc gccggccgta cccgattctg
55081 cggttacagc tcggtcctgg tcccgaccgg gctttgccag gcgggcggat ggaacgccaa
55141 gggggagcgc ggccgttgaa tggtgatcgc catcacgttt cattgacact tgaggggcgt
55201 ttagagcgag ccaggaaagc cgacccctc cttggagtaa aaacccttgc ggcgttgcag
55261 ccggcacgga tcttccgatc gggcgcggtg gtggccgcgt ctgtgaccta aaaagggggg
55321 agtccagagg ggcgcagccc ctttgggcat agcgcagcgt aatcggagac gtaattgagc
55381 atttccaggc gcttgcgcct ggtcaacgaa agagtcagcg ccgtaggcgc tgccattttt
55441 ggggtgaggc cgttcgcggc cgagggcgc agcccctggg gggatgggag gcccgcgtta
55501 gcgggccggg agggttcgag aagggggggc accccccttc ggcgtgcgcg gtcacgcgca
55561 cagggcgcag ccctggttaa aaacaaggtt tataaatatt ggtttaaaag caggttaaaa
55621 gacaggttag cggtggccga aaaacgggcg gaaacccttg caaatgctgg attttctgcc
55681 tgtggacagc ccctcaaatg tcaataggtg cgcccctcat ctgtcagcac tctgcccctc
55741 aagtgtcaag gatcgcgccc ctcatctgtc agtagtcgcg cccctcaagt gtcaataccg
55801 cagggcactt atccccaggc ttgtccacat catctgtggg aaactcgcgt aaaatcaggc
55861 gttttcgccg atttgcgagg ctggccagct ccacgtcgcc ggccgaaatc gagcctgccc
55921 ctcatctgtc aacgccgcgc cgggtgagtc ggcccctcaa gtgtcaacgt ccgcccctca
55981 tctgtcagtg agggccaagt tttccgcgag gtatccacaa cgccggcggc cgcggtgtct
56041 cgcacacggc ttcgacggcg tttctggcgc gtttgcaggg ccatagacgg ccgccagccc
56101 agcggcgagg gcaaccagcc cggtgagcgt cggaaaggcg ctggaagccc cgtagcgacg
56161 cggagagggg cgagacaagc caagggcgca ggctcgatgc gcagcacgac atagccggtt
56221 ctcgcaagga cgagaatttc cctgcggtgc ccctcaagtg tcaatgaaag tttccaacgc
56281 gagccattcg cgagagcctt gagtccacgc tagatctatc tcatctgcgc aaggcagaac
56341 gtgaagacgg ccgccctgga cctcgcccgc gagcgccagg cgcacgaggc cggcgcgcgg
56401 acccgcgcca cggcccacga gcggacgccg cagcaggagc gccagaaggc cgccagagag
56461 gccgagcgcg gccgtgaggc ttggacgcta gggcagggca tgaaaaagcc cgtagcgggc
56521 tgctacgggc gtctgacgcg gtggaaaggg ggaggggatg ttgtctacat ggctctgctg
56581 tagtgagtgg gttgcgctcc ggcagcggtc ctgatcaatc gtcacccttt ctcggtcctt
56641 caacgttcct gacaacgagc ctccttttcg ccaatccatc gacaatcacc gcgagtccct
56701 gctcgaacgc tgcgtccgga ccggcttcgt cgaaggcgtc tatcgcggcc cgcaacagcg
56761 gcgagagcgg agcctgttca acggtgccgc cgcgctcgcc ggcatcgctg tcgccggcct
56821 gctcctcaag cacggcccca acagtgaagt agctgattgt catcagcgca ttgacggcgt
56881 ccccggccga aaaacccgcc tcgcagagga agcgaagctg cgcgtcggcc gtttccatct
56941 gcggtgcgcc cggtcgcgtg ccggcatgga tgcgcgcgcc atcgcggtag gcgagcagcg
57001 cctgcctgaa gctgcgggca ttcccgatca gaaatgagcg ccagtcgtcg tcggctctcg
57061 gcaccgaatg cgtatgattc tccgccagca tggcttcggc cagtgcgtcg agcagcgccc
57121 gcttgttcct gaagtgccag taaagcgccg gctgctgaac cccaaccgt tccgccagtt
57181 tgcgtgtcgt cagaccgtct acgccgacct cgttcaacag gtccagggcg gcacggatca
57241 ctgtattcgg ctgcaacttt gtcatgcttg acactttatc actgataaac ataatatgtc
57301 caccaactta tcagtgataa agaatccgcg cgttcaatcg gaccagcgga ggctggtccg
```

57361 gaggccagac gtgaaaccca acatacccct gatcgtaatt ctgagcactg tcgcgctcga
57421 cgctgtcggc atcggcctga ttatgccggt gctgccgggc ctcctgcgcg atctggttca
57481 ctcgaacgac gtcaccgccc actatggcat tctgctggcg ctgtatgcgt tggtgcaatt
57541 tgcctgcgca cctgtgctgg gcgcgctgtc ggatcgtttc gggcggcggc caatcttgct
57601 cgtctcgctg gccggcgcca ctgtcgacta cgccatcatg gcgacagcgc ctttcctttg
57661 ggttctctat atcgggcgga tcgtggccgg catcaccggg gcgactgggg cggtagccgg
57721 cgcttatatt gccgatatca ctgatggcga tgagcgcgcg cggcacttcg gcttcatgag
57781 cgcctgtttc gggttcggga tggtcgcggg acctgtgctc ggtgggctga tgggcggttt
57841 ctccccccac gctccgttct tcgccgcggc agccttgaac ggcctcaatt tcctgacggg
57901 ctgtttcctt ttgccggagt cgcacaaagg cgaacgccgg ccgttacgcc gggaggctct
57961 caacccgctc gcttcgttcc ggtgggcccg gggcatgacc gtcgtcgccg ccctgatggc
58021 ggtcttcttc atcatgcaac ttgtcggaca ggtgccggcc gcgctttggg tcattttcgg
58081 cgaggatcgc tttcactggg acgcgaccac gatcggcatt tcgcttgccg catttggcat
58141 tctgcattca ctcgcccagg caatgatcac cggccctgta gccgcccggc tcggcgaaag
58201 gcgggcactc atgctcggaa tgattgccga cggcacaggc tacatcctgc ttgccttcgc
58261 gacacgggga tggatggcgt tcccgatcat ggtcctgctt gcttcgggtg gcatcggaat
58321 gccggcgctg caagcaatgt tgtccaggca ggtggatgag gaacgtcagg ggcagctgca
58381 aggctcactg gcggcgctca ccagcctgac ctcgatcgtc ggacccctcc tcttcacggc
58441 gatctatgcg gcttctataa caacgtggaa cgggtgggca tggattgcag gcgctgccct
58501 ctacttgctc tgcctgccgg cgctgcgtcg cgggctttgg agcggcgcag ggcaacgagc
58561 cgatcgctga tcgtggaaac gataggccta tgccatgcgg gtcaaggcga cttccggcaa
58621 gctatacgcg ccctaggagt gcggttggaa cgttggccca gccagatact cccgatcacg
58681 agcaggacgc cgatgatttg aagcgcactc agcgtctgat ccaagaacaa ccatcctagc
58741 aacacggcgg tccccgggct gagaaagccc agtaaggaaa caactgtagg ttcgagtcgc
58801 gagatccccc ggaaccaaag gaagtaggtt aaacccgctc cgatcaggcc gagccacgcc
58861 aggccgagaa cattggttcc tgtaggcatc gggattggcg gatcaaacac taaagctact
58921 ggaacgagca gaagtcctcc ggccgccagt tgccaggcgg taaaggtgag cagaggcacg
58981 ggaggttgcc acttgcgggt cagcacggtt ccgaacgcca tggaaaccgc cccgccagg
59041 cccgctgcga cgccgacagg atctagcgct gcgtttggtg tcaacaccaa cagcgccacg
59101 cccgcagttc cgcaaatagc cccaggacc gccatcaatc gtatcgggct acctagcaga
59161 gcggcagaga tgaacacgac catcagcggc tgcacagcgc ctaccgtcgc cgcgaccccg
59221 cccggcaggc ggtagaccga aataaacaac aagctccaga atagcgaaat attaagtgcg
59281 ccgaggatga agatgcgcat ccaccagatt cccgttggaa tctgtcggac gatcatcacg
59341 agcaataaac ccgccggcaa cgcccgcagc agcataccgg cgacccctcg gcctcgctgt
59401 tcgggctcca cgaaaacgcc ggacagatgc gccttgtgag cgtccttggg gccgtcctcc
59461 tgtttgaaga ccgacagccc aatgatctcg ccgtcgatgt aggcgccgaa tgccacggca
59521 tctcgcaacc gttcagcgaa cgcctccatg ggctttttct cctcgtgctc gtaaacggac
59581 ccgaacatct ctggagcttt cttcagggcc gacaatcgga tctcgcggaa atcctgcacg
59641 tcggccgctc caagccgtcg aatctgagcc ttaatcacaa ttgtcaattt taatcctctg
59701 tttatcggca gttcgtagag cgcgccgtgc gtcccgagcg atactgagcg aagcaagtgc
59761 gtcgagcagt gcccgcttgt tcctgaaatg ccagtaaagc gctggctgct gaacccccag
59821 ccggaactga ccccacaagg ccctagcgtt tgcaatgcac caggtcatca ttgacccagg
59881 cgtgttccac caggccgctg cctcgcaact cttcgcaggc ttcgccgacc tgctcgcgcc
59941 acttcttcac gcgggtggaa tccgatccgc acatgaggcg gaaggtttcc agcttgagcg
60001 ggtacggctc ccggtgcgag ctgaaatagt cgaacatccg tcgggccgtc ggcgacagct
60061 tgcggtactt ctcccatatg aatttcgtgt agtggtcgcc agcaaacagc acgacgattt
60121 cctcgtcgat caggacctgg caacgggacg ttttcttgcc acggtccagg acgcggaagc
60181 ggtgcagcag cgacaccgat tccaggtgcc caacgcggtc ggacgtgaag cccatcgccg
60241 tcgcctgtag gcgcgacagg cattcctcgg ccttcgtgta ataccggcca ttgatcgacc
60301 agcccaggtc ctggcaaagc tcgtagaacg tgaaggtgat cggctcgccg ataggggtgc
60361 gcttcgcgta ctccaacacc tgctgccaca ccagttcgtc atcgtcggcc cgcagctcga
60421 cgccggtgta ggtgatcttc acgtccttgt tgacgtggaa aatgaccttg ttttgcagcg
60481 cctcgcgcgg gattttcttg ttgcgcgtgg tgaacagggc agagcggcc gtgtcgtttg
60541 gcatcgctcg catcgtgtcc ggccacggcg caatatcgaa caaggaaagc tgcatttcct
60601 tgatctgctg cttcgtgtgt ttcagcaacg cggcctgctt ggcctcgctg acctgttttg
60661 ccaggtcctc gccggcggtt tttcgcttct tggtcgtcat agttcctcgc gtgtcgatgg
60721 tcatcgactt cgccaaacct gccgcctcct gttcgagacg acgcgaacgc tccacggcgg

```
    60781 ccgatggcgc gggcagggca gggggagcca gttgcacgct gtcgcgctcg atcttggccg
    60841 tagcttgctg gaccatcgag ccgacggact ggaaggtttc gcggggcgca cgcatgacgg
    60901 tgcggcttgc gatggtttcg gcatcctcgg cggaaaaccc cgcgtcgatc agttcttgcc
    60961 tgtatgcctt ccggtcaaac gtccgattca ttcaccctcc ttgcgggatt gccccgactc
    61021 acgccggggc aatgtgccct tattcctgat ttgacccgcc tggtgccttg gtgtccagat
    61081 aatccacctt atcggcaatg aagtcggtcc cgtagaccgt ctggccgtcc ttctcgtact
    61141 tggtattccg aatcttgccc tgcacgaata ccagcgaccc cttgcccaaa tacttgccgt
    61201 gggcctcggc ctgagagcca aaacacttga tgcggaagaa gtcggtgcgc tcctgcttgt
    61261 cgccggcatc gttgcgccac tcttcattaa ccgctatatc gaaaattgct tgcggcttgt
    61321 tagaattgcc atgacgtacc tcggtgtcac gggtaagatt accgataaac tggaactgat
    61381 tatggctcat atcgaaagtc tccttgagaa aggagactct agtttagcta aacattggtt
    61441 ccgctgtcaa gaactttagc ggctaaaatt ttgcgggccg cgaccaaagg tgcgaggggc
    61501 ggcttccgct gtgtacaacc agatattttt caccaacatc cttcgtctgc tcgatgagcg
//
```

**pMAS054**

```
LOCUS       pMAS054                 5904 bp ds-DNA     circular     02-APR-
2026
DEFINITION  .
Aignments
FEATURES             Location/Qualifiers
     misc_feature    120..140
                     /label="Probe binding site"
                     /ApEinfo_revcolor="#ff9ccd"
                     /ApEinfo_fwdcolor="#ff9ccd"
     primer          complement(120..141)
                     /label="KRG089"
                     /note="sequence: GCGGTCTCAgcgcCGGGAAAAGCATTGAACACCAT"
                     /ApEinfo_revcolor="#b7e6d7"
                     /ApEinfo_fwdcolor="#b7e6d7"
     primer          complement(120..141)
                     /label="oMJD105"
                     /note="sequence: GCGGTCTCAcactcGGGAAAAGCATTGAACACCAT"
                     /ApEinfo_revcolor="#b7e6d7"
                     /ApEinfo_fwdcolor="#b7e6d7"
     primer          complement(120..142)
                     /label="MAS75B"
                     /note="sequence: TCGGGAAAAGCATTGAACACCAT"
                     /ApEinfo_revcolor="#b7e6d7"
                     /ApEinfo_fwdcolor="#b7e6d7"
     misc_feature    120..642
                     /label="Barcode (sfGFP_2)"
                     /ApEinfo_revcolor="#84b0dc"
                     /ApEinfo_fwdcolor="#84b0dc"
     misc_feature    141..145
                     /label="Barcode 2"
                     /ApEinfo_revcolor="#b1ff67"
                     /ApEinfo_fwdcolor="#b1ff67"
     primer          143..166
                     /label="MAS74B"
                     /note="sequence: GTGGTTATCCGGATCACATGAAAC"
                     /ApEinfo_revcolor="#b7e6d7"
                     /ApEinfo_fwdcolor="#b7e6d7"
     primer          144..165
                     /label="KRG090"
                     /note="sequence: GCGGTCTCAgcgcGGTTATCCGGATCACATGAAA"
                     /ApEinfo_revcolor="#b4abac"
                     /ApEinfo_fwdcolor="#b4abac"
     primer          146..165
                     /label="oMJD104"
                     /note="sequence: GCGGTCTCAGAGTGGTTATCCGGATCACATGAAA"
                     /ApEinfo_revcolor="#b4abac"
                     /ApEinfo_fwdcolor="#b4abac"
     misc_feature    555..560
                     /label="BamHI-BglII Scar"
                     /ApEinfo_revcolor="#ac1dff"
                     /ApEinfo_fwdcolor="#ac1dff"
     terminator      647..699
                     /label="tVoigtS4 L3S3P21 51C>G"
                     /ApEinfo_revcolor="#f58a5e"
```

```
                    /ApEinfo_fwdcolor="#f58a5e"
     misc_feature    675..699
                    /label="homology tVoigtS5"
                    /ApEinfo_revcolor="#f58a5e"
                    /ApEinfo_fwdcolor="#f58a5e"
     primer          675..699
                    /label="AB17 ConL F seq"
                    /note="sequence: ggcctttttttgtttctggtctgcc"
                    /ApEinfo_revcolor="#75c6a9"
                    /ApEinfo_fwdcolor="#75c6a9"
     primer          complement(686..703)
                    /label="MAS65B"
                    /note="sequence: CAGCggCagaccagaaac"
                    /ApEinfo_revcolor="#b1ff67"
                    /ApEinfo_fwdcolor="#b1ff67"
     primer          689..731
                    /label="MAS62B"
                    /note="sequence:
tctggtctGccGCTGagttcaccgacaaacaacagataaaacg"
                    /ApEinfo_revcolor="#ffef86"
                    /ApEinfo_fwdcolor="#ffef86"
     misc_feature    complement(704..1071)
                    /label="TrrnB"
                    /ApEinfo_revcolor="#85dae9"
                    /ApEinfo_fwdcolor="#85dae9"
     misc_feature    704..1448
                    /label="from pMAS049"
                    /ApEinfo_revcolor="#75c6a9"
                    /ApEinfo_fwdcolor="#75c6a9"
     primer          complement(815..840)
                    /label="JEC20A | JC.A20"
                    /note="sequence: gaagtgaaacgccgtagcgccgatgg"
                    /ApEinfo_revcolor="#9eafd2"
                    /ApEinfo_fwdcolor="#9eafd2"
     misc_feature    963..980
                    /label="JEC71A"
                    /ApEinfo_revcolor="#b7e6d7"
                    /ApEinfo_fwdcolor="#b7e6d7"
     primer          963..980
                    /label="JEC71A"
                    /note="sequence: caaaacagccaagctgga"
                    /ApEinfo_revcolor="#b7e6d7"
                    /ApEinfo_fwdcolor="#b7e6d7"
     misc_feature    1030..1035
                    /label="BamHI-BglII Scar"
                    /ApEinfo_revcolor="#ac1dff"
                    /ApEinfo_fwdcolor="#ac1dff"
     primer          complement(1052..1077)
                    /label="AOS49A"
                    /note="sequence: ggatctgaagcttgggcccgaacaaa"
                    /ApEinfo_revcolor="#c7b0e3"
                    /ApEinfo_fwdcolor="#c7b0e3"
     misc_feature    complement(1060..1081)
                    /label="LCG38F"
                    /ApEinfo_revcolor="#85dae9"
                    /ApEinfo_fwdcolor="#85dae9"
     primer          complement(1060..1081)
```

```
                     /label="poscon temp fwd"
                     /note="sequence: TATGggatctgaagcttgggcc"
                     /ApEinfo_revcolor="#d59687"
                     /ApEinfo_fwdcolor="#d59687"
     primer          complement(1062..1096)
                     /label="MAS60B"
                     /note="sequence: ttctggggaatataaTATGggatctgaagcttggg"
                     /ApEinfo_revcolor="#c7b0e3"
                     /ApEinfo_fwdcolor="#c7b0e3"
     primer          1067..1109
                     /label="MAS59B"
                     /note="sequence:
gcttcagatccCATAttatattccccagaacatcaggttaatg"
                     /ApEinfo_revcolor="#f58a5e"
                     /ApEinfo_fwdcolor="#f58a5e"
     CDS             complement(1069..1081)
                     /label="Translation 1081-1069"
                     /translation="YGI*"
     misc_feature    1072..1077
                     /label="BamHI-BglII Scar"
                     /ApEinfo_revcolor="#ac1dff"
                     /ApEinfo_fwdcolor="#ac1dff"
     misc_feature    1078..1081
                     /label="LCG54G"
                     /ApEinfo_revcolor="#9eafd2"
                     /ApEinfo_fwdcolor="#9eafd2"
     CDS             complement(1082..1387)
                     /label="ccdB CDS (type II toxin-antitoxin system toxin
CcdB)"
                     /ApEinfo_revcolor="#b4abac"
                     /ApEinfo_fwdcolor="#b4abac"
     gene            complement(1082..1387)
                     /label="ccdB gene"
                     /ApEinfo_revcolor="#faac61"
                     /ApEinfo_fwdcolor="#faac61"
     primer          complement(1357..1403)
                     /label="MAS58B"
                     /note="sequence:
ctaggaggaatgACCTatgcagtttaaggtttacacctataaaagag"
                     /ApEinfo_revcolor="#f58a5e"
                     /ApEinfo_fwdcolor="#f58a5e"
     primer          1373..1414
                     /label="MAS61B"
                     /note="sequence:
aaccttaaactgcatAGGTcattcctcctagatccaaaatac"
                     /ApEinfo_revcolor="#c7b0e3"
                     /ApEinfo_fwdcolor="#c7b0e3"
     misc_feature    complement(1388..1388)
                     /label="Catalytic U"
                     /ApEinfo_revcolor="#c7b0e3"
                     /ApEinfo_fwdcolor="#c7b0e3"
     misc_feature    complement(1388..1393)
                     /label="IGS"
                     /ApEinfo_revcolor="#c7b0e3"
                     /ApEinfo_fwdcolor="#c7b0e3"
     primer          1388..1415
                     /label="MAS007A"
```

```
                     /note="sequence: AGGTcattcctcctagatccaaaatacg"
                     /ApEinfo_revcolor="#d59687"
                     /ApEinfo_fwdcolor="#d59687"
     misc_feature    complement(1394..1401)
                     /label="RBS-Strong"
                     /ApEinfo_revcolor="#9eafd2"
                     /ApEinfo_fwdcolor="#9eafd2"
     misc_feature    1402..1407
                     /label="BamHI-BglII Scar"
                     /ApEinfo_revcolor="#ac1dff"
                     /ApEinfo_fwdcolor="#ac1dff"
     misc_feature    complement(1408..1448)
                     /label="HP14 Stability Hairpin"
                     /ApEinfo_revcolor="#fa1839"
                     /ApEinfo_fwdcolor="#fa1839"
     primer          complement(1441..1465)
                     /label="MAS73B"
                     /note="sequence: ctaggtacaatgctagcacgtcgac"
                     /ApEinfo_revcolor="#75c6a9"
                     /ApEinfo_fwdcolor="#75c6a9"
     misc_feature    complement(1449..1483)
                     /label="J23114"
                     /ApEinfo_revcolor="#85dae9"
                     /ApEinfo_fwdcolor="#85dae9"
     primer          1466..1493
                     /label="MAS72B"
                     /note="sequence: gactgagctagccataaaCGTTtGAGAC"
                     /ApEinfo_revcolor="#75c6a9"
                     /ApEinfo_fwdcolor="#75c6a9"
     primer          1484..1506
                     /label="MAS64B"
                     /note="sequence: CGTTtGAGACGtactagtagcgg"
                     /ApEinfo_revcolor="#b1ff67"
                     /ApEinfo_fwdcolor="#b1ff67"
     misc_feature    complement(1484..1644)
                     /label="ConR2"
                     /ApEinfo_revcolor="#c6c9d1"
                     /ApEinfo_fwdcolor="#c6c9d1"
     misc_feature    complement(1489..1494)
                     /label="Esp3I"
                     /ApEinfo_revcolor="#f58a5e"
                     /ApEinfo_fwdcolor="#f58a5e"
     misc_feature    complement(1495..1515)
                     /label="BioBrick Suffix BBa_G00001"
                     /ApEinfo_revcolor="#d59687"
                     /ApEinfo_fwdcolor="#d59687"
     misc_feature    1496..1501
                     /label="SpeI"
                     /ApEinfo_revcolor="#d59687"
                     /ApEinfo_fwdcolor="#d59687"
     misc_feature    1503..1510
                     /label="NotI"
                     /ApEinfo_revcolor="#b7e6d7"
                     /ApEinfo_fwdcolor="#b7e6d7"
     misc_feature    1510..1515
                     /label="PstI"
                     /ApEinfo_revcolor="#d59687"
```

```
                     /ApEinfo_fwdcolor="#d59687"
     misc_feature    1516..1530
                     /label="6-frame STOP b"
                     /ApEinfo_revcolor="#c6c9d1"
                     /ApEinfo_fwdcolor="#c6c9d1"
     primer          complement(1531..1555)
                     /label="AB18 ConR R seq"
                     /note="sequence: ctcttttctggaatttggtaccgag"
                     /ApEinfo_revcolor="#85dae9"
                     /ApEinfo_fwdcolor="#85dae9"
     terminator      1531..1591
                     /label="tVoigtS1, L3S2P21"
                     /ApEinfo_revcolor="#c6c9d1"
                     /ApEinfo_fwdcolor="#c6c9d1"
     terminator      complement(1592..1644)
                     /label="tVoigtS19, L3S1P00"
                     /ApEinfo_revcolor="#c6c9d1"
                     /ApEinfo_fwdcolor="#c6c9d1"
     gene            complement(1649..2603)
                     /label="KanR.II, kanamycin resistance"
                     /ApEinfo_revcolor="#f6989d"
                     /ApEinfo_fwdcolor="#f6989d"
     CDS             complement(1650..2447)
                     /label="aph(3')-II / nptII / neo, aminoglycoside 3'-
phosphotransferase type II (KanR)"
                     /ApEinfo_revcolor="#f6989d"
                     /ApEinfo_fwdcolor="#f6989d"
     misc_feature    complement(2448..2603)
                     /label="aphIIp, KanR promoter/UTR"
                     /ApEinfo_revcolor="#f6989d"
                     /ApEinfo_fwdcolor="#f6989d"
     misc_feature    complement(2551..2559)
                     /label="aphIIp -10"
                     /ApEinfo_revcolor="#f6989d"
                     /ApEinfo_fwdcolor="#f6989d"
     misc_feature    complement(2574..2579)
                     /label="aphIIp -35"
                     /ApEinfo_revcolor="#f6989d"
                     /ApEinfo_fwdcolor="#f6989d"
     terminator      2608..2662
                     /label="tVoigtN17, tonBt (bidir), BBa_B0054"
                     /ApEinfo_revcolor="#f58a5e"
                     /ApEinfo_fwdcolor="#f58a5e"
     CDS             complement(2667..3332)
                     /label="rep"
                     /ApEinfo_revcolor="#c6c9d1"
                     /ApEinfo_fwdcolor="#c6c9d1"
     misc_feature    complement(2667..4190)
                     /label="pBBR1 rep region"
                     /ApEinfo_revcolor="#c6c9d1"
                     /ApEinfo_fwdcolor="#c6c9d1"
     misc_feature    2667..5328
                     /label="pBBR1-mob origin"
                     /ApEinfo_revcolor="#c6c9d1"
                     /ApEinfo_fwdcolor="#c6c9d1"
     misc_feature    3200..3208
                     /label="unstable repeat. 1e-5 rate"
```

```
                     /ApEinfo_revcolor="#b4abac"
                     /ApEinfo_fwdcolor="#b4abac"
     promoter        complement(3408..3442)
                     /label="repp"
                     /ApEinfo_revcolor="#c6c9d1"
                     /ApEinfo_fwdcolor="#c6c9d1"
     misc_feature    complement(3414..3422)
                     /label=-10
                     /ApEinfo_revcolor="#c6c9d1"
                     /ApEinfo_fwdcolor="#c6c9d1"
     misc_feature    complement(3437..3442)
                     /label=-35
                     /ApEinfo_revcolor="#c6c9d1"
                     /ApEinfo_fwdcolor="#c6c9d1"
     misc_feature    complement(3519..3531)
                     /label="IHF binding site"
                     /ApEinfo_revcolor="#c6c9d1"
                     /ApEinfo_fwdcolor="#c6c9d1"
     primer          4190..4206
                     /label="oPK133 pcr81 f1L"
                     /note="sequence: gtCGTCTCactcgACGGccgcagccgccgtagg"
                     /ApEinfo_revcolor="#ff9ccd"
                     /ApEinfo_fwdcolor="#ff9ccd"
     misc_feature    4191..5328
                     /label="pBBR1 mob region"
                     /ApEinfo_revcolor="#d6b295"
                     /ApEinfo_fwdcolor="#d6b295"
     misc_feature    4208..4259
                     /label="oriT region"
                     /ApEinfo_revcolor="#d6b295"
                     /ApEinfo_fwdcolor="#d6b295"
     misc_feature    4218..4223
                     /label=-35
                     /ApEinfo_revcolor="#d6b295"
                     /ApEinfo_fwdcolor="#d6b295"
     promoter        4218..4251
                     /label="mobp"
                     /ApEinfo_revcolor="#d6b295"
                     /ApEinfo_fwdcolor="#d6b295"
     misc_feature    4226..4248
                     /label="recombination site A (RSA)"
                     /ApEinfo_revcolor="#d6b295"
                     /ApEinfo_fwdcolor="#d6b295"
     misc_feature    complement(4240..4240)
                     /label="nicking site"
                     /ApEinfo_revcolor="#d6b295"
                     /ApEinfo_fwdcolor="#d6b295"
     misc_feature    4240..4245
                     /label=-10
                     /ApEinfo_revcolor="#d6b295"
                     /ApEinfo_fwdcolor="#d6b295"
     CDS             4327..5328
                     /label="mob"
                     /ApEinfo_revcolor="#d6b295"
                     /ApEinfo_fwdcolor="#d6b295"
     misc_feature    5290..5319
                     /label="AmpR promoter homology"
```

```
                     /ApEinfo_revcolor="#c6c9d1"
                     /ApEinfo_fwdcolor="#c6c9d1"
     terminator      5333..5371
                     /label="bidir. modified thrLt BBa_B1006 "
                     /ApEinfo_revcolor="#c6c9d1"
                     /ApEinfo_fwdcolor="#c6c9d1"
     misc_feature    5333..5482
                     /label="ConL2"
                     /ApEinfo_revcolor="#c6c9d1"
                     /ApEinfo_fwdcolor="#c6c9d1"
     terminator      5372..5420
                     /label="tVoigtS5, L3S3P22 47C>G"
                     /ApEinfo_revcolor="#c6c9d1"
                     /ApEinfo_fwdcolor="#c6c9d1"
     misc_feature    5396..5420
                     /label="homology tVoigtS4"
                     /ApEinfo_revcolor="#f58a5e"
                     /ApEinfo_fwdcolor="#f58a5e"
     primer          5396..5420
                     /label="AB17 ConL F seq"
                     /note="sequence: ggcctttttttgtttctggtctgcc"
                     /ApEinfo_revcolor="#75c6a9"
                     /ApEinfo_fwdcolor="#75c6a9"
     misc_feature    5421..5435
                     /label="6-frame STOP a"
                     /ApEinfo_revcolor="#c6c9d1"
                     /ApEinfo_fwdcolor="#c6c9d1"
     misc_feature    5436..5441
                     /label="EcoRI"
                     /ApEinfo_revcolor="#d59687"
                     /ApEinfo_fwdcolor="#d59687"
     misc_feature    complement(5436..5457)
                     /label="BioBrick Prefix BBa_G00000"
                     /ApEinfo_revcolor="#d59687"
                     /ApEinfo_fwdcolor="#d59687"
     misc_feature    5442..5449
                     /label="NotI"
                     /ApEinfo_revcolor="#b7e6d7"
                     /ApEinfo_fwdcolor="#b7e6d7"
     misc_feature    5451..5456
                     /label="XbaI"
                     /ApEinfo_revcolor="#d59687"
                     /ApEinfo_fwdcolor="#d59687"
     misc_feature    5458..5463
                     /label="Esp3I"
                     /ApEinfo_revcolor="#faac61"
                     /ApEinfo_fwdcolor="#faac61"
     primer          complement(5465..5486)
                     /label="oPK134 pcr81 f1R"
                     /note="sequence: caCGTCTCactctGAAGcctgcagggctagcAACG"
                     /ApEinfo_revcolor="#b4abac"
                     /ApEinfo_fwdcolor="#b4abac"
     misc_feature    5469..5474
                     /label="NheI"
                     /ApEinfo_revcolor="#c7b0e3"
                     /ApEinfo_fwdcolor="#c7b0e3"
     misc_feature    5475..5482
```

```
                     /label="SbfI"
                     /ApEinfo_revcolor="#85dae9"
                     /ApEinfo_fwdcolor="#85dae9"
     misc_feature    5487..5518
                     /label="CymR operator"
                     /ApEinfo_revcolor="#ffef86"
                     /ApEinfo_fwdcolor="#ffef86"
     promoter        5487..5576
                     /label="P.CymRC (Marionette)"
                     /ApEinfo_revcolor="#ffef86"
                     /ApEinfo_fwdcolor="#ffef86"
     misc_feature    5510..5515
                     /label=-35
                     /ApEinfo_revcolor="#ffef86"
                     /ApEinfo_fwdcolor="#ffef86"
     primer          complement(5532..5565)
                     /label="KRG107"
                     /note="sequence:
GGTCTCaagaccagattgtctgtttgttgaatctattatac"
                     /ApEinfo_revcolor="#ff9ccd"
                     /ApEinfo_fwdcolor="#ff9ccd"
     misc_feature    5533..5538
                     /label=-10
                     /ApEinfo_revcolor="#ffef86"
                     /ApEinfo_fwdcolor="#ffef86"
     misc_feature    5545..5545
                     /label="TSS"
                     /ApEinfo_revcolor="#ffef86"
                     /ApEinfo_fwdcolor="#ffef86"
     misc_feature    5545..5576
                     /label="CymR operator"
                     /ApEinfo_revcolor="#ffef86"
                     /ApEinfo_fwdcolor="#ffef86"
     primer          complement(5553..5580)
                     /label="KRG064"
                     /note="sequence: GGTCTCagtcgataatacaaacagaccagattgtc"
                     /ApEinfo_revcolor="#f8d3a9"
                     /ApEinfo_fwdcolor="#f8d3a9"
     misc_feature    5581..642
                     /label="U64 Ribozyme transcript"
                     /ApEinfo_revcolor="#d59687"
                     /ApEinfo_fwdcolor="#d59687"
                     /note="https://rnacentral.org/rna/URS0002349E27/5911"
     misc_feature    5581..5636
                     /label="U64 RNA Guide"
                     /ApEinfo_revcolor="#faac61"
                     /ApEinfo_fwdcolor="#faac61"
     misc_feature    5631..5636
                     /label="IGS"
                     /ApEinfo_revcolor="#c7b0e3"
                     /ApEinfo_fwdcolor="#c7b0e3"
     misc_feature    5637..119
                     /label="Group I Intron Ribozyme (from p-OiRS3GG)"
                     /ApEinfo_revcolor="#b4abac"
                     /ApEinfo_fwdcolor="#b4abac"
     primer          5637..5659
                     /label="KRG063"
```

```
                     /note="sequence: GGTCTCaaaaagttatcaggcatgcacctg"
                     /ApEinfo_revcolor="#9eafd2"
                     /ApEinfo_fwdcolor="#9eafd2"
     primer          5639..5660
                     /label="KRG106"
                     /note="sequence: GGTCTCaagttatcaggcatgcacctgg"
                     /ApEinfo_revcolor="#75c6a9"
                     /ApEinfo_fwdcolor="#75c6a9"
ORIGIN
        1 CTCATAAGAT ATAGTCGGAC CTCTCCTTAA TGGGAGCTAG CGGATGAAGT GATGCAACAC
       61 TGGAGCCGCT GGGAACTAAT TTGTATGCGA AAGTATATTG ATTAGTTTTG GAGTACTCGA
      121 TGGTGTTCAA TGCTTTTCCC GAGTGGTTAT CCGGATCACA TGAAACGGCA TGACTTTTTC
      181 AAGAGTGCCA TGCCCGAAGG TTATGTACAG GAACGCACTA TATCTTTCAA AGATGACGGG
      241 ACCTACAAGA CGCGTGCTGA AGTCAAGTTT GAAGGTGATA CCCTTGTTAA TCGTATCGAG
      301 TTAAAGGGTA TTGATTTTAA AGAAGATGGA AACATTCTTG GACACAAACT CGAGTACAAC
      361 TTTAACTCAC ACAATGTATA CATCACGGCA GACAAACAAA AGAATGGAAT CAAAGCTAAC
      421 TTCAAAATTC GCCACAACGT TGAAGATGGT TCCGTTCAAC TAGCAGACCA TTATCAACAA
      481 AATACTCCAA TTGGCGATGG CCCTGTCCTT TTACCAGACA ACCATTACCT GTCGACACAA
      541 TCTGTCCTTT CGAAAGATCC CAACGAAAAG CGTGACCACA TGGTCCTTCT TGAGTTTGTA
      601 ACTGCTGCTG GGATTACACA TGGCATGGAT GAGCTCTACA AATGGCccaa ttattgaagg
      661 cctccctaac gggggccttt tttttgtttc tggtctGccG CTGagttcac cgacaaacaa
      721 cagataaaac gaaaggccca gtctttcgac tgagcctttc gttttatttg atgcctggca
      781 gttccctact ctcgcatggg gagaccccac actaccatcg gcgctacggc gtttcacttc
      841 tgagttcggc atggggtcag gtgggaccac cgcgctactg ccgccaggca aattctgttt
      901 tatcagaccg cttctgcgtt ctgatttaat ctgtatcagg ctgaaaatct tctctcatcc
      961 gccaaaacag ccaagctgga gaccgtttaa actcaatgat gatgatgatg atggtcgacg
     1021 gcgctattca gatcctcttc tgagatgagt ttttgttcgg gcccaagctt cagatccCAT
     1081 Attatattcc ccagaacatc aggttaatgg cgtttttgat gtcattttcg cggtggctga
     1141 gatcagccac ttcttccccg ataacggaga ccggcacact ggccatatcg gtggtcatca
     1201 tgcgccagct ttcatccccg atatgcacca ccgggtaaag ttcacgggag actttatctg
     1261 acagcagacg tgcactggcc aggggggatca ccatccgtcg cccgggcgtg tcaataatat
     1321 cactctgtac atccacaaac agacgataac ggctctctct tttataggtg taaaccttaa
     1381 actgcatAGG Tcattcctcc tagatccaaa atacggtacc gtcaacaatc tcactcgaga
     1441 gtcgacgtgc tagcattgta cctaggactg agctagccat aaaCGTTtGA GACGtactag
     1501 tagcggccgc tgcagttaat cactgattaa ctcggtacca aattccagaa aagaggcctc
     1561 ccgaaagggg ggcctttttt cgttttggtc cttttgttat caataaaaaa ggggagcggt
     1621 ttcccgctcc ccttattgtt cgtcTACAat tattagaaga actcgtcaag aaggcgatag
     1681 aaggcgatgc gctgcgaatc gggagcggcg ataccgtaaa gcacgaggaa gcggtcagcc
     1741 cattcgccgc caagctcttc agcaatatca cgggtagcca acgctatgtc ctgatagcgg
     1801 tccgccacac ccagccggcc acagtcgatg aatccagaaa agcggccatt ttccaccatg
     1861 atattcggca agcaggcatc gccatgggtc acgacgagat cctcgccgtc gggcatgcgc
     1921 gccttgagcc tggcgaacag ttcggctggc gcgagcccct gatgctcttc gtccagatca
     1981 tcctgatcga caagaccggc ttccatccga gtacgtgctc gctcgatgcg atgtttcgct
     2041 tggtggtcga atgggcaggt agccggatca agcgtatgca gccgccgcat tgcatcagcc
     2101 atgatggata ctttctcggc aggagcaagg tgagatgaca ggagatcctg ccccggcact
     2161 tcgcccaata gcagccagtc ccttcccgct tcagtgacaa cgtcgagcac agctgcgcaa
     2221 ggaacgcccg tcgtggccag ccacgatagc cgcgctgcct cgtcctgcag ttcattcagg
     2281 gcaccggaca ggtcggtctt gacaaaaaga accgggcgcc cctgcgctga cagccggaac
     2341 acggcggcat cagagcagcc gattgtctgt tgtgcccagt catagccgaa tagcctctcc
     2401 acccaagcgg ccggagaacc tgcgtgcaat ccatcttgtt caatcatgcg aaacgatcct
     2461 catcctgtct cttgatcaga tcttgatccc ctgcgccatc agatccttgg cggcaagaaa
     2521 gccatccagt ttactttgca gggcttccca accttaccag agggcgcccc agctggcaat
     2581 tccggttcgc ttgctgtcca taaGAGTcct gttgagtaat agtcaaaagc ctccggtcgg
     2641 aggcttttga ctttctgctt acCCAAttac taccggcgcg gcagcgtgac ccgtgtcggc
     2701 ggctccaacg gctcgccatc gtccagaaaa cacggctcat cgggcatcgg caggcgctgc
     2761 tgcccgcgcc gttcccattc ctccgtttcg gtcaaggctg gcaggtctgg ttccatgccc
     2821 ggaatgccgg gctggctggg cggctcctcg ccggggccgg tcggtagttg ctgctcgccc
```

```
     2881 ggatacaggg tcgggatgcg gcgcaggtcg ccatgcccca acagcgattc gtcctggtcg
     2941 tcgtgatcaa ccaccacggc ggcactgaac accgacaggc gcaactggtc gcggggctgg
     3001 ccccacgcca cgcggtcatt gaccacgtag gccgacacgg tgccggggcc gttgagcttc
     3061 acgacggaga tccagcgctc ggccaccaag tccttgactg cgtattggac cgtccgcaaa
     3121 gaacgtccga tgagcttgga aagtgttttc tggctgacca ccacggcgtt ctggtggccc
     3181 atctgcgcca cgaggtgatg cagcagcatt gccgccgtgg gtttcctcgc aataagcccg
     3241 gcccacgcct catgcgcttt gcgttccgtt tgcacccagt gaccgggctt gttcttggct
     3301 tgaatgccga tttctctgga ctgcgtggcc atgcttatct ccatgcggta ggggtgccgc
     3361 acggttgcgg caccatgcgc aatcagctgc aacttttcgg cagcgcgaca acaattatgc
     3421 gttgcgtaaa agtggcagtc aattacagat ttctttaac ctacgcaatg agctattgcg
     3481 gggggtgccg caatgagctg ttgcgtaccc cccttttta agttgttgat ttttaagtct
     3541 ttcgcatttc gccctatatc tagttctttg gtgcccaaag aagggcaccc ctgcggggtt
     3601 cccccacgcc ttcggcgcgg ctccccctcc ggcaaaaagt ggcccctccg ggcttgttg
     3661 atcgactgcg cggccttcgg ccttgcccaa ggtggcgctg ccccccttgga acccccgcac
     3721 tcgccgccgt gaggctcggg gggcaggcgg gcgggcttcg ccttcgactg cccccactcg
     3781 cataggcttg ggtcgttcca ggcgcgtcaa ggccaagccg ctgcgcggtc gctgcgcgag
     3841 ccttgacccg ccttccactt ggtgtccaac cggcaagcga agcgcgcagg ccgcaggccg
     3901 gaggcttttc cccagagaaa attaaaaaaa ttgatggggc aaggccgcag gccgcgcagt
     3961 tggagccggt gggtatgtgg tcgaaggctg ggtagccggt gggcaatccc tgtggtcaag
     4021 ctcgtgggca ggcgcagcct gtccatcagc ttgtccagca gggttgtcca cgggccgagc
     4081 gaagcgagcc agccggtggc cgctcgcggc catcgtccac atatccacgg gctggcaagg
     4141 gagcgcagcg accgcgcagg gcgaagcccg gagagcaagc ccgtagggcg ccgcagccgc
     4201 cgtaggcggt cacgactttg cgaagcaaag tctagtgagt atactcaagc attgagtggc
     4261 ccgccggagg caccgccttg cgctgccccc gtcgagccgg ttggacacca aaagggaggg
     4321 gcaggcatgg cggcatacgc gatcatgcga tgcaagaagc tggcgaaaat gggcaacgtg
     4381 gcggccagtc tcaagcacgc ctaccgcgag cgcgagactc ccaacgctga cgccagcagg
     4441 acgccagaga acgagcactg ggcggccagc agcaccgatg aagcgatggg ccgactgcgc
     4501 gagttgctgc cagagaagcg gcgcaaggac gctgtgttgg cggtcgagta cgtcatgacg
     4561 gccagcccgg aatggtggaa gtcggccagc caagaacagc aggcggcgtt cttcgagaag
     4621 gcgcacaagt ggctggcgga caagtacggg gcggatcgca tcgtgacggc cagcatccac
     4681 cgtgacgaaa ccagcccgca catgaccgcg ttcgtggtgc cgctgacgca ggacggcagg
     4741 ctgtcggcca aggagttcat cggcaacaaa gcgcagatga cccgcgacca gaccacgttt
     4801 gcggcAgctg tggccgatct agggctgcaa cggggcatcg agggcagcaa ggcacgtcac
     4861 acgcgcattc aggcgttcta cgaggccctg gagcggccac cagtgggcca cgtcaccatc
     4921 agcccgcaag cggtcgagcc acgcgcctat gcaccgcagg gattggccga aaagctggga
     4981 atctcaaagc gcgttgaAac gccggaagcc gtggccgacc ggctgacaaa agcggttcgg
     5041 caggggtatg agcctgccct acaggccgcc gcaggagcgc gtgagatgcg caagaaggcc
     5101 gatcaagccc aagagacagc ccgagaTctt cgggagcgcc tgaagcccgt tctggacgcc
     5161 ctggggccgt tgaatcggga tatgcaggcc aaggccgccg cgatcatcaa ggccgtgggc
     5221 gaaaagctgc tgacggaaca gcgggaagtc cagcgccaga aacaggccca gcgccagcag
     5281 gaacgcgggc gcgcacattt ccccgaaaag tgccacctgg gctgataaAC GGaaaaaaaa
     5341 accccgcccc tgacagggcg gggttttttt tccaattatt gaaggccgct aacgcggcct
     5401 tttttgttt ctggtctgcc ttaatcaatg actaagaatt cgcggccgct tctagagCGT
     5461 CTCaCGTTgc tagccctgca ggCTTCaaca aacagacaat ctggtctgtt tgtattatgg
     5521 aaaattttc tgtataatag attcaacaaa cagacaatct ggtctgtttg tattatcgac
     5581 CAACCCACTC CCATGGTGTG ACGGGCGGTG TGTACAAGGC CCGGGAACGT gTTCACAAAA
     5641 GTTATCAGGC ATGCACCTGG TAGCTAGTCT TTAAACCAAT AGATTGCATC GGTTTAAAAG
     5701 GCAAGACCGT CAAATTGCGG GAAAGGGGTC AACAGCCGTT CAGTACCAAG TCTCAGGGGA
     5761 AACTTTGAGA TGGCCTTGCA AAGGGTATGG TAATAAGCTG ACGGACATGG TCCTAACCAC
     5821 GCAGCCAAGT CCTAAGTCAA CAGATCTTCT GTTGATATGG ATGCAGTTCA CAGACTAAAT
     5881 GTCGGTCGGG GAAGATGTAT TCTT
//
```

**pMAS053**

```
LOCUS       pMAS053                  5899 bp ds-DNA     circular     12-SEP-
2025
DEFINITION  .
Aignments
FEATURES             Location/Qualifiers
     terminator      1..39
                     /label="bidir. modified thrLt BBa_B1006 "
                     /ApEinfo_revcolor="#c6c9d1"
                     /ApEinfo_fwdcolor="#c6c9d1"
     misc_feature    1..150
                     /label="ConL2"
                     /ApEinfo_revcolor="#c6c9d1"
                     /ApEinfo_fwdcolor="#c6c9d1"
     terminator      40..88
                     /label="tVoigtS5, L3S3P22 47C>G"
                     /ApEinfo_revcolor="#c6c9d1"
                     /ApEinfo_fwdcolor="#c6c9d1"
     misc_feature    64..88
                     /label="homology tVoigtS4"
                     /ApEinfo_revcolor="#f58a5e"
                     /ApEinfo_fwdcolor="#f58a5e"
     primer          64..88
                     /label="AB17 ConL F seq"
                     /note="sequence: ggccttttttttgtttctggtctgcc"
                     /ApEinfo_revcolor="#75c6a9"
                     /ApEinfo_fwdcolor="#75c6a9"
     misc_feature    89..103
                     /label="6-frame STOP a"
                     /ApEinfo_revcolor="#c6c9d1"
                     /ApEinfo_fwdcolor="#c6c9d1"
     misc_feature    104..109
                     /label="EcoRI"
                     /ApEinfo_revcolor="#d59687"
                     /ApEinfo_fwdcolor="#d59687"
     misc_feature    complement(104..125)
                     /label="BioBrick Prefix BBa_G00000"
                     /ApEinfo_revcolor="#d59687"
                     /ApEinfo_fwdcolor="#d59687"
     misc_feature    110..117
                     /label="NotI"
                     /ApEinfo_revcolor="#b7e6d7"
                     /ApEinfo_fwdcolor="#b7e6d7"
     misc_feature    119..124
                     /label="XbaI"
                     /ApEinfo_revcolor="#d59687"
                     /ApEinfo_fwdcolor="#d59687"
     misc_feature    126..131
                     /label="Esp3I"
                     /ApEinfo_revcolor="#faac61"
                     /ApEinfo_fwdcolor="#faac61"
     primer          complement(133..154)
                     /label="oPK134 pcr81 f1R"
                     /note="sequence: caCGTCTCactctGAAGcctgcagggctagcAACG"
                     /ApEinfo_revcolor="#b4abac"
```

```
                     /ApEinfo_fwdcolor="#b4abac"
     misc_feature    137..142
                     /label="NheI"
                     /ApEinfo_revcolor="#c7b0e3"
                     /ApEinfo_fwdcolor="#c7b0e3"
     misc_feature    143..150
                     /label="SbfI"
                     /ApEinfo_revcolor="#85dae9"
                     /ApEinfo_fwdcolor="#85dae9"
     misc_feature    155..186
                     /label="CymR operator"
                     /ApEinfo_revcolor="#ffef86"
                     /ApEinfo_fwdcolor="#ffef86"
     promoter        155..244
                     /label="P.CymRC (Marionette)"
                     /ApEinfo_revcolor="#ffef86"
                     /ApEinfo_fwdcolor="#ffef86"
     misc_feature    178..183
                     /label=-35
                     /ApEinfo_revcolor="#ffef86"
                     /ApEinfo_fwdcolor="#ffef86"
     primer          complement(200..233)
                     /label="KRG107"
                     /note="sequence:
GGTCTCaagaccagattgtctgtttgttgaatctattatac"
                     /ApEinfo_revcolor="#ff9ccd"
                     /ApEinfo_fwdcolor="#ff9ccd"
     misc_feature    201..206
                     /label=-10
                     /ApEinfo_revcolor="#ffef86"
                     /ApEinfo_fwdcolor="#ffef86"
     misc_feature    213..213
                     /label="TSS"
                     /ApEinfo_revcolor="#ffef86"
                     /ApEinfo_fwdcolor="#ffef86"
     misc_feature    213..244
                     /label="CymR operator"
                     /ApEinfo_revcolor="#ffef86"
                     /ApEinfo_fwdcolor="#ffef86"
     primer          complement(221..248)
                     /label="KRG064"
                     /note="sequence: GGTCTCagtcgataatacaaacagaccagattgtc"
                     /ApEinfo_revcolor="#f8d3a9"
                     /ApEinfo_fwdcolor="#f8d3a9"
     misc_feature    249..304
                     /label="U64 RNA Guide"
                     /ApEinfo_revcolor="#faac61"
                     /ApEinfo_fwdcolor="#faac61"
     misc_feature    249..1209
                     /label="U64 Ribozyme transcript"
                     /ApEinfo_revcolor="#d59687"
                     /ApEinfo_fwdcolor="#d59687"
                     /note="https://rnacentral.org/rna/URS0002349E27/5911"
     misc_feature    299..304
                     /label="IGS"
                     /ApEinfo_revcolor="#c7b0e3"
                     /ApEinfo_fwdcolor="#c7b0e3"
```

```
     primer          305..327
                     /label="KRG063"
                     /note="sequence: GGTCTCaaaaagttatcaggcatgcacctg"
                     /ApEinfo_revcolor="#9eafd2"
                     /ApEinfo_fwdcolor="#9eafd2"
     misc_feature    305..691
                     /label="Group I Intron Ribozyme (from p-OiRS3GG)"
                     /ApEinfo_revcolor="#b4abac"
                     /ApEinfo_fwdcolor="#b4abac"
     primer          307..328
                     /label="KRG106"
                     /note="sequence: GGTCTCaagttatcaggcatgcacctgg"
                     /ApEinfo_revcolor="#75c6a9"
                     /ApEinfo_fwdcolor="#75c6a9"
     misc_feature    692..712
                     /label="Probe binding site"
                     /ApEinfo_revcolor="#ff9ccd"
                     /ApEinfo_fwdcolor="#ff9ccd"
     primer          complement(692..713)
                     /label="KRG089"
                     /note="sequence: GCGGTCTCAgcgcCGGGAAAAGCATTGAACACCAT"
                     /ApEinfo_revcolor="#b7e6d7"
                     /ApEinfo_fwdcolor="#b7e6d7"
     primer          complement(692..713)
                     /label="oMJD105"
                     /note="sequence: GCGGTCTCAcactcGGGAAAAGCATTGAACACCAT"
                     /ApEinfo_revcolor="#b7e6d7"
                     /ApEinfo_fwdcolor="#b7e6d7"
     misc_feature    692..1209
                     /label="Barcode (sfGFP_2)"
                     /ApEinfo_revcolor="#84b0dc"
                     /ApEinfo_fwdcolor="#84b0dc"
     primer          711..732
                     /label="KRG090"
                     /note="sequence: GCGGTCTCAgcgcGGTTATCCGGATCACATGAAA"
                     /ApEinfo_revcolor="#b4abac"
                     /ApEinfo_fwdcolor="#b4abac"
     primer          713..732
                     /label="oMJD104"
                     /note="sequence: GCGGTCTCAGAGTGGTTATCCGGATCACATGAAA"
                     /ApEinfo_revcolor="#b4abac"
                     /ApEinfo_fwdcolor="#b4abac"
     misc_feature    1122..1127
                     /label="BamHI-BglII Scar"
                     /ApEinfo_revcolor="#ac1dff"
                     /ApEinfo_fwdcolor="#ac1dff"
     terminator      1214..1266
                     /label="tVoigtS4 L3S3P21 51C>G"
                     /ApEinfo_revcolor="#f58a5e"
                     /ApEinfo_fwdcolor="#f58a5e"
     misc_feature    1242..1266
                     /label="homology tVoigtS5"
                     /ApEinfo_revcolor="#f58a5e"
                     /ApEinfo_fwdcolor="#f58a5e"
     primer          1242..1266
                     /label="AB17 ConL F seq"
                     /note="sequence: ggccttttttgtttctggtctgcc"
```

```
                     /ApEinfo_revcolor="#75c6a9"
                     /ApEinfo_fwdcolor="#75c6a9"
     primer          complement(1253..1270)
                     /label="MAS65B"
                     /note="sequence: CAGCggCagaccagaaac"
                     /ApEinfo_revcolor="#b1ff67"
                     /ApEinfo_fwdcolor="#b1ff67"
     primer          1256..1298
                     /label="MAS62B"
                     /note="sequence:
tctggtctGccGCTGagttcaccgacaaacaacagataaaacg"
                     /ApEinfo_revcolor="#ffef86"
                     /ApEinfo_fwdcolor="#ffef86"
     misc_feature    complement(1271..1638)
                     /label="TrrnB"
                     /ApEinfo_revcolor="#85dae9"
                     /ApEinfo_fwdcolor="#85dae9"
     misc_feature    1271..2015
                     /label="from pMAS049"
                     /ApEinfo_revcolor="#75c6a9"
                     /ApEinfo_fwdcolor="#75c6a9"
     primer          complement(1382..1407)
                     /label="JEC20A | JC.A20"
                     /note="sequence: gaagtgaaacgccgtagcgccgatgg"
                     /ApEinfo_revcolor="#9eafd2"
                     /ApEinfo_fwdcolor="#9eafd2"
     misc_feature    1530..1547
                     /label="JEC71A"
                     /ApEinfo_revcolor="#b7e6d7"
                     /ApEinfo_fwdcolor="#b7e6d7"
     primer          1530..1547
                     /label="JEC71A"
                     /note="sequence: caaaacagccaagctgga"
                     /ApEinfo_revcolor="#b7e6d7"
                     /ApEinfo_fwdcolor="#b7e6d7"
     misc_feature    1597..1602
                     /label="BamHI-BglII Scar"
                     /ApEinfo_revcolor="#ac1dff"
                     /ApEinfo_fwdcolor="#ac1dff"
     primer          complement(1619..1644)
                     /label="AOS49A"
                     /note="sequence: ggatctgaagcttgggcccgaacaaa"
                     /ApEinfo_revcolor="#c7b0e3"
                     /ApEinfo_fwdcolor="#c7b0e3"
     misc_feature    complement(1627..1648)
                     /label="LCG38F"
                     /ApEinfo_revcolor="#85dae9"
                     /ApEinfo_fwdcolor="#85dae9"
     primer          complement(1627..1648)
                     /label="poscon temp fwd"
                     /note="sequence: TATGggatctgaagcttgggcc"
                     /ApEinfo_revcolor="#d59687"
                     /ApEinfo_fwdcolor="#d59687"
     primer          complement(1629..1663)
                     /label="MAS60B"
                     /note="sequence: ttctggggaatataaTATGggatctgaagcttggg"
                     /ApEinfo_revcolor="#c7b0e3"
```

```
                     /ApEinfo_fwdcolor="#c7b0e3"
     primer          1634..1676
                     /label="MAS59B"
                     /note="sequence:
gcttcagatccCATAttatattccccagaacatcaggttaatg"
                     /ApEinfo_revcolor="#f58a5e"
                     /ApEinfo_fwdcolor="#f58a5e"
     CDS             complement(1636..1648)
                     /label="Translation 1648-1636"
                     /translation="YGI*"
     misc_feature    1639..1644
                     /label="BamHI-BglII Scar"
                     /ApEinfo_revcolor="#ac1dff"
                     /ApEinfo_fwdcolor="#ac1dff"
     misc_feature    1645..1648
                     /label="LCG54G"
                     /ApEinfo_revcolor="#9eafd2"
                     /ApEinfo_fwdcolor="#9eafd2"
     CDS             complement(1649..1954)
                     /label="ccdB CDS (type II toxin-antitoxin system toxin
CcdB)"
                     /ApEinfo_revcolor="#b4abac"
                     /ApEinfo_fwdcolor="#b4abac"
     gene            complement(1649..1954)
                     /label="ccdB gene"
                     /ApEinfo_revcolor="#faac61"
                     /ApEinfo_fwdcolor="#faac61"
     primer          complement(1924..1970)
                     /label="MAS58B"
                     /note="sequence:
ctaggaggaatgACCTatgcagtttaaggtttacacctataaaagag"
                     /ApEinfo_revcolor="#f58a5e"
                     /ApEinfo_fwdcolor="#f58a5e"
     primer          1940..1981
                     /label="MAS61B"
                     /note="sequence:
aaccttaaactgcatAGGTcattcctcctagatccaaaatac"
                     /ApEinfo_revcolor="#c7b0e3"
                     /ApEinfo_fwdcolor="#c7b0e3"
     misc_feature    complement(1955..1955)
                     /label="Catalytic U"
                     /ApEinfo_revcolor="#c7b0e3"
                     /ApEinfo_fwdcolor="#c7b0e3"
     misc_feature    complement(1955..1960)
                     /label="IGS"
                     /ApEinfo_revcolor="#c7b0e3"
                     /ApEinfo_fwdcolor="#c7b0e3"
     primer          1955..1982
                     /label="MAS007A"
                     /note="sequence: AGGTcattcctcctagatccaaaatacg"
                     /ApEinfo_revcolor="#d59687"
                     /ApEinfo_fwdcolor="#d59687"
     misc_feature    complement(1961..1968)
                     /label="RBS-Strong"
                     /ApEinfo_revcolor="#9eafd2"
                     /ApEinfo_fwdcolor="#9eafd2"
     misc_feature    1969..1974
```

```
                     /label="BamHI-BglII Scar"
                     /ApEinfo_revcolor="#ac1dff"
                     /ApEinfo_fwdcolor="#ac1dff"
     misc_feature    complement(1975..2015)
                     /label="HP14 Stability Hairpin"
                     /ApEinfo_revcolor="#fa1839"
                     /ApEinfo_fwdcolor="#fa1839"
     primer          complement(2008..2032)
                     /label="MAS73B"
                     /note="sequence: ctaggtacaatgctagcacgtcgac"
                     /ApEinfo_revcolor="#75c6a9"
                     /ApEinfo_fwdcolor="#75c6a9"
     misc_feature    complement(2016..2050)
                     /label="J23114"
                     /ApEinfo_revcolor="#85dae9"
                     /ApEinfo_fwdcolor="#85dae9"
     primer          2033..2060
                     /label="MAS72B"
                     /note="sequence: gactgagctagccataaaCGTTtGAGAC"
                     /ApEinfo_revcolor="#75c6a9"
                     /ApEinfo_fwdcolor="#75c6a9"
     primer          2051..2073
                     /label="MAS64B"
                     /note="sequence: CGTTtGAGACGtactagtagcgg"
                     /ApEinfo_revcolor="#b1ff67"
                     /ApEinfo_fwdcolor="#b1ff67"
     misc_feature    complement(2051..2211)
                     /label="ConR2"
                     /ApEinfo_revcolor="#c6c9d1"
                     /ApEinfo_fwdcolor="#c6c9d1"
     misc_feature    complement(2056..2061)
                     /label="Esp3I"
                     /ApEinfo_revcolor="#f58a5e"
                     /ApEinfo_fwdcolor="#f58a5e"
     misc_feature    complement(2062..2082)
                     /label="BioBrick Suffix BBa_G00001"
                     /ApEinfo_revcolor="#d59687"
                     /ApEinfo_fwdcolor="#d59687"
     misc_feature    2063..2068
                     /label="SpeI"
                     /ApEinfo_revcolor="#d59687"
                     /ApEinfo_fwdcolor="#d59687"
     misc_feature    2070..2077
                     /label="NotI"
                     /ApEinfo_revcolor="#b7e6d7"
                     /ApEinfo_fwdcolor="#b7e6d7"
     misc_feature    2077..2082
                     /label="PstI"
                     /ApEinfo_revcolor="#d59687"
                     /ApEinfo_fwdcolor="#d59687"
     misc_feature    2083..2097
                     /label="6-frame STOP b"
                     /ApEinfo_revcolor="#c6c9d1"
                     /ApEinfo_fwdcolor="#c6c9d1"
     primer          complement(2098..2122)
                     /label="AB18 ConR R seq"
                     /note="sequence: ctcttttctggaatttggtaccgag"
```

```
                     /ApEinfo_revcolor="#85dae9"
                     /ApEinfo_fwdcolor="#85dae9"
     terminator      2098..2158
                     /label="tVoigtS1, L3S2P21"
                     /ApEinfo_revcolor="#c6c9d1"
                     /ApEinfo_fwdcolor="#c6c9d1"
     terminator      complement(2159..2211)
                     /label="tVoigtS19, L3S1P00"
                     /ApEinfo_revcolor="#c6c9d1"
                     /ApEinfo_fwdcolor="#c6c9d1"
     gene            complement(2216..3170)
                     /label="KanR.II, kanamycin resistance"
                     /ApEinfo_revcolor="#f6989d"
                     /ApEinfo_fwdcolor="#f6989d"
     CDS             complement(2217..3014)
                     /label="aph(3')-II / nptII / neo, aminoglycoside 3'-
phosphotransferase type II (KanR)"
                     /ApEinfo_revcolor="#f6989d"
                     /ApEinfo_fwdcolor="#f6989d"
     misc_feature    complement(3015..3170)
                     /label="aphIIp, KanR promoter/UTR"
                     /ApEinfo_revcolor="#f6989d"
                     /ApEinfo_fwdcolor="#f6989d"
     misc_feature    complement(3118..3126)
                     /label="aphIIp -10"
                     /ApEinfo_revcolor="#f6989d"
                     /ApEinfo_fwdcolor="#f6989d"
     misc_feature    complement(3141..3146)
                     /label="aphIIp -35"
                     /ApEinfo_revcolor="#f6989d"
                     /ApEinfo_fwdcolor="#f6989d"
     terminator      3175..3229
                     /label="tVoigtN17, tonBt (bidir), BBa_B0054"
                     /ApEinfo_revcolor="#f58a5e"
                     /ApEinfo_fwdcolor="#f58a5e"
     CDS             complement(3234..3899)
                     /label="rep"
                     /ApEinfo_revcolor="#c6c9d1"
                     /ApEinfo_fwdcolor="#c6c9d1"
     misc_feature    complement(3234..4757)
                     /label="pBBR1 rep region"
                     /ApEinfo_revcolor="#c6c9d1"
                     /ApEinfo_fwdcolor="#c6c9d1"
     misc_feature    3234..5895
                     /label="pBBR1-mob origin"
                     /ApEinfo_revcolor="#c6c9d1"
                     /ApEinfo_fwdcolor="#c6c9d1"
     misc_feature    3767..3775
                     /label="unstable repeat. 1e-5 rate"
                     /ApEinfo_revcolor="#b4abac"
                     /ApEinfo_fwdcolor="#b4abac"
     promoter        complement(3975..4009)
                     /label="repp"
                     /ApEinfo_revcolor="#c6c9d1"
                     /ApEinfo_fwdcolor="#c6c9d1"
     misc_feature    complement(3981..3989)
                     /label=-10
```

```
                     /ApEinfo_revcolor="#c6c9d1"
                     /ApEinfo_fwdcolor="#c6c9d1"
     misc_feature    complement(4004..4009)
                     /label=-35
                     /ApEinfo_revcolor="#c6c9d1"
                     /ApEinfo_fwdcolor="#c6c9d1"
     misc_feature    complement(4086..4098)
                     /label="IHF binding site"
                     /ApEinfo_revcolor="#c6c9d1"
                     /ApEinfo_fwdcolor="#c6c9d1"
     primer          4757..4773
                     /label="oPK133 pcr81 f1L"
                     /note="sequence: gtCGTCTCactcgACGGccgcagccgccgtagg"
                     /ApEinfo_revcolor="#ff9ccd"
                     /ApEinfo_fwdcolor="#ff9ccd"
     misc_feature    4758..5895
                     /label="pBBR1 mob region"
                     /ApEinfo_revcolor="#d6b295"
                     /ApEinfo_fwdcolor="#d6b295"
     misc_feature    4775..4826
                     /label="oriT region"
                     /ApEinfo_revcolor="#d6b295"
                     /ApEinfo_fwdcolor="#d6b295"
     misc_feature    4785..4790
                     /label=-35
                     /ApEinfo_revcolor="#d6b295"
                     /ApEinfo_fwdcolor="#d6b295"
     promoter        4785..4818
                     /label="mobp"
                     /ApEinfo_revcolor="#d6b295"
                     /ApEinfo_fwdcolor="#d6b295"
     misc_feature    4793..4815
                     /label="recombination site A (RSA)"
                     /ApEinfo_revcolor="#d6b295"
                     /ApEinfo_fwdcolor="#d6b295"
     misc_feature    complement(4807..4807)
                     /label="nicking site"
                     /ApEinfo_revcolor="#d6b295"
                     /ApEinfo_fwdcolor="#d6b295"
     misc_feature    4807..4812
                     /label=-10
                     /ApEinfo_revcolor="#d6b295"
                     /ApEinfo_fwdcolor="#d6b295"
     CDS             4894..5895
                     /label="mob"
                     /ApEinfo_revcolor="#d6b295"
                     /ApEinfo_fwdcolor="#d6b295"
     misc_feature    5857..5886
                     /label="AmpR promoter homology"
                     /ApEinfo_revcolor="#c6c9d1"
                     /ApEinfo_fwdcolor="#c6c9d1"
ORIGIN
        1 aaaaaaaaac cccgcccctg acagggcggg gttttttttc caattattga aggccgctaa
       61 cgcggccttt ttttgtttct ggtctgcctt aatcaatgac taagaattcg cggccgcttc
      121 tagagCGTCT CaCGTTgcta gccctgcagg CTTCaacaaa cagacaatct ggtctgtttg
      181 tattatggaa aatttttctg tataatagat tcaacaaaca gacaatctgg tctgtttgta
      241 ttatcgacCA ACCCACTCCC ATGGTGTGAC GGGCGGTGTG TACAAGGCCC GGGAACGTgT
```

301 TCACAAAAGT TATCAGGCAT GCACCTGGTA GCTAGTCTTT AAACCAATAG ATTGCATCGG
361 TTTAAAAGGC AAGACCGTCA AATTGCGGGA AAGGGGTCAA CAGCCGTTCA GTACCAAGTC
421 TCAGGGGAAA CTTTGAGATG GCCTTGCAAA GGGTATGGTA ATAAGCTGAC GGACATGGTC
481 CTAACCACGC AGCCAAGTCC TAAGTCAACA GATCTTCTGT TGATATGGAT GCAGTTCACA
541 GACTAAATGT CGGTCGGGGA AGATGTATTC TTCTCATAAG ATATAGTCGG ACCTCTCCTT
601 AATGGGAGCT AGCGGATGAA GTGATGCAAC ACTGGAGCCG CTGGGAACTA ATTTGTATGC
661 GAAAGTATAT TGATTAGTTT TGGAGTACTC GATGGTGTTC AATGCTTTTC CCGTTATCCG
721 GATCACATGA AACGGCATGA CTTTTTCAAG AGTGCCATGC CCGAAGGTTA TGTACAGGAA
781 CGCACTATAT CTTTCAAAGA TGACGGGACC TACAAGACGC GTGCTGAAGT CAAGTTTGAA
841 GGTGATACCC TTGTTAATCG TATCGAGTTA AAGGGTATTG ATTTTAAAGA AGATGGAAAC
901 ATTCTTGGAC ACAAACTCGA GTACAACTTT AACTCACACA ATGTATACAT CACGGCAGAC
961 AAACAAAAGA ATGGAATCAA AGCTAACTTC AAAATTCGCC ACAACGTTGA AGATGGTTCC
1021 GTTCAACTAG CAGACCATTA TCAACAAAAT ACTCCAATTG GCGATGGCCC TGTCCTTTTA
1081 CCAGACAACC ATTACCTGTC GACACAATCT GTCCTTTCGA AAGATCCCAA CGAAAAGCGT
1141 GACCACATGG TCCTTCTTGA GTTTGTAACT GCTGCTGGGA TTACACATGG CATGGATGAG
1201 CTCTACAAAT GGCccaatta ttgaaggcct ccctaacggg gggccttttt ttgtttctgg
1261 tctGccGCTG agttcaccga caaacaacag ataaaacgaa aggcccagtc tttcgactga
1321 gcctttcgtt ttatttgatg cctggcagtt ccctactctc gcatggggag accccacact
1381 accatcggcg ctacggcgtt tcacttctga gttcggcatg gggtcaggtg ggaccaccgc
1441 gctactgccg ccaggcaaat tctgttttat cagaccgctt ctgcgttctg atttaatctg
1501 tatcaggctg aaaatcttct ctcatccgcc aaaacagcca agctggagac cgtttaaact
1561 caatgatgat gatgatgatg gtcgacggcg ctattcagat cctcttctga gatgagtttt
1621 tgttcgggcc caagcttcag atcccATAtt atattcccca gaacatcagg ttaatggcgt
1681 ttttgatgtc attttcgcgg tggctgagat cagccacttc ttccccgata acggagaccg
1741 gcacactggc catatcggtg gtcatcatgc gccagctttc atccccgata tgcaccaccg
1801 ggtaaagttc acgggagact ttatctgaca gcagacgtgc actggccagg gggatcacca
1861 tccgtcgccc gggcgtgtca ataatatcac tctgtacatc cacaaacaga cgataacggc
1921 tctctctttt ataggtgtaa accttaaact gcatAGGTca ttcctcctag atccaaaata
1981 cggtaccgtc aacaatctca ctcgagagtc gacgtgctag cattgtacct aggactgagc
2041 tagccataaa CGTTtGAGAC Gtactagtag cggccgctgc agttaatcac tgattaactc
2101 ggtaccaaat tccagaaaag aggcctcccg aaaggggggc ctttttcgt tttggtcctt
2161 ttgttatcaa taaaaaaggg gagcggtttc ccgctcccct tattgttcgt cTACAattat
2221 tagaagaact cgtcaagaag gcgatagaag gcgatgcgct gcgaatcggg agcggcgata
2281 ccgtaaagca cgaggaagcg gtcagcccat tcgccgccaa gctcttcagc aatatcacgg
2341 gtagccaacg ctatgtcctg atagcggtcc gccacaccca gccggccaca gtcgatgaat
2401 ccagaaaagc ggccattttc caccatgata ttcggcaagc aggcatcgcc atgggtcacg
2461 acgagatcct cgccgtcggg catgcgcgcc ttgagcctgg cgaacagttc ggctggcgcg
2521 agcccctgat gctcttcgtc cagatcatcc tgatcgacaa gaccggcttc catccgagta
2581 cgtgctcgct cgatgcgatg tttcgcttgg tggtcgaatg ggcaggtagc cggatcaagc
2641 gtatgcagcc gccgcattgc atcagccatg atggatactt tctcggcagg agcaaggtga
2701 gatgacagga gatcctgccc cggcacttcg cccaatagca gccagtccct tcccgcttca
2761 gtgacaacgt cgagcacagc tgcgcaagga acgcccgtcg tggccagcca cgatagccgc
2821 gctgcctcgt cctgcagttc attcagggca ccggacaggt cggtcttgac aaaaagaacc
2881 gggcgcccct gcgctgacag ccggaacacg gcggcatcag agcagccgat tgtctgttgt
2941 gcccagtcat agccgaatag cctctccacc caagcggccg gagaacctgc gtgcaatcca
3001 tcttgttcaa tcatgcgaaa cgatcctcat cctgtctctt gatcagatct tgatcccctg
3061 cgccatcaga tccttggcgg caagaaagcc atccagttta ctttgcaggg cttcccaacc
3121 ttaccagagg gcgccccagc tggcaattcc ggttcgcttg ctgtccataa GAGTcctgtt
3181 gagtaatagt caaaagcctc cggtcggagg cttttgactt tctgcttacC CAAttactac
3241 cggcgcggca gcgtgacccg tgtcggcggc tccaacggct cgccatcgtc cagaaaacac
3301 ggctcatcgg gcatcggcag gcgctgctgc ccgcgccgtt cccattcctc cgtttcggtc
3361 aaggctggca ggtctggttc catgcccgga atgccgggct ggctgggcgg ctcctcgccg
3421 gggccggtcg gtagttgctg ctcgcccgga tacagggtcg ggatgcggcg caggtcgcca
3481 tgccccaaca gcgattcgtc ctggtcgtcg tgatcaacca ccacggcggc actgaacacc
3541 gacaggcgca actggtcgcg gggctggccc cacgccacgc ggtcattgac cacgtaggcc
3601 gacacggtgc cggggccgtt gagcttcacg acggagatcc agcgctcggc caccaagtcc
3661 ttgactgcgt attggaccgt ccgcaaagaa cgtccgatga gcttggaaag tgttttctgg

```
3721 ctgaccacca cggcgttctg gtggcccatc tgcgccacga ggtgatgcag cagcattgcc
3781 gccgtgggtt tcctcgcaat aagcccggcc cacgcctcat gcgctttgcg ttccgtttgc
3841 acccagtgac cgggcttgtt cttggcttga atgccgattt ctctggactg cgtggccatg
3901 cttatctcca tgcggtaggg gtgccgcacg gttgcggcac catgcgcaat cagctgcaac
3961 ttttcggcag cgcgacaaca attatgcgtt gcgtaaaagt ggcagtcaat tacagatttt
4021 ctttaaccta cgcaatgagc tattgcgggg ggtgccgcaa tgagctgttg cgtacccccc
4081 tttttaagt tgttgatttt taagtctttc gcatttcgcc ctatatctag ttctttggtg
4141 cccaaagaag ggcacccctg cggggttccc ccacgccttc ggcgcggctc cccctccggc
4201 aaaaagtggc ccctccgggg cttgttgatc gactgcgcgg ccttcggcct tgcccaaggt
4261 ggcgctgccc ccttggaacc cccgcactcg ccgccgtgag gctcgggggg caggcgggcg
4321 ggcttcgcct tcgactgccc ccactcgcat aggcttgggt cgttccaggc gcgtcaaggc
4381 caagccgctg cgcggtcgct gcgcgagcct tgacccgcct tccacttggt gtccaaccgg
4441 caagcgaagc gcgcaggccg caggccggag gcttttcccc agagaaaatt aaaaaaattg
4501 atggggcaag gccgcaggcc gcgcagttgg agccggtggg tatgtggtcg aaggctgggt
4561 agccggtggg caatccctgt ggtcaagctc gtgggcaggc gcagcctgtc catcagcttg
4621 tccagcaggg ttgtccacgg gccgagcgaa gcgagccagc cggtggccgc tcgcggccat
4681 cgtccacata tccacgggct ggcaagggag cgcagcgacc gcgcagggcg aagcccggag
4741 agcaagcccg tagggcgccg cagccgccgt aggcggtcac gactttgcga agcaaagtct
4801 agtgagtata ctcaagcatt gagtggcccg ccggaggcac cgccttgcgc tgcccccgtc
4861 gagccggttg gacaccaaaa gggaggggca ggcatggcgg catacgcgat catgcgatgc
4921 aagaagctgg cgaaaatggg caacgtggcg gccagtctca agcacgccta ccgcgagcgc
4981 gagactccca acgctgacgc cagcaggacg ccagagaacg agcactgggc ggccagcagc
5041 accgatgaag cgatgggccg actgcgcgag ttgctgccag agaagcggcg caaggacgct
5101 gtgttggcgg tcgagtacgt catgacggcc agcccggaat ggtggaagtc ggccagccaa
5161 gaacagcagg cggcgttctt cgagaaggcg cacaagtggc tggcggacaa gtacggggcg
5221 gatcgcatcg tgacggccag catccaccgt gacgaaacca gcccgcacat gaccgcgttc
5281 gtggtgccgc tgacgcagga cggcaggctg tcggccaagg agttcatcgg caacaaagcg
5341 cagatgaccc gcgaccagac cacgtttgcg gcAgctgtgg ccgatctagg gctgcaacgg
5401 ggcatcgagg gcagcaaggc acgtcacacg cgcattcagg cgttctacga ggccctggag
5461 cggccaccag tgggccacgt caccatcagc ccgcaagcgg tcgagccacg cgcctatgca
5521 ccgcagggat tggccgaaaa gctgggaatc tcaaagcgcg ttgaAacgcc ggaagccgtg
5581 gccgaccggc tgacaaaagc ggttcggcag ggtatgagc ctgccctaca ggccgccgca
5641 ggagcgcgtg agatgcgcaa gaaggccgat caagcccaag agacagcccg agaTcttcgg
5701 gagcgcctga agcccgttct ggacgccctg ggccgttga atcgggatat gcaggccaag
5761 gccgccgcga tcatcaaggc cgtgggcgaa aagctgctga cggaacagcg ggaagtccag
5821 cgccagaaac aggcccagcg ccagcaggaa cgcgggcgcg cacatttccc cgaaaagtgc
5881 cacctgggct gataaACGG
//
```

**pMAS052**

```
LOCUS       pMAS052                  5899 bp ds-DNA     circular     20-AUG-
2025
DEFINITION  .
Aignments
FEATURES             Location/Qualifiers
     terminator      1..39
                     /label="bidir. modified thrLt BBa_B1006 "
                     /ApEinfo_revcolor="#c6c9d1"
                     /ApEinfo_fwdcolor="#c6c9d1"
     misc_feature    1..150
                     /label="ConL2"
                     /ApEinfo_revcolor="#c6c9d1"
                     /ApEinfo_fwdcolor="#c6c9d1"
     terminator      40..88
                     /label="tVoigtS5, L3S3P22 47C>G"
                     /ApEinfo_revcolor="#c6c9d1"
                     /ApEinfo_fwdcolor="#c6c9d1"
     misc_feature    64..88
                     /label="homology tVoigtS4"
                     /ApEinfo_revcolor="#f58a5e"
                     /ApEinfo_fwdcolor="#f58a5e"
     primer          64..88
                     /label="AB17 ConL F seq"
                     /note="sequence: ggcctttttttgtttctggtctgcc"
                     /ApEinfo_revcolor="#75c6a9"
                     /ApEinfo_fwdcolor="#75c6a9"
     misc_feature    89..103
                     /label="6-frame STOP a"
                     /ApEinfo_revcolor="#c6c9d1"
                     /ApEinfo_fwdcolor="#c6c9d1"
     misc_feature    104..109
                     /label="EcoRI"
                     /ApEinfo_revcolor="#d59687"
                     /ApEinfo_fwdcolor="#d59687"
     misc_feature    complement(104..125)
                     /label="BioBrick Prefix BBa_G00000"
                     /ApEinfo_revcolor="#d59687"
                     /ApEinfo_fwdcolor="#d59687"
     misc_feature    110..117
                     /label="NotI"
                     /ApEinfo_revcolor="#b7e6d7"
                     /ApEinfo_fwdcolor="#b7e6d7"
     misc_feature    119..124
                     /label="XbaI"
                     /ApEinfo_revcolor="#d59687"
                     /ApEinfo_fwdcolor="#d59687"
     misc_feature    126..131
                     /label="Esp3I"
                     /ApEinfo_revcolor="#faac61"
                     /ApEinfo_fwdcolor="#faac61"
     primer          complement(133..154)
                     /label="oPK134 pcr81 f1R"
                     /note="sequence: caCGTCTCactctGAAGcctgcagggctagcAACG"
                     /ApEinfo_revcolor="#b4abac"
```

```
                     /ApEinfo_fwdcolor="#b4abac"
     misc_feature    137..142
                     /label="NheI"
                     /ApEinfo_revcolor="#c7b0e3"
                     /ApEinfo_fwdcolor="#c7b0e3"
     misc_feature    143..150
                     /label="SbfI"
                     /ApEinfo_revcolor="#85dae9"
                     /ApEinfo_fwdcolor="#85dae9"
     misc_feature    155..186
                     /label="CymR operator"
                     /ApEinfo_revcolor="#ffef86"
                     /ApEinfo_fwdcolor="#ffef86"
     promoter        155..244
                     /label="P.CymRC (Marionette)"
                     /ApEinfo_revcolor="#ffef86"
                     /ApEinfo_fwdcolor="#ffef86"
     misc_feature    178..183
                     /label=-35
                     /ApEinfo_revcolor="#ffef86"
                     /ApEinfo_fwdcolor="#ffef86"
     primer          complement(200..233)
                     /label="KRG107"
                     /note="sequence:
GGTCTCaagaccagattgtctgtttgttgaatctattatac"
                     /ApEinfo_revcolor="#ff9ccd"
                     /ApEinfo_fwdcolor="#ff9ccd"
     misc_feature    201..206
                     /label=-10
                     /ApEinfo_revcolor="#ffef86"
                     /ApEinfo_fwdcolor="#ffef86"
     misc_feature    213..213
                     /label="TSS"
                     /ApEinfo_revcolor="#ffef86"
                     /ApEinfo_fwdcolor="#ffef86"
     misc_feature    213..244
                     /label="CymR operator"
                     /ApEinfo_revcolor="#ffef86"
                     /ApEinfo_fwdcolor="#ffef86"
     primer          complement(221..248)
                     /label="KRG064"
                     /note="sequence: GGTCTCagtcgataatacaaacagaccagattgtc"
                     /ApEinfo_revcolor="#f8d3a9"
                     /ApEinfo_fwdcolor="#f8d3a9"
     misc_feature    249..304
                     /label="U64 RNA Guide"
                     /ApEinfo_revcolor="#faac61"
                     /ApEinfo_fwdcolor="#faac61"
     misc_feature    249..1209
                     /label="U64 Ribozyme transcript"
                     /ApEinfo_revcolor="#d59687"
                     /ApEinfo_fwdcolor="#d59687"
                     /note="https://rnacentral.org/rna/URS0002349E27/5911"
     misc_feature    299..304
                     /label="IGS"
                     /ApEinfo_revcolor="#c7b0e3"
                     /ApEinfo_fwdcolor="#c7b0e3"
```

```
     primer          305..327
                     /label="KRG063"
                     /note="sequence: GGTCTCaaaaagttatcaggcatgcacctg"
                     /ApEinfo_revcolor="#9eafd2"
                     /ApEinfo_fwdcolor="#9eafd2"
     misc_feature    305..691
                     /label="Group I Intron Ribozyme (from p-OiRS3GG)"
                     /ApEinfo_revcolor="#b4abac"
                     /ApEinfo_fwdcolor="#b4abac"
     primer          307..328
                     /label="KRG106"
                     /note="sequence: GGTCTCaagttatcaggcatgcacctgg"
                     /ApEinfo_revcolor="#75c6a9"
                     /ApEinfo_fwdcolor="#75c6a9"
     misc_feature    692..712
                     /label="Probe binding site"
                     /ApEinfo_revcolor="#ff9ccd"
                     /ApEinfo_fwdcolor="#ff9ccd"
     primer          complement(692..713)
                     /label="oMJD105"
                     /note="sequence: GCGGTCTCAcactcGGGAAAAGCATTGAACACCAT"
                     /ApEinfo_revcolor="#b7e6d7"
                     /ApEinfo_fwdcolor="#b7e6d7"
     primer          complement(692..713)
                     /label="KRG089"
                     /note="sequence: GCGGTCTCAgcgcCGGGAAAAGCATTGAACACCAT"
                     /ApEinfo_revcolor="#b7e6d7"
                     /ApEinfo_fwdcolor="#b7e6d7"
     misc_feature    692..1209
                     /label="Barcode (sfGFP_2)"
                     /ApEinfo_revcolor="#84b0dc"
                     /ApEinfo_fwdcolor="#84b0dc"
     primer          711..732
                     /label="KRG090"
                     /note="sequence: GCGGTCTCAgcgcGGTTATCCGGATCACATGAAA"
                     /ApEinfo_revcolor="#b4abac"
                     /ApEinfo_fwdcolor="#b4abac"
     primer          713..732
                     /label="oMJD104"
                     /note="sequence: GCGGTCTCAGAGTGGTTATCCGGATCACATGAAA"
                     /ApEinfo_revcolor="#b4abac"
                     /ApEinfo_fwdcolor="#b4abac"
     misc_feature    1122..1127
                     /label="BamHI-BglII Scar"
                     /ApEinfo_revcolor="#ac1dff"
                     /ApEinfo_fwdcolor="#ac1dff"
     terminator      1214..1266
                     /label="tVoigtS4 L3S3P21 51C>G"
                     /ApEinfo_revcolor="#f58a5e"
                     /ApEinfo_fwdcolor="#f58a5e"
     misc_feature    1242..1266
                     /label="homology tVoigtS5"
                     /ApEinfo_revcolor="#f58a5e"
                     /ApEinfo_fwdcolor="#f58a5e"
     primer          1242..1266
                     /label="AB17 ConL F seq"
                     /note="sequence: ggcctttttttgtttctggtctgcc"
```

```
                     /ApEinfo_revcolor="#75c6a9"
                     /ApEinfo_fwdcolor="#75c6a9"
     primer          complement(1253..1270)
                     /label="MAS65B"
                     /note="sequence: CAGCggCagaccagaaac"
                     /ApEinfo_revcolor="#b1ff67"
                     /ApEinfo_fwdcolor="#b1ff67"
     primer          1256..1298
                     /label="MAS62B"
                     /note="sequence:
tctggtctGccGCTGagttcaccgacaaacaacagataaaacg"
                     /ApEinfo_revcolor="#ffef86"
                     /ApEinfo_fwdcolor="#ffef86"
     misc_feature    complement(1271..1638)
                     /label="TrrnB"
                     /ApEinfo_revcolor="#85dae9"
                     /ApEinfo_fwdcolor="#85dae9"
     misc_feature    1271..2015
                     /label="from pMAS049"
                     /ApEinfo_revcolor="#75c6a9"
                     /ApEinfo_fwdcolor="#75c6a9"
     primer          complement(1382..1407)
                     /label="JEC20A | JC.A20"
                     /note="sequence: gaagtgaaacgccgtagcgccgatgg"
                     /ApEinfo_revcolor="#9eafd2"
                     /ApEinfo_fwdcolor="#9eafd2"
     misc_feature    1530..1547
                     /label="JEC71A"
                     /ApEinfo_revcolor="#b7e6d7"
                     /ApEinfo_fwdcolor="#b7e6d7"
     primer          1530..1547
                     /label="JEC71A"
                     /note="sequence: caaaacagccaagctgga"
                     /ApEinfo_revcolor="#b7e6d7"
                     /ApEinfo_fwdcolor="#b7e6d7"
     misc_feature    1597..1602
                     /label="BamHI-BglII Scar"
                     /ApEinfo_revcolor="#ac1dff"
                     /ApEinfo_fwdcolor="#ac1dff"
     primer          complement(1619..1644)
                     /label="AOS49A"
                     /note="sequence: ggatctgaagcttgggcccgaacaaa"
                     /ApEinfo_revcolor="#c7b0e3"
                     /ApEinfo_fwdcolor="#c7b0e3"
     misc_feature    complement(1627..1648)
                     /label="LCG38F"
                     /ApEinfo_revcolor="#85dae9"
                     /ApEinfo_fwdcolor="#85dae9"
     primer          complement(1627..1648)
                     /label="poscon temp fwd"
                     /note="sequence: TATGggatctgaagcttgggcc"
                     /ApEinfo_revcolor="#d59687"
                     /ApEinfo_fwdcolor="#d59687"
     primer          complement(1629..1663)
                     /label="MAS60B"
                     /note="sequence: ttctggggaatataaTATGggatctgaagcttggg"
                     /ApEinfo_revcolor="#c7b0e3"
```

```
                     /ApEinfo_fwdcolor="#c7b0e3"
     primer          1634..1676
                     /label="MAS59B"
                     /note="sequence:
gcttcagatccCATAttatattccccagaacatcaggttaatg"
                     /ApEinfo_revcolor="#f58a5e"
                     /ApEinfo_fwdcolor="#f58a5e"
     CDS             complement(1636..1648)
                     /label="Translation 1648-1636"
                     /translation="YGI*"
     misc_feature    1639..1644
                     /label="BamHI-BglII Scar"
                     /ApEinfo_revcolor="#ac1dff"
                     /ApEinfo_fwdcolor="#ac1dff"
     misc_feature    1645..1648
                     /label="LCG54G"
                     /ApEinfo_revcolor="#9eafd2"
                     /ApEinfo_fwdcolor="#9eafd2"
     gene            complement(1649..1954)
                     /label="ccdB gene"
                     /ApEinfo_revcolor="#faac61"
                     /ApEinfo_fwdcolor="#faac61"
     CDS             complement(1649..1954)
                     /label="ccdB CDS (type II toxin-antitoxin system toxin
CcdB)"
                     /ApEinfo_revcolor="#b4abac"
                     /ApEinfo_fwdcolor="#b4abac"
     primer          complement(1924..1970)
                     /label="MAS58B"
                     /note="sequence:
ctaggaggaatgACCTatgcagtttaaggtttacacctataaaagag"
                     /ApEinfo_revcolor="#f58a5e"
                     /ApEinfo_fwdcolor="#f58a5e"
     primer          1940..1981
                     /label="MAS61B"
                     /note="sequence:
aaccttaaactgcatAGGTcattcctcctagatccaaaatac"
                     /ApEinfo_revcolor="#c7b0e3"
                     /ApEinfo_fwdcolor="#c7b0e3"
     misc_feature    complement(1955..1955)
                     /label="Catalytic U"
                     /ApEinfo_revcolor="#c7b0e3"
                     /ApEinfo_fwdcolor="#c7b0e3"
     misc_feature    complement(1955..1960)
                     /label="IGS"
                     /ApEinfo_revcolor="#c7b0e3"
                     /ApEinfo_fwdcolor="#c7b0e3"
     primer          1955..1982
                     /label="MAS007A"
                     /note="sequence: AGGTcattcctcctagatccaaaatacg"
                     /ApEinfo_revcolor="#d59687"
                     /ApEinfo_fwdcolor="#d59687"
     misc_feature    complement(1961..1968)
                     /label="RBS-Strong"
                     /ApEinfo_revcolor="#9eafd2"
                     /ApEinfo_fwdcolor="#9eafd2"
     misc_feature    1969..1974
```

```
                    /label="BamHI-BglII Scar"
                    /ApEinfo_revcolor="#ac1dff"
                    /ApEinfo_fwdcolor="#ac1dff"
     misc_feature   complement(1975..2015)
                    /label="HP14 Stability Hairpin"
                    /ApEinfo_revcolor="#fa1839"
                    /ApEinfo_fwdcolor="#fa1839"
     primer         complement(2006..2032)
                    /label="MAS71B"
                    /note="sequence: ctaggtactatgctagcacgtcgactc"
                    /ApEinfo_revcolor="#f8d3a9"
                    /ApEinfo_fwdcolor="#f8d3a9"
     misc_feature   complement(2016..2050)
                    /label="J23105"
                    /ApEinfo_revcolor="#b7e6d7"
                    /ApEinfo_fwdcolor="#b7e6d7"
     primer         complement(2021..2065)
                    /label="MAS67B"
                    /note="sequence:
agtaCGTCTCaAACGtttacggctagctcagtcctaggtactatg"
                    /ApEinfo_revcolor="#d59687"
                    /ApEinfo_fwdcolor="#d59687"
     primer         2033..2058
                    /label="MAS70B"
                    /note="sequence: gactgagctagccgtaaaCGTTtGAG"
                    /ApEinfo_revcolor="#f8d3a9"
                    /ApEinfo_fwdcolor="#f8d3a9"
     primer         2051..2073
                    /label="MAS64B"
                    /note="sequence: CGTTtGAGACGtactagtagcgg"
                    /ApEinfo_revcolor="#b1ff67"
                    /ApEinfo_fwdcolor="#b1ff67"
     misc_feature   complement(2051..2211)
                    /label="ConR2"
                    /ApEinfo_revcolor="#c6c9d1"
                    /ApEinfo_fwdcolor="#c6c9d1"
     misc_feature   complement(2056..2061)
                    /label="Esp3I"
                    /ApEinfo_revcolor="#f58a5e"
                    /ApEinfo_fwdcolor="#f58a5e"
     misc_feature   complement(2062..2082)
                    /label="BioBrick Suffix BBa_G00001"
                    /ApEinfo_revcolor="#d59687"
                    /ApEinfo_fwdcolor="#d59687"
     misc_feature   2063..2068
                    /label="SpeI"
                    /ApEinfo_revcolor="#d59687"
                    /ApEinfo_fwdcolor="#d59687"
     misc_feature   2070..2077
                    /label="NotI"
                    /ApEinfo_revcolor="#b7e6d7"
                    /ApEinfo_fwdcolor="#b7e6d7"
     misc_feature   2077..2082
                    /label="PstI"
                    /ApEinfo_revcolor="#d59687"
                    /ApEinfo_fwdcolor="#d59687"
     misc_feature   2083..2097
```

```
                     /label="6-frame STOP b"
                     /ApEinfo_revcolor="#c6c9d1"
                     /ApEinfo_fwdcolor="#c6c9d1"
     primer          complement(2098..2122)
                     /label="AB18 ConR R seq"
                     /note="sequence: ctcttttctggaatttggtaccgag"
                     /ApEinfo_revcolor="#85dae9"
                     /ApEinfo_fwdcolor="#85dae9"
     terminator      2098..2158
                     /label="tVoigtS1, L3S2P21"
                     /ApEinfo_revcolor="#c6c9d1"
                     /ApEinfo_fwdcolor="#c6c9d1"
     terminator      complement(2159..2211)
                     /label="tVoigtS19, L3S1P00"
                     /ApEinfo_revcolor="#c6c9d1"
                     /ApEinfo_fwdcolor="#c6c9d1"
     gene            complement(2216..3170)
                     /label="KanR.II, kanamycin resistance"
                     /ApEinfo_revcolor="#f6989d"
                     /ApEinfo_fwdcolor="#f6989d"
     CDS             complement(2217..3014)
                     /label="aph(3')-II / nptII / neo, aminoglycoside 3'-
phosphotransferase type II (KanR)"
                     /ApEinfo_revcolor="#f6989d"
                     /ApEinfo_fwdcolor="#f6989d"
     misc_feature    complement(3015..3170)
                     /label="aphIIp, KanR promoter/UTR"
                     /ApEinfo_revcolor="#f6989d"
                     /ApEinfo_fwdcolor="#f6989d"
     misc_feature    complement(3118..3126)
                     /label="aphIIp -10"
                     /ApEinfo_revcolor="#f6989d"
                     /ApEinfo_fwdcolor="#f6989d"
     misc_feature    complement(3141..3146)
                     /label="aphIIp -35"
                     /ApEinfo_revcolor="#f6989d"
                     /ApEinfo_fwdcolor="#f6989d"
     terminator      3175..3229
                     /label="tVoigtN17, tonBt (bidir), BBa_B0054"
                     /ApEinfo_revcolor="#f58a5e"
                     /ApEinfo_fwdcolor="#f58a5e"
     CDS             complement(3234..3899)
                     /label="rep"
                     /ApEinfo_revcolor="#c6c9d1"
                     /ApEinfo_fwdcolor="#c6c9d1"
     misc_feature    complement(3234..4757)
                     /label="pBBR1 rep region"
                     /ApEinfo_revcolor="#c6c9d1"
                     /ApEinfo_fwdcolor="#c6c9d1"
     misc_feature    3234..5895
                     /label="pBBR1-mob origin"
                     /ApEinfo_revcolor="#c6c9d1"
                     /ApEinfo_fwdcolor="#c6c9d1"
     misc_feature    3767..3775
                     /label="unstable repeat. 1e-5 rate"
                     /ApEinfo_revcolor="#b4abac"
                     /ApEinfo_fwdcolor="#b4abac"
```

```
     promoter        complement(3975..4009)
                     /label="repp"
                     /ApEinfo_revcolor="#c6c9d1"
                     /ApEinfo_fwdcolor="#c6c9d1"
     misc_feature    complement(3981..3989)
                     /label=-10
                     /ApEinfo_revcolor="#c6c9d1"
                     /ApEinfo_fwdcolor="#c6c9d1"
     misc_feature    complement(4004..4009)
                     /label=-35
                     /ApEinfo_revcolor="#c6c9d1"
                     /ApEinfo_fwdcolor="#c6c9d1"
     misc_feature    complement(4086..4098)
                     /label="IHF binding site"
                     /ApEinfo_revcolor="#c6c9d1"
                     /ApEinfo_fwdcolor="#c6c9d1"
     primer          4757..4773
                     /label="oPK133 pcr81 f1L"
                     /note="sequence: gtCGTCTCactcgACGGccgcagccgccgtagg"
                     /ApEinfo_revcolor="#ff9ccd"
                     /ApEinfo_fwdcolor="#ff9ccd"
     misc_feature    4758..5895
                     /label="pBBR1 mob region"
                     /ApEinfo_revcolor="#d6b295"
                     /ApEinfo_fwdcolor="#d6b295"
     misc_feature    4775..4826
                     /label="oriT region"
                     /ApEinfo_revcolor="#d6b295"
                     /ApEinfo_fwdcolor="#d6b295"
     misc_feature    4785..4790
                     /label=-35
                     /ApEinfo_revcolor="#d6b295"
                     /ApEinfo_fwdcolor="#d6b295"
     promoter        4785..4818
                     /label="mobp"
                     /ApEinfo_revcolor="#d6b295"
                     /ApEinfo_fwdcolor="#d6b295"
     misc_feature    4793..4815
                     /label="recombination site A (RSA)"
                     /ApEinfo_revcolor="#d6b295"
                     /ApEinfo_fwdcolor="#d6b295"
     misc_feature    complement(4807..4807)
                     /label="nicking site"
                     /ApEinfo_revcolor="#d6b295"
                     /ApEinfo_fwdcolor="#d6b295"
     misc_feature    4807..4812
                     /label=-10
                     /ApEinfo_revcolor="#d6b295"
                     /ApEinfo_fwdcolor="#d6b295"
     CDS             4894..5895
                     /label="mob"
                     /ApEinfo_revcolor="#d6b295"
                     /ApEinfo_fwdcolor="#d6b295"
     misc_feature    5857..5886
                     /label="AmpR promoter homology"
                     /ApEinfo_revcolor="#c6c9d1"
                     /ApEinfo_fwdcolor="#c6c9d1"
```

```
ORIGIN
        1 aaaaaaaaac cccgcccctg acagggcggg gttttttttc caattattga aggccgctaa
       61 cgcggccttt ttttgtttct ggtctgcctt aatcaatgac taagaattcg cggccgcttc
      121 tagagCGTCT CaCGTTgcta gccctgcagg CTTCaacaaa cagacaatct ggtctgtttg
      181 tattatggaa atttttctg tataatagat tcaacaaaca gacaatctgg tctgtttgta
      241 ttatcgacCA ACCCACTCCC ATGGTGTGAC GGGCGGTGTG TACAAGGCCC GGGAACGTgT
      301 TCACAAAAGT TATCAGGCAT GCACCTGGTA GCTAGTCTTT AAACCAATAG ATTGCATCGG
      361 TTTAAAAGGC AAGACCGTCA AATTGCGGGA AAGGGGTCAA CAGCCGTTCA GTACCAAGTC
      421 TCAGGGGAAA CTTTGAGATG GCCTTGCAAA GGGTATGGTA ATAAGCTGAC GGACATGGTC
      481 CTAACCACGC AGCCAAGTCC TAAGTCAACA GATCTTCTGT TGATATGGAT GCAGTTCACA
      541 GACTAAATGT CGGTCGGGGA AGATGTATTC TTCTCATAAG ATATAGTCGG ACCTCTCCTT
      601 AATGGGAGCT AGCGGATGAA GTGATGCAAC ACTGGAGCCG CTGGGAACTA ATTTGTATGC
      661 GAAAGTATAT TGATTAGTTT TGGAGTACTC GATGGTGTTC AATGCTTTTC CCGTTATCCG
      721 GATCACATGA AACGGCATGA CTTTTTCAAG AGTGCCATGC CCGAAGGTTA TGTACAGGAA
      781 CGCACTATAT CTTTCAAAGA TGACGGGACC TACAAGACGC GTGCTGAAGT CAAGTTTGAA
      841 GGTGATACCC TTGTTAATCG TATCGAGTTA AAGGGTATTG ATTTTAAAGA AGATGGAAAC
      901 ATTCTTGGAC ACAAACTCGA GTACAACTTT AACTCACACA ATGTATACAT CACGGCAGAC
      961 AAACAAAAGA ATGGAATCAA AGCTAACTTC AAAATTCGCC ACAACGTTGA AGATGGTTCC
     1021 GTTCAACTAG CAGACCATTA TCAACAAAAT ACTCCAATTG GCGATGGCCC TGTCCTTTTA
     1081 CCAGACAACC ATTACCTGTC GACACAATCT GTCCTTTCGA AAGATCCCAA CGAAAAGCGT
     1141 GACCACATGG TCCTTCTTGA GTTTGTAACT GCTGCTGGGA TTACACATGG CATGGATGAG
     1201 CTCTACAAAT GGCccaatta ttgaaggcct ccctaacggg gggccttttt ttgtttctgg
     1261 tctGccGCTG agttcaccga caaacaacag ataaaacgaa aggcccagtc tttcgactga
     1321 gcctttcgtt ttatttgatg cctggcagtt ccctactctc gcatggggag accccacact
     1381 accatcggcg ctacggcgtt tcacttctga gttcggcatg gggtcaggtg ggaccaccgc
     1441 gctactgccg ccaggcaaat tctgttttat cagaccgctt ctgcgttctg atttaatctg
     1501 tatcaggctg aaaatcttct ctcatccgcc aaaacagcca agctggagac cgtttaaact
     1561 caatgatgat gatgatgatg gtcgacggcg ctattcagat cctcttctga gatgagtttt
     1621 tgttcgggcc caagcttcag atcccATAtt atattcccca gaacatcagg ttaatggcgt
     1681 ttttgatgtc attttcgcgg tggctgagat cagccacttc ttccccgata acggagaccg
     1741 gcacactggc catatcggtg gtcatcatgc gccagctttc atccccgata tgcaccaccg
     1801 ggtaaagttc acgggagact ttatctgaca gcagacgtgc actggccagg gggatcacca
     1861 tccgtcgccc gggcgtgtca ataatatcac tctgtacatc cacaaacaga cgataacggc
     1921 tctctctttt ataggtgtaa accttaaact gcatAGGTca ttcctcctag atccaaaata
     1981 cggtaccgtc aacaatctca ctcgagagtc gacgtgctag catagtacct aggactgagc
     2041 tagccgtaaa CGTTtGAGAC Gtactagtag cggccgctgc agttaatcac tgattaactc
     2101 ggtaccaaat tccagaaaag aggcctcccg aaaggggggc ctttttcgt tttggtcctt
     2161 ttgttatcaa taaaaaaggg gagcggtttc ccgctcccct tattgttcgt cTACAattat
     2221 tagaagaact cgtcaagaag gcgatagaag gcgatgcgct gcgaatcggg agcggcgata
     2281 ccgtaaagca cgaggaagcg gtcagcccat tcgccgccaa gctcttcagc aatatcacgg
     2341 gtagccaacg ctatgtcctg atagcggtcc gccacaccca gccggccaca gtcgatgaat
     2401 ccagaaaagc ggccattttc caccatgata ttcggcaagc aggcatcgcc atgggtcacg
     2461 acgagatcct cgccgtcggg catgcgcgcc ttgagcctgg cgaacagttc ggctggcgcg
     2521 agcccctgat gctcttcgtc cagatcatcc tgatcgacaa gaccggcttc catccgagta
     2581 cgtgctcgct cgatgcgatg tttcgcttgg tggtcgaatg ggcaggtagc cggatcaagc
     2641 gtatgcagcc gccgcattgc atcagccatg atggatactt tctcggcagg agcaaggtga
     2701 gatgacagga gatcctgccc cggcacttcg cccaatagca gccagtccct tcccgcttca
     2761 gtgacaacgt cgagcacagc tgcgcaagga acgcccgtcg tggccagcca cgatagccgc
     2821 gctgcctcgt cctgcagttc attcagggca ccggacaggt cggtcttgac aaaaagaacc
     2881 gggcgcccct gcgctgacag ccggaacacg gcggcatcag agcagccgat tgtctgttgt
     2941 gcccagtcat agccgaatag cctctccacc caagcggccg gagaacctgc gtgcaatcca
     3001 tcttgttcaa tcatgcgaaa cgatcctcat cctgtctctt gatcagatct tgatcccctg
     3061 cgccatcaga tccttggcgg caagaaagcc atccagttta ctttgcaggg cttcccaacc
     3121 ttaccagagg gcgccccagc tggcaattcc ggttcgcttg ctgtccataa GAGTcctgtt
     3181 gagtaatagt caaaagcctc cggtcggagg cttttgactt tctgcttacC CAAttactac
     3241 cggcgcggca gcgtgacccg tgtcggcggc tccaacggct cgccatcgtc cagaaaacac
     3301 ggctcatcgg gcatcggcag gcgctgctgc ccgcgccgtt cccattcctc cgtttcggtc
```

```
     3361 aaggctggca ggtctggttc catgcccgga atgccgggct ggctgggcgg ctcctcgccg
     3421 gggccggtcg gtagttgctg ctcgcccgga tacagggtcg ggatgcggcg caggtcgcca
     3481 tgccccaaca gcgattcgtc ctggtcgtcg tgatcaacca ccacggcggc actgaacacc
     3541 gacaggcgca actggtcgcg gggctggccc cacgccacgc ggtcattgac cacgtaggcc
     3601 gacacggtgc cggggccgtt gagcttcacg acggagatcc agcgctcggc caccaagtcc
     3661 ttgactgcgt attggaccgt ccgcaaagaa cgtccgatga gcttggaaag tgttttctgg
     3721 ctgaccacca cggcgttctg gtggcccatc tgcgccacga ggtgatgcag cagcattgcc
     3781 gccgtgggtt tcctcgcaat aagcccggcc cacgcctcat gcgctttgcg ttccgtttgc
     3841 acccagtgac cgggcttgtt cttggcttga atgccgattt ctctggactg cgtggccatg
     3901 cttatctcca tgcggtaggg gtgccgcacg gttgcggcac catgcgcaat cagctgcaac
     3961 ttttcggcag cgcgacaaca attatgcgtt gcgtaaaagt ggcagtcaat tacagatttt
     4021 ctttaaccta cgcaatgagc tattgcgggg ggtgccgcaa tgagctgttg cgtacccccc
     4081 ttttttaagt tgttgatttt taagtctttc gcatttcgcc ctatatctag ttctttggtg
     4141 cccaaagaag ggcacccctg cggggttccc ccacgccttc ggcgcggctc cccctccggc
     4201 aaaaagtggc ccctccgggg cttgttgatc gactgcgcgg ccttcggcct tgcccaaggt
     4261 ggcgctgccc ccttggaacc cccgcactcg ccgccgtgag gctcgggggg caggcgggcg
     4321 ggcttcgcct tcgactgccc ccactcgcat aggcttgggt cgttccaggc gcgtcaaggc
     4381 caagccgctg cgcggtcgct gcgcgagcct tgacccgcct tccacttggt gtccaaccgg
     4441 caagcgaagc gcgcaggccg caggccggag gcttttcccc agagaaaatt aaaaaaattg
     4501 atggggcaag gccgcaggcc gcgcagttgg agccggtggg tatgtggtcg aaggctgggt
     4561 agccggtggg caatccctgt ggtcaagctc gtgggcaggc gcagcctgtc catcagcttg
     4621 tccagcaggg ttgtccacgg gccgagcgaa gcgagccagc cggtggccgc tcgcggccat
     4681 cgtccacata tccacgggct ggcaagggag cgcagcgacc gcgcagggcg aagcccggag
     4741 agcaagcccg tagggcgccg cagccgccgt aggcggtcac gactttgcga agcaaagtct
     4801 agtgagtata ctcaagcatt gagtggcccg ccggaggcac cgccttgcgc tgcccccgtc
     4861 gagccggttg gacaccaaaa gggaggggca ggcatggcgg catacgcgat catgcgatgc
     4921 aagaagctgg cgaaaatggg caacgtggcg gccagtctca agcacgccta ccgcgagcgc
     4981 gagactccca acgctgacgc cagcaggacg ccagagaacg agcactgggc ggccagcagc
     5041 accgatgaag cgatgggccg actgcgcgag ttgctgccag agaagcggcg caaggacgct
     5101 gtgttggcgg tcgagtacgt catgacggcc agcccggaat ggtggaagtc ggccagccaa
     5161 gaacagcagg cggcgttctt cgagaaggcg cacaagtggc tggcggacaa gtacggggcg
     5221 gatcgcatcg tgacggccag catccaccgt gacgaaacca gcccgcacat gaccgcgttc
     5281 gtggtgccgc tgacgcagga cggcaggctg tcggccaagg agttcatcgg caacaaagcg
     5341 cagatgaccc gcgaccagac cacgtttgcg gcAgctgtgg ccgatctagg gctgcaacgg
     5401 ggcatcgagg gcagcaaggc acgtcacacg cgcattcagg cgttctacga ggccctggag
     5461 cggccaccag tgggccacgt caccatcagc ccgcaagcgg tcgagccacg cgcctatgca
     5521 ccgcagggat tggccgaaaa gctgggaatc tcaaagcgcg ttgaAacgcc ggaagccgtg
     5581 gccgaccggc tgacaaaagc ggttcggcag ggtatgagc ctgccctaca ggccgccgca
     5641 ggagcgcgtg agatgcgcaa gaaggccgat caagcccaag agacagcccg agaTcttcgg
     5701 gagcgcctga agcccgttct ggacgccctg ggccgttga atcgggatat gcaggccaag
     5761 gccgccgcga tcatcaaggc cgtgggcgaa aagctgctga cggaacagcg ggaagtccag
     5821 cgccagaaac aggcccagcg ccagcaggaa cgcgggcgcg cacatttccc cgaaaagtgc
     5881 cacctgggct gataaACGG
//
```

**pMAS048**

```
LOCUS       pMAS048                 3211 bp ds-DNA     circular     10-OCT-
2025
DEFINITION  .
FEATURES             Location/Qualifiers
     promoter        131..158
                     /label="RNAIp (predicted)"
                     /ApEinfo_revcolor="#c6c9d1"
                     /ApEinfo_fwdcolor="#c6c9d1"
     misc_feature    167..270
                     /label="RNAI (predicted)"
                     /ApEinfo_revcolor="#c6c9d1"
                     /ApEinfo_fwdcolor="#c6c9d1"
     promoter        complement(284..310)
                     /label="RNAIIp (predicted)"
                     /ApEinfo_revcolor="#c6c9d1"
                     /ApEinfo_fwdcolor="#c6c9d1"
     terminator      332..370
                     /label="bidir. modified thrLt BBa_B1006 "
                     /ApEinfo_revcolor="#c6c9d1"
                     /ApEinfo_fwdcolor="#c6c9d1"
     misc_feature    332..487
                     /label="ConL'S"
                     /ApEinfo_revcolor="#c6c9d1"
                     /ApEinfo_fwdcolor="#c6c9d1"
     terminator      371..419
                     /label="tVoigtS5, L3S3P22 47C>G"
                     /ApEinfo_revcolor="#c6c9d1"
                     /ApEinfo_fwdcolor="#c6c9d1"
     primer          395..419
                     /label="AB17 ConL F seq"
                     /note="sequence: ggcctttttttgtttctggtctgcc"
                     /ApEinfo_revcolor="#75c6a9"
                     /ApEinfo_fwdcolor="#75c6a9"
     misc_feature    420..434
                     /label="6-frame STOP a"
                     /ApEinfo_revcolor="#c6c9d1"
                     /ApEinfo_fwdcolor="#c6c9d1"
     misc_feature    435..440
                     /label="EcoRI"
                     /ApEinfo_revcolor="#d59687"
                     /ApEinfo_fwdcolor="#d59687"
     misc_feature    complement(435..456)
                     /label="BioBrick Prefix BBa_G00000"
                     /ApEinfo_revcolor="#d59687"
                     /ApEinfo_fwdcolor="#d59687"
     misc_feature    441..448
                     /label="NotI"
                     /ApEinfo_revcolor="#b7e6d7"
                     /ApEinfo_fwdcolor="#b7e6d7"
     misc_feature    450..455
                     /label="XbaI"
                     /ApEinfo_revcolor="#d59687"
                     /ApEinfo_fwdcolor="#d59687"
     misc_feature    457..462
```

```
                     /label="BsaI"
                     /ApEinfo_revcolor="#b1ff67"
                     /ApEinfo_fwdcolor="#b1ff67"
     misc_feature    468..473
                     /label="XhoI"
                     /ApEinfo_revcolor="#c7b0e3"
                     /ApEinfo_fwdcolor="#c7b0e3"
     misc_feature    474..481
                     /label="AscI"
                     /ApEinfo_revcolor="#ffef86"
                     /ApEinfo_fwdcolor="#ffef86"
     misc_feature    480..487
                     /label="SbfI"
                     /ApEinfo_revcolor="#85dae9"
                     /ApEinfo_fwdcolor="#85dae9"
     misc_feature    492..497
                     /label="BamHI"
                     /ApEinfo_revcolor="#c7b0e3"
                     /ApEinfo_fwdcolor="#c7b0e3"
     promoter        518..563
                     /label="P.lacIq (Piq)"
                     /ApEinfo_revcolor="#ffef86"
                     /ApEinfo_fwdcolor="#ffef86"
     misc_feature    528..533
                     /label=-35
                     /ApEinfo_revcolor="#ffef86"
                     /ApEinfo_fwdcolor="#ffef86"
     misc_feature    551..556
                     /label=-10
                     /ApEinfo_revcolor="#ffef86"
                     /ApEinfo_fwdcolor="#ffef86"
     RBS             564..591
                     /label="cym1"
                     /ApEinfo_revcolor="#f8d3a9"
                     /ApEinfo_fwdcolor="#f8d3a9"
     CDS             592..1206
                     /label="cymR.AM"
                     /ApEinfo_revcolor="#c7b0e3"
                     /ApEinfo_fwdcolor="#c7b0e3"
     CDS             592..1206
                     /ApEinfo_revcolor="#c7b0e3"
                     /ApEinfo_fwdcolor="#c7b0e3"

/translation="MSPKRRTQAERAMETQGKLIAAALGVLREKGYAGFRIADVPGAAGVSRGAQSHHFPTKLELL
LATFEWLYEQITERSRARLAKLKPEDDVIQQMLDDAAEFFLDDDFSIGLDLIVAADRDPALREGIQRTVERNRFVV
EDMWLGVLVSRGLSRDDAEDILWLIFNSVRGLVVRSLWQKDKERFERVRNSTLEIARERYAKFKR**"
     misc_feature    919..921
                     /label="S110G"
                     /ApEinfo_revcolor="#c7b0e3"
                     /ApEinfo_fwdcolor="#c7b0e3"
     misc_feature    1102..1104
                     /label="A171V"
                     /ApEinfo_revcolor="#c7b0e3"
                     /ApEinfo_fwdcolor="#c7b0e3"
     misc_feature    1219..1225
                     /label="3' cleaved"
                     /ApEinfo_revcolor="#f8d3a9"
```

```
                     /ApEinfo_fwdcolor="#f8d3a9"
     misc_feature    1219..1293
                     /label="RiboJ00"
                     /ApEinfo_revcolor="#f8d3a9"
                     /ApEinfo_fwdcolor="#f8d3a9"
     RBS             1298..1322
                     /label="RBS, TIR=919k ±RiboJ"
                     /ApEinfo_revcolor="#f8d3a9"
                     /ApEinfo_fwdcolor="#f8d3a9"
     primer          complement(1302..1339)
                     /label="MAS55B"
                     /note="sequence:
gtaatacgctgcttcatTaaacctccttaactcgcgtg"
                     /ApEinfo_revcolor="#9eafd2"
                     /ApEinfo_fwdcolor="#9eafd2"
     primer          1307..1347
                     /label="MAS56B"
                     /note="sequence:
gagttaaggaggtttAatgaagcagcgtattacagtgacag"
                     /ApEinfo_revcolor="#b1ff67"
                     /ApEinfo_fwdcolor="#b1ff67"
     gene            1323..1541
                     /label="ccdA gene"
                     /ApEinfo_revcolor="#f8d3a9"
                     /ApEinfo_fwdcolor="#f8d3a9"
     CDS             1323..1541
                     /label="ccdA CDS (type II toxin-antitoxin system
antitoxin CcdA)"
                     /ApEinfo_revcolor="#b4abac"
                     /ApEinfo_fwdcolor="#b4abac"
     CDS             1323..1541
                     /label="ccdA CDS"
                     /ApEinfo_revcolor="#84b0dc"
                     /ApEinfo_fwdcolor="#84b0dc"

/translation="MKQRITVTVDSDSYQLLKAYDVNISGLVSTTMQNEARRLRAERWKAENQEGMAEVARFIEMN
GSFADENRDW*"
     primer          complement(1523..1564)
                     /label="MAS57B"
                     /note="sequence:
cttaagtttttggctgaaGCCAtcaccagtccctgttctcg"
                     /ApEinfo_revcolor="#b1ff67"
                     /ApEinfo_fwdcolor="#b1ff67"
     primer          1526..1564
                     /label="MAS54B"
                     /note="sequence:
gaacagggactggtgaTGGCttcagccaaaaaacttaag"
                     /ApEinfo_revcolor="#9eafd2"
                     /ApEinfo_fwdcolor="#9eafd2"
     terminator      1546..1635
                     /label="tVoigtN1 spyt"
                     /ApEinfo_revcolor="#f58a5e"
                     /ApEinfo_fwdcolor="#f58a5e"
     misc_feature    1644..1649
                     /label="SphI"
                     /ApEinfo_revcolor="#c7b0e3"
                     /ApEinfo_fwdcolor="#c7b0e3"
```

```
     misc_feature    1650..1657
                     /label="AscI"
                     /ApEinfo_revcolor="#ffef86"
                     /ApEinfo_fwdcolor="#ffef86"
     misc_feature    complement(1663..1668)
                     /label="BsaI"
                     /ApEinfo_revcolor="#b1ff67"
                     /ApEinfo_fwdcolor="#b1ff67"
     misc_feature    1667..1674
                     /label="SbfI"
                     /ApEinfo_revcolor="#85dae9"
                     /ApEinfo_fwdcolor="#85dae9"
     misc_feature    complement(1675..1695)
                     /label="BioBrick Suffix BBa_G00001"
                     /ApEinfo_revcolor="#d59687"
                     /ApEinfo_fwdcolor="#d59687"
     misc_feature    1676..1681
                     /label="SpeI"
                     /ApEinfo_revcolor="#d59687"
                     /ApEinfo_fwdcolor="#d59687"
     misc_feature    1683..1690
                     /label="NotI"
                     /ApEinfo_revcolor="#b7e6d7"
                     /ApEinfo_fwdcolor="#b7e6d7"
     misc_feature    1690..1695
                     /label="PstI"
                     /ApEinfo_revcolor="#d59687"
                     /ApEinfo_fwdcolor="#d59687"
     misc_feature    1696..1710
                     /label="6-frame STOP b"
                     /ApEinfo_revcolor="#c6c9d1"
                     /ApEinfo_fwdcolor="#c6c9d1"
     terminator      1711..1771
                     /label="tVoigtS1, L3S2P21"
                     /ApEinfo_revcolor="#c6c9d1"
                     /ApEinfo_fwdcolor="#c6c9d1"
     terminator      complement(1772..1824)
                     /label="tVoigtS19, L3S1P00"
                     /ApEinfo_revcolor="#c6c9d1"
                     /ApEinfo_fwdcolor="#c6c9d1"
     terminator      complement(1829..1884)
                     /label="λ t0"
                     /ApEinfo_revcolor="#82ca9d"
                     /ApEinfo_fwdcolor="#82ca9d"
     gene            complement(1829..2649)
                     /label="ChlR, chloramphenicol resistance (Tn9)"
                     /ApEinfo_revcolor="#82ca9d"
                     /ApEinfo_fwdcolor="#82ca9d"
     CDS             complement(1885..2544)
                     /label="cat, chloramphenicol O-acetyltransferase A-1
(ChlR)"
                     /ApEinfo_revcolor="#82ca9d"
                     /ApEinfo_fwdcolor="#82ca9d"
     RBS             complement(2545..2557)
                     /label="cat RBS"
                     /ApEinfo_revcolor="#82ca9d"
                     /ApEinfo_fwdcolor="#82ca9d"
```

```
     misc_feature    complement(2545..2649)
                     /label="catp, ChlR promoter / 5' UTR"
                     /ApEinfo_revcolor="#82ca9d"
                     /ApEinfo_fwdcolor="#82ca9d"
     misc_feature    complement(2583..2591)
                     /label="catp -10 box"
                     /ApEinfo_revcolor="#82ca9d"
                     /ApEinfo_fwdcolor="#82ca9d"
     misc_feature    complement(2607..2612)
                     /label="catp -35 box"
                     /ApEinfo_revcolor="#82ca9d"
                     /ApEinfo_fwdcolor="#82ca9d"
     terminator      2654..2708
                     /label="tVoigtN17, tonBt (bidir), BBa_B0054"
                     /ApEinfo_revcolor="#f58a5e"
                     /ApEinfo_fwdcolor="#f58a5e"
     misc_feature    complement(2713..327)
                     /label="p15A ori"
                     /ApEinfo_revcolor="#b4abac"
                     /ApEinfo_fwdcolor="#b4abac"
     misc_feature    complement(2976..273)
                     /label="RNAII (predicted)"
                     /ApEinfo_revcolor="#c6c9d1"
                     /ApEinfo_fwdcolor="#c6c9d1"
     misc_feature    complement(2977..2977)
                     /label="replication origin (predicted)"
                     /ApEinfo_revcolor="#ff9ccd"
                     /ApEinfo_fwdcolor="#ff9ccd"
ORIGIN
        1 atgcacgaac cccccgttca gtccgaccgc tgcgccttat ccggtaacta tcgtcttgag
       61 tccaacccgg aaagacatgc aaaagcacca ctggcagcag ccactggtaa ttgatttaga
      121 ggagttagtc ttgaagtcat gcgccggtta aggctaaact gaaaggacaa gttttggtga
      181 ctgcgctcct ccaagccagt tacctcggtt caaagagttg gtagctcaga gaaccttcga
      241 aaaaccgccc tgcaaggcgg ttttttcgtt ttcagagcaa gagattacgc gcagaccaaa
      301 acgatctcaa gaagatcatc ttattaaACG Gaaaaaaaaa ccccgcccct gacagggcgg
      361 ggtttttttt ccaattattg aaggccgcta acgcggcctt tttttgtttc tggtctgcct
      421 taatcaatga ctaagaattc gcggccgctt ctagagGGTC TCaCTTCctc gagggcgcgc
      481 ctgcaggCTT Cggatccgct gcaggCTTCg cggcgcgcca tcgaatggtg caaaaccttt
      541 cgcggtatgg catgatagcg cccggaagag agtcaattca gggtggtgaa tatgagcccg
      601 aaacgtcgta cccaggcaga acgtgcaatg gaaacccagg gtaaactgat tgcagcagca
      661 ctgggtgttc tgcgtgaaaa aggttatgca ggttttcgta ttgcagatgt tccgggtgca
      721 gccggtgtta gccgtggtgc acagagccat cattttccga ccaaactgga actgctgctg
      781 gcaacctttg aatggctgta tgagcagatt accgaacgta gccgtgcacg tctggcaaaa
      841 ctgaaaccgg aagatgatgt tattcagcag atgctggatg atgcagcaga atttttctg
      901 gatgatgatt ttagcatcgg cctggatctg attgttgcag cagatcgtga tccggcactg
      961 cgtgaaggta ttcagcgtac cgttgaacgt aatcgttttg ttgttgaaga tatgtggctg
     1021 ggtgtgctgg tgagccgtgg tctgagccgt gatgatgccg aagatattct gtggctgatt
     1081 tttaacagcg ttcgtggtct ggtagttcgt agcctgtggc agaaagataa agaacgtttt
     1141 gaacgtgtgc gtaatagcac cctggaaatt gcacgtgaac gttatgcaaa attcaaacgt
     1201 tgataaTGGC TTCCCGACag ctgtcaccgg atgtgctttc cggtctgatg agtccgtgag
     1261 gacgaaacag cctctacaaa taattttgtt taaTCAGagg tcacgcgagt taaggaggtt
     1321 tAatgaagca gcgtattaca gtgacagttg acagcgacag ctatcagttg ctcaaggcat
     1381 atgatgtcaa tatctccggt ctggtaagca caaccatgca gaatgaagcc cgtcgtctgc
     1441 gtgccgaacg ctggaaagcg gaaaatcagg aagggatggc tgaggtcgcc cggtttattg
     1501 aaatgaacgg ctcttttgct gacgagaaca gggactggtg aTGGCttcag ccaaaaaact
     1561 taagaccgcc ggtcttgtcc actaccttgc agtaatgcgg tggacaggat cggcggtttt
     1621 cttttctctt ctcaaGCTGG CTGgcatgcg gcgcgccGCT GtGAGACCtg caggtactag
```

```
     1681 tagcggccgc tgcagttaat cactgattaa ctcggtacca aattccagaa aagaggcctc
     1741 ccgaaagggg ggcctttttt cgttttggtc cttttgttat caataaaaaa ggggagcggt
     1801 ttcccgctcc ccttattgtt cgtcTACAac caataaaaaa cgcccggcgg caaccgagcg
     1861 ttctgaacaa atccagatgg agtattacgc cccgccctgc cactcatcgc agtactgttg
     1921 taattcatta agcattctgc cgacatggaa gccatcacaa acggcatgat gaacctgaat
     1981 cgccagcggc atcagcacct tgtcgccttg cgtataatat ttgcccatgg tgaaaacggg
     2041 ggcgaagaag ttgtccatat tggccacgtt taaatcaaaa ctggtgaaac tcacccaggg
     2101 attggctgaa acgaaaaaca tattctcaat aaacccttta gggaaatagg ccaggttttc
     2161 accgtaacac gccacatctt gcgaatatat gtgtagaaac tgccggaaat cgtcgtggta
     2221 ttcactccag agcgatgaaa acgtttcagt ttgctcatgg aaaacggtgt aacaagggtg
     2281 aacactatcc catatcacca gctcaccgtc tttcattgcc atacgaaatt ccggatgagc
     2341 attcatcagg cgggcaagaa tgtgaataaa ggccggataa aacttgtgct tatttttctt
     2401 tacggtcttt aaaaaggccg taatatccag ctgaacggtc tggttatagg tacattgagc
     2461 aactgactga aatgcctcaa aatgttcttt acgatgccat tgggatatat caacggtggt
     2521 atatccagtg attttttct ccattttagc ttccttagct cctgaaaatc tcgataactc
     2581 aaaaaatacg cccggtagtg atcttatttc attatggtga aagttggaac ctcttacgtg
     2641 cccgatcaaG AGTcctgttg agtaatagtc aaaagcctcc ggtcggaggc ttttgacttt
     2701 ctgcttacCC AAtagcggag tgtatactgg cttactatgt tggcactgat gagggtgtca
     2761 gtgaagtgct tcatgtggca ggagaaaaaa ggctgcaccg gtgcgtcagc agaatatgtg
     2821 atacaggata tattccgctt cctcgctcac tgactcgcta cgctcggtcg ttcgactgcg
     2881 gcgagcggaa atggcttacg aacggggcgg agatttcctg gaagatgcca ggaagatact
     2941 taacagggaa gtgagagggc cgcggcaaag ccgtttttcc ataggctccg cccccctgac
     3001 aagcatcacg aaatctgacg ctcaaatcag tggtggcgaa acccgacagg actataaaga
     3061 taccaggcgt ttccccctgg cggctccctc gtgcgctctc ctgttcctgc ctttcggttt
     3121 accggtgtca ttccgctgtt atggccgcgt ttgtctcatt ccacgcctga cactcagttc
     3181 cgggtaggca gttcgctcca agctggactg t
//
```

**pMAS047**

```
LOCUS       pMAS047                12562 bp ds-DNA     circular     22-MAY-
2025
DEFINITION  .
FEATURES             Location/Qualifiers
     rep_origin      4..744
                     /label="oriR101"
                     /ApEinfo_revcolor="#ff9ccd"
                     /ApEinfo_fwdcolor="#ff9ccd"
     misc_feature    745..842
                     /label="CmR Promoter"
                     /ApEinfo_revcolor="#b4abac"
                     /ApEinfo_fwdcolor="#b4abac"
     misc_feature    745..1608
                     /label="CmR"
                     /ApEinfo_revcolor="#faac61"
                     /ApEinfo_fwdcolor="#faac61"
     CDS             843..1502
                     /label="CmR CDS"
                     /ApEinfo_revcolor="#faac61"
                     /ApEinfo_fwdcolor="#faac61"
     primer          1468..1487
                     /label="DSZ26A"
                     /note="sequence: AACAGTACTGCGATGAGTGG"
                     /ApEinfo_revcolor="#9eafd2"
                     /ApEinfo_fwdcolor="#9eafd2"
     misc_feature    1536..1541
                     /label="BamHI-BglII Scar"
                     /ApEinfo_revcolor="#ac1dff"
                     /ApEinfo_fwdcolor="#ac1dff"
     misc_feature    1609..1643
                     /label="J23119 Promoter"
                     /ApEinfo_revcolor="#b4abac"
                     /ApEinfo_fwdcolor="#b4abac"
                     /note="ApEinfo_revcolor: #b4abac"
     misc_feature    1667..1694
                     /label="CRISPR repeat (VCHE45 Tn6677)"
                     /ApEinfo_revcolor="#f58a5e"
                     /ApEinfo_fwdcolor="#f58a5e"
     misc_feature    1695..1726
                     /label="bla CDS 5 guide"
                     /ApEinfo_revcolor="#c7b0e3"
                     /ApEinfo_fwdcolor="#c7b0e3"
     misc_feature    1727..1754
                     /label="CRISPR repeat (VCHE45 Tn6677)"
                     /ApEinfo_revcolor="#f58a5e"
                     /ApEinfo_fwdcolor="#f58a5e"
     RBS             1772..1777
                     /label="RBS"
                     /ApEinfo_revcolor="#b4abac"
                     /ApEinfo_fwdcolor="#b4abac"
                     /note="ApEinfo_revcolor: #b4abac"
     CDS             1786..2970
                     /label="tniQ (pQCascade)"
                     /ApEinfo_revcolor="#85dae9"
```

```
                     /ApEinfo_fwdcolor="#85dae9"
     CDS             2971..4893
                     /label="cas8-5 fusion (pQCascade)"
                     /ApEinfo_revcolor="#bc8dbf"
                     /ApEinfo_fwdcolor="#bc8dbf"
     CDS             4865..5923
                     /label="cas7/csy3 (pQCascade)"
                     /ApEinfo_revcolor="#f8d3a9"
                     /ApEinfo_fwdcolor="#f8d3a9"
     CDS             5926..6525
                     /label="cas6f (pQCascade)"
                     /ApEinfo_revcolor="#c7b0e3"
                     /ApEinfo_fwdcolor="#c7b0e3"
     RBS             6534..6539
                     /label="RBS"
                     /ApEinfo_revcolor="#b4abac"
                     /ApEinfo_fwdcolor="#b4abac"
                     /note="ApEinfo_revcolor: #b4abac"
     CDS             6547..7239
                     /label="tnsA (pTnsABC)"
                     /ApEinfo_revcolor="#f58a5e"
                     /ApEinfo_fwdcolor="#f58a5e"
     CDS             7232..9043
                     /label="tnsB (pTnsABC)"
                     /ApEinfo_revcolor="#ffef86"
                     /ApEinfo_fwdcolor="#ffef86"
     CDS             9055..10047
                     /label="tnsC (pTnsABC)"
                     /ApEinfo_revcolor="#75c6a9"
                     /ApEinfo_fwdcolor="#75c6a9"
     terminator      10085..10132
                     /label="T7 terminator"
                     /ApEinfo_revcolor="#c6c9d1"
                     /ApEinfo_fwdcolor="#c6c9d1"
                     /note="ApEinfo_revcolor: #c6c9d1"
     misc_feature    10153..10160
                     /label="END"
                     /ApEinfo_revcolor="#84b0dc"
                     /ApEinfo_fwdcolor="#84b0dc"
                     /note="ApEinfo_revcolor: #84b0dc"
     misc_feature    10153..10277
                     /label="VchINT R-end"
                     /ApEinfo_revcolor="#ffef86"
                     /ApEinfo_fwdcolor="#ffef86"
                     /note="ApEinfo_revcolor: #ffef86"
     misc_feature    10161..10180
                     /label="R1 tnsB binding site (VCHE45 Tn6677)"
                     /ApEinfo_revcolor="#c7b0e3"
                     /ApEinfo_fwdcolor="#c7b0e3"
     misc_feature    10161..10263
                     /label="RE (partial) (VCHE45 Tn6677)"
                     /ApEinfo_revcolor="#c6c9d1"
                     /ApEinfo_fwdcolor="#c6c9d1"
     misc_feature    10181..10200
                     /label="R2 tnsB binding site (VCHE45 Tn6677)"
                     /ApEinfo_revcolor="#c7b0e3"
                     /ApEinfo_fwdcolor="#c7b0e3"
```

```
     misc_feature    10204..10223
                     /label="R3 tnsB binding site (VCHE45 Tn6677)"
                     /ApEinfo_revcolor="#c7b0e3"
                     /ApEinfo_fwdcolor="#c7b0e3"
     primer          10263..10300
                     /label="DSZ79C"
                     /note="sequence:
aatgaagactatttcCCAATTATTGAAGGCCGCTAACG"
                     /ApEinfo_revcolor="#b4abac"
                     /ApEinfo_fwdcolor="#b4abac"
     primer          10264..10300
                     /label="DSZ11E"
                     /note="sequence: aatggtctcaaagcCCAATTATTGAAGGCCGCTAACG"
                     /ApEinfo_revcolor="#ffef86"
                     /ApEinfo_fwdcolor="#ffef86"
     primer          10274..10326
                     /label="DSZ51C"
                     /note="sequence:
tttcCCAATTATTGAAGGCCGCTAACGCGGCCTTTTTTTGTTTCTGGTCTGCC"
                     /ApEinfo_revcolor="#ff9ccd"
                     /ApEinfo_fwdcolor="#ff9ccd"
     terminator      10278..10326
                     /label="tVoigtS5, L3S3P22 47C>G"
                     /ApEinfo_revcolor="#c6c9d1"
                     /ApEinfo_fwdcolor="#c6c9d1"
     primer          complement(10278..10330)
                     /label="DSZ52C"
                     /note="sequence:
aggaGGCAGACCAGAAACAAAAAAAGGCCGCGTTAGCGGCCTTCAATAATTGG"
                     /ApEinfo_revcolor="#9eafd2"
                     /ApEinfo_fwdcolor="#9eafd2"
     primer          10327..10352
                     /label="DSZ76B"
                     /note="sequence: aatgaagacattcctCCCTGCAGGCTTCAACAAACAG"
                     /ApEinfo_revcolor="#84b0dc"
                     /ApEinfo_fwdcolor="#84b0dc"
     misc_feature    10332..10339
                     /label="SbfI"
                     /ApEinfo_revcolor="#85dae9"
                     /ApEinfo_fwdcolor="#85dae9"
     misc_feature    10344..10375
                     /label="CymR operator"
                     /ApEinfo_revcolor="#ffef86"
                     /ApEinfo_fwdcolor="#ffef86"
     promoter        10344..10433
                     /label="P.CymRC (Marionette)"
                     /ApEinfo_revcolor="#ffef86"
                     /ApEinfo_fwdcolor="#ffef86"
     misc_feature    10367..10372
                     /label=-35
                     /ApEinfo_revcolor="#ffef86"
                     /ApEinfo_fwdcolor="#ffef86"
     misc_feature    10390..10395
                     /label=-10
                     /ApEinfo_revcolor="#ffef86"
                     /ApEinfo_fwdcolor="#ffef86"
     misc_feature    10402..10402
```

```
                     /label="TSS"
                     /ApEinfo_revcolor="#ffef86"
                     /ApEinfo_fwdcolor="#ffef86"
     misc_feature    10402..10433
                     /label="CymR operator"
                     /ApEinfo_revcolor="#ffef86"
                     /ApEinfo_fwdcolor="#ffef86"
     misc_feature    10438..10493
                     /label="U64 RNA Guide"
                     /ApEinfo_revcolor="#faac61"
                     /ApEinfo_fwdcolor="#faac61"
     misc_feature    10438..11403
                     /label="U64 Ribozyme transcript"
                     /ApEinfo_revcolor="#d59687"
                     /ApEinfo_fwdcolor="#d59687"
     primer          10475..10493
                     /label="MAS52B"
                     /note="sequence: ggcccgggaacgtgttcac"
                     /ApEinfo_revcolor="#faac61"
                     /ApEinfo_fwdcolor="#faac61"
     misc_feature    10488..10493
                     /label="IGS"
                     /ApEinfo_revcolor="#c7b0e3"
                     /ApEinfo_fwdcolor="#c7b0e3"
     misc_feature    10494..10880
                     /label="Group I Intron Ribozyme (from p-OiRS3GG)"
                     /ApEinfo_revcolor="#b4abac"
                     /ApEinfo_fwdcolor="#b4abac"
     primer          complement(10874..10901)
                     /label="MAS51B"
                     /note="sequence: GGGAAAAGCATTGAACACCATCGAGTAC"
                     /ApEinfo_revcolor="#b1ff67"
                     /ApEinfo_fwdcolor="#b1ff67"
     misc_feature    10881..11403
                     /label="Barcode (sfGFP_2)"
                     /ApEinfo_revcolor="#84b0dc"
                     /ApEinfo_fwdcolor="#84b0dc"
     misc_feature    10902..10906
                     /label="Internal Barcode"
                     /ApEinfo_revcolor="#b1ff67"
                     /ApEinfo_fwdcolor="#b1ff67"
     primer          10902..10929
                     /label="MAS50B"
                     /note="sequence: GAGTGGTTATCCGGATCACATGAAACGG"
                     /ApEinfo_revcolor="#b1ff67"
                     /ApEinfo_fwdcolor="#b1ff67"
     primer          complement(10902..10929)
                     /label="MAS53B"
                     /note="sequence: ccgtttcatgtgatccggataacCACTC"
                     /ApEinfo_revcolor="#faac61"
                     /ApEinfo_fwdcolor="#faac61"
     misc_feature    11316..11321
                     /label="BamHI-BglII Scar"
                     /ApEinfo_revcolor="#ac1dff"
                     /ApEinfo_fwdcolor="#ac1dff"
     terminator      11408..11460
                     /label="tVoigtS4 L3S3P21 51C>G"
```

```
                     /ApEinfo_revcolor="#f58a5e"
                     /ApEinfo_fwdcolor="#f58a5e"
     primer          complement(11447..11473)
                     /label="DSZ40A"
                     /note="sequence: aatgaagacattaagCGCAGCGGCAGACCAGAAAC"
                     /ApEinfo_revcolor="#85dae9"
                     /ApEinfo_fwdcolor="#85dae9"
     primer          complement(11459..11490)
                     /label="DSZ12E"
                     /note="sequence: aatggtctcattcgCTGCAGTAAGCGCAGCGG"
                     /ApEinfo_revcolor="#84b0dc"
                     /ApEinfo_fwdcolor="#84b0dc"
     primer          complement(11459..11491)
                     /label="DSZ80C"
                     /note="sequence: aatgaagactaaggaCTGCAGTAAGCGCAGCGG"
                     /ApEinfo_revcolor="#d6b295"
                     /ApEinfo_fwdcolor="#d6b295"
     misc_feature    complement(11467..11611)
                     /label="VchINT L-end"
                     /ApEinfo_revcolor="#ffef86"
                     /ApEinfo_fwdcolor="#ffef86"
                     /note="ApEinfo_revcolor: #ffef86"
     misc_feature    complement(11467..11611)
                     /label="LE (partial) (VCHE45 Tn6677)"
                     /ApEinfo_revcolor="#b4abac"
                     /ApEinfo_fwdcolor="#b4abac"
     misc_feature    complement(11504..11523)
                     /label="L3 tnsB binding site (VCHE45 Tn6677)"
                     /ApEinfo_revcolor="#f58a5e"
                     /ApEinfo_fwdcolor="#f58a5e"
     misc_feature    complement(11530..11549)
                     /label="L2 tnsB binding site (VCHE45 Tn6677)"
                     /ApEinfo_revcolor="#f58a5e"
                     /ApEinfo_fwdcolor="#f58a5e"
     misc_feature    complement(11584..11603)
                     /label="L1 tnsB binding site (VCHE45 Tn6677)"
                     /ApEinfo_revcolor="#f58a5e"
                     /ApEinfo_fwdcolor="#f58a5e"
     misc_feature    complement(11604..11611)
                     /label="END"
                     /ApEinfo_revcolor="#84b0dc"
                     /ApEinfo_fwdcolor="#84b0dc"
                     /note="ApEinfo_revcolor: #84b0dc"
     CDS             complement(11612..12562)
                     /label="repA101 protein"
                     /ApEinfo_revcolor="#ff9ccd"
                     /ApEinfo_fwdcolor="#ff9ccd"
     CDS             complement(11612..12562)
                     /label="repA101"
                     /ApEinfo_revcolor="#84b0dc"
                     /ApEinfo_fwdcolor="#84b0dc"

/translation="MSELVVFKANELAISRYDLTEHETKLILCCVALLNPTIENPTRKERTVSFTYNQYAQMMNIS
RENAYGVLAKATRELMTRTVEIRNPLVKGFEIFQWTNYAKFSSEKLELVFSEEILPYLFQLKKFIKYNLEHVKSFE
NKYSMRIYEWLLKELTQKKTHKANIEISLDEFKFMLMLENNYHEFKRLNQWVLKPISKDLNTYSNMKLVVDKRGRP
TDTLIFQVELDRQMDLVTELENNQIKMNGDKIPTTITSDSYLRNGLRKTLHDALTAKIQLTSFEAKFLSDMQSKYD
LNGSFSWLTQKQRTTLENILAKYGRI*"
```

```
ORIGIN
        1 acatctcaat tggtctaggt gattttaatc actataccaa ttgagatggg ctagtcaatg
       61 ataattacta gtccttttcc tttgagttgt gggtatctgt aaattctgct agacctttgc
      121 tggaaaactt gtaaattctg ctagaccctc tgtaaattcc gctagacctt tgtgtgtttt
      181 ttttgtttat attcaagtgg ttataattta tagaataaag aaagaataaa aaaagataaa
      241 aagaatagat cccagccctg tgtataactc actactttag tcagttccgc agtattacaa
      301 aaggatgtcg caaacgctgt ttgctcctct acaaaacaga ccttaaaacc ctaaaggctt
      361 aagtagcacc ctcgcaagct cggttgcggc cgcaatcggg caaatcgctg aatattcctt
      421 ttgtctccga ccatcaggca cctgagtcgc tgtctttttc gtgacattca gttcgctgcg
      481 ctcacggctc tggcagtgaa tggggtaaa tggcactaca ggcgcctttt atggattcat
      541 gcaaggaaac tacccataat acaagaaaag cccgtcacgg gcttctcagg gcgttttatg
      601 gcgggtctgc tatgtggtgc tatctgactt tttgctgttc agcagttcct gccctctgat
      661 tttccagtct gaccacttcg gattatcccg tgacaggtca ttcagactgg ctaatgcacc
      721 cagtaaggca gcggtatcat caacggcacg taagaggttc caactttcac cataatgaaa
      781 taagatcact accgggcgta ttttttgagt tatcgagatt ttcaggagct aaggaagcta
      841 aaatggagaa aaaaatcact ggatatacca ccgttgatat atcccaatgg catcgtaaag
      901 aacattttga ggcatttcag tcagttgctc aatgtaccta taaccagacc gttcagctgg
      961 atattacggc ctttttaaag accgtaaaga aaaataagca caagttttat ccggccttta
     1021 ttcacattct tgcccgcctg atgaatgctc atccggaatt tcgtatggca atgaaagacg
     1081 gtgagctggt gatatgggat agtgttcacc cttgttacac cgttttccat gagcaaactg
     1141 aaacgttttc atcgctctgg agtgaatacc acgacgattt ccggcagttt ctacacatat
     1201 attcgcaaga tgtggcgtgt tacggtgaaa acctggccta tttccctaaa gggtttattg
     1261 agaatatgtt tttcgtctca gccaatccct gggtgagttt caccagtttt gatttaaacg
     1321 tggccaatat ggacaacttc ttcgcccccg ttttcaccat gggcaaatat tatacgcaag
     1381 gcgacaaggt gctgatgccg ctggcgattc aggttcatca tgccgtttgt gatggcttcc
     1441 atgtcggcag aatgcttaat gaattacaac agtactgcga tgagtggcag ggcggggcgt
     1501 aatttgatat cgagctcgct tggactcctg ttgatagatc cagtaatgac ctcagaactc
     1561 catctggatt tgttcagaac gctcggttgc cgccgggcgt ttttttattt gacagctagc
     1621 tcagtcctag gtataatact agcgtcgacG TGGAGATATA CCATGGGTGA ACTGCCGAGT
     1681 AGGTAGCTGA TAACAAGTCA TTCTGAGAAT AGTGTATGCG GCGACCGTGA ACTGCCGAGT
     1741 AGGTAGCTGA TAACGGATCC GAATTCGAGC GAAGGAGATA TACATATGTT TTTGCAAAGA
     1801 CCTAAACCTT ACAGCGATGA AAGtTTAGAA AGtTTCTTTA TCCGAGTGGC TAACAAAAAT
     1861 GGCTACGGTG ATGTCCATCG CTTCCTAGAA GCCACTAAAC GATTCCTTCA AGACATTGAC
     1921 CATAATGGCT ATCAAACCTT TCCGACTGAT ATAACTCGGA TAAACCCATA CTCAGCTAAA
     1981 AACAGTTCCA GCGCACGAAC TGCGTCATTC CTGAAaCTTG CACAATTGAC ATTTAATGAA
     2041 CCGCCAGAGC TACTTGGGTT GGCAATTAAC AGAACAAACA TGAAATACTC GCCGTCAACT
     2101 AGCGCGGTTG TTAGAGGTGC AGAAGTCTTT CCTCGCAGTT TACTACGGAC GCACTCCATC
     2161 CCCTGCTGTC CTTTGTGTCT GCGAGAAAAT GGCTACGCCT CCTACCTTTG GCACTTTCAG
     2221 GGGTACGAAT ACTGCCACAG CCATAACGTA CCTTTAATTA CCACTTGTAG CTGTGGTAAG
     2281 GAGTTTGACT ACCGAGTATC TGGGTTAAAG GGCATTTGCT GCAAATGCAA GGAGCCTATC
     2341 ACCTTAACCA GCAGGGAGAA CGGTCATGAG GCAGCGTGTA CTGTTTCAAA CTGGCTTGCT
     2401 GGCCATGAAT CTAAACCTCT GCCAAATCTT CCTAAAAGCT ACCGATGGGG TTTAGTTCAT
     2461 TGGTGGATGG GTATTAAAGA TAGCGAgTTC GATCACTTTT CGTTCGTTCA ATTTTTCTCA
     2521 AACTGGCCAA GGTCATTCCA CTCGATAATC GAAGATGAAG TAGAGTTCAA CCTTGAGCAT
     2581 GCTGTTGTCA GCACGTCTGA ATTACGACTA AAAGATCTTC TTGGTCGATT GTTTTTCGGT
     2641 TCAATTCGGT TACCTGAGCG GAATCTTCAA CACAATATCA TCCTTGGTGA GCTTCTCTGC
     2701 TATTTAGAAA ATCGTTTATG GCAAGACAAG GGATTAATCG CCAACCTCAA AATGAACGCG
     2761 TTAGAGGCGA CTGTAATGTT AAATTGTAGC CTCGATCAGA TTGCATCAAT GGTTGAACAA
     2821 CGCATCTTGA AGCCAAATCG AAAAAGCAAG CCCAACAGCC CTCTTGATGT TACCGATTAT
     2881 CTATTTCATT TCGGCGATAT TTTCTGTCTT TGGTTAGCTG AgTTCCAAAG CGATGAGTTT
     2941 AACCGTTCGT TTTATGTGTC GAGGTGGTAA ATGCAAACTC TGAAAGAACT AATCGCATCC
     3001 AATCCTGACG ACTTAACAAC TGAACTTAAG AGAGCATTTC GTCCACTCAC ACCGCATATT
     3061 GCAATTGATG GTAATGAACT TGACGCACTG ACGATATTAG TCAATTTAAC CGATAAGACT
     3121 GATGATCAGA AAGACCTGCT CGATCGAGCC AAATGCAAGC AAAAACTTCG AGATGAAAAA
     3181 TGGTGGGCTA GCTGCATAAA TTGCGTTAAt TACAGACAAA GCCATAACCC AAAATTCCCG
     3241 GATATACGTT CTGAAGGCGT GATTCGAACC CAAGCCCTgG GTGAATTACC GAGTTTCCTA
     3301 CTCTCGTCcT CAAAAATCCC ACCATACCAT TGGTCATATA GCCATGATTC AAAATACGTC
```

3361 AACAAAAGCG CATTCCTCAC CAATGAGTTT TGTTGGGATG GTGAGATCTC ATGTTTGGGT
3421 GAGCTTCTTA AAGATGCAGA TCACCCACTT TGGAATACTT TGAAAAAGTT AGGTTGTTCT
3481 CAAAAAACTT GCAAAGCAAT GGCAAAACAA CTAGCTGATA TTACTCTCAC GACTATCAAT
3541 GTCACACTCG CACCAAATTA CCTGACTCAA ATATCTCTCC CCGATAGTGA TACATCCTAC
3601 ATTTCACTCT CTCCTGTTGC ATCGCTATCG ATGCAAAGCC ACTTTCATCA GAGGCTCCAA
3661 GATGAAAATC GGCATAGCGC GATAACGCGG TTTAGCCGAA CTACCAACAT GGGGGTCACT
3721 GCGATGACAT GTGGCGGCGC ATTTAGAATG TTAAAGTCTG GCGCTAAGTT TTCCAGCCCC
3781 CCTCATCACC GATTAAACAG TAAACGAAGT TGGTTgACGT CAGAGCATGT TCAGTCATTA
3841 AAACAGTACC AGCGCCTCAA CAAAAGCCTC ATACCTGAAA ACTCTCGGAT TGCACTCCGT
3901 AGAAAATACA AAATCGAGCT TCAAAATATG GTCAGATCTT GGTTTGCAAT GCAAGACCAT
3961 ACCCTTGATT CGAACATACT TATCCAACAT TTGAATCATG ACCTATCTTA CTTAGGAGCC
4021 ACAAAACGTT TTGCATACGA TCCAGCGATG ACCAAGCTCT TTACTGAGCT TTTGAAACGA
4081 GAGTTATCAA ATTCAATCAA TAATGGTGAG CAACACACTA ATGGATCGTT TTTAGTCCTA
4141 CCGAATATCA GGGTTTGTGG CGCAACAGCT TTAAGCTCCC CGGTAACGGT GGGGATtCCA
4201 TCACTTACAG CTTTCTTTGG CTTCGTTCAC GCATTTGAAC GGAATATAAA TCGCACCACC
4261 TCATCGTTTC GTGTTGAATC CTTTGCGATA TGCGTCCATC AACTACATGT CGAGAAGCGA
4321 GGTTTGACAG CAGAGTTTGT GGAAAAAGGC GACGGGACTA TATCCGCTCC CGCGACCCGG
4381 GATGACTGGC AGTGTGATGT CGTATTTAGC CTTATTTTGA ACACCAACTT TGCTCAACAT
4441 ATTGACCAAG ATACGTTAGT TACATCACTA CCAAAGCGAT TGGCTCGgGG TTCAGCAAAA
4501 ATTGCGATTG ATGACTTTAA ACATATCAAC TCATTCTCGA CATTAGAAAC AGCGATCGAA
4561 TCTCTGCCAA TAGAAGCTGG TAGGTGGTTA TCACTTTACG CACAGTCAAA CAATAATCTA
4621 AGTGATCTAT TAGCAGCCAT GACAGAGGAC CATCAGCTCA TGGCAAGCTG CGTCGGTTAC
4681 CACTTGTTAG AAGAGCCCAA AGATAAACCA AACTCCCTCA GAGGTTACAA ACACGCTATC
4741 GCCGAGTGCA TCATTGGACT CATTAACTCA ATCACCTTTA GCTCAGAGAC TGATCCCAAC
4801 ACAATCTTTT GGTCGCTAAA GAACTATCAA AACTACCTAG TGGTACAGCC AAGGAGTATC
4861 AACGATGAAA CTACCGACAA ATCTAGCCTA TGAGCGCTCT ATCGACCCAT CAGATGTCTG
4921 TTTTTTTGTC GTCTGGCCCG ATGATAGAAA AACACCTTTA ACCTACAATT CTCGTACTCT
4981 GCTCGGGCAA ATGGAAGCGG CATCATTAGC CTATGATGTC TCAGGTCAAC CAATAAAAAG
5041 TGCCACCGCT GAGGCGTTAG CTCAAGGGAA CCCTCATCAA GTTGATTTCT GCCACGTTCC
5101 ATAcGGTGCG AGTCATATTG AATGCAGTTT CTCCGTCTCG TTTTCTTCTG AACTACGTCA
5161 ACCATATAAG TGTAACTCAA GCAAAGTTAA ACAAACGCTA GTGCAATTAG TCGAGCTCTA
5221 CGAAACGAAA ATCGGCTGGA CTGAGCTAGC AACCCGATAT TTGATGAAcA TTTGCAACGG
5281 TAAATGGCTG TGGAAAAATA CCCGTAAAGC cTATTGCTGG AACATTGTAC TTACACCTTG
5341 GCCtTGGAAC GGGGAAAAGG TTGGATTTGA AGATATtCGT ACTAACTACA CCTCACGGCA
5401 AGACTTTAAA AATAATAAAA ATTGGTCTGC TATAGTTGAA ATGATCAAAA CCGCATTTTC
5461 TAGTACTGAT GGGCTGGCGA TATTTGAAGT CAGGGCCACC TTGCACTTGC CAACGAATGC
5521 TATGGTGCGG CCAAGCCAAG TTTTCACAGA AAAAGAAAGT GGCAGTAAAA GTAAATCTAA
5581 AACTCAAAAC AGTCGAGTTT TTCAGAGTAC AACTATTGAT GGTGAACGAT CGCCAATACT
5641 AGGGGCCTTT AAAACGGGAG CAGCTATTGC AACCATTGAC GACTGGTATC CTGAAGCCAC
5701 TGAGCCACTA AGGGTCGGAC GGTTTGGGGT TCATCGCGAA GATGTCACTT GCTACCGTCA
5761 TCCGTCTACC GGAAAAGATT TTTTCTCGAT ATTACAACAA GCAGAGCACT ATATTGAAGT
5821 GTTGAGCGCC AACAAAACTC CCGCTCAAGA AACTATCAAC GACATGCACT TTTTAATGGC
5881 TAACCTGATT AAGGGTGGGA TGTTCCAGCA TAAAGGAGAC TGACTGTGAA ATGGTATTAT
5941 AAGACAATCA CCTTTCTGCC AGAGTTGTGC AACAACGAGT CACTGGCTGC AAAGTGTCTc
6001 CGCGTTCTGC ATGGATTTAA CTATCAGTAT GAGACACGAA ATATcGGCGT TTCATTTCCG
6061 CTTTGGTGTG ATGCAACGGT TGGAAAAAAG ATTTCATTTG TCAGCAAGAA CAAGATAGAA
6121 CTCGACTTAC TACTTAAACA ACACTATTTC GTCCAAATGG AACAACTTCA ATAcTTTCAT
6181 ATATCCAACA CTGTTCTCGT CCCAGAAGAT TGTACATACG TTTCCTTTAG ACGCTGTCAA
6241 TCTATAGATA AGCTCACAGC AGCAGGGCTG GCAAGGAAAA TCAGACGCCT GGAGAAACGT
6301 GCTCTATCTA GAGGCGAGCA ATTTGACCCA TCATCTTTTG CTCAAAAAGA GCATACTGCA
6361 ATAGCGCACT ACCACTCACT TGGGGAGTCC AGCAAACAGA CGAACCGCAA CTTTCGACTC
6421 AATATCAGGA TGCTCTCGGA GCAACCCCGT GAGGGGAACT CGATTTTTAG TAGCTATGGC
6481 TTATCAAATT CAGAAAACTC GTTTCAGCCT GTACCCTTAA TCTGACCTTT AATAAGGAGA
6541 TATACCATGG cgACAAGtTT ACCTACGCCC TCAGCAATTA CGACTTCGGC GTTAGAGTAT
6601 GCATTCCATA CTCCCGCTCG CAATCTAACG AAATCTCGCG GAAAAAAcAT TCATCGTTAT
6661 GTCAGTGTAA AGATGAGTAA GAGGATTACG GTAGAATCTA CTCTAGAGTG TGATGCCTGC
6721 TATCACTTTG ATTTTGAGCC AAGTATTGTT CGCTTTTGCG CTCAACCGAT TCGATTTTTA

6781 TATTATCTCA ATGGTCAGTC TCACTCCTAT GTTCCTGACT TTCTAGTTCA ATTTGATACC
6841 AACGAGTTTG TTCTATATGA AGTAAAGTCA GCTTATGCTA AGAACAAACC TGATTTTGAT
6901 GTTGAATGGG AGGCGAAAGT AAAAGCAGCA ACTGAACTAG GGCTAGAATT GGAGCTTGTT
6961 GAAGAGAGTG ATATTAGGGA TACGGTTGTA TTAAATAATC TTAAGCGCAT GCATCGTTAT
7021 GCTTCGAAAG ATGAGCTGAA TAACGTACAT AACTCTCTCT TAAAAATAAT AAAGTACAAT
7081 GGCGCCCAAT CTGCAAGATG CTTGGGAGAA CAGTTGGGTT TAAAAGGCCG AACTGTTTTA
7141 CCAATTTTGT GCGATTTGCT GTCAAGGTGT TTACTCGATA CACGTTTGGA TAAGCCTCTA
7201 TCTCTTGAAT CTCGATTTGA GTTGGCCAGT TATGGCTAAG AAAGGGTTCT CaAGTTTCCA
7261 TAGAAAAGCA GTCTCTTCCC AAGATACGCT TGAATCGATA GAGCTTGTCT CTAGCGCTAA
7321 TTGTCTAGAA AGTGTTACGT ATCAAGATAT ATCAGCATTT CCCGAAACAA TTGCGGTAGA
7381 GATTAATTTC CGATTAAGCA TTCTTCGCTT TTTAGCGCGA AAGTGCGAAA CCATTGTGGC
7441 CAAATCAATT GAACCACATC GTGTAGAGCT ACAGCAAAAC TATAGTAGAA AAATACCCAG
7501 TGCAATAACG ATATATCGAT GGTGGCTTGC TTTTCGAAAA TCAGACTACA ACCCCATTAG
7561 CTTAGCACCT AATATCAAGG ATAGAGGTAA TAGAGAGACA AAAGTGTCAA CAGTTGTTGA
7621 TTCTATTATG GAACAGGCAG TTGAAAGAGT TATATCTGGA CGAAAAGTCA ATGTTAGCTC
7681 TGCATATAAA CGTGTTCGAC GAAAAGTTCG TCAATACAAT CTAACTCATG GAACGAAATA
7741 CACGTATCCT AAGTACGAAT CTGTAAGAAA GCGAGTAAAA AAGAAAACCC CATTTGAGTT
7801 ATTAGCCGCA GGGAAAGGGG AGAGAGTAGC TAAGAGAGAG TTTCGCCGAA TGGGAAAAAA
7861 GATCCTCACG TCTAGCGTGC TAGAGAGGGT TGAAATAGAT CACACTGTCG TTGACCTTTT
7921 TGCAGTACAC GAAGAGTATC GAATcCCATT GGGCCGACCT TGGCTTACTC AATTGGTTGA
7981 TTGTTACAGT AAAGCTGTAA TCGGTTTTTA TTTAGGTTTC GAGCCTCCTA GCTATGTGTC
8041 GGTTTCCCTT GCACTTAAGA ATGCAATACA ACGCAAAGAT GACTTAATCT CCTCGTATGA
8101 ATCGATCGAG AATGAATGGC TATGTTATGG cATCCCAGAC CTACTCGTAA CTGATAATGG
8161 TAAAGAGTTT TTGTCGAAAG CATTTGATCA AGCATGTGAA TCACTATTGA TCAATGTGCA
8221 TCAAAATAAA GTTGAGACGC CCGACAACAA ACCTCATGTT GAACGTAACT ACGGGACTAT
8281 TAATACTTCT CTGTTAGACG ATTTACCTGG GAAATCCTTC AGCCAGTACC TTCAAAGAGA
8341 AGGGTACGAC TCTGTGGGAG AAGCTACCCT TACACTCAAT GAGATTAGAG AAATTTACTT
8401 AATTTGGTTG GTGGATATTT ATCATAAAAA ACCCAATCAG AGAGGCACTA ATTGTCCTAA
8461 TGTTGCGTGG AAAAAGGGTT GTCAAGAATG GGAACCAGAG GAGTTCTCTG GTTCTAAAGA
8521 CGAATTAGAC TTTAAATTTG CTATTGTTGA TTACAAACAA CTTACTAAAG TAGGGATAAC
8581 TGTCTACAAA GAACTGAGTT ATAGCAATGA CCGTTTAGCT GAATATAGAG GGAAGAAAGG
8641 AAACCATAAA GTTCAGTTCA AGTATAACCC TGAGTGTATG GCAGTTATTT GGGTGTTGGA
8701 TGAGGATATG AATGAGTACT TTACAGTTAA TGCGATTGAC TACGAATATG CAAGTAGAGT
8761 ATCACTTTGG CAACATAAAT ATAACATGAA ATATCAAGCA GAACTAAATT CAGCAGAATA
8821 TGATGAgGAC AAGGAAATTG ATGCAGAAAT AAAAATTGAA GAAATCGCAG ATCGTTCAAT
8881 TGTTAAGACT AACAAAATCA GAGCTCGGAG GCGTGGCGCT AGGCATCAAG AGAATAGCGC
8941 AAGGGCTAAG TCAATCAGTA ATGCGAACCC GGCCTCGATA CAAAAACATG AAGATGAAAT
9001 CGTTAGTGCA GATAATGACG ATTGGGATAT TGATTATGTC TGAGAAATCG TCAAATGAGT
9061 GAAACGCGTG AGGCTCGTAT ATCAAGAGCT AAAAGGGCAT TTGTATCCAC ACCGAGCGTT
9121 AGGAAAATCT TGAGTTACAT GGATAGATGT AGAGAtCTAT CAGACCTAGA GTCTGAGCCT
9181 ACATGCATGA TGGTCTATGG TGCTTCGGGT GTAGGTAAAA CGACCGTCAT CAAGAAATAC
9241 TTAAATCAAA ACAGAAGAGA GTCCGAAGCC GGGGGCGATA TAATACCGGT TTTGCATATT
9301 GAGTTGCCAG ACAATGCGAA GCCAGTAGAT GCAGCAAGGG AATTGCTGGT TGAAATGGGT
9361 GACCCGCTAG CACTTTATGA AACTGACTTA GCTAGATTGA CGAAAAGACT GACTGAATTA
9421 ATCCCTGCGG TCGGCGTGAA GCTGATTATT ATCGATGAGT TCCAACATTT GGTGGAAGAA
9481 AGGTCAAATC GGGTTCTTAC CCAAGTAGGT AATTGGCTAA AAATGATACT TAACAAAACG
9541 AAATGTCCAA TTGTTATATT TGGTATGCCA TACTCAAAAG TTGTACTGCA AGCAAACTCG
9601 CAACTTCACG GGCGATTTTC CATTCAGGTT GAACTGCGCC CCTTTAGCTA CCAGGGAGGT
9661 AGAGGTGTAT TTAAAACTTT TTTGGAATAC CTTGATAAAG CCCTACCTTT TGAAAAACAG
9721 GCTGGCTTAG CCAACGAAAG TTTGCAGAAA AAATTGTATG CATTCTCTCA GGGAAACATG
9781 CGTTCGTTGA GAAACCTTAT TTATCAAGCA TCTATCGAAG CAATTGATAA TCAGCATGAG
9841 ACGATAACCG AAGAAGATTT CGTTTTTGCA TCGAAGTTGA CATCGGGCGA TAAACCCAAC
9901 TCATGGAAAA ATCCTTTTGA GGAGGGTGTT GAGGTAACAG AAGATATGTT ACGACCGCCA
9961 CCAAAAGATA TTGGTTGGGA gGACTATTTG AGACATTCAA CCCCGAGAGT GAGTAAACCA
10021 GGTAGAAATA AAAACTTTTT CGAATAACCT AGGCTGCTGC GCTGCTGCCA CCGCTGAGCA
10081 ATAACTAGCA TAACCCCTTG GGGCCTCTAA ACGGGTCTTG AGGGGTTTTT TGCTGAAACC
10141 TCAGGCATTT GTTGTTGATA CAACCATAAA ATGATAATTA CACCCATAAA TTGATAATTA

```
    10201 TCACACCCAT AAATTGATAT TGCCTCTTCA TGGTCTAAAC TTCAGTAAGT TTACGACATT
    10261 TTCCTCGAGG TCATTTCcca attattgaag gccgctaacg cggccttttt ttgtttctgg
    10321 tctgcctcct CCCTGCAGGC TTCAACAAAC AGacaatctg gtctgtttgt attatggaaa
    10381 atttttctgt ataatagatt caacaaacag acaatctggt ctgtttgtat tatcgacCAA
    10441 CCCACTCCCA TGGTGTGACG GGCGGTGTGT ACAAGGCCCG GGAACGTgTT CACAAAAGTT
    10501 ATCAGGCATG CACCTGGTAG CTAGTCTTTA AACCAATAGA TTGCATCGGT TTAAAAGGCA
    10561 AGACCGTCAA ATTGCGGGAA AGGGGTCAAC AGCCGTTCAG TACCAAGTCT CAGGGGAAAC
    10621 TTTGAGATGG CCTTGCAAAG GGTATGGTAA TAAGCTGACG GACATGGTCC TAACCACGCA
    10681 GCCAAGTCCT AAGTCAACAG ATCTTCTGTT GATATGGATG CAGTTCACAG ACTAAATGTC
    10741 GGTCGGGGAA GATGTATTCT TCTCATAAGA TATAGTCGGA CCTCTCCTTA ATGGGAGCTA
    10801 GCGGATGAAG TGATGCAACA CTGGAGCCGC TGGGAACTAA TTTGTATGCG AAAGTATATT
    10861 GATTAGTTTT GGAGTACTCG ATGGTGTTCA ATGCTTTTCC CGAGTGGTTA TCCGGATCAC
    10921 ATGAAACGGC ATGACTTTTT CAAGAGTGCC ATGCCCGAAG GTTATGTACA GGAACGCACT
    10981 ATATCTTTCA AAGATGACGG GACCTACAAG ACGCGTGCTG AAGTCAAGTT TGAAGGTGAT
    11041 ACCCTTGTTA ATCGTATCGA GTTAAAGGGT ATTGATTTTA AAGAAGATGG AAACATTCTT
    11101 GGACACAAAC TCGAGTACAA CTTTAACTCA CACAATGTAT ACATCACGGC AGACAAACAA
    11161 AAGAATGGAA TCAAAGCTAA CTTCAAAATT CGCCACAACG TTGAAGATGG TTCCGTTCAA
    11221 CTAGCAGACC ATTATCAACA AAATACTCCA ATTGGCGATG GCCCTGTCCT TTTACCAGAC
    11281 AACCATTACC TGTCGACACA ATCTGTCCTT TCGAAAGATC CCAACGAAAA GCGTGACCAC
    11341 ATGGTCCTTC TTGAGTTTGT AACTGCTGCT GGGATTACAC ATGGCATGGA TGAGCTCTAC
    11401 AAATGGCcca attattgaag gcctccctaa cggggggcct ttttttGTTT CTGGTCTGCC
    11461 GCTGCGCTTA CTGCAGTAGT TTTGCTGAAA TACTCGATTC ACAAAAATAT CAACTTATGG
    11521 TTGTTTTGTG AGATATCAAT ATATGGTTGT TTTGTGGTTA AGTTGCTGAT TATAAATAAT
    11581 TATTAAATAT CACTTTATGG TTGCATCAAC Atcagatcct tccgtattta gccagtatgt
    11641 tctctagtgt ggttcgttgt ttttgcgtga gccatgagaa cgaaccattg agatcatact
    11701 tactttgcat gtcactcaaa aattttgcct caaaactggt gagctgaatt tttgcagtta
    11761 aagcatcgtg tagtgttttt cttagtccgt tacgtaggta ggaatctgat gtaatggttg
    11821 ttggtatttt gtcaccattc atttttatct ggttgttctc aagttcggtt acgagatcca
    11881 tttgtctatc tagttcaact tggaaaatca acgtatcagt cgggcggcct cgcttatcaa
    11941 ccaccaattt catattgctg taagtgttta aatctttact tattggtttc aaaacccatt
    12001 ggttaagcct tttaaactca tggtagttat tttcaagcat taacatgaac ttaaattcat
    12061 caaggctaat ctctatattt gccttgtgag ttttcttttg tgttagttct tttaataacc
    12121 actcataaat cctcatagag tatttgtttt caaaagactt aacatgttcc agattatatt
    12181 ttatgaattt ttttaactgg aaaagataag gcaatatctc ttcactaaaa actaattcta
    12241 atttttcgct tgagaacttg gcatagtttg tccactggaa aatctcaaag cctttaacca
    12301 aaggattcct gatttccaca gttctcgtca tcagctctct ggttgcttta gctaatacac
    12361 cataagcatt ttccctactg atgttcatca tctgagcgta ttggttataa gtgaacgata
    12421 ccgtccgttc tttccttgta gggttttcaa tcgtggggtt gagtagtgcc acacagcata
    12481 aaattagctt ggtttcatgc tccgttaagt catagcgact aatcgctagt tcatttgctt
    12541 tgaaaacaac taattcagac at
//
```

**pESCccdB**

```
LOCUS       pEscapeCcdB            9110 bp ds-DNA     circular     09-APR-
2026
DEFINITION  .
KEYWORDS    "Author" "Plasmid" "Freezer" "Rack" "Box"
FEATURES             Location/Qualifiers
     misc_feature    1..3211
                     /label="From pMAS048"
                     /ApEinfo_revcolor="#ff9ccd"
                     /ApEinfo_fwdcolor="#ff9ccd"
     misc_feature    5..160
                     /label="ConL 'S"
                     /ApEinfo_revcolor="#84b0dc"
                     /ApEinfo_fwdcolor="#84b0dc"
     misc_feature    3216..3369
                     /label="conL2"
                     /ApEinfo_revcolor="#9eafd2"
                     /ApEinfo_fwdcolor="#9eafd2"
     misc_feature    3370..3459
                     /label="Pcym"
                     /ApEinfo_revcolor="#ffef86"
                     /ApEinfo_fwdcolor="#ffef86"
     misc_feature    3464..4424
                     /label="Ribozyme"
                     /ApEinfo_revcolor="#ffef86"
                     /ApEinfo_fwdcolor="#ffef86"
     misc_feature    4864..5169
                     /label="ccdB"
                     /ApEinfo_revcolor="#faac61"
                     /ApEinfo_fwdcolor="#faac61"
     misc_feature    5170..5230
                     /label="ccdb promoter and such"
                     /ApEinfo_revcolor="#d6b295"
                     /ApEinfo_fwdcolor="#d6b295"
     misc_feature    5231..5431
                     /label="con L2 "
                     /ApEinfo_revcolor="#b4abac"
                     /ApEinfo_fwdcolor="#b4abac"
     misc_feature    5432..6229
                     /label="neomycin"
                     /ApEinfo_revcolor="#b1ff67"
                     /ApEinfo_fwdcolor="#b1ff67"
ORIGIN
        1 ACGGAAAAAA AAACCCCGCC CCTGACAGGG CGGGGTTTTT TTTCCAATTA TTGAAGGCCG
       61 CTAACGCGGC CTTTTTTTGT TTCTGGTCTG CCTTAATCAA TGACTAAGAA TTCGCGGCCG
      121 CTTCTAGAGG GTCTCACTTC CTCGAGGGCG CGCCTGCAGG CTTCGGATCC GCTGCAGGCT
      181 TCGCGGCGCG CCATCGAATG GTGCAAAACC TTTCGCGGTA TGGCATGATA GCGCCCGGAA
      241 GAGAGTCAAT TCAGGGTGGT GAATATGAGC CCGAAACGTC GTACCCAGGC AGAACGTGCA
      301 ATGGAAACCC AGGGTAAACT GATTGCAGCA GCACTGGGTG TTCTGCGTGA AAAAGGTTAT
      361 GCAGGTTTTC GTATTGCAGA TGTTCCGGGT GCAGCCGGTG TTAGCCGTGG TGCACAGAGC
      421 CATCATTTTC CGACCAAACT GGAACTGCTG CTGGCAACCT TTGAATGGCT GTATGAGCAG
      481 ATTACCGAAC GTAGCCGTGC ACGTCTGGCA AAACTGAAAC CGGAAGATGA TGTTATTCAG
      541 CAGATGCTGG ATGATGCAGC AGAATTTTTT CTGGATGATG ATTTTAGCAT CGGCCTGGAT
      601 CTGATTGTTG CAGCAGATCG TGATCCGGCA CTGCGTGAAG GTATTCAGCG TACCGTTGAA
      661 CGTAATCGTT TTGTTGTTGA AGATATGTGG CTGGGTGTGC TGGTGAGCCG TGGTCTGAGC
```

721 CGTGATGATG CCGAAGATAT TCTGTGGCTG ATTTTTAACA GCGTTCGTGG TCTGGTAGTT
781 CGTAGCCTGT GGCAGAAAGA TAAAGAACGT TTTGAACGTG TGCGTAATAG CACCCTGGAA
841 ATTGCACGTG AACGTTATGC AAAATTCAAA CGTTGATAAT GGCTTCCCGA CAGCTGTCAC
901 CGGATGTGCT TTCCGGTCTG ATGAGTCCGT GAGGACGAAA CAGCCTCTAC AAATAATTTT
961 GTTTAATCAG AGGTCACGCG AGTTAAGGAG GTTTAATGAA GCAGCGTATT ACAGTGACAG
1021 TTGACAGCGA CAGCTATCAG TTGCTCAAGG CATATGATGT CAATATCTCC GGTCTGGTAA
1081 GCACAACCAT GCAGAATGAA GCCCGTCGTC TGCGTGCCGA ACGCTGGAAA GCGGAAAATC
1141 AGGAAGGGAT GGCTGAGGTC GCCCGGTTTA TTGAAATGAA CGGCTCTTTT GCTGACGAGA
1201 ACAGGGACTG GTGATGGCTT CAGCCAAAAA ACTTAAGACC GCCGGTCTTG TCCACTACCT
1261 TGCAGTAATG CGGTGGACAG GATCGGCGGT TTTCTTTTCT CTTCTCAAGC TGGCTGGCAT
1321 GCGGCGCGCC GCTGTGAGAC CTGCAGGTAC TAGTAGCGGC CGCTGCAGTT AATCACTGAT
1381 TAACTCGGTA CCAAATTCCA GAAAAGAGGC CTCCCGAAAG GGGGGCCTTT TTTCGTTTTG
1441 GTCCTTTTGT TATCAATAAA AAAGGGGAGC GGTTTCCCGC TCCCCTTATT GTTCGTCTAC
1501 AACCAATAAA AAACGCCCGG CGGCAACCGA GCGTTCTGAA CAAATCCAGA TGGAGTATTA
1561 CGCCCCGCCC TGCCACTCAT CGCAGTACTG TTGTAATTCA TTAAGCATTC TGCCGACATG
1621 GAAGCCATCA CAAACGGCAT GATGAACCTG AATCGCCAGC GGCATCAGCA CCTTGTCGCC
1681 TTGCGTATAA TATTTGCCCA TGGTGAAAAC GGGGGCGAAG AAGTTGTCCA TATTGGCCAC
1741 GTTTAAATCA AAACTGGTGA AACTCACCCA GGGATTGGCT GAAACGAAAA ACATATTCTC
1801 AATAAACCCT TTAGGGAAAT AGGCCAGGTT TTCACCGTAA CACGCCACAT CTTGCGAATA
1861 TATGTGTAGA AACTGCCGGA AATCGTCGTG GTATTCACTC CAGAGCGATG AAAACGTTTC
1921 AGTTTGCTCA TGGAAAACGG TGTAACAAGG GTGAACACTA TCCCATATCA CCAGCTCACC
1981 GTCTTTCATT GCCATACGAA ATTCCGGATG AGCATTCATC AGGCGGGCAA GAATGTGAAT
2041 AAAGGCCGGA TAAAACTTGT GCTTATTTTT CTTTACGGTC TTTAAAAAGG CCGTAATATC
2101 CAGCTGAACG GTCTGGTTAT AGGTACATTG AGCAACTGAC TGAAATGCCT CAAAATGTTC
2161 TTTACGATGC CATTGGGATA TATCAACGGT GGTATATCCA GTGATTTTTT TCTCCATTTT
2221 AGCTTCCTTA GCTCCTGAAA ATCTCGATAA CTCAAAAAAT ACGCCCGGTA GTGATCTTAT
2281 TTCATTATGG TGAAAGTTGG AACCTCTTAC GTGCCCGATC AAGAGTCCTG TTGAGTAATA
2341 GTCAAAAGCC TCCGGTCGGA GGCTTTTGAC TTTCTGCTTA CCCAATAGCG GAGTGTATAC
2401 TGGCTTACTA TGTTGGCACT GATGAGGGTG TCAGTGAAGT GCTTCATGTG GCAGGAGAAA
2461 AAAGGCTGCA CCGGTGCGTC AGCAGAATAT GTGATACAGG ATATATTCCG CTTCCTCGCT
2521 CACTGACTCG CTACGCTCGG TCGTTCGACT GCGGCGAGCG GAAATGGCTT ACGAACGGGG
2581 CGGAGATTTC CTGGAAGATG CCAGGAAGAT ACTTAACAGG GAAGTGAGAG GGCCGCGGCA
2641 AAGCCGTTTT TCCATAGGCT CCGCCCCCCT GACAAGCATC ACGAAATCTG ACGCTCAAAT
2701 CAGTGGTGGC GAAACCCGAC AGGACTATAA AGATACCAGG CGTTTCCCCC TGGCGGCTCC
2761 CTCGTGCGCT CTCCTGTTCC TGCCTTTCGG TTTACCGGTG TCATTCCGCT GTTATGGCCG
2821 CGTTTGTCTC ATTCCACGCC TGACACTCAG TTCCGGGTAG GCAGTTCGCT CCAAGCTGGA
2881 CTGTATGCAC GAACCCCCCG TTCAGTCCGA CCGCTGCGCC TTATCCGGTA ACTATCGTCT
2941 TGAGTCCAAC CCGGAAAGAC ATGCAAAAGC ACCACTGGCA GCAGCCACTG GTAATTGATT
3001 TAGAGGAGTT AGTCTTGAAG TCATGCGCCG GTTAAGGCTA AACTGAAAGG ACAAGTTTTG
3061 GTGACTGCGC TCCTCCAAGC CAGTTACCTC GGTTCAAAGA GTTGGTAGCT CAGAGAACCT
3121 TCGAAAAACC GCCCTGCAAG GCGGTTTTTT CGTTTTCAGA GCAAGAGATT ACGCGCAGAC
3181 CAAAACGATC TCAAGAAGAT CATCTTATTA AACGGAAAAA AAAACCCCGC CCCTGACAGG
3241 GCGGGGTTTT TTTTCCAATT ATTGAAGGCC GCTAACGCGG CCTTTTTTTG TTTCTGGTCT
3301 GCCTTAATCA ATGACTAAGA ATTCGCGGCC GCTTCTAGAG CGTCTCACGT TGCTAGCCCT
3361 GCAGGCTTCA ACAAACAGAC AATCTGGTCT GTTTGTATTA TGGAAAATTT TTCTGTATAA
3421 TAGATTCAAC AAACAGACAA TCTGGTCTGT TTGTATTATC GACCAACCCA CTCCCATGGT
3481 GTGACGGGCG GTGTGTACAA GGCCCGGGAA CGTGTTCACA AAGTTATCA GGCATGCACC
3541 TGGTAGCTAG TCTTTAAACC AATAGATTGC ATCGGTTTAA AAGGCAAGAC CGTCAAATTG
3601 CGGGAAAGGG GTCAACAGCC GTTCAGTACC AAGTCTCAGG GGAAACTTTG AGATGGCCTT
3661 GCAAAGGGTA TGGTAATAAG CTGACGGACA TGGTCCTAAC CACGCAGCCA AGTCCTAAGT
3721 CAACAGATCT TCTGTTGATA TGGATGCAGT TCACAGACTA AATGTCGGTC GGGGAAGATG
3781 TATTCTTCTC ATAAGATATA GTCGGACCTC TCCTTAATGG GAGCTAGCGG ATGAAGTGAT
3841 GCAACACTGG AGCCGCTGGG AACTAATTTG TATGCGAAAG TATATTGATT AGTTTTGGAG
3901 TACTCGATGG TGTTCAATGC TTTTCCCGTT ATCCGGATCA CATGAAACGG CATGACTTTT
3961 TCAAGAGTGC CATGCCCGAA GGTTATGTAC AGGAACGCAC TATATCTTTC AAAGATGACG
4021 GGACCTACAA GACGCGTGCT GAAGTCAAGT TTGAAGGTGA TACCCTTGTT AATCGTATCG
4081 AGTTAAAGGG TATTGATTTT AAAGAAGATG GAAACATTCT TGGACACAAA CTCGAGTACA

4141 ACTTTAACTC ACACAATGTA TACATCACGG CAGACAAACA AAAGAATGGA ATCAAAGCTA
4201 ACTTCAAAAT TCGCCACAAC GTTGAAGATG GTTCCGTTCA ACTAGCAGAC CATTATCAAC
4261 AAAATACTCC AATTGGCGAT GGCCCTGTCC TTTTACCAGA CAACCATTAC CTGTCGACAC
4321 AATCTGTCCT TTCGAAAGAT CCCAACGAAA AGCGTGACCA CATGGTCCTT CTTGAGTTTG
4381 TAACTGCTGC TGGGATTACA CATGGCATGG ATGAGCTCTA CAAATGGCCC AATTATTGAA
4441 GGCCTCCCTA ACGGGGGGCC TTTTTTTGTT TCTGGTCTGC CGCTGAGTTC ACCGACAAAC
4501 AACAGATAAA ACGAAAGGCC CAGTCTTTCG ACTGAGCCTT TCGTTTTATT TGATGCCTGG
4561 CAGTTCCCTA CTCTCGCATG GGGAGACCCC ACACTACCAT CGGCGCTACG GCGTTTCACT
4621 TCTGAGTTCG GCATGGGGTC AGGTGGGACC ACCGCGCTAC TGCCGCCAGG CAAATTCTGT
4681 TTTATCAGAC CGCTTCTGCG TTCTGATTTA ATCTGTATCA GGCTGAAAAT CTTCTCTCAT
4741 CCGCCAAAAC AGCCAAGCTG GAGACCGTTT AAACTCAATG ATGATGATGA TGATGGTCGA
4801 CGGCGCTATT CAGATCCTCT TCTGAGATGA GTTTTTGTTA GGGCCCAAGC TTCAGATCCC
4861 ATATTATATT CCCCAGAACA TCAGGTTAAT GGCGTTTTTG ATGTCATTTT CGCGGTGGCT
4921 GAGATCAGCC ACTTCTTCCC CGATAACGGA GACCGGCACA CTGGCCATAT CGGTGGTCAT
4981 CATGCGCCAG CTTTCATCCC CGATATGCAC CACCGGGTAA AGTTCACGGG AGACTTTATC
5041 TGACAGCAGA CGTGCACTGG CCAGGGGGAT CACCATCCGT CGCCCGGGCG TGTCAATAAT
5101 ATCACTCTGT ACATCCACAA ACAGACGATA ACGGCTCTCT CTTTTATAGG TGTAAACCTT
5161 AAACTGCATA GGTCATTCCT CCTAGATCCA AAATACGGTA CCGTCAACAA TCTCACTCGA
5221 GAGTCGACGT GCTAGCATTG TACCTAGGAC TGAGCTAGCC ATAAACGTTT GAGACGTACT
5281 AGTAGCGGCC GCTGCAGTTA ATCACTGATT AACTCGGTAC CAAATTCCAG AAAAGAGGCC
5341 TCCCGAAAGG GGGGCCTTTT TTCGTTTTGG TCCTTTTGTT ATCAATAAAA AAGGGGAGCG
5401 GTTTCCCGCT CCCCTTATTG TTCGTCTACA ATTATTAGAA GAACTCGTCA AGAAGGCGAT
5461 AGAAGGCGAT GCGCTGCGAA TCGGGAGCGG CGATACCGTA AAGCACGAGG AAGCGGTCAG
5521 CCCATTCGCC GCCAAGCTCT TCAGCAATAT CACGGGTAGC CAACGCTATG TCCTGATAGC
5581 GGTCCGCCAC ACCCAGCCGG CCACAGTCGA TGAATCCAGA AAAGCGGCCA TTTTCCACCA
5641 TGATATTCGG CAAGCAGGCA TCGCCATGGG TCACGACGAG ATCCTCGCCG TCGGGCATGC
5701 GCGCCTTGAG CCTGGCGAAC AGTTCGGCTG GCGCGAGCCC CTGATGCTCT TCGTCCAGAT
5761 CATCCTGATC GACAAGACCG GCTTCCATCC GAGTACGTGC TCGCTCGATG CGATGTTTCG
5821 CTTGGTGGTC GAATGGGCAG GTAGCCGGAT CAAGCGTATG CAGCCGCCGC ATTGCATCAG
5881 CCATGATGGA TACTTTCTCG GCAGGAGCAA GGTGAGATGA CAGGAGATCC TGCCCCGGCA
5941 CTTCGCCCAA TAGCAGCCAG TCCCTTCCCG CTTCAGTGAC AACGTCGAGC ACAGCTGCGC
6001 AAGGAACGCC CGTCGTGGCC AGCCACGATA GCCGCGCTGC CTCGTCCTGC AGTTCATTCA
6061 GGGCACCGGA CAGGTCGGTC TTGACAAAAA GAACCGGGCG CCCCTGCGCT GACAGCCGGA
6121 ACACGGCGGC ATCAGAGCAG CCGATTGTCT GTTGTGCCCA GTCATAGCCG AATAGCCTCT
6181 CCACCCAAGC GGCCGGAGAA CCTGCGTGCA ATCCATCTTG TTCAATCATG CGAAACGATC
6241 CTCATCCTGT CTCTTGATCA GATCTTGATC CCCTGCGCCA TCAGATCCTT GGCGGCAAGA
6301 AAGCCATCCA GTTTACTTTG CAGGGCTTCC CAACCTTACC AGAGGGCGCC CCAGCTGGCA
6361 ATTCCGGTTC GCTTGCTGTC CATAAGAGTC CTGTTGAGTA ATAGTCAAAA GCCTCCGGTC
6421 GGAGGCTTTT GACTTTCTGC TTACCCAATT ACTACCGGCG CGGCAGCGTG ACCCGTGTCG
6481 GCGGCTCCAA CGGCTCGCCA TCGTCCAGAA AACACGGCTC ATCGGGCATC GGCAGGCGCT
6541 GCTGCCCGCG CCGTTCCCAT TCCTCCGTTT CGGTCAAGGC TGGCAGGTCT GGTTCCATGC
6601 CCGGAATGCC GGGCTGGCTG GGCGGCTCCT CGCCGGGGCC GGTCGGTAGT TGCTGCTCGC
6661 CCGGATACAG GGTCGGGATG CGGCGCAGGT CGCCATGCCC CAACAGCGAT TCGTCCTGGT
6721 CGTCGTGATC AACCACCACG GCGGCACTGA ACACCGACAG GCGCAACTGG TCGCGGGGCT
6781 GGCCCCACGC CACGCGGTCA TTGACCACGT AGGCCGACAC GGTGCCGGGG CCGTTGAGCT
6841 TCACGACGGA GATCCAGCGC TCGGCCACCA AGTCCTTGAC TGCGTATTGG ACCGTCCGCA
6901 AAGAACGTCC GATGAGCTTG GAAAGTGTTT TCTGGCTGAC CACCACGGCG TTCTGGTGGC
6961 CCATCTGCGC CACGAGGTGA TGCAGCAGCA TTGCCGCCGT GGGTTTCCTC GCAATAAGCC
7021 CGGCCCACGC CTCATGCGCT TTGCGTTCCG TTTGCACCCA GTGACCGGGC TTGTTCTTGG
7081 CTTGAATGCC GATTTCTCTG GACTGCGTGG CCATGCTTAT CTCCATGCGG TAGGGGTGCC
7141 GCACGGTTGC GGCACCATGC GCAATCAGCT GCAACTTTTC GGCAGCGCGA CAACAATTAT
7201 GCGTTGCGTA AAAGTGGCAG TCAATTACAG ATTTTCTTTA ACCTACGCAA TGAGCTATTG
7261 CGGGGGGTGC CGCAATGAGC TGTTGCGTAC CCCCCTTTTT TAAGTTGTTG ATTTTTAAGT
7321 CTTTCGCATT TCGCCCTATA TCTAGTTCTT TGGTGCCCAA AGAAGGGCAC CCCTGCGGGG
7381 TTCCCCCACG CCTTCGGCGC GGCTCCCCCT CCGGCAAAAA GTGGCCCCTC CGGGGCTTGT
7441 TGATCGACTG CGCGGCCTTC GGCCTTGCCC AAGGTGGCGC TGCCCCCTTG GAACCCCCGC
7501 ACTCGCCGCC GTGAGGCTCG GGGGGCAGGC GGGCGGGCTT CGCCCTTCGA CTGCCCCCAC

```
     7561 TCGCATAGGC TTGGGTCGTT CAGGCGCGTC AAGGCCAAGC CGCTGCGCGG TCGCTGCGCG
     7621 AGCCTTGACC CGCCTTCCAC TTGGTGTCCA ACCGGCAAGC GAAGCGCGCA GGCCGCAGGC
     7681 CGGAGGCTTT TCCCCAGAGA AAATTAAAAA AATTGATGGG GCAAGGCCGC AGGCCGCGCA
     7741 GTTGGAGCCG GTGGGTATGT GGTCGAAGGC TGGGTAGCCG GTGGGCAATC CCTGTGGTCA
     7801 AGCTCGTGGG CAGGCGCAGC CTGTCCATCA GCTTGTCCAG CAGGGTTGTC CACGGGCCGA
     7861 GCGAAGCGAG CCAGCCGGTG GCCGCTCGCG GCCATCGTCC ACATATCCAC GGGCTGGCAA
     7921 GGGAGCGCAG CGACCGCGCA GGGCGAAGCC CGGAGAGCAA GCCCGTAGGG CGCCGCAGCC
     7981 GCCGTAGGCG GTCACGACTT TGCGAAGCAA AGTCTAGTGA GTATACTCAA GCATTGAGTG
     8041 GCCCGCCGGA GGCACCGCCT TGCGCTGCCC CCGTCGAGCC GGTTGGACAC CAAAAGGGAG
     8101 GGGCAGGCAT GGCGGCATAC GCGATCATGC GATGCAAGAA GCTGGCGAAA ATGGGCAACG
     8161 TGGCGGCCAG TCTCAAGCAC GCCTACCGCG AGCGCGAGAC TCCCAACGCT GACGCCAGCA
     8221 GGACGCCAGA GAACGAGCAC TGGGCGGCCA GCAGCACCGA TGAAGCGATG GGCCGACTGC
     8281 GCGAGTTGCT GCCAGAGAAG CGGCGCAAGG ACGCTGTGTT GGCGGTCGAG TACGTCATGA
     8341 CGGCCAGCCC GGAATGGTGG AAGTCGGCCA GCCAAGAACA GCAGGCGGCG TTCTTCGAGA
     8401 AGGCGCACAA GTGGCTGGCG GACAAGTACG GGGCGGATCG CATCGTGACG GCCAGCATCC
     8461 ACCGTGACGA AACCAGCCCG CACATGACCG CGTTCGTGGT GCCGCTGACG CAGGACGGCA
     8521 GGCTGTCGGC CAAGGAGTTC ATCGGCAACA AAGCGCAGAT GACCCGCGAC CAGACCACGT
     8581 TTGCGGCAGC TGTGGCCGAT CTAGGGCTGC AACGGGGCAT CGAGGGCAGC AAGGCACGTC
     8641 ACACGCGCAT TCAGGCGTTC TACGAGGCCC TGGAGCGGCC ACCAGTGGGC CACGTCACCA
     8701 TCAGCCCGCA AGCGGTCGAG CCACGCGCCT ATGCACCGCA GGGATTGGCC GAAAAGCTGG
     8761 GAATCTCAAA GCGCGTTGAA ACGCCGGAAG CCGTGGCCGA CCGGCTGACA AAAGCGGTTC
     8821 GGCAGGGGTA TGAGCCTGCC CTACAGGCCG CCGCAGGAGC GCGTGAGATG CGCAAGAAGG
     8881 CCGATCAAGC CCAAGAGACA GCCCGAGATC TTCGGGAGCG CCTGAAGCCC GTTCTGGACG
     8941 CCCTGGGGCC GTTGAATCGG GATATGCAGG CCAAGGCCGC CGCGATCATC AAGGCCGTGG
     9001 GCGAAAAGCT GCTGACGGAA CAGCGGGAAG TCCAGCGCCA GAAACAGGCC CAGCGCCAGC
     9061 AGGAACGCGG GCGCGCACAT TTCCCCGAAA AGTGCCACCT GGGCTGATAA
//
```

**pMAS045**

```
LOCUS       pMAS045                 4372 bp ds-DNA     circular     22-MAY-
2025
DEFINITION  .
KEYWORDS    "Desc:pCas/pTargetF linear donor as plasmid, CmR and
promoterless
            RFP" "Cloning entry:[pAMG046, pAMG047, donor] Promoterless RFP
            strain, pCas/pTargetF" "Origin:ColE1`" "Resistance:Ampicillin,
            Chloramphenicol" "MoClo" "Glycerol stock(s) (mm/dd/yy)" "JEC
alt"
FEATURES             Location/Qualifiers
     rep_origin      1..683
                     /label="ColE1 origin"
                     /ApEinfo_revcolor="#c7b0e3"
                     /ApEinfo_fwdcolor="#c7b0e3"
     misc_feature    461..568
                     /label="RNAI transcript"
                     /ApEinfo_revcolor="#b4abac"
                     /ApEinfo_fwdcolor="#b4abac"
     misc_feature    complement(599..604)
                     /label="BamHI-BglII Scar"
                     /ApEinfo_revcolor="#ac1dff"
                     /ApEinfo_fwdcolor="#ac1dff"
     misc_feature    610..615
                     /label="BamHI-BglII Scar"
                     /ApEinfo_revcolor="#ac1dff"
                     /ApEinfo_fwdcolor="#ac1dff"
     misc_feature    complement(696..701)
                     /label="BamHI-BglII Scar"
                     /ApEinfo_revcolor="#ac1dff"
                     /ApEinfo_fwdcolor="#ac1dff"
     misc_feature    708..713
                     /label="BamHI-BglII Scar"
                     /ApEinfo_revcolor="#ac1dff"
                     /ApEinfo_fwdcolor="#ac1dff"
     CDS             complement(781..1440)
                     /label="AmpR (aminoglycoside n(3)-acetyltransferas)"
                     /ApEinfo_revcolor="#f58a5e"
                     /ApEinfo_fwdcolor="#f58a5e"
     CDS             complement(781..1638)
                     /label="AmpR"
                     /ApEinfo_revcolor="#ffef86"
                     /ApEinfo_fwdcolor="#ffef86"
     misc_feature    complement(1476..1481)
                     /label="BamHI-BglII Scar"
                     /ApEinfo_revcolor="#ac1dff"
                     /ApEinfo_fwdcolor="#ac1dff"
     misc_feature    1493..1498
                     /label="BamHI-BglII Scar"
                     /ApEinfo_revcolor="#ac1dff"
                     /ApEinfo_fwdcolor="#ac1dff"
     misc_feature    1680..1708
                     /label="AmpR promoter"
                     /ApEinfo_revcolor="#ffef86"
                     /ApEinfo_fwdcolor="#ffef86"
```

```
     primer          1761..1778
                     /label="JEC58A | JC.A58"
                     /note="sequence: TGCCACCTGACGTCTAAG"
                     /ApEinfo_revcolor="#85dae9"
                     /ApEinfo_fwdcolor="#85dae9"
     misc_feature    1894..1928
                     /label="Promoter-J23119(SpeI)"
                     /ApEinfo_revcolor="#ffef86"
                     /ApEinfo_fwdcolor="#ffef86"
     primer          complement(1915..1966)
                     /label="AMG34C"
                     /note="sequence:
CTGATTAAATATGATGAAAACGGCAACCCGTCAGATCCACTAGTATTATACC"
                     /ApEinfo_revcolor="#c7b0e3"
                     /ApEinfo_fwdcolor="#c7b0e3"
     misc_feature    complement(1929..1934)
                     /label="BamHI-BglII Scar"
                     /ApEinfo_revcolor="#ac1dff"
                     /ApEinfo_fwdcolor="#ac1dff"
     CDS             complement(1937..1950)
                     /label="lacZ CDS"
                     /ApEinfo_revcolor="#84b0dc"
                     /ApEinfo_fwdcolor="#84b0dc"
                     /note="locus_tag: b0344 gene_synonym: ECK0341
EC_number: 3.2.1.108 transl_table: 11 product: beta-galactosidase
protein_id: NP_414878.1 db_xref: GeneID:945006 translation:
MTMITDSLAVVLQRRDWENPGVTQLNRLAAHPPFASWRNSEEARTDRPSQQLRSLNGEWRFAWFPAPEAVPESWLE
CDLPEADTVVVPSNWQMHGYDAPIYTNVTYPITVNPPFVPTENPTGCYSLTFNVDESWLQEGQTRIIFDGVNSAFH
LWCNGRWVGYGQDSRLPSEFDLSAFLRAGENRLAVMVLRWSDGSYLEDQDMWRMSGIFRDVSLLHKPTTQISDFHV
ATRFNDDFSRAVLEAEVQMCGELRDYLRVTVSLWQGETQVASGTAPFGGEIIDERGGYADRVTLRLNVENPKLWSA
EIPNLYRAVVELHTADGTLIEAEACDVGFREVRIENGLLLLNGKPLLIRGVNRHEHHPLHGQVMDEQTMVQDILLM
KQNNFNAVRCSHYPNHPLWYTLCDRYGLYVVDEANIETHGMVPMNRLTDDPRWLPAMSERVTRMVQRDRNHPSVII
WSLGNESGHGANHDALYRWIKSVDPSRPVQYEGGGADTTATDIICPMYARVDEDQPFPAVPKWSIKKWLSLPGETR
PLILCEYAHAMGNSLGGFAKYWQAFRQYPRLQGGFVWDWVDQSLIKYDENGNPWSAYGGDFGDTPNDRQFCMNGLV
FADRTPHPALTEAKHQQQFFQFRLSGQTIEVTSEYLFRHSDNELLHWMVALDGKPLASGEVPLDVAPQGKQLIELP
ELPQPESAGQLWLTVRVVQPNATAWSEAGHISAWQQWRLAENLSVTLPAASHAIPHLTTSEMDFCIELGNKRWQFN
RQSGFLSQMWIGDKKQLLTPLRDQFTRAPLDNDIGVSEATRIDPNAWVERWKAAGHYQAEAALLQCTADTLADAVL
ITTAHAWQHQGKTLFISRKTYRIDGSGQMAITVDVEVASDTPHPARIGLNCQLAQVAERVNWLGLGPQENYPDRLT
AACFDRWDLPLSDMYTPYVFPSENGLRCGTRELNYGPHQWRGDFQFNISRYSQQQLMETSHRHLLHAEEGTWLNID
GFHMGIGGDDSWSPSVSAEFQLSAGRYHYQLVWCQK"
     primer          1937..1972
                     /label="AMG15C (upstream_fwd)"
                     /note="sequence: cgggttgccgttttcatcatatttaatcagcgactg"
                     /ApEinfo_revcolor="#7bcdc8"
                     /ApEinfo_fwdcolor="#7bcdc8"
     primer          1951..1972
                     /label="AMG30C (seq)"
                     /note="sequence: catcatatttaatcagcgactg"
                     /ApEinfo_revcolor="#d59687"
                     /ApEinfo_fwdcolor="#d59687"
     misc_feature    1951..2457
                     /label="upstream homology"
                     /ApEinfo_revcolor="#85dae9"
                     /ApEinfo_fwdcolor="#85dae9"
     CDS             complement(1951..2457)
                     /label="lacZ CDS"
                     /ApEinfo_revcolor="#84b0dc"
```

```
                     /ApEinfo_fwdcolor="#84b0dc"
                     /note="locus_tag: b0344 gene_synonym: ECK0341
EC_number: 3.2.1.108 transl_table: 11 product: beta-galactosidase
protein_id: NP_414878.1 db_xref: GeneID:945006 translation:
MTMITDSLAVVLQRRDWENPGVTQLNRLAAHPPFASWRNSEEARTDRPSQQLRSLNGEWRFAWFPAPEAVPESWLE
CDLPEADTVVVPSNWQMHGYDAPIYTNVTYPITVNPPFVPTENPTGCYSLTFNVDESWLQEGQTRIIFDGVNSAFH
LWCNGRWVGYGQDSRLPSEFDLSAFLRAGENRLAVMVLRWSDGSYLEDQDMWRMSGIFRDVSLLHKPTTQISDFHV
ATRFNDDFSRAVLEAEVQMCGELRDYLRVTVSLWQGETQVASGTAPFGGEIIDERGGYADRVTLRLNVENPKLWSA
EIPNLYRAVVELHTADGTLIEAEACDVGFREVRIENGLLLLNGKPLLIRGVNRHEHHPLHGQVMDEQTMVQDILLM
KQNNFNAVRCSHYPNHPLWYTLCDRYGLYVVDEANIETHGMVPMNRLTDDPRWLPAMSERVTRMVQRDRNHPSVII
WSLGNESGHGANHDALYRWIKSVDPSRPVQYEGGGADTTATDIICPMYARVDEDQPFPAVPKWSIKKWLSLPGETR
PLILCEYAHAMGNSLGGFAKYWQAFRQYPRLQGGFVWDWVDQSLIKYDENGNPWSAYGGDFGDTPNDRQFCMNGLV
FADRTPHPALTEAKHQQQFFQFRLSGQTIEVTSEYLFRHSDNELLHWMVALDGKPLASGEVPLDVAPQGKQLIELP
ELPQPESAGQLWLTVRVVQPNATAWSEAGHISAWQQWRLAENLSVTLPAASHAIPHLTTSEMDFCIELGNKRWQFN
RQSGFLSQMWIGDKKQLLTPLRDQFTRAPLDNDIGVSEATRIDPNAWVERWKAAGHYQAEAALLQCTADTLADAVL
ITTAHAWQHQGKTLFISRKTYRIDGSGQMAITVDVEVASDTPHPARIGLNCQLAQVAERVNWLGLGPQENYPDRLT
AACFDRWDLPLSDMYTPYVFPSENGLRCGTRELNYGPHQWRGDFQFNISRYSQQQLMETSHRHLLHAEEGTWLNID
GFHMGIGGDDSWSPSVSAEFQLSAGRYHYQLVWCQK"
     primer          2433..2480
                     /label="MAS46B"
                     /note="sequence:
ccgtgggtttcaatattggcttcatccaattattgaaggccgctaacg"
                     /ApEinfo_revcolor="#faac61"
                     /ApEinfo_fwdcolor="#faac61"
     primer          complement(2437..2481)
                     /label="MAS45B"
                     /note="sequence:
gcgttagcggccttcaataattggatgaagccaatattgaaaccc"
                     /ApEinfo_revcolor="#b1ff67"
                     /ApEinfo_fwdcolor="#b1ff67"
     terminator      2458..2506
                     /label="tVoigtS5, L3S3P22 47C>G"
                     /ApEinfo_revcolor="#c6c9d1"
                     /ApEinfo_fwdcolor="#c6c9d1"
     misc_feature    2458..3639
                     /label="RAM CymR from pDSZ114"
                     /ApEinfo_revcolor="#ff9ccd"
                     /ApEinfo_fwdcolor="#ff9ccd"
     misc_feature    2482..2506
                     /label="homology tVoigtS4"
                     /ApEinfo_revcolor="#f58a5e"
                     /ApEinfo_fwdcolor="#f58a5e"
     primer          2482..2506
                     /label="AB17 ConL F seq"
                     /note="sequence: ggcctttttttgtttctggtctgcc"
                     /ApEinfo_revcolor="#75c6a9"
                     /ApEinfo_fwdcolor="#75c6a9"
     misc_feature    2512..2519
                     /label="SbfI"
                     /ApEinfo_revcolor="#85dae9"
                     /ApEinfo_fwdcolor="#85dae9"
     misc_feature    2524..2555
                     /label="CymR operator"
                     /ApEinfo_revcolor="#ffef86"
                     /ApEinfo_fwdcolor="#ffef86"
     promoter        2524..2613
                     /label="P.CymRC (Marionette)"
```

```
                     /ApEinfo_revcolor="#ffef86"
                     /ApEinfo_fwdcolor="#ffef86"
     misc_feature    2547..2552
                     /label=-35
                     /ApEinfo_revcolor="#ffef86"
                     /ApEinfo_fwdcolor="#ffef86"
     misc_feature    2570..2575
                     /label=-10
                     /ApEinfo_revcolor="#ffef86"
                     /ApEinfo_fwdcolor="#ffef86"
     misc_feature    2582..2582
                     /label="TSS"
                     /ApEinfo_revcolor="#ffef86"
                     /ApEinfo_fwdcolor="#ffef86"
     misc_feature    2582..2613
                     /label="CymR operator"
                     /ApEinfo_revcolor="#ffef86"
                     /ApEinfo_fwdcolor="#ffef86"
     misc_feature    2618..2673
                     /label="U64 RNA Guide"
                     /ApEinfo_revcolor="#faac61"
                     /ApEinfo_fwdcolor="#faac61"
     misc_feature    2618..3578
                     /label="U64 Ribozyme transcript"
                     /ApEinfo_revcolor="#d59687"
                     /ApEinfo_fwdcolor="#d59687"
                     /note="https://rnacentral.org/rna/URS0002349E27/5911"
     misc_feature    2668..2673
                     /label="IGS"
                     /ApEinfo_revcolor="#c7b0e3"
                     /ApEinfo_fwdcolor="#c7b0e3"
     misc_feature    2674..3060
                     /label="Group I Intron Ribozyme (from p-OiRS3GG)"
                     /ApEinfo_revcolor="#b4abac"
                     /ApEinfo_fwdcolor="#b4abac"
     misc_feature    3061..3081
                     /label="Probe binding site"
                     /ApEinfo_revcolor="#ff9ccd"
                     /ApEinfo_fwdcolor="#ff9ccd"
     primer          complement(3061..3082)
                     /label="KRG089"
                     /note="sequence: GCGGTCTCAgcgcCGGGAAAAGCATTGAACACCAT"
                     /ApEinfo_revcolor="#b7e6d7"
                     /ApEinfo_fwdcolor="#b7e6d7"
     primer          complement(3061..3082)
                     /label="oMJD105"
                     /note="sequence: GCGGTCTCAcactcGGGAAAAGCATTGAACACCAT"
                     /ApEinfo_revcolor="#b7e6d7"
                     /ApEinfo_fwdcolor="#b7e6d7"
     misc_feature    3061..3578
                     /label="Barcode (sfGFP_2)"
                     /ApEinfo_revcolor="#84b0dc"
                     /ApEinfo_fwdcolor="#84b0dc"
     primer          3080..3101
                     /label="KRG090"
                     /note="sequence: GCGGTCTCAgcgcGGTTATCCGGATCACATGAAA"
                     /ApEinfo_revcolor="#b4abac"
```

```
                     /ApEinfo_fwdcolor="#b4abac"
     primer          3082..3101
                     /label="oMJD104"
                     /note="sequence: GCGGTCTCAGAGTGGTTATCCGGATCACATGAAA"
                     /ApEinfo_revcolor="#b4abac"
                     /ApEinfo_fwdcolor="#b4abac"
     misc_feature    3491..3496
                     /label="BamHI-BglII Scar"
                     /ApEinfo_revcolor="#ac1dff"
                     /ApEinfo_fwdcolor="#ac1dff"
     terminator      3583..3635
                     /label="tVoigtS4 L3S3P21 51C>G"
                     /ApEinfo_revcolor="#f58a5e"
                     /ApEinfo_fwdcolor="#f58a5e"
     misc_feature    3611..3635
                     /label="homology tVoigtS5"
                     /ApEinfo_revcolor="#f58a5e"
                     /ApEinfo_fwdcolor="#f58a5e"
     primer          3611..3635
                     /label="AB17 ConL F seq"
                     /note="sequence: ggccttttttgtttctggtctgcc"
                     /ApEinfo_revcolor="#75c6a9"
                     /ApEinfo_fwdcolor="#75c6a9"
     primer          3620..3665
                     /label="MAS36A"
                     /note="sequence:
gccgccgggcgtttttatttaaagttgttctgcttcatcagcagg"
                     /ApEinfo_revcolor="#75c6a9"
                     /ApEinfo_fwdcolor="#75c6a9"
     primer          complement(3622..3639)
                     /label="MAS49B"
                     /note="sequence: CAGCggCagaccagaaac"
                     /ApEinfo_revcolor="#faac61"
                     /ApEinfo_fwdcolor="#faac61"
     primer          complement(3622..3655)
                     /label="MAS47B"
                     /note="sequence: gaacctcttacgtgcccCAGCggCagaccagaaac"
                     /ApEinfo_revcolor="#faac61"
                     /ApEinfo_fwdcolor="#faac61"
     primer          3640..3665
                     /label="MAS48B"
                     /note="sequence: taaagttgttctgcttcatcagcagg"
                     /ApEinfo_revcolor="#f8d3a9"
                     /ApEinfo_fwdcolor="#f8d3a9"
     misc_feature    3640..4147
                     /label="downstream homology"
                     /ApEinfo_revcolor="#85dae9"
                     /ApEinfo_fwdcolor="#85dae9"
     CDS             complement(3640..4147)
                     /label="lacZ CDS"
                     /ApEinfo_revcolor="#84b0dc"
                     /ApEinfo_fwdcolor="#84b0dc"
                     /note="locus_tag: b0344 gene_synonym: ECK0341
EC_number: 3.2.1.108 transl_table: 11 product: beta-galactosidase
protein_id: NP_414878.1 db_xref: GeneID:945006 translation:
MTMITDSLAVVLQRRDWENPGVTQLNRLAAHPPFASWRNSEEARTDRPSQQLRSLNGEWRFAWFPAPEAVPESWLE
CDLPEADTVVVPSNWQMHGYDAPIYTNVTYPITVNPPFVPTENPTGCYSLTFNVDESWLQEGQTRIIFDGVNSAFH
```

```
LWCNGRWVGYGQDSRLPSEFDLSAFLRAGENRLAVMVLRWSDGSYLEDQDMWRMSGIFRDVSLLHKPTTQISDFHV
ATRFNDDFSRAVLEAEVQMCGELRDYLRVTVSLWQGETQVASGTAPFGGEIIDERGGYADRVTLRLNVENPKLWSA
EIPNLYRAVVELHTADGTLIEAEACDVGFREVRIENGLLLLNGKPLLIRGVNRHEHHPLHGQVMDEQTMVQDILLM
KQNNFNAVRCSHYPNHPLWYTLCDRYGLYVVDEANIETHGMVPMNRLTDDPRWLPAMSERVTRMVQRDRNHPSVII
WSLGNESGHGANHDALYRWIKSVDPSRPVQYEGGGADTTATDIICPMYARVDEDQPFPAVPKWSIKKWLSLPGETR
PLILCEYAHAMGNSLGGFAKYWQAFRQYPRLQGGFVWDWVDQSLIKYDENGNPWSAYGGDFGDTPNDRQFCMNGLV
FADRTPHPALTEAKHQQQFFQFRLSGQTIEVTSEYLFRHSDNELLHWMVALDGKPLASGEVPLDVAPQGKQLIELP
ELPQPESAGQLWLTVRVVQPNATAWSEAGHISAWQQWRLAENLSVTLPAASHAIPHLTTSEMDFCIELGNKRWQFN
RQSGFLSQMWIGDKKQLLTPLRDQFTRAPLDNDIGVSEATRIDPNAWVERWKAAGHYQAEAALLQCTADTLADAVL
ITTAHAWQHQGKTLFISRKTYRIDGSGQMAITVDVEVASDTPHPARIGLNCQLAQVAERVNWLGLGPQENYPDRLT
AACFDRWDLPLSDMYTPYVFPSENGLRCGTRELNYGPHQWRGDFQFNISRYSQQQLMETSHRHLLHAEEGTWLNID
GFHMGIGGDDSWSPSVSAEFQLSAGRYHYQLVWCQK"
     primer          complement(4127..4147)
                     /label="AMG31C (seq)"
                     /note="sequence: cataaaccgactacacaaatc"
                     /ApEinfo_revcolor="#c7b0e3"
                     /ApEinfo_fwdcolor="#c7b0e3"
     primer          complement(4127..4160)
                     /label="AMG22C (downstream_rev)"
                     /note="sequence: cgtctcgttgctgcataaaccgactacacaaatc"
                     /ApEinfo_revcolor="#7bcdc8"
                     /ApEinfo_fwdcolor="#7bcdc8"
     primer          4141..4180
                     /label="AMG33C"
                     /note="sequence:
gtttatgcagcaacgagacggttttatctgttgtttgtcg"
                     /ApEinfo_revcolor="#c7b0e3"
                     /ApEinfo_fwdcolor="#c7b0e3"
     CDS             complement(4148..4160)
                     /label="lacZ CDS"
                     /ApEinfo_revcolor="#84b0dc"
                     /ApEinfo_fwdcolor="#84b0dc"
                     /note="locus_tag: b0344 gene_synonym: ECK0341
EC_number: 3.2.1.108 transl_table: 11 product: beta-galactosidase
protein_id: NP_414878.1 db_xref: GeneID:945006 translation:
MTMITDSLAVVLQRRDWENPGVTQLNRLAAHPPFASWRNSEEARTDRPSQQLRSLNGEWRFAWFPAPEAVPESWLE
CDLPEADTVVVPSNWQMHGYDAPIYTNVTYPITVNPPFVPTENPTGCYSLTFNVDESWLQEGQTRIIFDGVNSAFH
LWCNGRWVGYGQDSRLPSEFDLSAFLRAGENRLAVMVLRWSDGSYLEDQDMWRMSGIFRDVSLLHKPTTQISDFHV
ATRFNDDFSRAVLEAEVQMCGELRDYLRVTVSLWQGETQVASGTAPFGGEIIDERGGYADRVTLRLNVENPKLWSA
EIPNLYRAVVELHTADGTLIEAEACDVGFREVRIENGLLLLNGKPLLIRGVNRHEHHPLHGQVMDEQTMVQDILLM
KQNNFNAVRCSHYPNHPLWYTLCDRYGLYVVDEANIETHGMVPMNRLTDDPRWLPAMSERVTRMVQRDRNHPSVII
WSLGNESGHGANHDALYRWIKSVDPSRPVQYEGGGADTTATDIICPMYARVDEDQPFPAVPKWSIKKWLSLPGETR
PLILCEYAHAMGNSLGGFAKYWQAFRQYPRLQGGFVWDWVDQSLIKYDENGNPWSAYGGDFGDTPNDRQFCMNGLV
FADRTPHPALTEAKHQQQFFQFRLSGQTIEVTSEYLFRHSDNELLHWMVALDGKPLASGEVPLDVAPQGKQLIELP
ELPQPESAGQLWLTVRVVQPNATAWSEAGHISAWQQWRLAENLSVTLPAASHAIPHLTTSEMDFCIELGNKRWQFN
RQSGFLSQMWIGDKKQLLTPLRDQFTRAPLDNDIGVSEATRIDPNAWVERWKAAGHYQAEAALLQCTADTLADAVL
ITTAHAWQHQGKTLFISRKTYRIDGSGQMAITVDVEVASDTPHPARIGLNCQLAQVAERVNWLGLGPQENYPDRLT
AACFDRWDLPLSDMYTPYVFPSENGLRCGTRELNYGPHQWRGDFQFNISRYSQQQLMETSHRHLLHAEEGTWLNID
GFHMGIGGDDSWSPSVSAEFQLSAGRYHYQLVWCQK"
     primer          complement(4356..1)
                     /label="JEC14A | JC.A14.Rev_seq"
                     /note="sequence: ctttttacggttcctggc"
                     /ApEinfo_revcolor="#d59687"
                     /ApEinfo_fwdcolor="#d59687"
ORIGIN
        1 ggccgcgttg ctggcgtttt tccacaggct ccgcccccct gacgagcatc acaaaaatcg
       61 acgctcaagt cagaggtggc gaaacccgac aggactataa agataccagg cgtttccccc
```

```
 121 tggaagctcc ctcgtgcgct ctcctgttcc gaccctgccg cttaccggat acctgtccgc
 181 ctttctccct tcgggaagcg tggcgctttc tcatagctca cgctgtaggt atctcagttc
 241 ggtgtaggtc gttcgctcca agctgggctg tgtgcacgaa ccccccgttc agcccgaccg
 301 ctgcgcctta tccggtaact atcgtcttga gtccaacccg gtaagacacg acttatcgcc
 361 actggcagca gccactggta acaggattag cagagcgagg tatgtaggcg gtgctacaga
 421 gttcttgaag tggtggccta actacggcta cactagaaga acagtatttg gtatctgcgc
 481 tctgctgaag ccagttacct tcggaaaaag agttggtagc tcttgatccg gcaaacaaac
 541 caccgctggt agcggtggtt tttttgtttg caagcagcag attacgcgca gaaaaaaagg
 601 atctcaagaa gatcctttga tcttttctac ggggtctgac gctcagtgga acgaaaactc
 661 acgttaaggg attttggtca tgagattatc aaaaaggatc ttcacctaga tccttttaaa
 721 ttaaaaatga agttttaaat caatctaaag tatatatgag taaacttggt ctgacagtta
 781 ccaatgctta atcagtgagg cacctatctc agcgatctgt ctatttcgtt catccatagt
 841 tgcctgactc cccgtcgtgt agataactac gatacgggag ggcttaccat ctggccccag
 901 tgctgcaatg ataccgcgag acccacgctc accggctcca gatttatcag caataaacca
 961 gccagccgga agggccgagc gcagaagtgg tcctgcaact ttatccgcct ccatccagtc
1021 tattaattgt tgccgggaag ctagagtaag tagttcgcca gttaatagtt tgcgcaacgt
1081 tgttgccatt gctacaggca tcgtggtgtc acgctcgtcg tttggtatgg cttcattcag
1141 ctccggttcc caacgatcaa ggcgagttac atgatccccc atgttgtgca aaaaagcggt
1201 tagctccttc ggtcctccga tcgttgtcag aagtaagttg gccgcagtgt tatcactcat
1261 ggttatggca gcactgcata attctcttac tgtcatgcca tccgtaagat gcttttctgt
1321 gactggtgag tactcaacca agtcattctg agaatagtgt atgcggcgac cgagttgctc
1381 ttgcccggcg tcaatacggg ataataccgc gccacatagc agaactttaa aagtgctcat
1441 cattggaaaa cgttcttcgg ggcgaaaact ctcaaggatc ttaccgctgt tgagatccag
1501 ttcgatgtaa cccactcgtg cacccaactg atcttcagca tcttttactt tcaccagcgt
1561 ttctgggtga gcaaaaacag gaaggcaaaa tgccgcaaaa aagggaataa gggcgacacg
1621 gaaatgttga atactcatac tcttcctttt tcaatattat tgaagcattt atcagggtta
1681 ttgtctcatg agcggataca tatttgaatg tatttagaaa aataaacaaa taggggttcc
1741 gcgcacattt ccccgaaaag tgccacctga cgtctaagaa accattatta tcatgacatt
1801 aacctataaa aataggcgta tcacgaggca gaatttcaga taaaaaaaat ccttagcttt
1861 cgctaaggat gatttctgga attctaaaga tctttgacag ctagctcagt cctaggtata
1921 atactagtgg atctgacggg ttgccgtttt catcatattt aatcagcgac tgatccaccc
1981 agtcccagac gaagccgccc tgtaaacggg gatactgacg aaacgcctgc cagtatttag
2041 cgaaaccgcc aagactgtta cccatcgcgt gggcgtattc gcaaaggatc agcgggcgcg
2101 tctctccagg tagcgaaagc catttttga tggaccattt cggcacagcc gggaagggct
2161 ggtcttcatc cacgcgcgcg tacatcgggc aaataatatc ggtggccgtg gtgtcggctc
2221 cgccgccttc atactgcacc gggcgggaag gatcgacaga tttgatccag cgatacagcg
2281 cgtcgtgatt agcgccgtgg cctgattcat tccccagcga ccagatgatc acactcgggt
2341 gattacgatc gcgctgcacc attcgcgtta cgcgttcgct catcgccggt agccagcgcg
2401 gatcatcggt cagacgattc attggcacca tgccgtgggt ttcaatattg gcttcatcca
2461 attattgaag gccgctaacg cggccttttt ttgtttctgg tctgcctcct CcctgcaggC
2521 TTCaacaaac agacaatctg gtctgtttgt attatggaaa atttttctgt ataatagatt
2581 caacaaacag acaatctggt ctgtttgtat tatcgacCAA CCCACTCCCA TGGTGTGACG
2641 GGCGGTGTGT ACAAGGCCCG GGAACGTgTT CACAAAAGTT ATCAGGCATG CACCTGGTAG
2701 CTAGTCTTTA AACCAATAGA TTGCATCGGT TTAAAAGGCA AGACCGTCAA ATTGCGGGAA
2761 AGGGGTCAAC AGCCGTTCAG TACCAAGTCT CAGGGGAAAC TTTGAGATGG CCTTGCAAAG
2821 GGTATGGTAA TAAGCTGACG GACATGGTCC TAACCACGCA GCCAAGTCCT AAGTCAACAG
2881 ATCTTCTGTT GATATGGATG CAGTTCACAG ACTAAATGTC GGTCGGGGAA GATGTATTCT
2941 TCTCATAAGA TATAGTCGGA CCTCTCCTTA ATGGGAGCTA GCGGATGAAG TGATGCAACA
3001 CTGGAGCCGC TGGGAACTAA TTTGTATGCG AAAGTATATT GATTAGTTTT GGAGTACTCG
3061 ATGGTGTTCA ATGCTTTTCC CGTTATCCGG ATCACATGAA ACGGCATGAC TTTTTCAAGA
3121 GTGCCATGCC CGAAGGTTAT GTACAGGAAC GCACTATATC TTTCAAAGAT GACGGGACCT
3181 ACAAGACGCG TGCTGAAGTC AAGTTTGAAG GTGATACCCT TGTTAATCGT ATCGAGTTAA
3241 AGGGTATTGA TTTTAAAGAA GATGGAAACA TTCTTGGACA CAAACTCGAG TACAACTTTA
3301 ACTCACACAA TGTATACATC ACGGCAGACA AACAAAAGAA TGGAATCAAA GCTAACTTCA
3361 AAATTCGCCA CAACGTTGAA GATGGTTCCG TTCAACTAGC AGACCATTAT CAACAAAATA
3421 CTCCAATTGG CGATGGCCCT GTCCTTTTAC CAGACAACCA TTACCTGTCG ACACAATCTG
3481 TCCTTTCGAA AGATCCCAAC GAAAAGCGTG ACCACATGGT CCTTCTTGAG TTTGTAACTG
```

```
     3541 CTGCTGGGAT TACACATGGC ATGGATGAGC TCTACAAATG GCccaattat tgaaggcctc
     3601 cctaacgggg ggcctttttt tgtttctggt ctGccGCTGt aaagttgttc tgcttcatca
     3661 gcaggatatc ctgcaccatc gtctgctcat ccatgacctg accatgcaga ggatgatgct
     3721 cgtgacggtt aacgcctcga atcagcaacg gcttgccgtt cagcagcagc agaccatttt
     3781 caatccgcac ctcgcggaaa ccgacatcgc aggcttctgc ttcaatcagc gtgccgtcgg
     3841 cggtgtgcag ttcaaccacc gcacgataga gattcgggat ttcggcgctc cacagtttcg
     3901 ggttttcgac gttcagacgt agtgtgacgc gatcggcata accaccacgc tcatcgataa
     3961 tttcaccgcc gaaaggcgcg gtgccgctgg cgacctgcgt ttcaccctgc cataaagaaa
     4021 ctgttacccg taggtagtca cgcaactcgc cgcacatctg aacttcagcc tccagtacag
     4081 cgcggctgaa atcatcatta aagcgagtgg caacatggaa atcgctgatt tgtgtagtcg
     4141 gtttatgcag caacgagacg gttttatctg ttgtttgtcg gtgaactgga tccttactcg
     4201 agtctagact gcaggcttcc tcgctcactg actcgctgcg ctcggtcgtt cggctgcggc
     4261 gagcggtatc agctcactca aaggcggtaa tacggttatc cacagaatca ggggataacg
     4321 caggaaagaa catgtgagca aaaggccagc aaaaggccag gaaccgtaaa aa
//
```
